# The complete sequence of a human Y chromosome

**DOI:** 10.1101/2022.12.01.518724

**Authors:** Arang Rhie, Sergey Nurk, Monika Cechova, Savannah J. Hoyt, Dylan J. Taylor, Nicolas Altemose, Paul W. Hook, Sergey Koren, Mikko Rautiainen, Ivan A. Alexandrov, Jamie Allen, Mobin Asri, Andrey V. Bzikadze, Nae-Chyun Chen, Chen-Shan Chin, Mark Diekhans, Paul Flicek, Giulio Formenti, Arkarachai Fungtammasan, Carlos Garcia Giron, Erik Garrison, Ariel Gershman, Jennifer L. Gerton, Patrick G.S. Grady, Andrea Guarracino, Leanne Haggerty, Reza Halabian, Nancy F. Hansen, Robert Harris, Gabrielle A. Hartley, William T. Harvey, Marina Haukness, Jakob Heinz, Thibaut Hourlier, Robert M. Hubley, Sarah E. Hunt, Stephen Hwang, Miten Jain, Rupesh K. Kesharwani, Alexandra P. Lewis, Heng Li, Glennis A. Logsdon, Julian K. Lucas, Wojciech Makalowski, Christopher Markovic, Fergal J. Martin, Ann M. Mc Cartney, Rajiv C. McCoy, Jennifer McDaniel, Brandy M. McNulty, Paul Medvedev, Alla Mikheenko, Katherine M. Munson, Terence D. Murphy, Hugh E. Olsen, Nathan D. Olson, Luis F. Paulin, David Porubsky, Tamara Potapova, Fedor Ryabov, Steven L. Salzberg, Michael E.G. Sauria, Fritz J. Sedlazeck, Kishwar Shafin, Valery A. Shepelev, Alaina Shumate, Jessica M. Storer, Likhitha Surapaneni, Angela M. Taravella Oill, Françoise Thibaud-Nissen, Winston Timp, Marta Tomaszkiewicz, Mitchell R. Vollger, Brian P. Walenz, Allison C. Watwood, Matthias H. Weissensteiner, Aaron M. Wenger, Melissa A. Wilson, Samantha Zarate, Yiming Zhu, Justin M. Zook, Evan E. Eichler, Rachel J. O’Neill, Michael C. Schatz, Karen H. Miga, Kateryna D. Makova, Adam M. Phillippy

**Author notes:** These authors contributed equally (Arang Rhie, Sergey Nurk, Monika Cechova, Savannah J. Hoyt, Dylan J. Taylor). Department of Anatomy and Anthropology and Department of Human Molecular Genetics and Biochemistry, Sackler Faculty of Medicine, Tel Aviv University, Israel (Ivan A. Alexandrov).

## Abstract

The human Y chromosome has been notoriously difficult to sequence and assemble because of its complex repeat structure including long palindromes, tandem repeats, and segmental duplications^1–3^. As a result, more than half of the Y chromosome is missing from the GRCh38 reference sequence and it remains the last human chromosome to be finished^4, 5^. Here, the Telomere-to-Telomere (T2T) consortium presents the complete 62,460,029 base pair sequence of a human Y chromosome from the HG002 genome (T2T-Y) that corrects multiple errors in GRCh38-Y and adds over 30 million base pairs of sequence to the reference, revealing the complete ampliconic structures of *TSPY*, *DAZ*, and *RBMY* gene families; 41 additional protein-coding genes, mostly from the *TSPY* family; and an alternating pattern of human satellite 1 and 3 blocks in the heterochromatic Yq12 region. We have combined T2T-Y with a prior assembly of the CHM13 genome^4^ and mapped available population variation, clinical variants, and functional genomics data to produce a complete and comprehensive reference sequence for all 24 human chromosomes.

---

The human Y chromosome plays critical roles in fertility and hosts genes important for spermatogenesis, as well as *SRY*, the mammalian sex-determining locus^6^. However, in the human reference genome, GRCh38, the Y chromosome remains the most incomplete chromosome, with >50% of bases represented by gaps. These multi-megabase gaps have persisted for decades and represent sequences flanking the endogenous model centromere, parts of the ampliconic regions, and large heterochromatic regions. The architecture of the Y chromosome, specifically the presence of large tandemly arrayed and inverted repeats (i.e. palindromes)^1^, makes assembly difficult and hinders the study of rearrangements, inversions, duplications, and deletions in several critical regions such as AZFa, AZFb, and AZFc (azoospermia factor), which are linked to clinical phenotypes, including infertility^7^.

Following the first complete assemblies of chromosomes X^8^ and 8^9^, the Telomere-to-Telomere (T2T) consortium successfully assembled all chromosomes of the CHM13 cell line^4^. This first complete human genome assembly was enabled by innovative technological improvements in generating Pacific Biosciences (PacBio) high-fidelity reads (HiFi)^10^ and Oxford Nanopore ultra-long reads (ONT)^11^, the development of better assembly algorithms for utilizing HiFi reads and generating assembly graphs^12^, the use of ONT reads for better resolving the graph^13^, new methods for validating and polishing^14–18^, and a coordinated curation effort to finish the assembly. Having been derived from a complete hydatidiform mole, CHM13 has a 46,XX karyotype but is almost entirely homozygous. This simplified assembly of its genome, but prevented assembly of a Y chromosome.

In parallel, with the goal of including broader genomic diversity across populations^19^, the Human Pangenome Reference Consortium (HPRC) has evaluated various methods for generating high-quality diploid genome assemblies^20^ using a well characterized human genome, HG002, which has been previously assembled^21^ and is commonly used for benchmarking by the Genome in a Bottle consortium^22^. Using this rich set of data, and integrating the lessons learned from assembling CHM13, we successfully reconstructed the complete sequence of the HG002 Y chromosome, hereafter referred to as T2T-Y.

Here we analyze the composition of the newly assembled pseudoautosomal regions (PARs), ampliconic and palindromic sequences, centromeric satellites, and q-arm heterochromatin of a complete Y chromosome. We have annotated T2T-Y and combined it with the prior T2T-CHM13 assembly to form a new, complete reference for all human chromosomes, T2T-CHM13+Y. To enable the use of this new reference sequence, we have lifted over available variation datasets from ClinVar^23^, GWAS^24^, dbSNP^25^ and gnomAD^26^. In addition, we have recalled variants from 1000 Genomes Project (1KGP)^27^ and Simons Genome Diversity Panel (SGDP)^28^ data, as well as epigenetic profiles from ENCODE data^29^. These experiments demonstrate improved mappability and variant calling for XY individuals when using T2T-Y as a reference.

## Assembly and validation of T2T-Y

Assembly of the HG002 Y chromosome followed the strategy used for the T2T-CHM13 genome^4^ (**Supplementary Table 1** and **Supplementary Fig. 1**). We used PacBio HiFi reads (60× haploid genome coverage) and ONT ultra-long reads (90× in reads longer than 100 kb) generated from HG002. An assembly string graph was first constructed for the whole HG002 genome using PacBio HiFi reads. The ChrX and ChrY string graph components shared connections to one another at the PARs, but to no other chromosomes in the genome and could be independently analyzed (**Extended Data Fig. 1a**). The remaining tangles in these XY subgraphs were resolved using ONT reads (**Extended Data Fig. 1b**). ChrX and ChrY chromosomal walks were identified using haplotype-specific k-mers from parental Illumina reads (**Extended Data Fig. 1c**), and a consensus sequence was computed for each. PAR1 was enriched for GA-microsatellites, which reduced HiFi coverage in this region and led to a more fragmented graph (due to a known HiFi sequencing bias^12^). These gaps were manually patched using a *de novo* assembly of trio-binned parental ONT reads^14^.

**Fig. 1.**
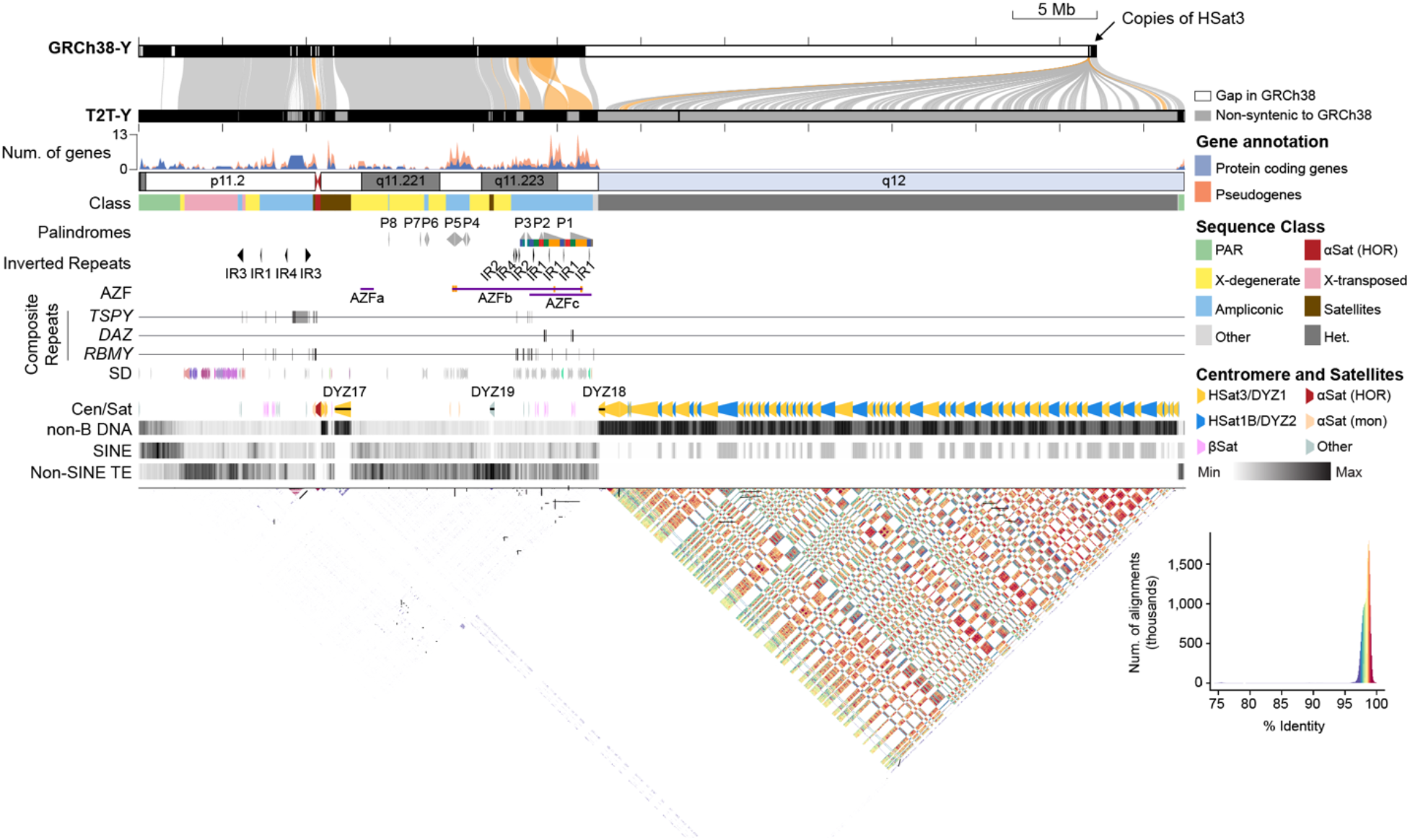
The structure of a complete Y chromosome. From top to bottom: Alignment of GRCh38-Y and T2T-Y. Regions with sequence identity over 95% are connected and colored by alignment direction (gray, forward; orange, reverse). Gene density plot shows enriched protein coding genes in ampliconic sequences. Sequence class, palindromes, inverted repeats (IR), and Azoospermia factor (AZF) a-c are annotated. Composite repeat arrays are named after the contained ampliconic genes. Segmental duplications (SDs) are colored by duplication types defined in DupMasker^35^. Centromere and satellite annotations (Cen/Sat) highlight the alternating HSat1 and HSat3 pattern comprising Yq12. Non-B DNA track shows regions forming alternate sequence structures are enriched in centromeric and satellite repeats. Short-interspersed repeat elements (SINE), including *AluY*, are highly enriched in the pseudo autosomal region 1 (PAR1). All other non-SINE transposable elements (TEs) are only found in the euchromatin. All repeats within T2T-Y are visualized by StainedGlass^36^ with similar repeats colored by % identity in the style of an alignment dotplot.

The ChrY draft assembly was further polished and validated using sequencing reads from Illumina (66× haploid genome coverage), HiFi (84×), and ONT (250×). During four rounds of polishing, 1,520 small and 10 large (>50 base) errors were detected and corrected (**Extended Data Fig. 2a**). Conservatively filtered long-read alignments identified two potential assembly issues remaining in the satellite (HSat) arrays around positions 40 Mb and 59.1 Mb, and Strand-seq^30, 31^ identified one inversion error within palindromic sequence P5 around position 18.8 Mb (**Extended Data Fig. 2b-c, Supplementary Table 2, Supplementary Figs. 2-4**). The validation signal at the two HSat positions was ambiguous, and the P5 inversion appears as a true recurrent inversion^32^, so these regions were noted but left uncorrected in this release. The remaining sequences showed no signs of collapse or false duplication, with even HiFi coverage (mean 39.3x ± SD 12.5 on ChrXY) except for regions associated with known sequencing biases^17^, all of which had supporting ONT coverage (reads over 25 kb, mean 78.1x ± SD 13.6 on ChrXY). The base error is estimated as less than 1 error per 10 Mb (Phred Q73.8, **Supplementary Table 3**). Mapped HiFi and ONT reads from the paternal HG003 genome are also consistent with the HG002 T2T-Y assembly, suggesting no large, structural variants were introduced during cell line immortalization and culture (**Supplementary Fig 5**).

**Fig. 2.**
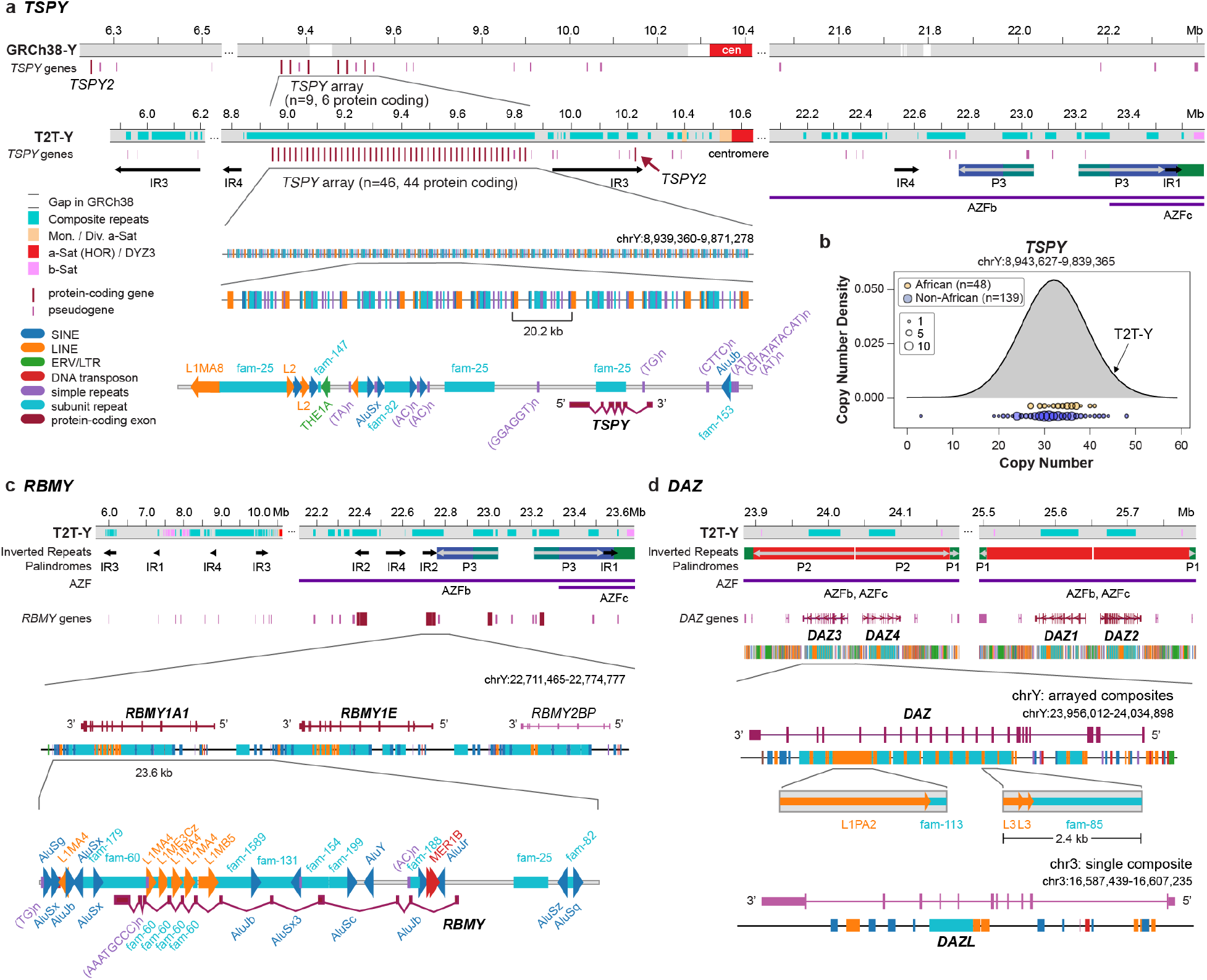
Ampliconic genes forming composite repeats. **a.** T2T-Y has 44 *TSPY* protein-coding genes organized in a single continuous array and a single *TSPY2* copy, compared to GRCh38-Y which has a gap in the *TSPY* array. T2T-Y shows a more regularized array and recovers additional *TSPY* pseudogenes not present in GRCh38-Y. **b**. Copy number differences of the *TSPY* protein-coding copies found in the SGDP. **c**, Repeat composition of the *RBMY* gene family. **d**. Repeat composition of the *DAZ* gene family, with one extra copy annotated on Chr3 that is missing L1PA2. While *TSPY* and *RBMY* genes are found within repeat composites forming arrays, *DAZ*-associated composites are embedded within the introns of the gene.

**Fig. 3.**
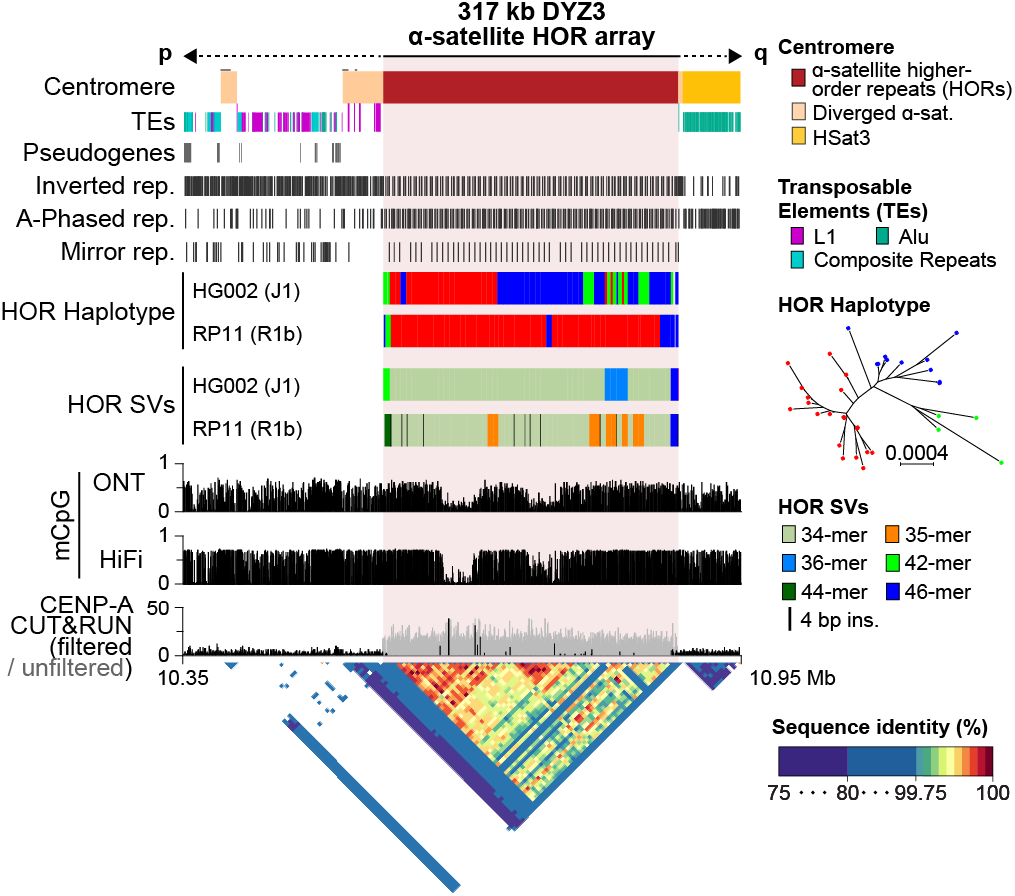
The structure of the T2T-Y centromere. No TEs were found within the DYZ3 array, while L1s (proximal) and *Alu*s (distal) were found within the diverged alpha satellites (drawn taller than the other TEs). A periodic non-B DNA motif pattern is shown within the HOR array. The HG002 (T2T-Y) HOR haplotypes and SVs reveal a different long-range structure and organization compared to a previously assembled centromere from RP11^47^. Three major HOR haplotypes were identified in HG002-Y based on their phylogenetic distance (red, blue, and green). RP11-Y has no 36-mer variants, but does have a number of 35-mers containing internal duplications. Histograms show the fraction of methylated CpG sites called by both ONT and HiFi, with two hypo-methylated centromeric dip regions (CDR) supported by CENP-A binding signal from CUT&RUN^50^. A StainedGlass dotplot illustrates high similarity within the HOR array (99.5– 100%).

**Fig. 4.**
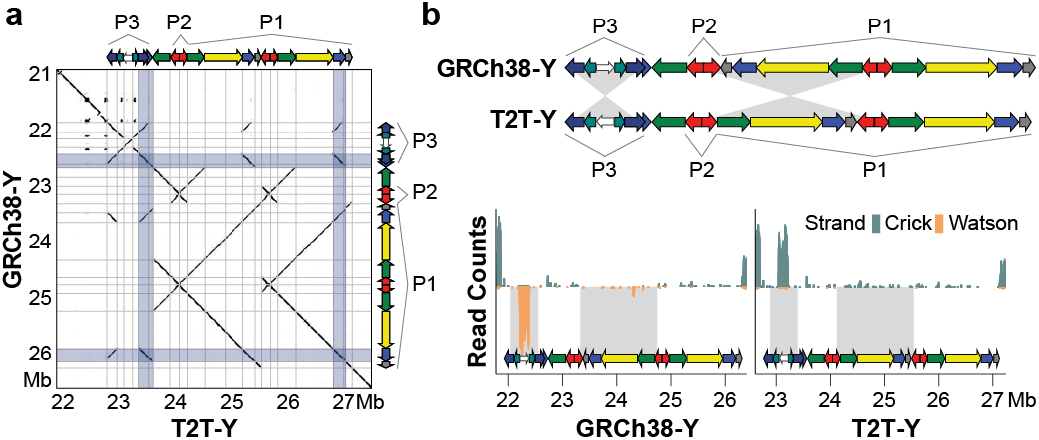
Comparison of the palindromic structure of the P1–P3 region. **a**. GRCh38-Y and T2T-Y alignment dotplot and schematics of the palindromes. Frequently recombining inverted repeats (IRs) in Azoospermia factor c (AZFc) region are highlighted in light blue. Deletion of AZFc between the IRs are known to cause spermatogenic failure^52^. A self-dotplot of the T2T-Y with AZFb and AZFc annotation is available as **Supplementary Fig. 15**. **b**. Top, a schematic of the palindromes. Two inversions are found, one in P3 and one between P1-P2. Below, Strand-seq signal from HG002 confirms the inverted orientation of P3 and P1 in T2T-Y compared to GRCh38-Y.

The resulting T2T-Y assembly is 62,460,029 bases in length with no gaps or model sequences, revealing the previously uncharacterized ∼30 Mb of sequence within the heterochromatic region of the q-arm (**Table 1**). In comparison, ChrY in the human reference genome (GRCh38-Y) consists of two sequences, with the longer sequence totaling 57.2 Mb (NC_000024.10), for which 53.8% (30.8 Mb) of the bases are unresolved gaps. The shorter GRCh38-Y sequence (NT_187395.1) is 37.2 kb in length, not placed in the primary Y assembly, and has been omitted from most prior genomic studies. The PAR1 (2.77 Mb) and PAR2 (329.5 kb) sequences in GRCh38-Y are duplicated from ChrX rather than assembled *de novo*, and the centromere is represented by a 227 kb model sequence. Direct sequence comparison between T2T-Y and GRCh38-Y yields an average sequence identity of ∼99.8% in the alignable regions, but with multiple structural differences including an incorrectly oriented centromere model for GRCh38-Y (**Fig. 1** and **Extended Data Fig. 3**). We identified the Y-chromosome haplogroup of HG002 as J-L816 (J1) and that of GRCh38 as R-L20 (R1b). These haplogroups are most commonly found among Ashkenazi Jews^33^ and Europeans^34^, respectively, consistent with the established ancestry of these genomes. T2T-Y was combined with the T2T-CHM13v1.1 assembly to create a new Y-bearing reference, T2T-CHM13v2.0, referred to here as T2T-CHM13+Y.

**Table 1.**
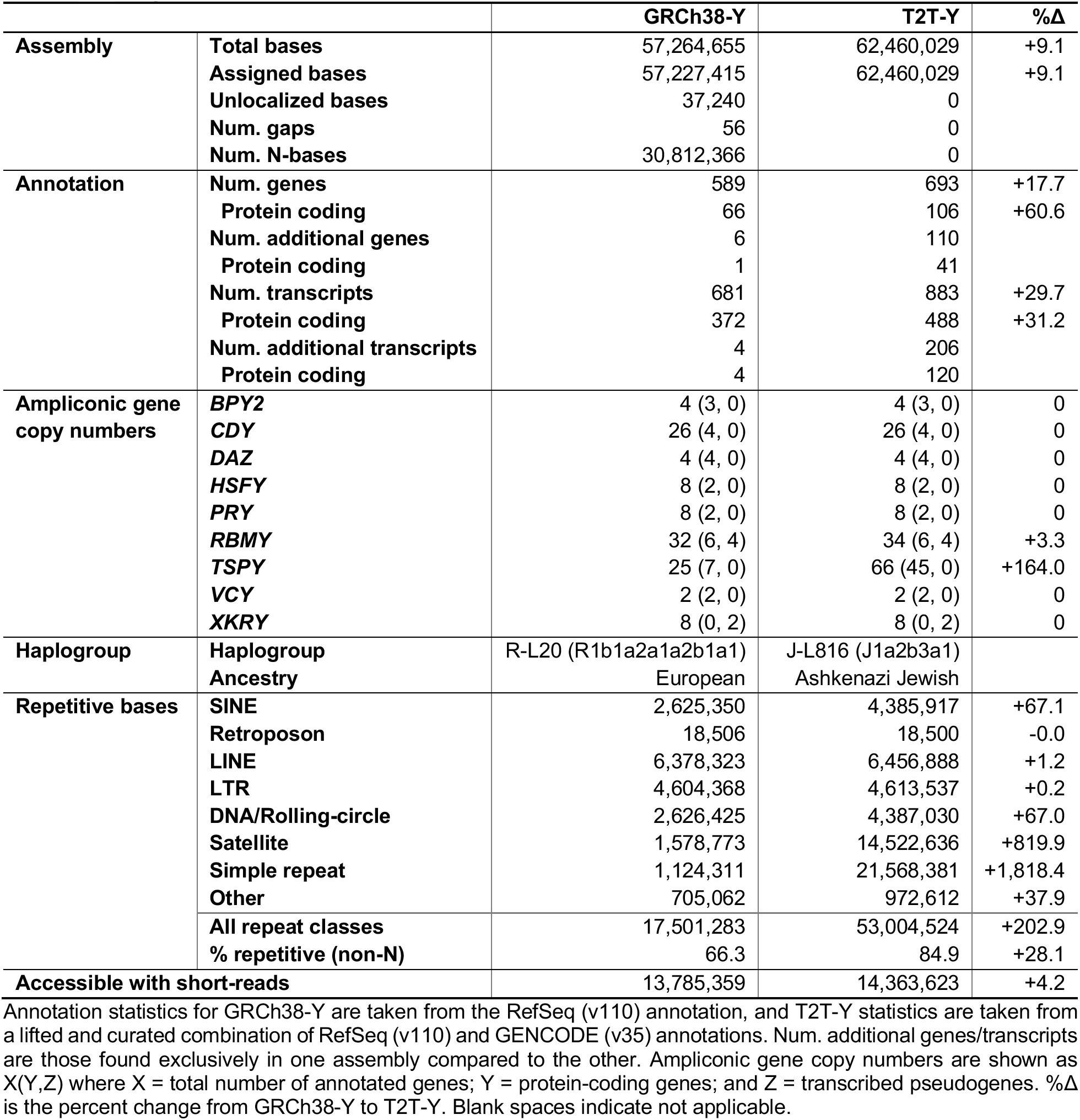
Comparison of GRCh38-Y and T2T-Y.

## Comprehensive annotation of the Y

### Gene annotation

We annotated T2T-CHM13+Y by mapping RefSeq (v110) and GENCODE (v35) annotations from GRCh38 and performed hand-curation of the ampliconic gene arrays (**Fig. 1** and **Supplementary Table 4-5**). NCBI RefSeq and EBI Ensembl generated additional *de novo* annotations using HG002 Iso-Seq transcriptomes from B-Lymphocyte and induced pluripotent stem cell (iPS) lines as well as tissue-specific expression data from other publicly available sources (**Supplementary Table 1, Supplementary Fig. 6-7**).

Our annotation of T2T-Y totals 693 genes and 883 transcripts, of which 106 genes (488 transcripts) are predicted to be protein-coding (**Table 1** and **Supplementary Table 4**). In addition to containing all genes annotated in GRCh38-Y, T2T-Y contains an additional 110 genes, among which 41 are predicted to be protein coding. The majority of these protein-coding genes (38 of 41) are additional copies of *TSPY*, one of the nine ampliconic gene families, filling the corresponding gap in GRCh38-Y (**Table 1**). The annotated ampliconic gene copies in T2T-Y were largely concordant with copy numbers estimated from Illumina reads and droplet digital PCR (ddPCR)^37^, confirming the accurate copy number representation of the ampliconic genes in T2T-Y (**Supplementary Table 6-9**). RNA-Seq data confirmed expression of the annotated ampliconic genes in testis^38^. Only six genes differed in their annotation between GRCh38-Y and T2T-Y, due to presumed Y haplogroup differences (**Supplementary Table 10**).

### Repeat annotation

We generated comprehensive repeat annotations, incorporating repeat models previously updated with CHM13^39^, as well as 29 previously unknown repeats identified in T2T-Y (**Extended Data Fig. 4a, Supplementary Table 11**). The newly added sequences increased the percentage of identifiable repeats on the Y chromosome from 66.3% to 84.9%, or 17.5 Mb of non-N bases in GRCh38-Y compared to 53 Mb of bases in T2T-Y (**Table 1, Supplementary Tables 12-13** and **Supplementary Fig. 8**). While short interspersed nuclear elements (SINEs), specifically *Alu*s, are found embedded as part of the human satellite 1 (HSat1) units across most of the q-arm, other transposable elements (TEs: long-interspersed nuclear elements (LINEs), long-terminal repeats (LTRs), SINE-VNTR-*Alu*s (SVAs), DNA transposons, and Rolling circles) are completely absent (**Fig. 1**). Moreover, TE distribution biases typify different subregions of ChrY, as *Alu*s are enriched in the PAR1 region, while other TEs (particularly L1s) are more abundant in the X-transposed region (XTR)^1^ (**Extended Data Fig. 4b-c** and **Supplementary Table 14**). The DYZ19 region is annotated by RepeatMasker entirely as LTRs (**Extended Data Fig. 4c**), but further analyses indicate this is a satellite array spanning 265 kb whose 125 base monomeric consensus is derived from an expanded portion of a LTR12B sequence^40^. Repeat discovery and annotation of T2T-Y also allowed for improved annotation of ChrX in both HG002 and CHM13, particularly in the PAR regions, adding ∼33 kb of satellite annotations per ChrX (**Supplementary Table 15**).

In addition, we searched for TE driven transductions mediated by L1s and SVAs. We detected six potential 3’ L1 transductions within the T2T-Y, yet no SVA-driven DNA transductions (**Supplementary Table 16**). Despite a genome-wide investigation of both T2T-CHM13+Y and GRCh38, we were not able to locate any potential donor elements, which confirms a prior analysis that found no evidence for DNA transduction between the Y and the rest of the genome^41^. The transduction rate in T2T-Y was also much lower (0.096 per 1 Mb) than the transduction rate observed in the CHM13 autosomes (avg. 6.9 per 1 Mb) and ChrX (10.19 per 1 Mb)^39^ (**Supplementary Note 1**).

In the T2T-Y, we identified a total of 825,526 repetitive sequence motifs capable of forming alternative DNA structures (non-B DNA), compared to only 138,640 in GRCh38-Y (**Supplementary Table 17**, **Supplementary Note 2**). This nearly 6-fold increase is largely attributed to our use of novel and improved experimental and computational methodology, as non-B DNA motifs, which might form structures during sequencing, are notoriously difficult to sequence through^42^. We found a particular enrichment of these motifs at the newly sequenced centromeric region (see below) and heterochromatic region on the Yq arm (**Fig. 1**).

### Ampliconic genes in composite repeats

Composite repeats are a type of segmental duplication that are typically arranged in tandem arrays, likely derived through unequal crossing over that contributed to their increased copy numbers^1, 37^. The *TSPY, RBMY*, and *DAZ* ampliconic gene families are all associated with composite repeats on the Y chromosome, and the T2T-Y assembly provides an opportunity to analyze the complete structure of these arrays (**Fig. 2**).

*TSPY* contains the largest number of protein-coding copies on the Y chromosome and is only expressed in testis. Expression level of this gene is dosage dependent and the copy number is polymorphic between individuals^43^. In GRCh38-Y, the *TSPY* array includes a 40 kb gap and a limited number of intact protein-coding copies. Our T2T-Y assembly resolved 45 protein-coding *TSPY* copies, including *TSPY2*, which was found downstream of the *TSPY* array in the distal part of the proximal inverted repeat IR3, in contrast to GRCh38-Y where it is located upstream, possibly due to translocations between the IR3 pairs. The distal positioning of *TSPY2* in HG002 was confirmed among all other Y haplogroups except R and Q, which match the proximal positioning of GRCh38-Y^32^. All 44 protein-coding copies in the *TSPY* array are embedded in an array of composite repeat units (∼20.2 kb in size, matching prior reports^1, 43^), with one composite unit per gene (**Fig. 2a** and **Supplementary Table 18**). Each unit includes five new repeat annotations (fam-*), several retroelements in the LINE, SINE, and LTR classes, and simple repeats. This 931 kb array is the largest gene-containing composite repeat array in the T2T-CHM13+Y assembly outside of the rDNA locus, and the third largest overall (the first being the rDNA arrays followed by an LSAU-BSAT composite array on Chr22^39^).

Data from 187 SGDP samples confirmed high *TSPY* sequence conservation but copy number varied from 10–40 copies (**Fig. 2b**). Phylogenetic analysis using protein-coding *TSPY*s from a Sumatran orangutan (*Pongo abelii*) and a Silvery gibbon (*Hylobates moloch*) as outgroups confirmed that all protein-coding *TSPY* copies (including *TSPY2*) originated from the same branch, which is separated from the majority (all but one) of *TSPY* pseudogenes (**Extended Data Fig. 6**). This result contradicts earlier findings, which concluded that *TSPY2* originated from a different lineage^44^.

The composite structure of *RBMY* is similar to that of *TSPY* (one composite unit per gene), is comparable in size (with *RBMY* at 23.6 kb), and includes LINEs, SINEs, simple repeats, and eight new repeat annotations (**Fig. 2c**). In contrast, the *DAZ* locus is structured such that the entire repeat array, consisting of 2.4 kb composite units each containing a new repeat annotation and a fragmented L3, falls within one gene annotation (**Fig. 2d**). Out of the three composite arrays described here, *DAZ* is the only one also found on an autosome (Chr3, *DAZL*)^45^, although as a single unit and lacking the young LINE1 (L1PA2) insertion of the ChrY *DAZ* copies.

### Centromere

Normal human centromeres are enriched for an AT-rich satellite family (∼171 base monomer), known as alpha satellite, typically arranged into higher-order repeat (HOR) structures and surrounded by more diverged alpha and other satellite classes^46^. Each HOR copy is nearly identical and comprises a tandemly arrayed set of monomers. We annotated 366 kb of alpha satellite in T2T-Y, spanning 317 kb of the DYZ3 HOR array. While the individual units within the HOR array are highly similar (99.5–100%), three HOR subtypes were identified from the full-length repeat units based on their monomer structure (red, blue, and green HOR haplotypes in **Fig. 3, Supplementary Figs. 9-13 and Supplementary Tables 19-20)**. The majority of the T2T-Y centromeric array is composed of 34-mer HORs with a small expansion of a 36-mer, and with longer HOR variants observed in the flanking p-arm (42-mer) and q-arm (46-mer). These variants are structurally different from the RP11 centromere, which is the basis for the GRCh38-Y centromere model and was recently finished by ONT sequencing^47^ (**Fig. 3**).

Methylated CpG sites called by both HiFi and ONT reads reveal two adjacent regions of hypomethylation (separated by approximately 100 kb) in the centromeric dip region (CDR) (**Fig. 3**), which has been reported to coincide with the CENP-A binding and is the putative site of kinetochore assembly^46^. In the T2T-Y centromere, the presence of two distinct hypomethylated dips per chromatin fiber was confirmed by inspection of single-molecule ONT reads (**Supplementary Fig. 14**). A similar pattern of multiple methylation dips within a single centromere was observed in other T2T-CHM13 chromosomes such as Chr11 and Chr20^48^. In addition, the HORs contained abundant inverted, A-phased, and mirror repeat motifs, forming a periodic pattern occurring every 5.7 kb (**Fig. 3** and **Supplementary Table 17**). Such non-B DNA motifs, inverted repeats in particular, potentially forming cruciforms, are hypothesized to play a functional role in defining human Y centromeres^49^ and their presence is confirmed here at the sequence level.

### Sequence classes and palindromes

We annotated sequence classes on the T2T-Y as ampliconic, X-degenerate, X-transposed, pseudoautosomal, heterochromatic, and other, in accordance with Skaletsky *et al*^1^. In addition, we were able to classify a more precise annotation for the satellites (including DYZ17 and DYZ19) and the centromere (**Fig. 1** and **Supplementary Table 21**). The X-degenerate and ampliconic regions were estimated to be 8.67 Mb and 10.08 Mb in length, in concordance with previous findings^1^. The T2T-Y ampliconic region contains eight palindromes, with palindromes P4–P8 highly concordant with GRCh38-Y (i.e. in terms of arm, spacer lengths, and sequence identity). Arm-to-arm identity of these five T2T-Y palindromes nested within X-degenerate regions ranged from 99.84–99.96% (**Supplementary Table 22-23**). Palindromes P1–P3 harbor the AZFc region, which contains genes critical for sperm production^51^. We discovered a large polymorphic inversion (>1.9 Mb) in respect to GRCh38-Y that likely arose from a single non-allelic homologous recombination event. Using Strand-seq, we were able to locate the breakpoints at two “red” amplicons (naming according to Kuroda-Kawaguchi *et al.*^52^): one forming the P2 palindrome and the other inside the P1 palindrome (**Fig. 4**). This rearrangement was previously annotated as the “gr/rg” (green-red/red-green) inversion with variable breakpoints and was confirmed to be present across six Y-chromosome haplogroups out of 44 genealogical branches^53^. Another inversion was detected in P3, which was recently reported as a recurrent variation in human^54^ (**Extended Data Fig. 7a**). Although inversions between amplicons are believed to serve as substrates for subsequent AZFc deletions and duplications that might affect sperm production^53, 55–57^, pinpointing the breakpoints and measuring the frequency of the polymorphic inversions was difficult because of the large size and high identity of the palindromic arms.

### Composition of the q-arm heterochromatin

The human Y chromosome contains a large heterochromatic region at the distal end of the q arm (Yq12), which consists almost entirely of two interspersed satellite sequences classically referred to as DYZ1 and DYZ2^58–61^. The single largest gap in GRCh38-Y is at Yq12, with minimal representation of DYZ1 and DYZ2, mostly in unplaced scaffolds. Here, we uncovered the detailed structure of the Yq12 region at single-base resolution, characterizing over 20 Mb of DYZ1 and 14 Mb of DYZ2 repeats. In T2T-Y, DYZ1 and DYZ2 are interspersed in 86 large blocks, with DYZ1 blocks ranging from 80–1,600 kb (median of 370 kb) and DYZ2 blocks ranging from 20–1,200 kb (median of 230 kb). DYZ2 blocks appear more abundant at the distal end of Yq12, and this trend is also visible in metaphase chromosome spreads with fluorescence *in situ* hybridization (FISH) (**Fig. 5a-b** and **Fig. 1** Cen/Sat track). Yq12 is highly variable in size and sequence structure between individuals^62–64^, and the number and size of these satellite blocks is expected to vary considerably.

**Fig. 5.**
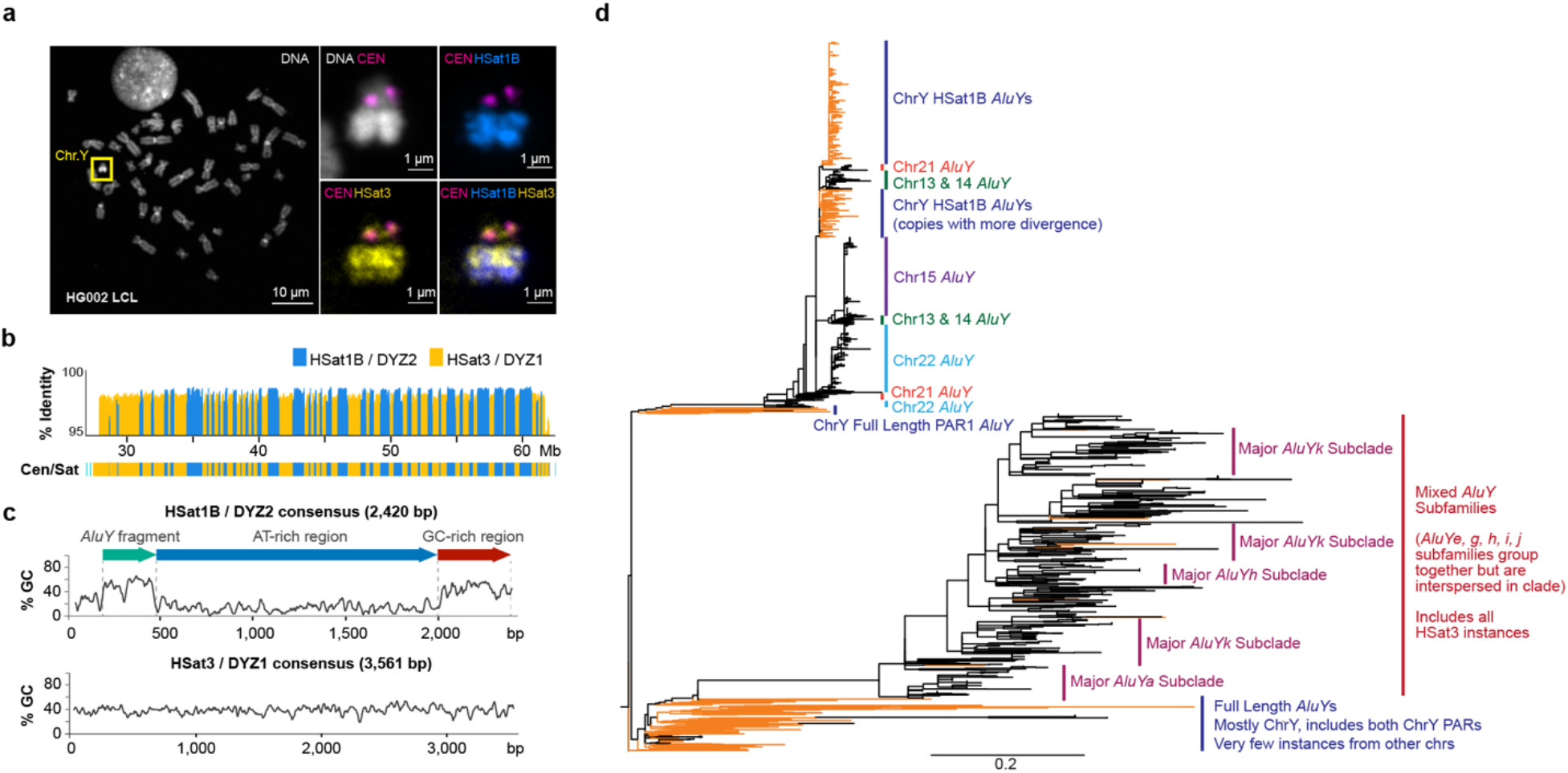
Heterochromatic region of the distal Y q-arm (Yq12). **a.** FISH painting of the Y chromosome, centromere/DYZ3 (magenta), HSat1B (blue), and HSat3 (yellow). Top-left, overall chromosome labeling by DNA dye (DAPI) with ChrY highlighted in an HG002-derived lymphoblastoid cell line (GM24385). The right panels show ChrY labeled with FISH probes recognizing centromeric alpha satellite/DYZ3 (magenta), HSat3/DYZ1 (yellow), and HSat1B/DYZ2 (blue). In concordance with the T2T-Y assembly, HSat3 probes indicate the presence at DYZ17 (close to centromere) as well as a slight enrichment to the proximal part of the Yq12 (DYZ1), while HSat1B is only present in the Yq12 and is more enriched towards the distal part (DYZ2). Maximum intensity projections are shown in all panels. The results of this experiment were replicated using two different sets of PCR probes. Fifteen large-field images containing at least 20 spreads were analyzed per condition. **b.** % identity of each DYZ2/DYZ1 repeat unit to its consensus sequence. **c**. % GC sequence composition of the HSat1B/DYZ2 and HSat3/DYZ1 repeat units and the position of an ancient *Alu*Y fragment in DYZ2. **d**. Phylogenetic tree of *Alu*Y sequences associated with HSat1B and HSat3, rooted on *Alu*Sc8. Tree represents subsampling of *Alu*Y elements, both full length (FL) and truncated, including *Alu*Y sequences found within HSat1B units and associated with HSat3 arrays. Elements located on ChrY are denoted with orange branches. The scale bar represents 0.2 substitutions per site on a branch of the same length.

DYZ1 is composed of a Y-specific subfamily of HSat3 sequences that occurs primarily as ∼3.6 kb nested tandem repeats derived from an ancestral tandem repeat of the pentamer CATTC^65^. DYZ2 is composed of an unrelated satellite family, HSat1B, and comprises a ∼2.5 kb tandem repeat made up of three parts: an ancient *Alu*Y subunit (20% diverged from the *Alu*Y consensus), an extremely AT-rich region (>85% AT), and a more GC-rich region^61, 65, 66^. The vast majority of repeat instances were over 98% identical, with slightly higher divergence at the more peripheral satellite blocks (**Fig. 5c**). A detailed comparison of the sequences within T2T-Y revealed recent structural rearrangements including iterative, tandem duplications as large as 5 Mb, which span multiple blocks of DYZ1 and DYZ2 (**Extended Data Fig. 8**). These structural rearrangement patterns are consistent with evolution by unequal exchange mechanisms. In addition, approximately 15% of Strand-seq libraries showed sister chromatid exchanges within the Yq12 heterochromatic region (**Extended Data Fig. 7b**).

While HSat3 is present across multiple chromosomes including the acrocentric short arms, HSat1B is present almost exclusively on the Y and the acrocentric short arms in smaller amounts, with the exception of Chr10^67^. While HSat1B carries an *Alu*Y-derived subunit as part of its composite repeat unit, some HSat3 arrays are tightly associated with *Alu* sequences, with blocks of HSat3 intermingled with *Alu* fragments, including *Alu*Y. Phylogenetic analyses place the ChrY HSat1B *Alu*Y subunits in a cluster with *Alu*Y subunits found in HSat1B sequences on the acrocentric chromosomes, with the highly homogenized ChrY copies appearing as a single cluster. Given the topology of this tree, it appears that the HSat1B sequences found on the acrocentric chromosomes were derived from the Y-linked HSat1B, with seeding events leading to local expansion and homogenization (**Fig. 5d**, upper branches). The *Alu*Y fragments found interspersed with HSat3 on the Y chromosome also phylogenetically cluster with *Alu*Y fragments associated with HSat3 on the acrocentric chromosomes. However, there is no evidence for local homogenization of HSat3-*Alu* fragments; likewise, there is no support for phylogenetic clustering by subgroup nor by chromosome. Based on the deep divide between the HSat1B and HSat3 clades in the tree for both ChrY and the acrocentric chromosomes, we conclude that the initial seeding events that created these arrays were independent of one another, yet were derived from *Alu*Y elements from PAR1 (**Fig. 5d**, lower branches).

## Improved variant calling for XY samples

We performed short-read alignment and variant calling for 3,202 samples (1,603 XX; 1,599 XY) from the 1KGP Phase 3, including 1,233 unrelated XY samples averaging at least 30× coverage of 150 bp paired-end reads^27^. This set of 1,233 XY samples spans all 26 geographically diverse 1KGP populations and 35 distinct Y-chromosome haplogroups (**Supplementary Table 24**). To more accurately represent the diploid nature of the PARs, we completely hard-masked ChrY in XX samples and ChrY PARs in XY samples, thereby forcing any reads originating from the ChrY PAR to align to the ChrX PAR (**Supplementary Tables 25-28** and **Extended Data Fig. 9**). Diploid genotypes were then called within the PAR for both XX and XY samples^68^ (**Extended Data Fig. 10a**). Aside from this modification, the alignment and variant calling pipeline mirrored our previous analysis based on GRCh38-Y^69^.

Across all 1,233 unrelated XY samples, we observed improved alignment to T2T-Y, including a higher number of mapped reads (increase of 1.4 million reads on average, SD: 432,115; **Fig. 6a**), a higher proportion of properly paired reads (increase of 1.4% on average, SD: 1.4%; **Fig. 6b**), and a lower proportion of mis-matched bases (decrease of 0.6% on average, SD: 0.06%; **Fig. 6c**) per sample relative to GRCh38-Y (**Supplementary Table 29**). Within syntenic regions of the two Y chromosome assemblies, the number of variants per sample declined for samples from all Y haplogroups with the exception of haplogroup R (haplogroup of GRCh38-Y) and with the greatest reduction observed for samples of haplogroup J1 (haplogroup of T2T-Y) (**Fig. 6d, Extended Data Fig. 10b-c**). Selecting one individual each from the J1, R1b, and E1b haplogroups, we compared per-variant read depth and allele balance for both references (**Fig. 6e**). In all three samples, we observed more variants with excessive read depth and abnormal allele balance on GRCh38-Y, corresponding with putative collapsed duplications (**Supplementary Table 30**, **Fig. 6f-g**). We replicated these analyses using an additional 279 samples across 142 populations from the SGDP^28^ and found similarly improved mappings and variant discovery using T2T-Y (**Extended Data Fig. 10d-e, Supplementary Figs. 16-18**).

**Fig. 6.**
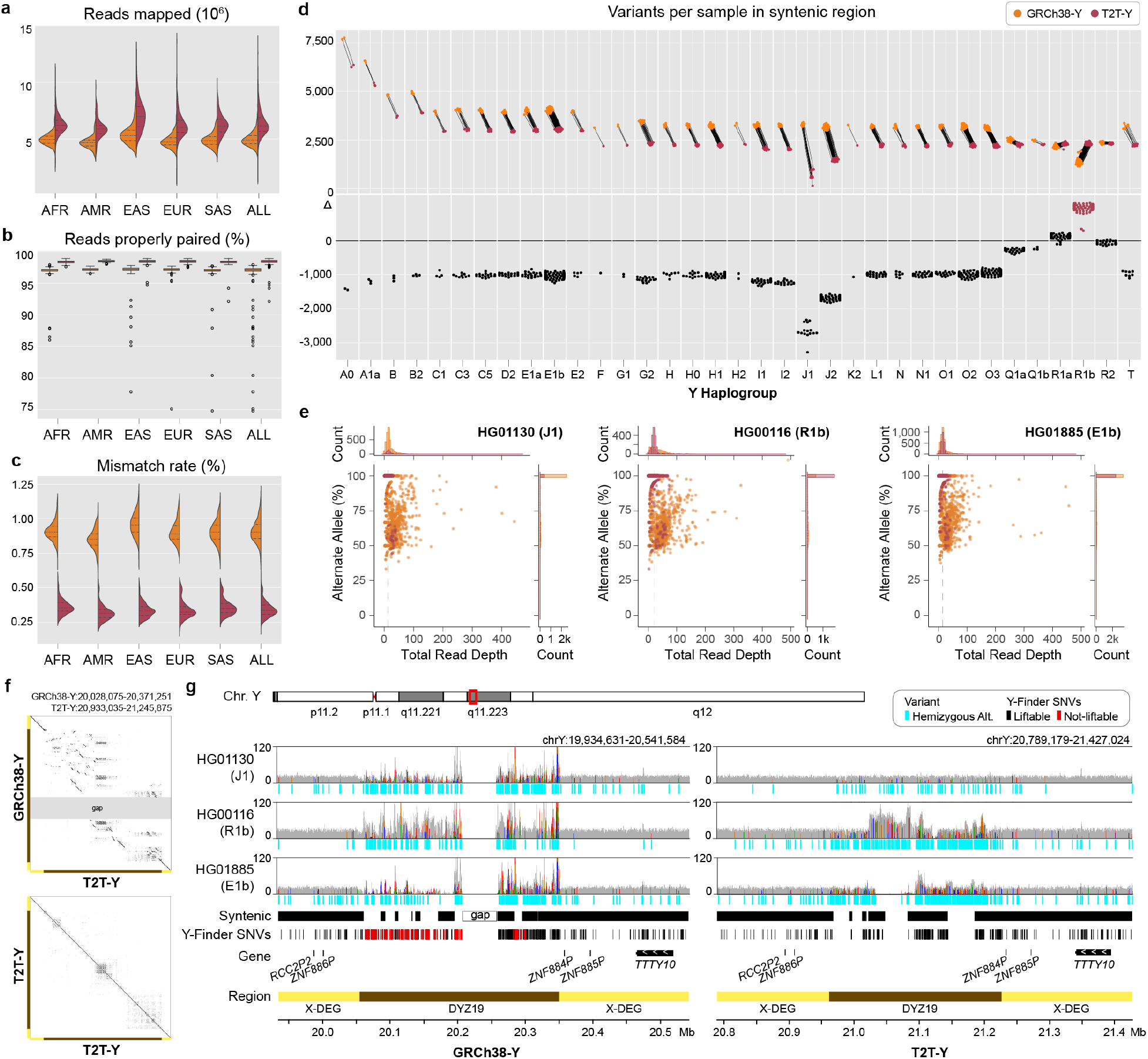
Short-read mappability and variant calling improvements on T2T-Y. In all plots, GRCh38-Y is colored orange and T2T-Y is maroon. The complete sequence of T2T-Y improves short-read alignment of the 1KGP dataset by **a.** increased number of reads mapped, **b**. higher portion of reads properly paired, and **c.** lower mismatch rate compared to GRCh38-Y. Bar in the box plot represents the 1st, 2nd (median), and 3rd quartile of the data. Whiskers are bound to the 1.5 × interquartile range. Data outside of the whisker ranges are shown as dots. **d.** The number of called variants within syntenic regions is reduced on T2T-Y for all haplogroups except R1 (haplogroup of GRCh38-Y). **e.** Further investigation on 3 samples (J1, R1b, and E1b) shows a higher number of variants called with excessive read depth and variable alternate allele fractions for GRCh38-Y. Each dot represents a variant, with the % alternate alleles as a function of total read depth. Dotted line represents the median coverage on T2T-Y, close to the expected 1-copy coverage. **f.** Dotplot of the DYZ19 array between GRCh38-Y and T2T-Y and self-dotplot of T2T-Y. Large rearrangements are observed, with multiple inversions proximal to the gap in GRCh38-Y with respect to T2T-Y (top), while more identical, tandem duplications are visible in T2T-Y (bottom). **g.** Read pile-ups and variants on DYZ19 for GRCh38-Y (left) and T2T-Y (right) as shown with IGV^71^. Gray histogram shows the mapped read coverage, with colored lines indicating non-reference bases with >60% allele frequency. Regardless of the haplogroup, the incomplete DYZ19 array in GRCh38-Y hinders interpretation. Syntenic regions between the two Ys are marked, and SNV sites used to identify Y haplogroup lineages in Y-Finder are shown below, with variants liftable from GRCh38-Y to T2T-Y in black, not-liftable in red, respectively.

Due to genomic repeats, accuracy of short-read variant calling is heterogeneous across the genome. One approach for improving reliability is to restrict analysis to “accessible” regions based on various alignment metrics. To this end, we followed published protocols to generate a short-read accessibility mask for T2T-Y based on patterns of normalized read depth, mapping quality, and base-calling quality^70^. Our masks reveal that while the heterochromatic long arm (Yq) remains largely inaccessible to short-read analysis, T2T-Y still adds 578 kb of accessible sequence compared to GRCh38-Y (increase of 4.2%, **Table 1**).

Taken together, these analyses indicate the complete T2T-Y reference improves short-read alignment and variant calling across populations and corrects errors in GRCh38-Y, but acknowledging the rich resources available on GRCh38, we also curated a 1-to-1 whole-genome alignment between each GRCh reference (GRCh37 and GRCh38) and T2T-CHM13+Y to enable lifting annotations in either direction. The vast majority of genetic variants in ClinVar (2022-03-13 release), dbSNP (build 155), and GWAS Catalog (v1.0 release) were successfully lifted to T2T-CHM13+Y (99.2%/97.8%/98.9% overall and 100%/95.0%/100% for ChrY, respectively, **Supplementary Table 31**). Accessibility masks and lifted annotations are provided along with variant calls as a resource for future studies (**Data Availability**).

## Contamination of genomic databases

Human DNA sequences can sometimes appear as contaminants in the assembled genomes of other species. In microbial studies, the human reference sequence has been used to screen out contaminating human DNA; however, due to the incomplete nature of the current reference, some human fragments are missed and mistakenly annotated as bacterial proteins, leading to thousands of spurious proteins in public databases^72, 73^. For example, a recent analysis of nearly 5,000 human whole-genome data sets found an unexpected linkage between multiple bacterial species and human samples of XY karyotype, including 77,647 100-mers that were significantly enriched in the XY samples^74^. The authors hypothesized that these bacterial genomes were not actually present in the samples, but rather the effect was caused by real human ChrY sequences matching to contaminated bacterial genome database entries. We compared XY-enriched 100-mers from the Chrisman *et al.* study^74^ to the T2T-Y chromosome and found that, as predicted, more than 95% of them had near-perfect matches to the complete T2T-Y sequence.

We further tested the entire NCBI RefSeq bacterial genome database (release 213, July 2022, totalling 69,122 species with 40,758,769 contig or scaffold accessions) and identified all 64-mers that appeared in both the bacterial database and T2T-Y. We found 4,179 and 5,148 potentially contaminated sequences matching GRCh38-Y and T2T-Y, respectively. (**Extended Data Fig. 11a,** top and middle). The sequences were relatively short in length (<1 kb), as is typical of contaminating genomic segments (**Extended Data Fig. 11b**). The vast majority of contaminated sequences found only with T2T-Y localized to the newly added HSat1B and HSat3 repeats (**Extended Data Fig. 11c**, **Supplementary Table 32**). Such repetitive sequences are common sources of contamination because of their high copy-number. We predict this human-derived sequence contamination issue includes sequence from all human chromosomes and extends to all sequence databases, including non-microbial genomes.

## Discussion

Owing to its highly repetitive structure, the human Y chromosome is the last of the human chromosomes to be completed from telomere to telomere. Here, we have presented T2T-Y, a complete and gapless assembly of the Y chromosome from the HG002 benchmarking genome, along with a full annotation of its gene, repeat, and organizational structure. We have combined T2T-Y with the prior T2T-CHM13 assembly to construct a new reference, T2T-CHM13+Y, that is inclusive of all human chromosomes. This assembly, along with all of the annotation resources presented here, is available for use as an alternative reference via NCBI and the UCSC Genome Browser^75^ (**Data Availability**).

Our analysis of the T2T-CHM13+Y reference assembly reveals a reduction in false-positive variant calls for XY-bearing samples due to the correction of collapsed, incomplete, misassembled, or otherwise inaccurate sequences in GRCh38-Y. Given the history of the GRCh38-Y assembly and its reliance on BAC libraries, we see no feasible means for its completion and suggest T2T-Y as a more suitable ChrY reference going forward. We recommend the use of T2T-CHM13 when mapping reads from XX samples and ChrY-PAR-masked T2T-CHM13+Y when mapping XY samples (**Supplementary Note 3**).

The completion of ampliconic and otherwise highly repetitive regions of ChrY will also require updates to existing gene annotations that are based on the incomplete GRCh38-Y assembly. How to label and refer to genes within variable-size ampliconic arrays, like *TSPY*, is an open question. Moreover, the highly repetitive sequences pose new challenges to computational tools developed on GRCh38. One example is the inconsistent methylation pattern observed in the satellite enriched heterochromatin region, in which both HiFi and ONT are prone to sequencing biases, hindering accurate biological interpretation (**Supplementary Note 4** and **Supplementary Fig. 19**). Lastly, we have noted the improved detection of human contamination in genomic databases using T2T-CHM13+Y and recommend a full contamination audit of public genome databases using this updated human reference. Taken together, these results illustrate the importance of using a complete human reference genome for bioinformatic analyses.

Construction of the T2T-Y assembly challenged the assembly methods developed for the haploid CHM13 genome and spurred the development of new, automated methods for diploid human genome assembly. In particular, the PARs of the HG002 sex chromosomes required phasing akin to heterozygous, diploid haplotypes, and the palindromic and heterochromatic regions of ChrY required expert curation of the initial assembly string graph. Lessons learned from our assembly of T2T-Y informed the development of the Verkko assembler^76^, which automates the integration of HiFi and ONT data for diploid human genome assembly. The companion study of Hallast *et al.*^32^ successfully used Verkko to generate 43 near-T2T assemblies from a diverse panel of human Y chromosomes, revealing dynamic structural changes within this chromosome over the past 180,000 years of human evolution. Ultimately, as the complete, accurate, and gapless assembly of diploid human genomes becomes routine, we expect “reference genomes” will become known as simply “genomes”.

Projects such as the Human Pangenome Reference Consortium^77^ are in the process of generating high-coverage HiFi and ONT sequencing for hundreds of additional human samples, and the assembly of these diverse, complete human genomes, along with similar quality assemblies of the non-human primates, will provide an unparalleled view of human variation and evolution. With the availability of complete, diploid human genome assemblies, association between phenotype and genotype will finally move beyond small variants alone and be made inclusive of all complex, structural genome variation.

## Online Methods

This section provides a brief summary of the methods. Refer to the **Supplementary Methods** for more details.

### Sequencing

Seventeen PacBio HiFi WGS runs were generated from GM24385 using the SMRTbell Express Template Prep Kit 2.0 on the Sequel II platform, after size-selection for 15 to 25 kb fragments. All of the ONT WGS runs are from the Jarvis *et al*.^20^ study, which were generated using protocols from Shafin *et al*.^78^ and Jain *et al*.^11^.

RNA was extracted from three cell lines to generate Iso-Seq reads: EBV-immortalized lymphoblastoid cell line (GM24385), iPSC of the EBV-immortalized lymphoblastoid cell line (GM26105), and iPSC derived directly from Peripheral Blood Mononuclear Cells (GM27730). Iso-Seq data was generated on the Sequel II platform and was processed using Lima and IsoSeq3.

Specific runs used either in assembly or validation along with their accessions are available in **Supplementary Table 1**, and DNA preparation and library generation information is available in **Supplementary Methods**.

### Assembly and validation

An assembly string graph was first constructed using PacBio HiFi reads (∼60x) and processed using custom pruning procedures. Due to high sequence similarity within PAR1 and PAR2, the HG002 ChrX and ChrY string graph components shared connections to one another in the PARs, but to no other chromosomes in the genome. The remaining tangles in these sex-chromosome subgraphs were resolved using ONT reads longer than 100 kb (∼90x). A semi-automated repeat resolution strategy utilized GraphAligner^13^ to map the ultra-long ONT data to the HiFi assembly graph and identify the correct traversals. To resolve the PAR regions, ChrX and ChrY chromosomal walks were identified using homopolymer compressed trio-binned k-mers from parental Illumina reads^79^, and a consensus sequence was computed for each. Remaining coverage gaps caused by HiFi sequencing biases were patched using a *de novo* Flye assembly of trio-binned paternal ONT reads^14, 80^. For new projects, we now recommend the automated Verkko pipeline^76^, which is able to replicate the semi-manual T2T-Y assembly presented here.

For polishing, the ChrXY draft assemblies were combined with the T2T-CHM13v1.1 autosomes to prevent mapping biases caused by the incompletely resolved autosomes of HG002 (Hereby T2T-CHM13+XY). For further polishing and validation, we used 66× Illumina, 84× HiFi, and 250× ONT (being haploid, the effective coverage on X and Y is half those depths). Small corrections were identified with DeepVariant^81, 82^ and filtered with Merfin^14^. Large errors were identified with Sniffles^83^, cuteSV^84^, and through a comparison to the HPRC-HG002v1 assembly^20^. All of the large errors were patched using marker-assisted HiFi and ONT reads. Assembly issues were identified using repeat-aware long-read alignments from Winnowmap2^16^ (filtered with globally unique markers^17^) and VerityMap^85^ (guided by locally unique markers). Coverage summaries were obtained using scripts from the T2T-CHM13 assembly evaluation^17^. Putative collapses and inversion errors were identified using Strand-seq data. Raw sequencing reads from 65 Strand-seq libraries^30, 31^ were aligned to both GRCh38 and T2T-CHM13+XY with BWA^86^, then processed with breakpointR^30, 87^ to identify inversion errors. Recurrent inversions were identified by comparing to results from Porubsky *et al.*^54^. To further confirm integrity of ChrY in the HG002 cell-line, we aligned publicly available GIAB^22^ HiFi and ONT reads from the paternal HG003 genome (including from the PacBio Revio platform^88^), and performed the same coverage analysis. Base error rate was measured by Merqury using a hybrid k-mer set from Illumina and HiFi reads^17, 18^ (**Supplementary Table 3**).

### Comparison to GRCh38-Y

#### Y haplogroup identification

The Y-chromosome haplogroup of HG002 and GRCh38 was identified using yhaplo^89^, which builds a tree from phylogenetically informative SNPs that accumulate in the non-recombining portion of the Y. The Y-haplogroups of the 1KGP samples were identified using Y-Finder^90^, using SNP calls on GRCh38 from Aganezov et al^69^.

#### Alignments between GRCh38 and HG002 Y assemblies

Alignments between the GRCh38-Y and T2T-Y assemblies for the purposes of visualization with SafFire were generated with minimap2^91^. The PAF was then processed with rustybam and visualized with SafFire. DupMaske^35^ and dna-brnn^92^ annotations were generated using Rhodonite (10.5281/zenodo.6036498).

A complementary alignment was generated with LASTZ^93^ after softmasking repeats from WindowMasker^94^. The alignment dotplot and best identity were plotted using R (https://github.com/arangrhie/T2T-HG002Y/tree/main/alignments/lastz). Regions along T2T-Y were colored according to their class.

To visualize three big structural differences of the three ChrY assemblies (GRCh37-Y, GRCh38-Y, and T2T-Y), we used the Pangenomics Research Tool Kit^95^ to construct principal bundles representing contiguous and conserved sequences among the pangenome contigs.

### Gene annotation

#### GENCODEv35 CAT/Liftoff annotation

Preliminary gene annotation was performed by mapping GENCODEv35^96^ annotations from GRCh38-Y to T2T-Y using a Cactus^97^ alignment with Chimp as an outgroup. Iso-Seq reads were aligned and assembled with Stringtie2^98^, aligned to the assembly with TransMap^99^, and used as input for CAT^100^ along with the GENCODEv35 annotation. GENCODEv35 Y annotations were mapped with Liftoff^101^, then intersected with Bedtools^102^ to isolate genes that Liftoff mapped to ChrY that were not in the CAT annotation.

#### *De novo* RefSeqv110 and GENCODEv38 annotation

In the meantime, a *de novo* RefSeq annotation was performed on both GRCh38 and T2T-CHM13+Y and released (v110) as previously described for other vertebrate genomes^103, 104^. A total of 82,862 curated RefSeq transcripts, 345,700 cDNAs, 8.65 million ESTs, 9.7 billion RNA-Seq reads, and 83 million PacBio IsoSeq and Oxford Nanopore reads from over 30 distinct tissues were retrieved from SRA and tentatively aligned to the assembly using Splign^105^ or minimap2^91^.

Simultaneously, an Ensembl gene annotation was performed by a mapping subset of the genes from GENCODEv38^96^ using minimap2^91^ and MAFFT^106^. Transcripts with low coverage or identity (<98%) were re-aligned using Exonerate^107^. Genes in potential recent duplications or collapsed paralogues were adjusted accordingly.

#### RefSeq/Liftoff, curated ampliconic gene annotation

Because the additional copies of the ampliconic genes hindered comparison to known genes in GRCh38-Y with differing gene IDs and names, we performed one more annotation by mapping GRCh38 RefSeq v110 annotations with Liftoff to T2T-CHM13+Y. We compared ampliconic gene family annotation results from that of GENCODE CAT/Liftoff and assigned gene names followed by best gene coverage and identity, including introns. Later, based on discussions with authors of a companion paper^32^, we adjusted the gene names for 3 protein-coding annotations based on exon sequence identity (**Supplementary Table 5**).

#### Validation of the ampliconic protein-coding genes

Copy numbers for each ampliconic gene family in both the GRCh38-Y and T2T-Y assemblies were estimated using an adapted application of AmpliCoNE^37^. Copy numbers of these gene families were previously estimated for HG002 using Illumina reads from GIAB^108^ and droplet digital PCR (ddPCR)^37^. The only notable difference was in the *TSPY* copy number, in which we identified 45 intact protein-coding copies. The copy number was slightly higher in the assembly than the estimates derived from Illumina reads and ddPCR (45 vs. 40 and 42, respectively). The *in silico* PCR primer search matched all 44 protein coding copies in the *TSPY* gene array and *TSPY2*, and two pseudogenes at the 3’ end of the *TSPY* array which we were unable to avoid in the ddPCR primer design. We conclude that our AmpliCoNE, ddPCR, and *in silico* PCR estimates agree with the ampliconic gene annotations in the T2T-Y assembly (**Table 1**).

### Repeat annotation

#### Segmental duplications

Segmental duplication (SD) annotations were created using the same methods as in Vollger *et al.* without modification^3^. In brief, SDs in T2T-CHM13+Y were identified using SEDEF^109^ after repeat masking with Tandem Repeats Finder^110^ and RepeatMasker^111^.

#### Repeat model discovery and annotation

A three-step repeat annotation was performed to annotate new repeat models on ChrY. First, RepeatMasker was performed on the T2T-Y assembly using the Dfam 3.3 library^112^, hard-masking previously annotated repeats. Second, RepeatModeler analysis was performed on the remaining unmasked regions to identify new repeat model consensuses, which were subjected to extension and filtering, and used as a library for a secondary RepeatMasker run. Two methods were primarily used to identify new satellite repeats: ULTRA^113^ and NTRprism^46^. Unannotated regions>5 kb were identified via Bedtools^114^ by subtracting repeat annotations from first and last steps above. The regions were manually curated in UCSC Genome Browser to check for any feature overlap (e.g. gene annotations). Tandemly repeated sequences were detected and assessed with a combination of ULTRA, NTRprism, and the TRF GUI version^110^ to determine the best monomer consensus for a given satellite model. Lastly, the compilation pipeline laid out in Hoyt *et al.*^39^ was followed to avoid potential false positives by simply masking with a combined library of new repeat models and known repeat models (Dfam library). The same three-step repeat annotation pipeline was applied to GRCh38-Y as well. Repeats were summarized using buildSummary.pl^115^ at the class and family level (**Table 1**, **Supplementary Table 12**) and at the subfamily level for new repeats (**Supplementary Table 11**) in both T2T-Y and GRCh38-Y.

#### Composite repeats

Composite elements were defined and characterized as described in Hoyt *et al.*^39^ as repeating units consisting of three or more repeated sequences, including TEs, simple repeats, composite subunits, and/or satellites, that are found as a tandem array in at least one location in the genome. BLAT^116^ was used to locate other composite unit copies across T2T-Y and cross-reference them with their associated gene annotations (CAT/liftoff). Identification of the potentially active, full length TEs (SINEs, LINEs, and retroposons are AluY, L1Hs, and SVA_E/F) across T2T-Y and GRCh38-Y was done by following the methods of Hoyt *et al.*^39^.

#### Satellite annotation

Centromeric Satellite (Cen/Sat) annotations were generated as in Altemose *et al.*^46^, with a few refinements tailored to include annotations of the entire ChrY. Major satellite types were extracted from the RepeatMasker track, with features merged for the same satellite type within 10 kb of each other. For HSat2 and HSat3, a specialized annotation tool was used (https://github.com/altemose/chm13_hsat)^46^. DYZ19 and HSat1B were annotated using RepeatMasker annotations. Exact boundaries between HSat3 and HSat1B (aka DYZ1 and DYZ2) were manually refined.

#### Transduction analysis

We utilized the same approach as Hoyt *et al.*^39^ to identify putative DNA transductions mediated by retroelements. Briefly, L1s and SVAs were identified in T2T-Y to detect the target site duplications and 3’ transduction signatures using a modified version of TSDfinder^117^. Then, we removed transductions residing in SDs and masked the transduced sequences using RepeatMasker^111^. To find the potential progenitor of each transduction within T2T-CHM13+Y and GRCh38, the offspring sequences were aligned to the corresponding databases of full-length L1s and SVAs using BLAST^118^.

#### Non-B DNA motif annotation

To predict sequence motifs with the potential to form alternative DNA structures (non-B DNA), we used nBMST^119^ for repeat motifs (A-phased, direct, inverted, and mirror repeats and STRs) and Z-DNA motifs^120, 121^. G4-motifs were detected with Quadron^122^, which also yields a score that quantifies the stability of a predicted G4 structure based on a machine-learning algorithm trained on empirical datasets. Motifs with a Quadron score ≥19 were considered stable, and thus used throughout our analysis. The non-B motif were intersected with other existing annotations of T2T-Y (gene annotations, satellite repeats, and CpG islands) using Bedtools^114^. Rideogram^123^ was used to generate these visualized tracks as well as the three composite repeat tracks. GraphPad Prism^124^ was used to generate the TE composition per sequence class plots.

### Data visualization

For **Fig. 1** and **Fig. 3**, the alignment of GRCh38-Y and T2T-Y was visualized with SafFire^125^. Segmental duplications (SDs) are colored by duplication types defined in DupMasker^35^. IGV^71^ was used to draw ideograms, sequence classes, palindromes, inverted repeats, and AZF. Bedtools^114^ was used to calculate density (across each gene type), bp coverage (across each repeat class) and average CpG methylation frequency per 100 kb window. Dotplots colored by identity were generated with StainedGlass^36^.

### TSPY gene family analysis

#### TSPY copy number estimation from SGDP

Copy number of the TSPY gene was estimated as in Vollger *et al*.^3^. In brief, we applied the fastCN pipeline^126^, which uses read-depth as a proxy. Short-read sequence data were processed into 36 bp non-overlapping fragments and mapped using mrsFAST^127^ to a T2T-CHM13+Y reference masked with TRF and RepeatMasker. Read-depth across the genome was corrected for GC bias and copy number was determined using linear regression on read-depth versus known fixed copy number control regions. Finally, integer genotypes for *TSPY* were generated by taking a weighted average of the copy number estimates from windows overlapping the locus.

#### Phylogenetic tree analysis of the TSPY genes

All curated protein-coding and pseudogene *TSPY* copies (including introns) from the CAT/Liftoff and RefSeq/Liftoff annotations were used. For outgroup rooting of the tree, *TSPY* sequences were used from *Hylobates moloch* (NW_022611649.1)^128^ and *Pongo abelii* (KP141780.1)^129^. Alignment was carried out in MAFFT^106^. Phylogenetic analysis was run in RAxML-NG^130^ with 200 bootstrap replicates with rapid bootstrap approximation. Consensus bootstrap values were then mapped to the highest likelihood phylogeny in Geneious^131^ and visualized in FigTree^132^.

### Centromere analysis

The T2T-Y was processed using the standard alpha-satellite (AS) tools as described in Altemose *et al.*^46^. The S4CYH1L (DYZ3) AS HOR was re-examined and re-defined for this paper to take into account its polymorphic variants both known from the old literature^133, 134^ and revealed by the recent complete centromere assembly of RP11^47^.

The CENP-A CUT&RUN data was aligned to the T2T-CHM13+Y assembly as previously described in Altemose *et al.* ^46^. The alignments were filtered using the single-copy k-mer locus filtering method as described in Hoyt *et al.* ^39^ through the use of the UCSC GenomeBrowser tool overlapSelect.

HG002 ONT UL data was re-basecalled using Guppy v6.1.2, Remora to obtain CpG methylation data (Supplementary Table 1). Modbams were converted to FASTQ files and aligned with Winnowmap2^16^. HG002 ONT nanoNOMe data was generated in Gershman *et al*.^48^ and analyzed with nanopolish^135^. The probability of methylation for each CpG site in PacBio HiFi reads was assigned using primrose (https://github.com/PacificBiosciences/primrose). Reads were aligned with pbmm2 (https://github.com/PacificBiosciences/pbmm2). The percent of methylated reads at each reference genome position was calculated using pb-CpG-tools (https://github.com/PacificBiosciences/pb-CpG-tools). Resulting modbams were re-processed identically to Remora-called ONT data to collect comparable aggregated native CpG methylation data. The CDR was manually annotated as the area where CpG methylation is lower than the flanking, active, alpha satellite (**Supplementary Fig. 14)**.

### Sequence classes on the Y chromosome

The X-degenerate and ampliconic regions were annotated using either exact boundaries of palindromes or intrachromosomal identity as defined in Skaletsky *et al.*^1^ with adjusted borders based on the gene annotations. T2T-Y was split into 5 kb sliding windows (step size 1 kb) and these sequences were mapped back to T2T-Y using Winnowmap2^16^. After excluding self-alignments, windows with identity >50% were considered indicative of ampliconic regions if present consecutively.

For the schematic representations in **Fig. 4**, amplicons from Teitz *et al.*^55^ were mapped to GRCh38-Y and T2T-Y assemblies with Winnowmap2^16^ to identify homologous regions.

Approximate boundaries of the palindrome P4–P8 arms were manually selected using Gepard^136^ and further refined based on aligning palindromic arms and adjacent flanks against each other (arm1 to the reverse complement of arm2) using global alignment with Stretcher^137^.

For AZFa, sequences between two HERV15 genes (including genes USP9Y and DDX3Y) were used to determine the AZFa boundaries^138^. Boundaries of AZFb and AZFc were defined by the amplicon units P5/proximal P1 deletion (yel3/yel1) and by the b2/b4 deletion. A self-dotplot of the T2T-Y assembly was used with word size of 100 in Gepard^136^. Breakpoints were identified as illustrated in Fig. 2 of Navarro-Costa *et al.*^57^ as shown in **Supplementary Fig. 15**.

The PAR and X-transposed regions were initially identified using LASTZ^93^ alignments between HG002-X, HG002-Y and CHM13-X. Exact boundaries were later refined using Minimap2^91^ alignments.

### Yq12 heterochromatin region

#### DYZ1 and DYZ2

DYZ1 and DYZ2 consensus sequences were generated using multiple sequence alignment using kalign^139^ and converted to a profile HMM using HMMER^140^. Dotplots in the **Extended Data Fig. 8** were produced using dottup of the EMBOSS software package^137^.

#### Phylogenetic tree analysis of the AluY

The AluY tree was rooted on the RepeatMasker/Dfam derived consensus sequence for AluSc8. Analysis was run on a MAFFT^106^ derived alignment using RAxML-NG^130^ with 100 non-parametric bootstrap replications. Note that in the AluY subfamily clade (“Mixed AluY Subfamilies”) there are scattered elements across the group even though the majority are represented in the labeled subclades.

### Short-read variant calling on T2T-CHM13+Y

#### Impact of masking PAR and XTR in variant calling

Simulated paired-end sequence reads were generated using NEAT^141^. Variants from 10 XY and 10 XX European individuals were collected from high coverage variant calls of 1KGP^27^ and used for benchmarking. Reads were processed with bbduk^142^ and mapped using BWA^86^ to two versions of GRCh38: X and Y both unmasked (default), and sex chromosome aware (SCC aware^68^). Masking was performed on PAR^143^ or both PAR and XTR^68, 144^. Mapping quality (MAPQ) was assessed on ChrX in each 50 kb windows, sliding 10 kb using Bedtools^114^. Variant calling was performed with GATK^145^ and compared against the chosen variants used in simulating the reads.

#### Mappability comparison and variant calling in 1KGP samples

Using the NHGRI Genomic Data Science Analysis, Visualization, and Informatics Lab-Space (AnVIL)^146^, we performed short-read alignment and variant calling for the 3,202 samples in 1KGP^27^ using the T2T-CHM13+Y assembly as a reference. These samples were sequenced to at least 30× coverage by the New York Genome Center (NYGC), and alignment and variant calling was previously performed on the GRCh38 reference. We largely followed the short-read alignment and variant calling pipeline used for analysis of T2T-CHM13v1.0^69^, except that we used SCC references for each XX and XY individuals using XYalign^68^. In the XX-specific reference, the entire Y chromosome is masked, whereas in the XY-specific reference, only the Y-PARs are masked. For all analyses, measures of mappability (reads mapped, reads properly paired, mismatch rate) were assessed with Samtools^147^, and variant counts and allele frequencies were assessed with bcftools^147^. Variants in syntenic regions between GRCh38-Y and T2T-Y were further subsetted with Bedtools^114^.

#### Putative collapsed regions in GRCh38-Y

Three individual’s variant calls and the corresponding bam files from the 1KGP dataset were downloaded from AnVIL: one individual each from the J1, R1b and E1b haplogroups (HG01130, HG00116 and HG01885, respectively). Variant calls on ChrY syntenic region were subsetted using bcftools^147^. From the VCF file, allelic read depth (defined as AD field) and reference allele depth (1st value in the DP field) were extracted using a custom script along with each variant’s chromosomal position and visualized with R. Coverage tracks of the bam files were collected with IGVtoolkit^71^ and samtools^147^. Variants from HG00116 on GRCh38-Y (R1b, thus least structural variations expected) were further aggregated as “excessive variant region” when non-reference alleles were present, merged within 50 kb. Coverage, variant calls, and the excessive variant regions were manually inspected on GRCh38-Y, and marked as a “putative collapse” if the region: 1) had an excessive number of variants called for all three samples, 2) overlapped with a known gap in GRCh38, and 3) did not overlap the palindromic region (where there were substantial rearrangements between the GRCh38-Y and T2T-Y).

#### Mapping and variant calling of the SGDP samples

The SGDP includes 279 open-access, high-coverage genomes from 130 diverse populations^28^. Compared to 1KGP, SGDP includes 118 additional populations with samples sequenced to an average of 43× coverage using a shared PCR-free Illumina library. The SGDP samples were aligned and genotyped to T2T-CHM13+Y and GRCh38 on AnVIL^146^ following the same pipeline as the analysis of 1KGP samples.

#### Curated syntenic region and liftover chains

The initial chain file was generated using nf-LO^148^ with minimap2^91^ alignments. The alignments were filtered and converted to PAF using chaintools. Alignments of nonhomologous chromosomes were removed. Overlapping alignments in the query sequence was removed with rustybam to create 1:1 alignments. PAF alignments were converted back to chain format.

In addition to the minimap2-based whole genome alignment, we applied a wfmash-based pipeline^149^ to validate the chain file. This pipeline starts with a wfmash^149^ whole-genome alignment of T2T-CHM13+Y and the masked and filtered GRCh38 assembly, and identifies 1-to-1 homologous regions at least 5 kb long with a nucleotide identity of at least 95%. Similarly, the resulting chain was post-processed to obtain 1:1 alignments using rustybam and the paf2chain tool. All PAF files with full CIGAR strings were then inspected with SafFire for quality investigation. The minimap2- and wfmash-based chains showed high consistency over the genomes.

### Datasets and resources for T2T-CHM13+Y

#### Lifting over resources from GRCh38 to CHM13+Y

Using the curated chain file, we lifted over dbSNP build 155^150^, the March 13, 2022 release of Clinvar^23, 151^, and GWAS Catalog v1.0^24, 152^ from the GRCh38 primary assembly to T2T- CHM13v2.0 (T2T-CHM13+Y). Liftover was performed as previously described^69^ using GATK Picard^153^ LiftoverVcf and the alignment chain described above.

#### ENCODE

Reads were obtained from the ENCODE dataset^29^ and mapped with Bowtie2^154^. Alignments were filtered using Samtools to remove unmapped or single end mapped reads and those with a mapping quality score <2. PCR duplicates were identified and removed with the Picard tools “mark duplicates”. Alignments were then filtered for the presence of unique k-mers. Bigwig coverage tracks and enrichment tracks were created using deepTools2 bamCoverage^155^.

#### gnomAD

Genome wide variant data from the Genome Aggregation Database (gnomAD) release v3.1.2 was lifted over from GRCh38 to each assembly using CrossMap^156^. The chain files used were created from the GRCh38-based HAL file, downloaded from the cactus-minigraph alignment of Liao *et al.*^77^. The resulting VCFs were annotated with predicted molecular consequence and transcript-specific variant deleteriousness scores from PolyPhen-2 and SIFT using Ensembl Variant Effect Predictor.

### Human Y chromosome contamination in bacterial genomes

#### Screening against Chrisman *et al.* study

We used the MUMmer^157^ to compare 73,691 bacterial 100-mers reported as enriched in human males by Chrisman *et al.*^74^ to the T2T-Y assembly. We found that, as predicted, more than 95% of the 100-mers had near-perfect matches, defined as an exact match of 50 bp or longer, to the complete T2T-Y sequence. The nucmer program from MUMmer was run with default options, except to specify -l 50 for an exact match length of 50 or more, and -c 50 so that it reported matches as short as 50 bp.

#### Screening with 64-mers

Meryl^18^ was used to compare 64-mers from NCBI RefSeq release 213 (July 2022) to T2T-Y and GRCh38-Y. Each bacterial contig was annotated with the number of matching k-mers in T2T-Y, GRCh38-Y, and the number of k-mers in the contig with a match. Each position in the reference chromosomes was annotated with the multiplicity of the k-mer at that position in the RefSeq contigs, and with the number of contigs containing the k-mer. Hits per query were filtered to retain only contigs with more than 20 k-mer matches or with more than 10% of the contig sequence covered by k-mer matches. The queries at each reference position were combined and accumulated into 10 kb windows and converted to an interval wiggle file for visualization. RefSeq sequence entries with hits were retrieved using seqrequester and categorized using 64-mers built from HSat1B and HSat3 annotations. The first and second words in the sequence entry names were extracted to visualize the taxonomic abundance of the microbial genomes in a pie chart using Kronatools^158^ (**Extended Data Fig. 11c)**.

## Data Availability

The T2T-CHM13v2.0 (T2T-CHM13+Y) assembly, reference analysis set, complete list of resources including gene annotation, repeat annotation, epigenetic profiles, variant calling results from 1KGP and SGDP, gnomAD, ClinVar, GWAS, and dbSNP datasets are available for download at https://github.com/marbl/CHM13. The assembly is also available from NCBI and EBI with GenBank accession GCA_009914755.4. Annotation and associated resources are also browsable as “hs1” from the UCSC Genome Browser http://genome.ucsc.edu/cgi-bin/hgTracks?db=hub_3671779_hs1, the Ensembl Genome Browser https://projects.ensembl.org/hprc/ (assembly name T2T-CHM13v2.0) and NCBI data-hub https://www.ncbi.nlm.nih.gov/data-hub/genome/GCF_009914755.1/. Potential assembly issues are listed and tracked at https://github.com/marbl/CHM13-issues. 1KGP and SGDP short read alignments and variant calls are available within AnVIL at https://anvil.terra.bio/#workspaces/anvil-datastorage/AnVIL_T2T_CHRY. Sequencing data used in this study is listed in **Supplementary Table 1**.

## Code Availability

Custom codes developed for data analysis and visualization are available at https://github.com/arangrhie/T2T-HG002Y, https://github.com/snurk/sg_sandbox, and https://github.com/schatzlab/t2t-chm13-chry and deposited on Zenodo along with https://github.com/marbl/CHM13 and https://github.com/marbl/CHM13-issues160. Software and parameters used are stated in the **Supplementary Methods** with more details.

## Acknowledgements

We thank P. Hallast, M. C. Loftus, M. K. Konkel, P. Ebert, T. Marschall, and C. Lee for coordination and discussions, J.C.-I. Lee for sharing the GRCh38-Y coordinates used in Y-Finder, and members of the Telomere-to-Telomere consortium and Human Pangenome Reference Consortium for constructive feedback. This work utilized the computational resources of the NIH HPC Biowulf cluster (https://hpc.nih.gov). Computational resources were partially provided by the e-INFRA CZ project (ID:90140), supported by the Ministry of Education, Youth and Sports of the Czech Republic and Computational Biology Core, Institute for Systems Genomics, University of Connecticut. Certain commercial equipment, instruments, or materials are identified to specify adequately experimental conditions or reported results. Such identification does not imply recommendation or endorsement by the National Institute of Standards and Technology, nor does it imply that the equipment, instruments, or materials identified are necessarily the best available for the purpose.

## Author contributions

Assembly: S.N., S.K., M.R. Validation: A.R., S.K., M.A., A.V.B., G.F., A.F., A.M.M., J.M., A.M., L.F.P., D.P., F.J.S., K.S., P.M., J.M.Z., K.D.M. ChrY haplogroups: A.R., A.C.W. Alignment: C.-S.C., M.D., R.Har., M.R.V., K.D.M. Satellite annotation: N.A., I.A.A., G.A.L., F.R., V.A.S., K.H.M. FISH: N.A., J.G., T.P. Repeat annotation: S.J.H., P.G.S.G., G.A.H., R.Hu., J.M.S., R.O. Retroelements: R.Hal., W.M. Non-B DNA: M.H.W., K.D.M. Gene annotation: A.R., M.D., P.F., C.G.G., L.H., M.H., J.H., T.H., F.J.M., T.D.M., S.L.S., A.S., F.T.-N. Ampliconic genes: A.R., R.Har., W.T.H., P.M., M.T., K.D.M. Structural annotation: A.R., M.C., H.L., P.M., K.D.M. Epigenetics: A.R., P.W.H., A.Ge., W.T., A.M.W. Mappability: A.M.T.O., M.A.W., J.M.Z. Non-B DNA: M.H.W., K.D.M. Variants and Liftover: A.R., D.J.T., S.K., J.A., N.-C.C., M.D., E.G., A.Gu., N.F.H., W.T.H., S.E.H., S.H., R.C.M., N.D.O., M.E.G.S., L.S., M.R.V., S.Z., J.M.Z., E.E.E., A.M.P. Contamination: A.R., S.L.S., B.P.W., A.M.P. Data generation: M.J., R.K.K., A.P.L., J.K.L., C.M., B.M.M., K.M.M., H.E.O., F.J.S., Y.Z. Data management: A.R., M.D., M.J., J.K.L. Computational resources: R.O., M.C.S., A.M.P. Manuscript draft: A.R., S.N., M.C., S.J.H., D.J.T., N.A., I.A.A., N.-C.C., E.G., J.G., P.G.S.G., A.Gu., R.Hal., W.M., J.M., T.P., F.R., S.L.S., J.M.S., A.M.T.O., A.C.W., M.A.W., S.Z., J.M.Z., E.E.E., R.O., M.C.S., K.H.M., K.D.M., A.M.P. Editing: A.R., A.M.P., with the assistance of all authors. Supervision: J.M.Z., E.E.E., R.O., M.C.S., K.H.M., K.D.M., A.M.P. Conceptualization: A.R., S.N., M.C., E.E.E., K.H.M., K.D.M., A.M.P.

## Additional information

**SupplementaryTables.xlsx** file contains **Table 1** and **Supplementary Tables 1-32. SupplementaryInformation.pdf** file contains **Supplementary Methods, Supplementary Figs. 1-19**, and **Supplementary Notes 1-4**. **SupplementaryFiles.zip** contains **Supplementary Files 1-3**.

## Extended Data Figures

**Extended Data Fig. 1.**
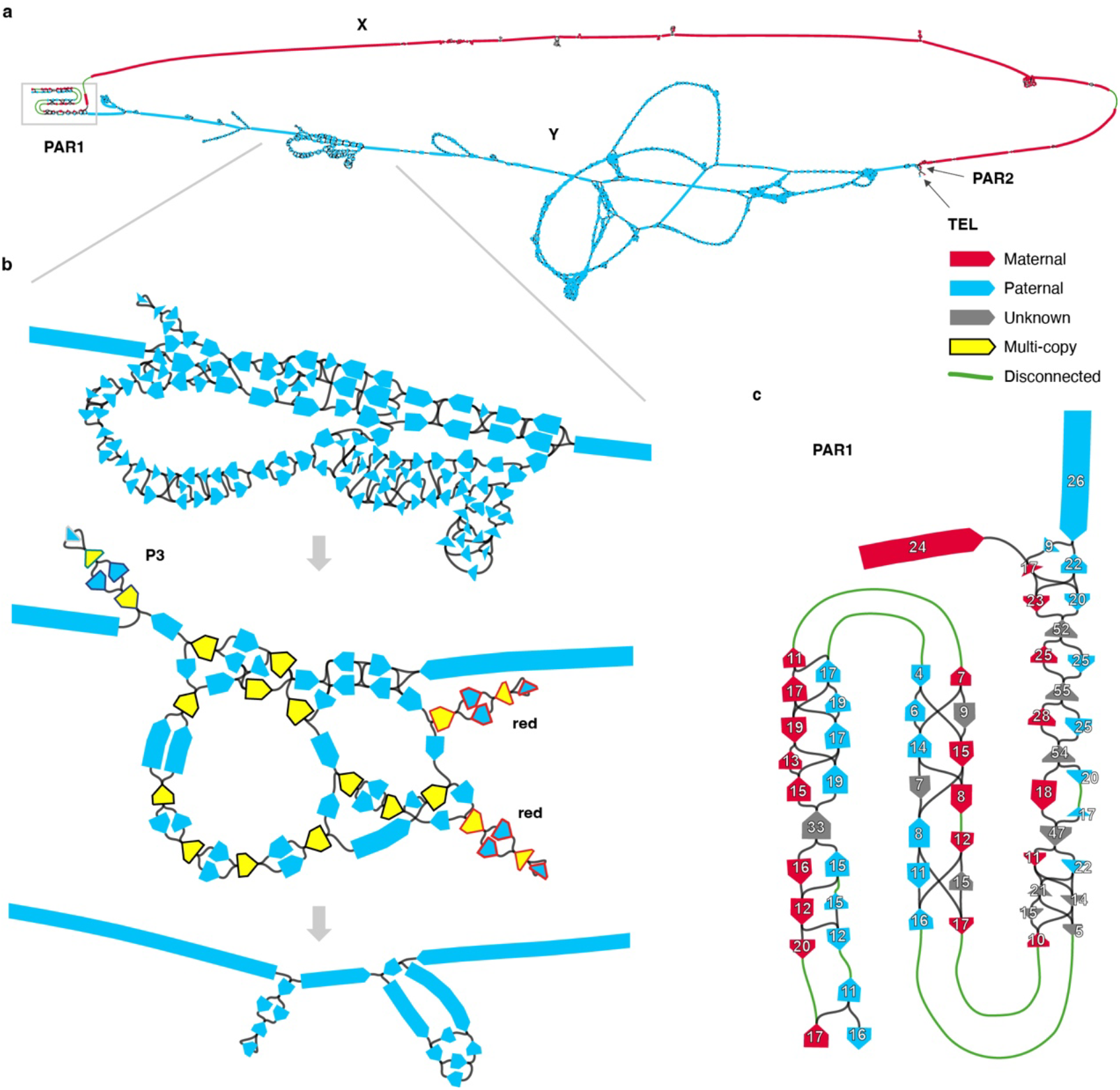
Assembling the X and Y chromosomes of HG002. **a**. Chromosome X and Y components of the assembly string graph built from HiFi reads, detected based on node sequence alignments to T2T-CHM13 and GRCh38 references. Each node is colored according to the excess of paternal-specific (blue) and maternal-specific (red) k-mers, obtained from parental Illumina reads, indicating if they exclusively belong to chromosome Y or X, respectively. Most complicated tangles are localized within the heterochromatic satellite region on the Y q-arm. The X and Y subgraphs are connected in PAR1 and PAR2. Graph discontinuities are due to a lack of HiFi sequence coverage in these regions caused by contextual sequencing bias, with 9 out of 11 observed breaks falling within PAR1 on either chromosome (5 out of 5 for chromosome Y). Note that for visualization purposes the length of shorter nodes is artificially increased making the extent of the tangles appear larger than reality. **b**. The effects of manual pruning and semi-automated ONT read integration is illustrated from top to bottom. Top, zoomed in view of a tangle encoding the P1–P3 palindromic region in Y (approx. 22.86–27.08 Mb, see **Fig. 4**). Middle, corresponding subgraph following the manual pruning and recompaction. Nodes excluded from the curated “single-copy” list for automated ONT-based repeat resolution are shown in yellow. Three hairpin structures are highlighted, which form almost-perfect inverted tandem repeats encompassing the entire P3 and two P2 (red) palindromes. Node outlines in the palindromes are colored according to the palindromic arms as in **Fig. 4**. Bottom, corresponding subgraph following the repeat resolution using ONT read-to-graph alignments. Remaining ambiguities were resolved by evaluating ONT read alignments to all candidate reconstructions of the corresponding sub-regions. **c**. PAR1 subgraph labeled with HiFi read coverage on each node. Gaps (green edges) and uneven node coverage estimates indicate biases in HiFi sequencing across the region. **Fig. 1** shows an enrichment of SINE repeats and non-B DNA motifs in PAR1 that may underlie the sequencing gaps in this region.

**Extended Data Fig. 2.**
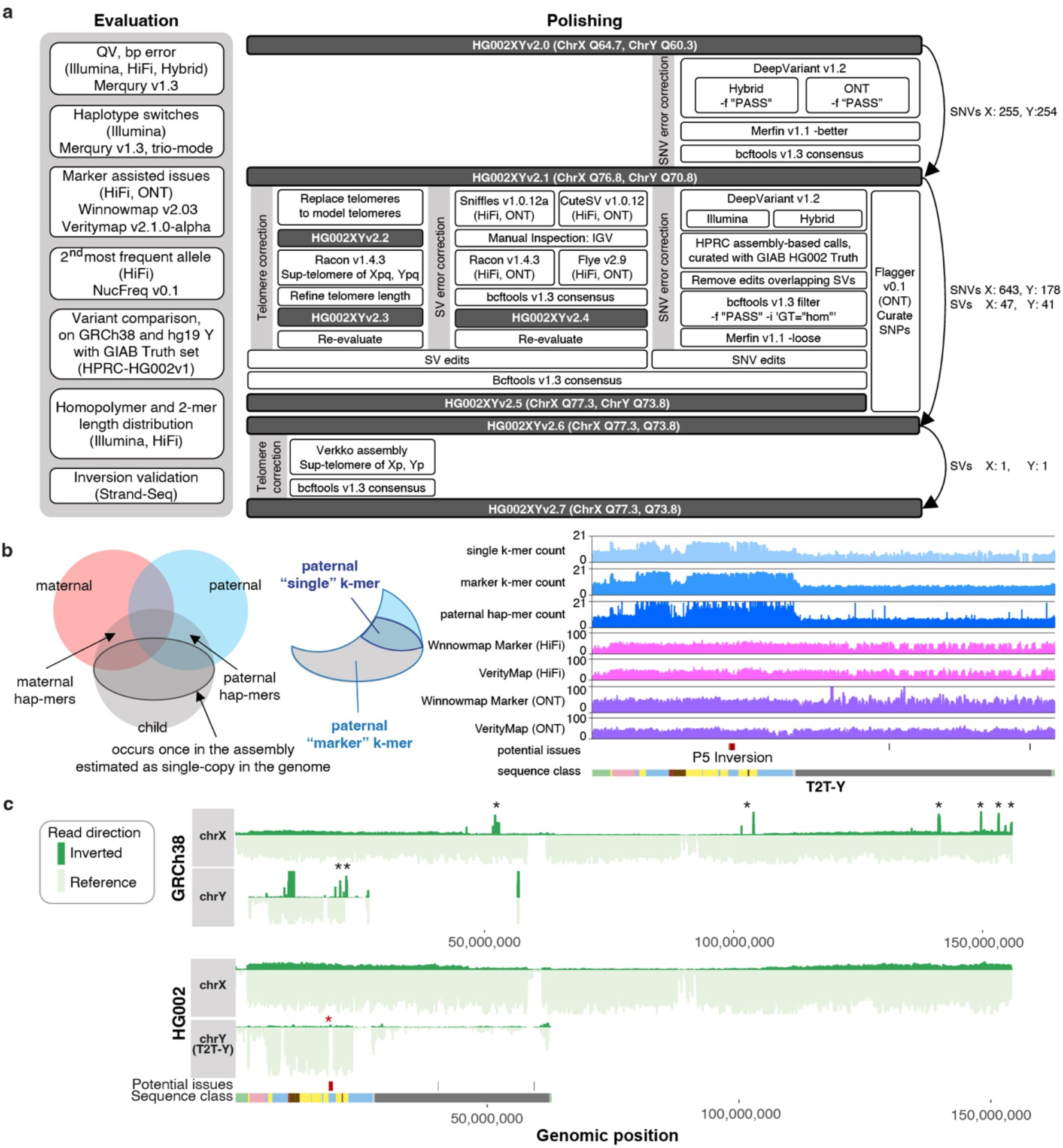
Validation and polishing of the T2T-Y. **a.** Evaluation and polishing workflow performed on T2T-CHM13v1.1 autosomes + HG002 XY assemblies. **b**. Venn diagram of the k-mers from the parents and child. On the left, hap-mers^18^ represent haplotype specific k-mers inherited by the child. The darker outlined circle inside the child k-mers represent single-copy k-mers (k-mers occurring once in the assembly and single-copy in the child’s genome). Right figure shows an example of the paternal specific, “single-copy” and “marker” k-mers. The marker set includes both multi-copy and single-copy k-mers specific to the paternal haplotype that were inherited by the child. Unlike polishing the nearly haploid CHM13 assembly^17^, both single-copy k-mers and marker k-mers were used for the marker-assisted alignments to HG002 XY. This helped align more reads within repetitive regions to the correct chromosome for evaluation during polishing. Right panel shows counts of the k-mers and coverage of HiFi and ONT reads using the marker-assisted Winnowmap2 alignment, in addition to alignments from VerityMap, which uses locally unique k-mers for anchoring the reads. **c.** Aggregated Strand-seq coverage profile across all 65 libraries on GRCh38-Y (top) and T2T- Y (bottom). Each bar represents read counts in every 20 kb bin supporting the reference in forward direction (light green) or reverse direction (dark green). Multiple spikes in reverse direction (black asterisks) in GRCh38-Y indicate inversion polymorphisms relative to HG002, likely due to differences between the haplogroups. Such spikes in coverage are not observed on T2T-X and T2T-Y, which confirm the structural and directional accuracy of the HG002 assemblies. A 3 kb inversion of the unique sequence between the P5 palindromic arms was identified as erroneous in T2T-Y (red asterisk), but was confirmed to be polymorphic in the population and left uncorrected in this version of the assembly.

**Extended Data Fig. 3.**
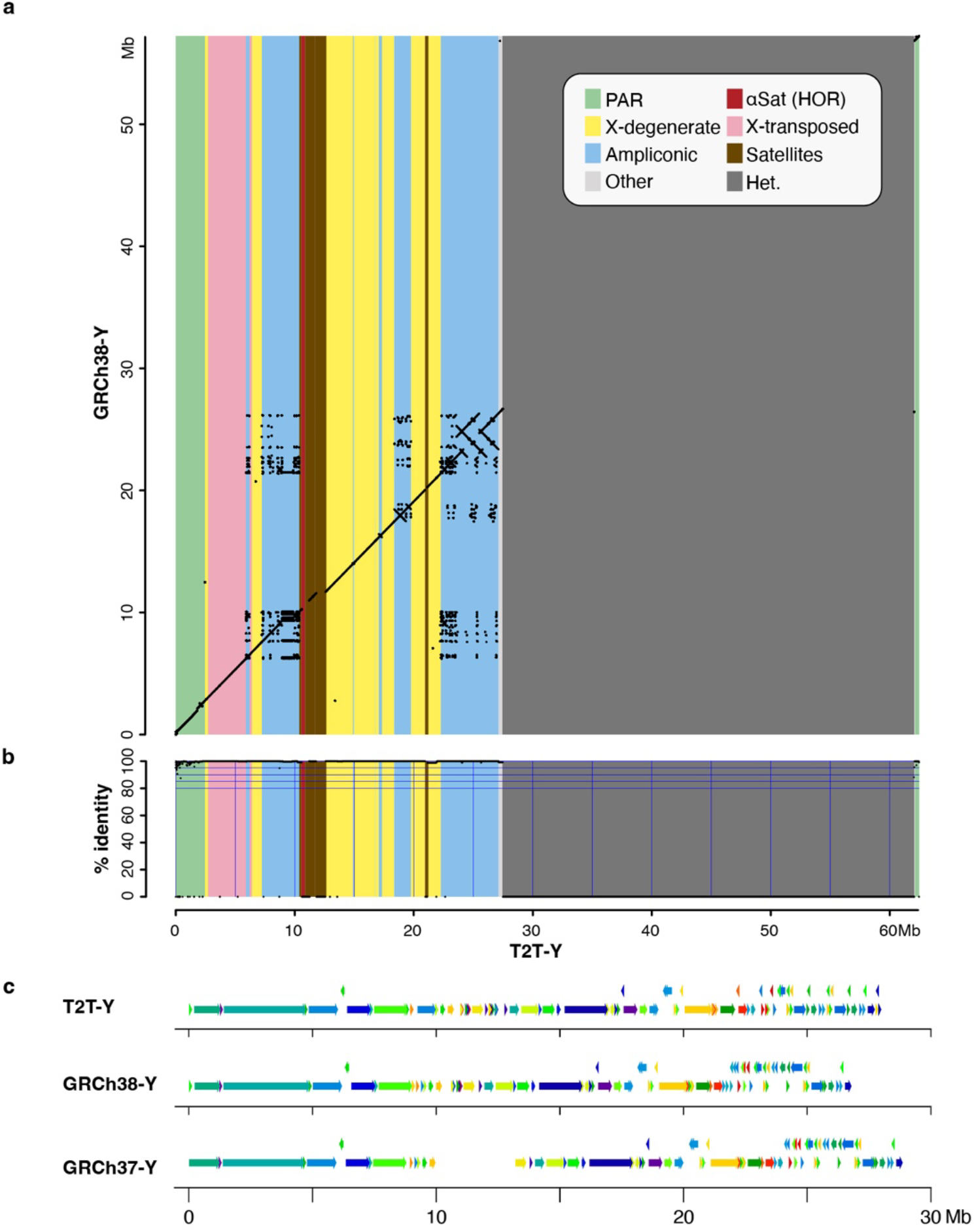
Large structural differences between T2T-Y and previous GRCh Y assemblies. **a-b.** Ampliconic genes and X-degenerate sequences revealed from alignments between GRCh38-Y (Y-axis) and T2T-Y (X-axis). **a.** Dotplot generated using LastZ^93^ after softmasking with WindowMasker^94^. **b.** Identity was computed from matches and mismatches over positions with alignments, excluding gaps. **c.** Structural differences revealed using PRG-TK^95^ against GRCh38-Y and GRCh37-Y in the euchromatic region of the Y chromosome.

**Extended Data Fig. 4.**
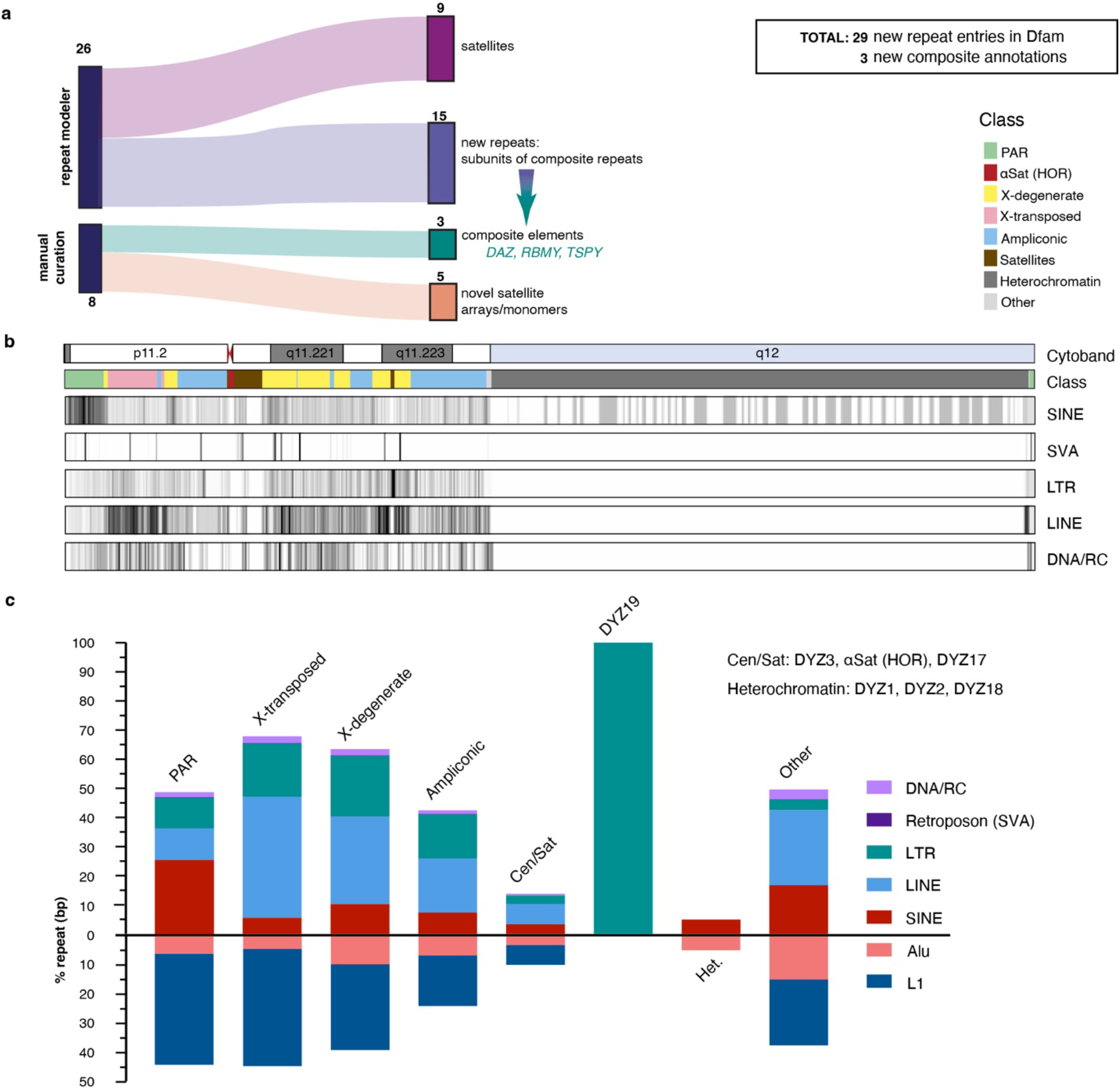
Repeat discovery and annotation of T2T-Y. **a.** Assembly completion allowed for a full assessment of repeats and resulted in the identification of previously unknown satellite arrays (predominantly in the PAR1) and subunit repeats that fall within one of three composite repeat units (*TSPY*, *RBMY*, *DAZ*). **b.** Ideogram of TE density (per 100 kb bin). This is an extension of **Fig**. **1** with non-SINEs expanded into separate TE classes (SVA, LTR, LINE, DNA/RC). Density scale ranges from low (white, zero) to high (black, relative to total density) and sequence classes are denoted by color. **c.** Summary (in terms of base coverage per region) across all five TE classes and two specific families: *Alu*/SINE and L1/LINE. The satellites in (**b**) were kept separate as two categories; Cen/Sat as the left satellite block including alpha satellites and DYZ19, while all other categories were combined per sequence classes.

**Extended Data Fig. 5.**
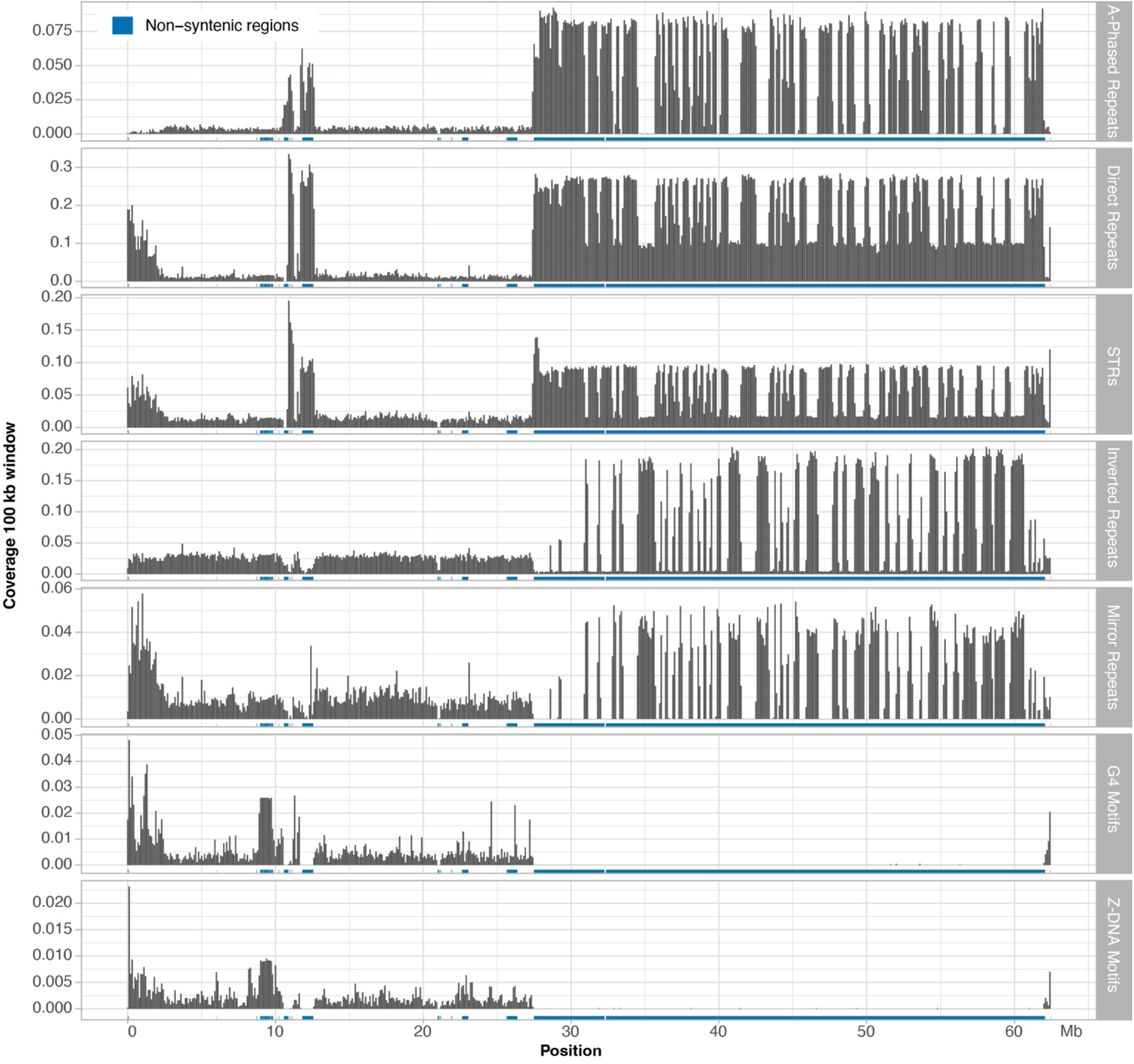
Non-B DNA motifs along the T2T-Y. HSat3 on the Yq and satellite sequences around the centromere are more enriched with A-phased repeats, direct repeats and STRs, while HSat1B is more enriched with inverted repeats and mirror repeats. Enrichment of non-B DNA sequences were also observed in the PAR region. Notably, the *TSPY* gene array is enriched for G4 and Z-DNA motifs, as shown in **Extended Data Fig. 6b**.

**Extended Data Fig. 6.**
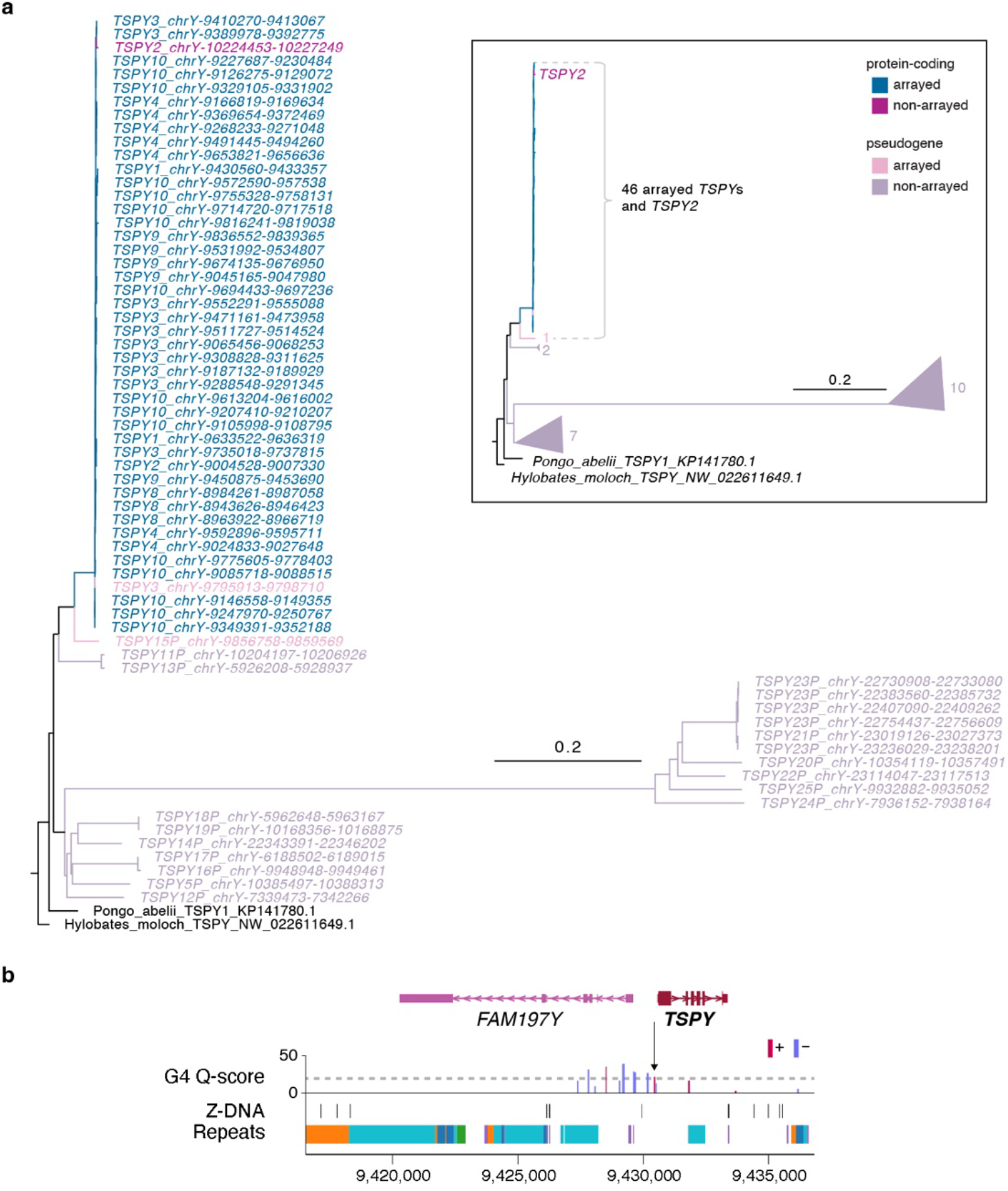
Phylogenetic tree analysis of the ampliconic *TSPY* gene family and pattern of non-B DNA structure. **a.** Phylogenetic tree analysis using protein-coding *TSPY*s from a Sumatran Orangutan (*Pongo abelii*) and a Silvery gibbon (*Hylobates moloch*) as outgroups confirmed *TSPY2* (distal to the array) and *TSPY* copies within the array originated from the same branch, distinguished from the rest of the *TSPY* pseudogenes. Rectangular inset shows a cartoon representation of the simplified tree. Numbers next to the triangles indicate the number of *TSPY* genes in the same branch. **b.** G4 and Z-DNA structures predicted for a typical *TSPY* copy inside the *TSPY* array. All *TSPY* copies in the array have the same signature, with one G4 peak present ∼500 bases upstream of the *TSPY* (arrow). Higher Quadron score^122^ (Q-score) indicates a more stable G4 structure, with scores over 19 considered stable (dotted line).

**Extended Data Fig. 7.**
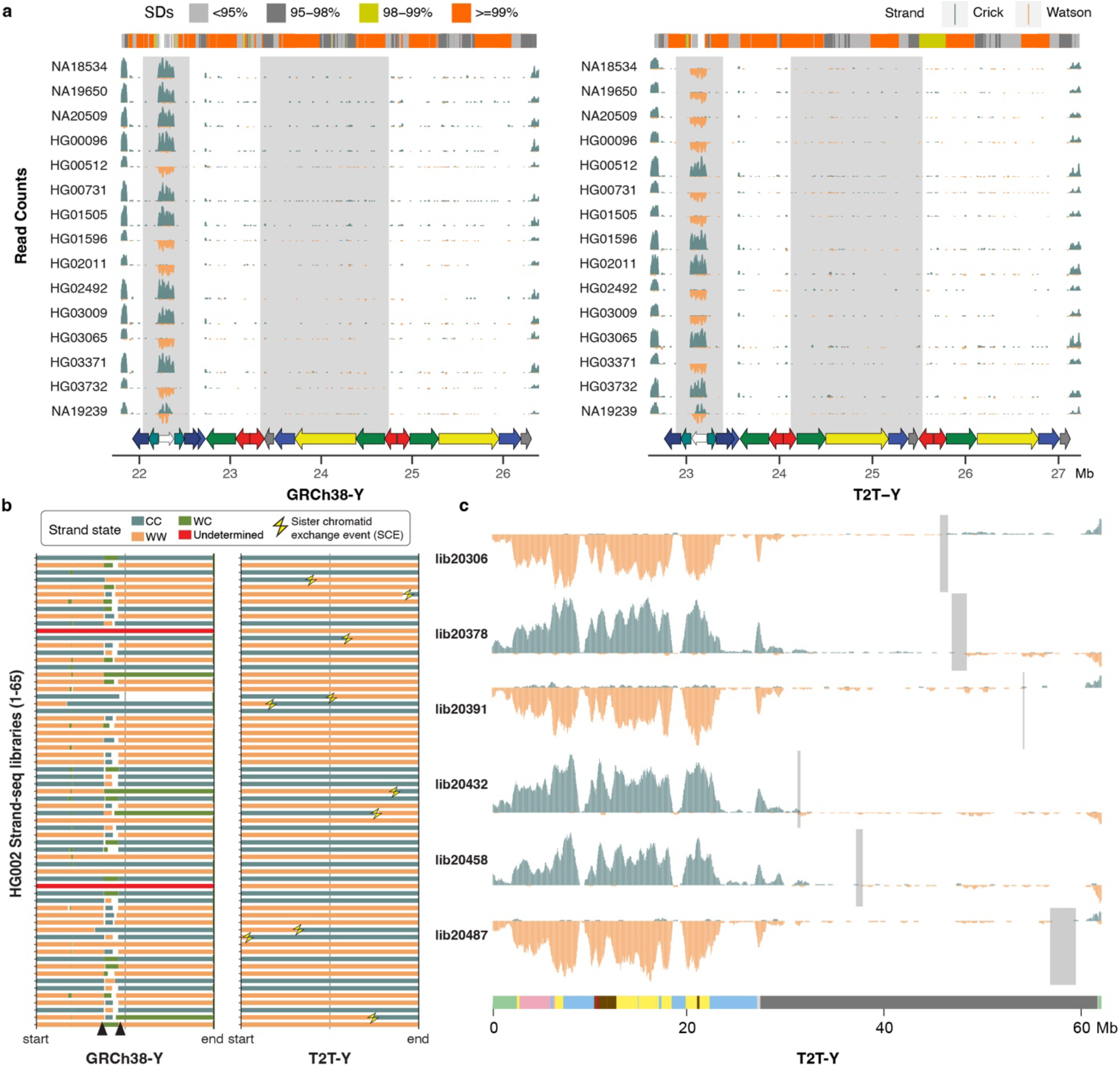
Recurrent inversions identified with Strand-seq. **a.** Five out of 15 individuals have the inverted variant as present in HG002 at the P3 palindrome (white arrow). Although inversions across P1–P2 (yellow and red arrows) are difficult to confirm with Strand-seq because of the high sequence similarity between the palindromic arms, different orientations are observable in these samples. **b.** Strand states for 65 Strand-seq libraries of HG002. Depending on the mappings of directional Strand-seq reads (+ reads: ‘Crick’, C, - reads: ‘Watson’, W), reference sequence was assigned in three states: WC, WW, and CC. WC, roughly equal mixture of plus and minus reads; WW, all reads mapped in minus orientation; CC, all reads mapped in plus orientation. Changes in strand state along a single chromosome are normally caused by a double-strand-break (DSBs) that occurred during DNA replication^159^ in a random fashion and we refer to them as sister-chromatid-exchanges (SCEs, yellow thunderbolts). Recurrent change in strand state over the same region in multiple Strand-seq cells indicates misassembly. Similarly, collapsed or incomplete assembly of a certain genomic region will result in a recurrent strand state change as observed for GRCh38-Y (black arrowheads). In contrast, T2T-Y shows strand state changes randomly distributed along each Strand-seq library with no evidence of misassembly or collapse. **c.** Strand-seq profile of selected libraries over T2T-Y summarized in bins (bin size: 500 kb, step size: 50 kb). Teal, Crick read counts; orange, Watson read counts. As ChrY is haploid, reads are expected to map only in Watson or Crick orientation. Light gray rectangles highlight regions where SCEs were detected in the heterochromatic Yq12 despite a lower coverage of Strand-seq reads. A modified breakpointR parameter was used (windowsize = 500000 minReads = 20) in order to refine detected SCEs presented in panel **b** and **c**.

**Extended Data Fig. 8.**
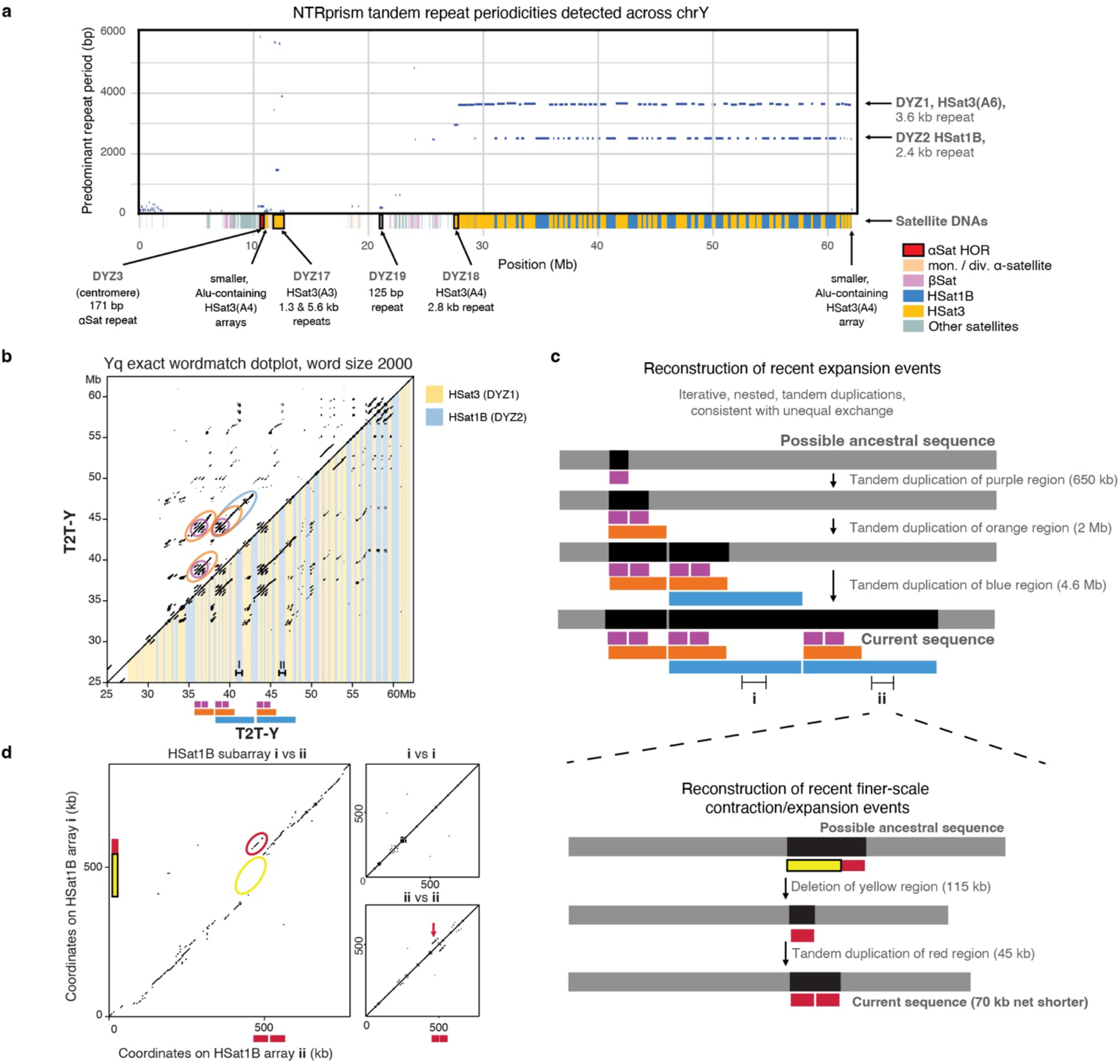
Satellite annotation and recent expansion events in the Yq heterochromatin. **a.** A plot showing the top repeat periodicities detected by NTRprism^46^ in 50 kb blocks tiled across T2T-Y, with centromeric satellite annotations overlaid on the X axis. Large arrays are labeled with their historic nomenclature^1^, HSat subfamilies^64^, and predominant repeat periodicities. **b.** An exact 2000-mer match dotplot of the Yq region (a dot is plotted when an identical 2000 base sequence is found at positions X and Y). The lower triangle has DYZ1/DYZ2 annotations overlaid as yellow and blue bars, respectively. Circled patterns in the upper triangle correspond to recent iterative duplication events, which are illustrated below the X axis. **c.** A reconstruction of a possible sequence of recent iterative duplications that could explain the observed dotplot patterns. **d**. A 2000-mer dotplot comparison of two ∼800 kb HSat1B sub-arrays that were part of a recent large duplication event, along with self-self comparisons of the same arrays, revealing sites of more recent and smaller-scale deletions and expansions (annotated in yellow and red, with a possible sequence of events illustrated by the schematic on the right).

**Extended Data Fig. 9.**
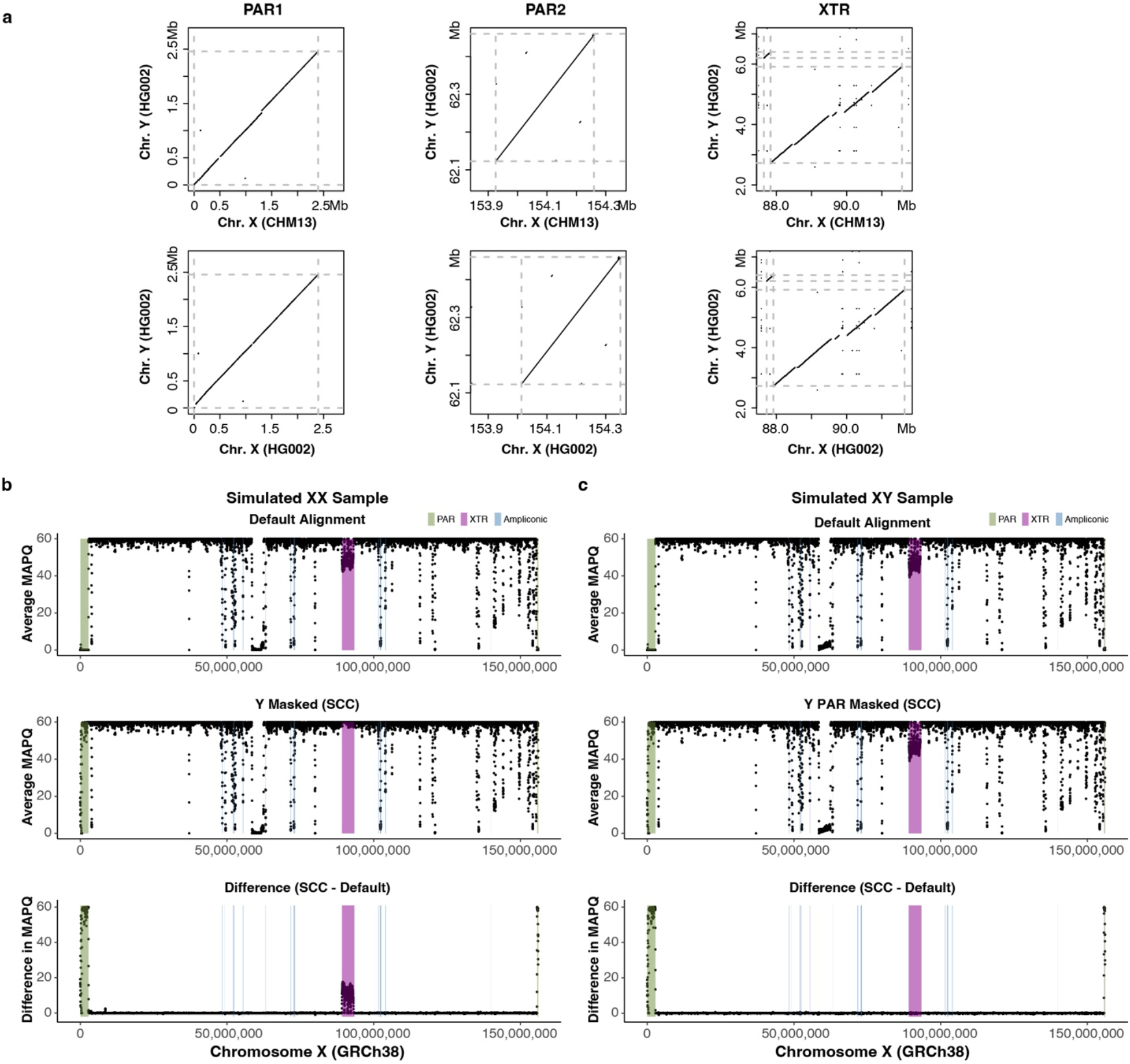
Genomic similarity in PARs and XTR and improved MAPQ of the PARs through informed sex chromosome complement reference. **a.** Dotplots from LASTZ alignments of the CHM13-X, HG002-X, and HG002-Y (T2T-Y) over 96% sequence identity. Dashed gray lines represent the start and end of the approximate PARs or XTR boundaries. Disconnected diagonal lines indicate the presence of genomic diversity between each paired region. More genomic differences are observed in the PAR1 between the HG002-Y and CHM13-X. **b-c.** Average mapping quality (MAPQ) across GRCh38-X from simulated reads of an XX **(b)** and XY **(c)** sample. Top, a default version of GRCh38 (with two copies of identical PARs on XY). Middle, a version of GRCh38 informed on the sex chromosome complement (SCC) of the sample (entire Y hard-masked for the XX sample vs. only PARs on the Y hard-masked for the XY sample). Bottom, the difference in average MAPQ between the SCC and default approaches. MAPQ was averaged in 50 kb windows, sliding 10 kb across the chromosome. A positive value means MAPQ score is higher with SCC reference alignment compared to default alignment.

**Extended Data Fig. 10.**
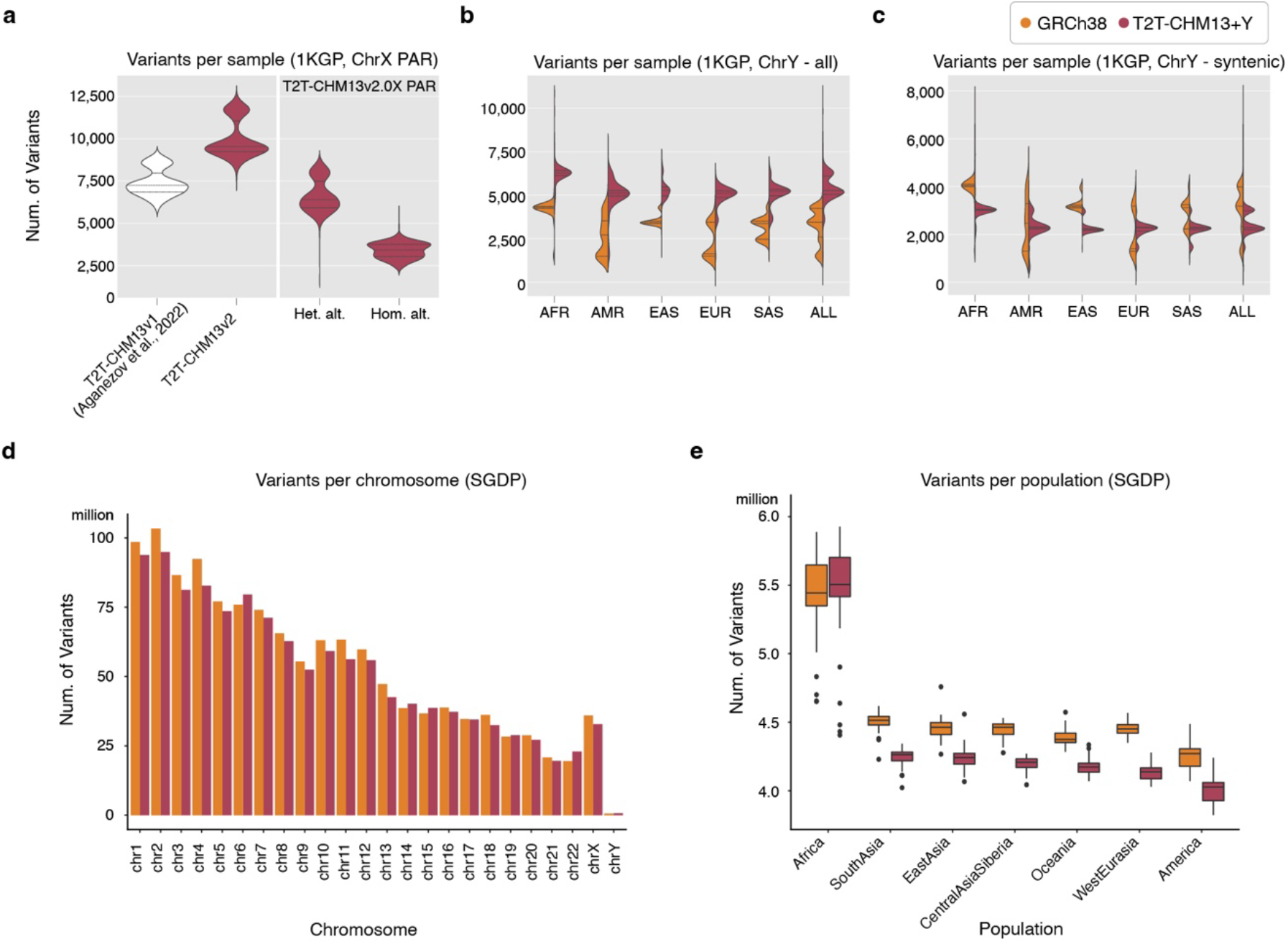
Number of variants called from 1KGP and SGDP individuals. **a.** More variants are called on the X-PARs when using the sex chromosome complement reference approach (calling variants in diploid mode on PARs) than the non-masked approach (calling variants in haploid mode on PARs). The 1KGP results for GRCh38-Y are from Aganezov et al.^69^, which was performed on CHM13v1.0+GRCh38-Y. **b.** Num. of variants called from each 1KGP XY sample on chromosome GRCh38- Y and T2T-Y **c.** Num. of variants called in the syntenic region between the two Ys. A large num. of additional variants are called on each sample attributed to the newly added, non-syntenic sequences on T2T-Y. Within the syntenic regions, a reduction in the number of variants is observed for each population except for samples from R1 haplogroups as shown in **Fig. 6c**. **d.** Aggregated total number of variants for the 279 SGDP samples per chromosome. **e.** SGDP genome-wide counts of variants per-sample (n=279) demonstrate increased variation in African samples regardless of reference. Each bar in the box plot represents the 1st, 2nd (median), and 3rd quartile of the number of variants in each population. Whiskers are bound to the 1.5 × interquartile range. Data outside of the whisker ranges are shown as dots. For the SGDP samples, variants were called using T2T-CHM13+Y or GRCh38 as the reference. All variants shown in this figure were filtered for “high quality (PASS)”.

**Extended Data Fig. 11.**
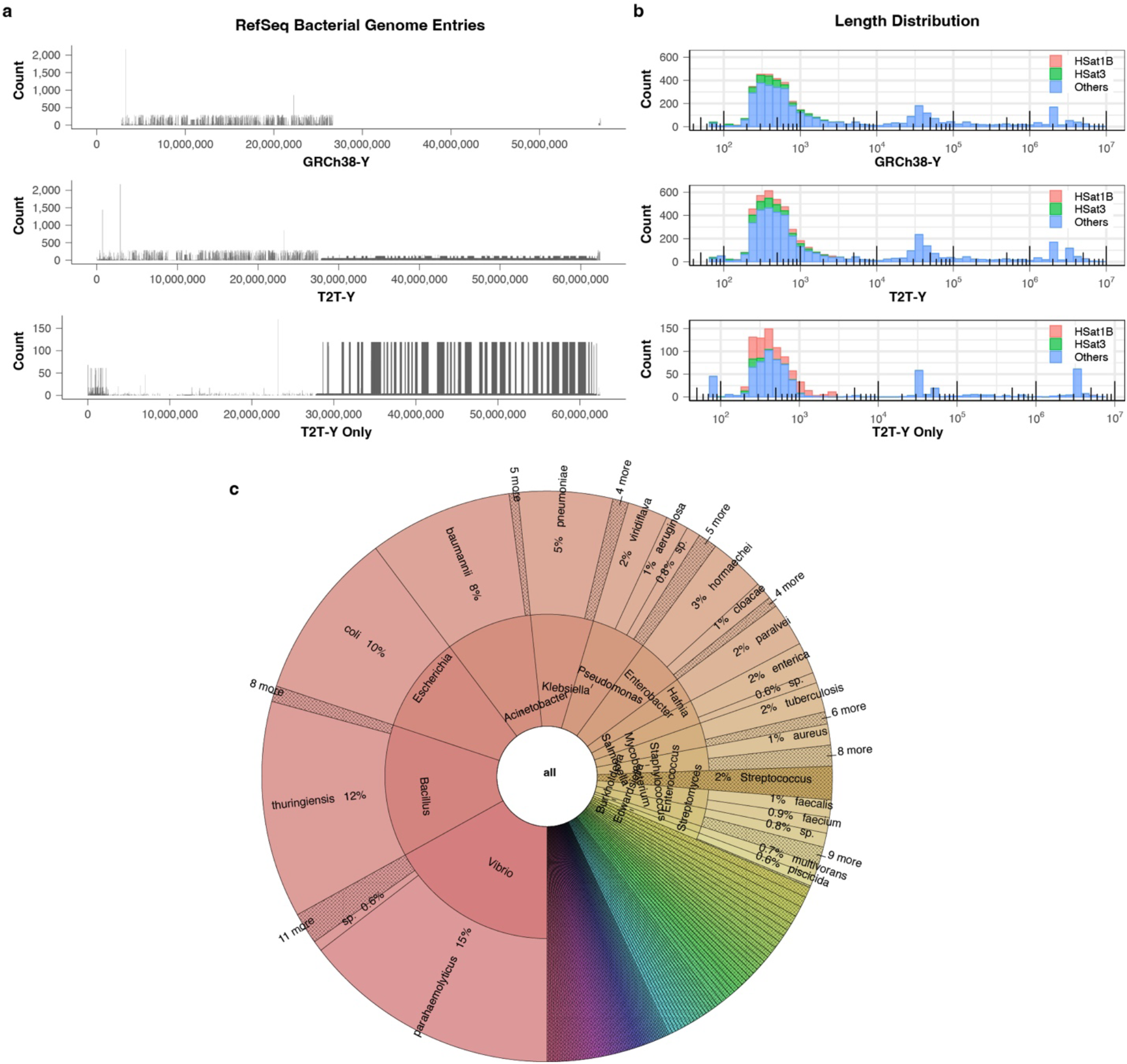
Human contaminants in bacterial reference genomes. **a.** Number of distinct RefSeq accessions in every 10 kb window containing 64-mers of GRCh38-Y (top), T2T-Y (middle), and in T2T-Y only (bottom). Here, RefSeq sequences with more than 20 64-mers or matching over 10% of the Y chromosome are included. **b.** Length distribution of the sequences from (**a**) in log scale. Majority of the shorter (<1 kb) sequences contain 64-mers found in HSat1B or HSat3. **c.** Number of bacterial RefSeq entries by strain identified to contain sequences of T2T-Y and not GRCh38-Y, visualized with Krona^158^.

