## Supplementary Methods and Notes for "The complete sequence of a human Y chromosome"

Rhie *et al*., 2023

**Supplementary Methods** **4**
Sequencing and Data Generation 4
PacBio HiFi whole-genome sequencing 4
At Pacific Biosciences 4
At University of Washington 4
At Washington University in St. Louis 5
ONT whole-genome sequencing 5
Pacbio Iso-Seq cDNA sequencing 6
Assembly 8
Graph construction and pruning 8
Using ONT read alignments for tangle resolution 10
Using parent-specific markers to inform PAR1 reconstruction 11
Gap patching and contig stitching 12
Error detection and polishing 12
Sequencing read alignment 13
Primary alignments 13
Marker assisted alignments 13
Polishing 15
Small (SNV) error correction on v2.0 16
Telomere correction on HG002v2.1 19
Large (SV) error correction on HG002v2.1 20
Variants called from HPRC assembly with respect to GRCh38 and GRCh37 23
Homopolymer and 2-mer evaluation 23
Less conservative filtering with Merfin 24
HPRC Flagger pipeline on v2.1 25
Telomere correction on v2.6 26
Evaluation 26
Merqury 26
Coverage analysis 27
VerityMap 27
Second-most frequent allele with NucFreq 30
Strand-seq evaluation 30

Integrity with HG003Y 30
Chromosome spreads and Fluorescent In-Situ Hybridization (FISH) 32
Comparison to GRCh38Y 33
Y haplogroup identification 33
Alignments between GRCh38 and HG002 Y assemblies 33
SafFire 33
LASTZ 33
Pangenomics Research Tool Kit 34
Gene annotation 34
CAT and Liftoff gene annotation 34
RefSeq gene annotation 35
RefSeq Liftoff gene annotation 35
Ensembl gene annotation 36
Iso-Seq analysis 37
Ampliconic gene copy number validation 39
Repeat annotation 40
Segmental duplications 40
Repeat model discovery and annotation 40
Repeat model discovery with RepeatModeler and loci identification with RepeatMasker 40
Discovery of new satellite models 40
Manual curation of previously unknown repeat models 41
Compilation and polishing of T2T-Y repeat annotations 42
Composite repeats 42
Identification of full-length TEs 43
T2T-Y liftOver analysis 43
Satellite annotation 44
Cytoband annotation 45
Transduction analysis 45
Non-B DNA motif annotation 46
Data visualization 46
TSPY gene family analysis 46
TSPY copy number estimation from SGDP 46
Phylogenetic tree analysis of the TSPY genes 47
Centromere analysis 47
HOR haplotype and SVs 47
HG002 cenY analysis 47
RP11 cenY analysis 51
CENP-A 54
Stained glass plot of the DYZ3 array 55
Epigenetic profile 55

ONT NanoNOMe sequencing data 55
ONT NanoNOMe alignments 55
ONT Nanopolish CpG and GpC methylation calling 56
ONT Remora CpG methylation calling and processing 57
PacBio methylation calling 58
Whole genome bisulfite sequencing (WGBS) and Enzymatic Methyl-seq (EM-seq) processing 58
Analysis of methylation detection technologies 58
Centromeric Dip Region (CDR) 59
Sequence classes on the Y chromosome 61
Palindrome structure, P1-P3 61
Palindrome structure, P4-P8 61
Azoospermia Factor (AZF) region 61
Pseudoautosomal region (PAR) and X-transposed region (XTR) 62
Yqh heterochromatin region 64
Yqh DYZ1/DYZ2 64
Phylogenetic analyses of AluY repeats 65
Short-read variant calling on T2T-CHM13+Y 65
Impact of masking PAR and XTR in variant calling 65
Mappability comparison and variant calling in 1KGP samples 67
Putative collapsed regions in GRCh38-Y 68
Mapping and variant calling of the SGDP samples 69
Curated syntenic region and liftover chains 71
GRCh38 pre-processing 71
minimap2-based pipeline 71
wfmash-based pipeline 72
Structurally variable region 72
Variants on GRCh38 that will disappear when calling on T2T-CHM13v2.0 72
Datasets and resources for T2T-CHM13v2.0 72
Lifting over resources from GRCh38 to CHM13v2.0 72
ENCODE 73
gnomAD 74
Human Y chromosome contamination in bacterial genomes 74
Screening against Chrisman et al. study 74
Screening with 64-mers 74
**Supplementary Notes 77**
1. Transduced genomic segments 77
2. Non-B DNA motifs 78
3. PAR and XTR masking for sex chromosome complement analysis 78
4. Different CpG Methylation pattern in HiFi and ONT at Yq12 79
**References 81**

### Supplementary Methods

#### Sequencing and Data Generation

##### PacBio HiFi whole-genome sequencing

###### At Pacific Biosciences

DNA was sheared to 20 kb with a Megaruptor 3. Libraries were prepared with SMRTbell Express Template Prep Kit 2.0 and size-selected with SageELF to the target size (15 kb, 19 kb, 20 kb). Libraries were sequenced on the Sequel II System with Chemistry 2.0.

###### At University of Washington

At all steps, DNA quantity was checked with fluorometry on the DS-11 FX instrument (DeNovix) with the Qubit dsDNA HS Assay Kit (Thermo Fisher) and sizes were examined on FEMTO Pulse (Agilent Technologies) using the Genomic DNA 165 kb Kit. Libraries were all prepared for sequencing according to the protocol 'Procedure & Checklist – Preparing HiFi SMRTbell Libraries using the SMRTbell Express Template Prep Kit 2.0' and loaded on Sequel II instruments (PacBio).

**Run m64076_201016_191536.ccs and m64076_201013_225902.ccs**

DNA was isolated from Coriell cell line NA24385 using a modified Gentra Puregene method and sheared using gTUBE (Covaris, Inc.) to 25 kb mode size. After SMRTbell generation, material was size-selected on a SageELF system (Sage Science) using the "0.75% 1-18kb" program and fraction 3 (mode size 20 kb) supplemented with fraction 4 (mode size 28 kb) selected for sequencing. The pooled library was bound with Sequencing Primer v2 and Sequel II Polymerase v2.0 and sequenced on two SMRT Cells 8M using Sequencing Plate v2.0, diffusion loading, four-hour pre-extension, and 30 hour movie times.

**Run m54329U_201103_231616.ccs**

DNA was obtained from PacBio and sheared using Megaruptor 3 (Diagenode, Inc.) twice using settings 30 and 31 to a mode size of 25 kb. After SMRTbell generation, material was size-selected on a SageELF system (Sage Science) using the "Waveform 200" program and fraction 3 with a mode size of 19.5 kb was selected for sequencing. The library was bound with Sequencing Primer v2 and Sequel II Polymerase v2.0 and sequenced on one SMRT Cell 8M using Sequencing Plate v2.0, diffusion loading, two-hour pre-extension, and 30 hour movie time.

**Run m64076_210309_014547**

DNA was obtained from PacBio and sheared using Megaruptor 3 (Diagenode, Inc.) twice using settings 30 and 31 to a mode size of 21 kb. After SMRTbell generation, material was size-selected on a PippinHT system (Sage Science) using the "0.75%, DF 6-10kb high-pass 75E" high-pass program with a 10 kb lower cut. Recovered library with a mode size of 20 kb was bound with

Sequencing Primer v5 and Sequel II Polymerase v2.2 and sequenced on one SMRT Cell 8M using Sequencing Plate v2.0, predictive loading target 0.85, no pre-extension, and 30 hour movie time.

**Run m64076_210310_104300**

HiFi SMRTbell library was prepared at PacBio following protocol 'Procedure & Checklist – Preparing HiFi SMRTbell Libraries using the SMRTbell Express Template Prep Kit 2.0'. Genomic DNA was fragmented with a mode of ~18 kb using the Diagenode Megaruptor 3. Library was size selected using Sage Science Pippin HT. Polymerase-bound SMRTbell complexes were generated using Sequencing Primer v5 and Sequel II Polymerase v2.2. Library was sequenced on a PacBio Sequel II with no pre-extension, 30 hour movies, and a 0.85 predictive loading target using Sequel II sequencing Plate 2.0.

##### At Washington University in St. Louis

**Run m64043_210311_174418**

HiFi SMRTbell library was prepared following protocol 'Procedure & Checklist – Preparing HiFi SMRTbell Libraries the SMRTbell Express Template Prep Kit 2.0'. Genomic DNA was fragmented with a mode of ~18 kb using the Diagenode Megaruptor 1. Library was size selected using Sage Science ELF 1-18 kb cassette. ELF fractions 2 and 3 were combined for sequencing. Polymerase bound SMRTbell complexes were generated using Sequencing Primer v5 and Sequel II Polymerase v2.2. Library was sequenced on a PacBio Sequel II with no pre-extension, 30 hour movies, and a 0.85 predictive loading target using Sequel II sequencing Plate 2.0.

**Run m64043_210313_013127**

HiFi SMRTbell library was prepared following protocol 'Procedure & Checklist – Preparing HiFi SMRTbell Libraries the SMRTbell Express Template Prep Kit 2.0'. Genomic DNA was fragmented with a mode of ~18 kb using the Diagenode Megaruptor 3. Library was size selected using Sage Science Pippin HT. Polymerase bound SMRTbell complexes were generated using Sequencing Primer v5 and Sequel II Polymerase v2.2. Library was sequenced on a PacBio Sequel II with no pre-extension, 30 hour movies, and a 0.85 predictive loading target using Sequel II sequencing Plate 2.0.

#### ONT whole-genome sequencing

All of the Oxford Nanopore Technologies (ONT) sequencing generated in Jarvis et al. (Jarvis et al. 2022) were re-used in this assembly and validation. In brief, HG0002 was run on PromethION and GridION sequencing instruments. The GridION uses MinION flow cells and the PromethION uses PromethION flow cells. Both flow cells employed the same ONT R9.4.1 sequencing chemistry. PromethION sequencing prepared libraries with the unsheared sequencing library prep protocol and used 28 PromethION flow cells to generate a total of 658x coverage (assuming 3.1 Gb genome size) and ~51x coverage with 100kb+ reads (Shafin et al. 2020). GridION sequencing prepared libraries with the Ultra-Long sequencing library prep protocol and used 106 MinION flow cells to generate a total of ~52x coverage (assuming 3.1 Gb genome size) and ~15x coverage of 100kb+ reads (M. Jain, Koren, et al. 2018). Later, remora methylation calling was performed on some of the runs (labeled in **STable 1**) using Guppy 6.1.2 using the following command line:

```
guppy_basecaller -i {input} -s {save} -c
dna_r9.4.1_450bps_modbases_5mc_cg_sup_prom.cfg -x "cuda:all" -r
```

#### Pacbio Iso-Seq cDNA sequencing

RNA was extracted from three cell lines from HG002 were used to generate Iso-Seq reads; from EBV-immortalized lymphoblastoid cell line (GM24385), iPSC of the EBV-immortalized lymphoblastoid cell line (GM26105), and iPSC derived directly from Peripheral Blood Mononuclear Cells (GM27730). All of these cells were made by and are available from the Coriell Institute for Medical Research. The corresponding Iso-Seq data is available in the NCBI Sequence Read Archive with accession numbers SRR18074967, SRR18074968, and SRR18074969, respectively.

Raw subreads were generated on Pacbio Sequel II instrument using chemistry version 10.1.0.119528 and SMRTLink version 10.1.0.119588. Each sample was generated with a 2 hour pre-extension and a total movie length of 30 hours. The subreads were collapsed and processed with CCS tool v6.0.0 to generate the consensus reads. Subsequently, adapters were removed from the collapsed reads using Lima v1.10.0 with --dump-clips option to output the clipped barcode regions in a separate file. An average of 85% of the CCS reads passed lima filters. The processed reads were assembled using IsoSeq3 v3.2.2 with --require-polya flag to require full length (FL) reads that have a poly(A) tail with at least 20 bp. The primer sequence is used in this step to detect concatemers. An average of 99% of the demultiplexed reads passed refine filters. Subsequently, IsoSeq3's cluster tool was applied to produce unpolished isoforms. Two FL reads were considered the same isoform if they have less than 100 bp difference in their 5' start, less than 30 bp difference in 3' end, and less than 10 bp difference in internal gap with no limit on the number of gaps. Finally, IsoSeq3's polish tool produces polished isoforms. The polish step generates consensus sequences that are divided into high quality (HQ) and low quality (LQ). HQ isoforms have predicted accuracy of 99% or more, while LQ isoforms have predicted accuracy of less than 99%. Both HQ and LQ isoforms must be supported by two or more FL reads. After polishing, more than 99.6% of the unpolished isoforms were HQ.

The CCS reads were generated using the following pbcromwell v1.2.0 command:

```
pbcromwell run pb_ccs \
      -e {input.subreadset.xml} \
      --nproc 8 \
      --config {cromwell.conf} \
      --output-dir {results_dir} \
      --overwrite \
      --tmp-dir /space1/tmp/$PBS_JOBID
```

The demultiplexed reads were generated using the following Lima v1.10.0 command:

```
lima \
      --isoseq \
      --dump-clips \
      -j 24 \
      {input.hifi_reads.bam} \    # input CCS reads
      {input.IsoSeqPrimers_Express_SMRTLink6.0.fasta} \
     {output.demux.bam}
```

IsoSeqPrimers_Express_SMRTLink6.0.fasta is the following:

```
>5p IsoSeq Express Primer
GCAATGAAGTCGCAGGGTT
>3p IsoSeq Express Primer
AAGCAGTGGTATCAACGCAGAGTAC
```

The full-length, non-chimeric (FLNC) reads were generated using the following IsoSeq3 v3.2.2 refine command:

```
isoseq3 refine \
      --require-polya \
      {input.demux.5p--3p.bam} \
      {input.IsoSeqPrimers_Express_SMRTLink6.0.fasta} \
      {output.flnc.bam}
```

The unpolished clustered reads were generated using the following command:

```
isoseq3 cluster \
      -j 32 \
      {input.flnc.bam} \
      {output.unpolished.bam} \
```

The polished clustered reads were generated using the following command:

```
isoseq3 polish \
      -j 16 \
      {input.unpolished.bam} \
      {input.subreads.bam} \
      {ouput.polished.bam}
```

### Assembly

This section gives an overview of the assembly pipeline used in this study, focusing on modifications to the methods presented in Nurk *et al.* (Nurk et al. 2022). While the integration of the “single-copy node bridging” strategy for ONT-based repeat resolution (described below) represents the only major modification to our original approach, a few improvements were also introduced throughout the pipeline (including graph construction and pruning procedures).

#### Graph construction and pruning

As in Nurk *et al.*, reads were first subjected to homopolymer compression and (most) residual errors self-corrected based on >99% identity overlaps. The overlaps were then recomputed and alignment differences falling within microsatellite arrays were masked, after which any overlaps with remaining observed differences were discarded.

Reads longer than 4 kb (after homopolymer compression) and “perfect overlaps” (with no non-masked differences) at least 2 kb long were then used to build Myers’ string graph with modified code from the miniasm codebase.

Instead of further applying miniasm graph processing, estimated coverage values were then assigned to the graph nodes, and a custom set of pruning procedures was applied to obtain the assembly graph. Notable changes in this phase compared to the version used in CHM13 study (Supplements of  (Nurk et al. 2022)) include:

- *Improved accuracy of the HiFi self-correction procedure* with updates to the codebase.
- *Exclusion of reads longer than 18 kb* (after homopolymer compression) from the string graph construction to prevent formation of gaps due to exclusion of reads contained within uncharacteristically long reads (see Fig. 5 of Hui et al. 2016 (Hui et al. 2016) and Fig. S11 of (Nurk et al. 2022)). This step excluded 3% of all considered reads (those longer than 4 kb)
- *Added condition to prevent the removal of genomic links during “weak edge pruning”* (Nurk et al. 2022). The pipeline used progressive 3-4-5-6 kb thresholds on the overlap size with the last two iterations enabling an additional check that blocks the removal of any weak edges leading in/out of the node if all the edges leading in/out of this node were going to be removed (leaving the node a dead-end).
- *Additional heuristics for non-genomic edge removal.* We implemented additional local criteria for the removal of “unusable” edges, i.e. edges that can not be used by any candidate traversal, satisfying node multiplicity constraints. Similar criteria were used for the manual graph pruning in the CHM13 study (see **FigS4** and its caption in Nurk et al. 2022). The procedure is as follows: First, some graph nodes are classified as “single-copy”, meaning their sequence occurs in the genome exactly once, and some as “reliable” meaning that they are unlikely to represent sequencing or methodological artifacts. This is done primarily based on the node length or estimated coverage (with some additional basic checks for consistency between adjacent node assignments). We assume that the assembly graph is complete (in the sense that genomic traversal exists) and that node classification is correct, and consider an edge *l* connecting a single-copy node *u* with a reliable node *v* (**Supplementary Fig. 1a**). If *l* is also the only edge incident to that side of *v,* then any other edge *e* incident to the same side of *u* as *l* is necessarily unusable and can be removed from the graph. Indeed, the genomic path must traverse *v*, which can be

done only via edge *l*, which in turn means that edge *e* can not be genomic, since node *u* is used by the underlying genome traversal only once. To make the procedure more robust to graph artifacts and incorrect node assignments, we only removed an edge *e* if it was deemed unusable while considering both of its incident nodes (unique nodes *u* and *u'* on **Supplementary Fig. 1b**), leaving application of the basic "one-sided" criteria to the discretion of the manual pruning stage. **Supplementary Fig. 1c** illustrates the application of the "two-sided" criteria to the common case of a subgraph corresponding to a 'spanned' copy number two repeat.

- *More conservative handling of bubble structures.* Bubble removal procedures are useful for cleaning the assembly graph from the impacts of recurrent sequencing errors and polymorphic differences within the cell population. At the same time they can lead to a loss of genomic variation between the repeat copies. To prevent such repeat "homogenization", automated bubble handling was mostly limited to cases where estimated coverage of the start/end nodes of the bubble suggested that the subgraph represented a single-copy genomic region. We also switched to analysis of "simple bubbles", where one of the alternatives is represented by a single node, as a primary strategy for handling bubbles encoding non-trivial sequence differences. It allowed us to naturally introduce extra conditions on the ratio of estimated coverages between the alternatives. The basic super-bubble removal procedure (Nurk et al. 2022), not using coverage estimates, was still launched to handle some residual artifacts of the string graph construction process, but was triggered only if the start/end bubble nodes overlapped (based on node length and overlap sizes) and the length difference between alternative paths didn't exceed 50 bp.

Automated procedures were too conservative (and unsophisticated) to effectively remove many non-genomic edges and nodes, in particular within the areas affected by considerable sequencing biases (including some HSAT arrays). Relevant graph components were subjected to extensive manual pruning, which took into account node lengths and estimated coverage, overlap sizes, as well as the broader graph context.

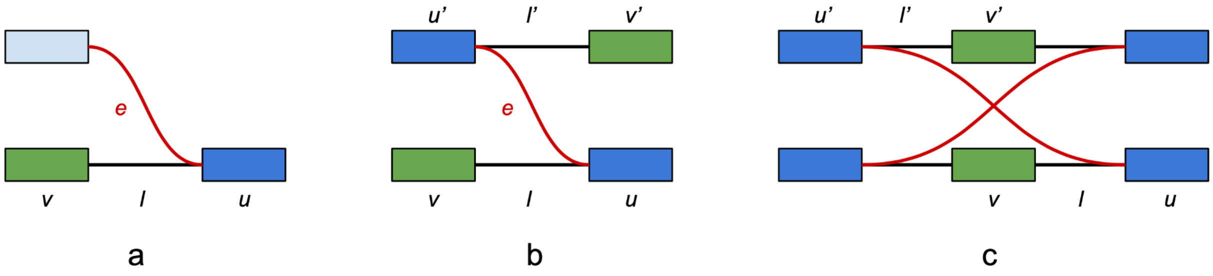


**Supplementary Fig. 1 | Basic heuristics for unusable edge removal around single-copy graph nodes**. Bidirected (Bandage-style) representation of the assembly graph is used in which undirected edges can be incident to either of the two sides of the node. "Reliable" nodes are shown in green, "single-copy" nodes in dark blue, and "unusable" edges in red. **a.** Minimal subgraph providing the evidence for edge *e* being unusable. Edge *l* is the only edge incident to its side of *v*, otherwise additional incident edges are not shown. Note that no requirements are imposed on the subgraph on the other side of *e*. **b.** Subgraph providing "two-sided" evidence for edge *e* being non-genomic. Many cases with only "one-sided" evidence were later processed during manual pruning. **c.** Example of a repeat with copy number two, where each copy is spanned by some reads (forming the reliable nodes *v* and *v'*). Nodes and edges that allow removal of one of the unusable cross-edges are labeled consistently with panel **b**. The other unusable red edge can be removed in the same way by considering the unlabeled pair of single-copy nodes.

#### Using ONT read alignments for tangle resolution

As a first step toward resolving the remaining graph tangles, we aligned homopolymer-compressed ONT reads directly onto the assembly graph using GraphAligner v1.0.13 (Rautiainen and Marschall 2020). GraphAligner's behavior and parameters were fine-tuned to produce more accurate alignments ("-x vg -b 50 --multiseed-DP 1 --X-drop 1000000 --precise-clipping 0.9 --multimap-score-fraction 1 --min-alignment-score 10000"). Alignments covering less than 90% of the read sequence or having average identity below 90% were discarded. In an attempt to ignore typically less reliable alignment margins, each alignment path was then 'trimmed' on both sides to ensure 5 kb minimal extension beyond the overlaps into the first/last remaining path nodes. Note that path trimming has a side effect of "normalizing" the alignment path representation. Indeed, while aligning to the graphs with non-trivial node overlaps, GraphAligner can represent the same alignment path in different ways.

Filtered ONT alignment paths were then used to untangle the graph by an approach we refer to as "*single-copy* node bridging" (reminder that a node is called "single-copy" if it appears in genomic graph traversal exactly once). The same procedure is also integrated in Verkko, with the only major difference being that the reliable single-copy node classification procedure had not yet been implemented when this assembly was performed (and it also might need considerable modifications to work reliably with the string graphs rather than multiplexed MBG graphs). Instead, here we identified the single-copy node candidates using simple criteria based on node length and estimated coverage values, and then curated the list by inspecting the assembly graph.

The bridging procedure analyzes how ONT alignment paths traverse each of the subgraphs enclosed between the set of single-copy nodes, which we refer to as "tangles". For a more detailed description, see the section **"ONT resolution"** in Rautiainen *et al.* (Rautiainen et al. 2022). Briefly, it first computes the number of times each source/sink pair is "bridged" by the ONT alignment paths (i.e. the pair appears as adjacent in the alignment paths, while ignoring non-single-copy nodes). If the count is less than half of the support of some other pair involving the same source or sink, then this evidence is dropped as alignment noise. Otherwise, we consider the nodes to be potentially adjacent in the genome and identify the connecting path with the highest frequency across the ONT alignments. This path, as well as any other connecting path with at least half as much support, is considered to be a potential bridging path for those single-copy nodes. If all single-copy nodes bordering the tangle belong to at least one bridging path then the tangle is resolved: all its "inner" (not single-copy) nodes are removed and the bordering nodes are connected via new nodes corresponding to the bridging paths.

The updated procedure used in Verkko improves the resolvable component condition, additionally checking if all the nodes and edges of the component that seem reliable are used by some bridging path. Here we instead manually checked that all the reliable-looking nodes of the manually pruned graph were incorporated into the result of the repeat resolution process.

Due to various reasons (insufficient node sequence quality, suboptimal alignments, etc.), ONT read alignments produced by GraphAligner were sometimes insufficient to provide reliable evidence for resolving repeats in the regions of very high underlying sequence similarity. Examples include some of the P3 ampliconic palindrome regions in **Extended Data Fig. 1b**.

Similarly to the CHM13 study, those remaining regions were resolved by a brute-force strategy, which evaluated all candidate paths from a particular graph region. To make the analysis more reliable, instead of straightforwardly concatenating graph node sequences (in turn obtained by concatenating individual homopolymer-compressed read sequences), we generated the consensus sequences for all candidate reconstructions (followed by homopolymer compression). ONT reads exhibiting substantial differences in alignment quality to different candidate reconstructions were identified using minimap2 (-x map-ont) (Li 2018, 2). Alignment reports for these reads were then inspected to select the best-supported candidate. In the most ambiguous cases, edlib-aligner (Šošić and Šikić 2017) (with parameters -l -m HW -k 40000) was used to find optimal semi-global alignments of the relevant reads onto the candidate reconstructions or vice versa, depending on the situation.

#### Using parent-specific markers to inform PAR1 reconstruction

Identification of chromosome walks through the PAR1 subgraph presented a unique challenge. A high level of fragmentation in the subgraph representing PAR1 led to unreliable alignments of ONT reads onto the graph (as well as unreliable scoring of the alignments to alternative reconstructions). In particular, when two underlying homologous regions were similar, with one being broken in the graph, GraphAligner would prefer threading all the reads through the single, continuous alternative. Luckily, since the two walks that we needed to recover belonged to ChrX and ChrY we were able to use parent-specific markers, instead of ONT read alignments, to inform their reconstruction. Indeed, we expect ChrY sequences to have only paternal-specific markers and ChrX sequences to have only maternal-specific ones. Parent-specific k-mers (k=30) were collected from (homopolymer-compressed) parental Illumina reads using meryl and then identified within node sequences of the assembly graph using Merqury hampers.sh (Rhie et al. 2020). Nodes having >90% of the overall assigned markers from the particular parental class were assigned as maternal or paternal. Along with the graph structure, those node assignments were enough to identify the walks through each of the components shown on **Extended Data Fig. 1c**. After consensus sequences for all walks were generated, the relation between the draft contigs were identified based on alignments to the ONT-based assembly (see next section).

#### Gap patching and contig stitching

Both ChrX and ChrY were split across multiple connected components in the assembly graph. Discontinuities observed here originated from the issues that HiFi sequencing exhibit in certain microsatellite array regions, in particular ones enriched with (GA)-di-nucleotide repeats (Nurk et al. 2020). HiCanu's consensus module was used to generate contigs for the sub-walks identified across all relevant connected components. To combine those draft contigs reliably into the complete reconstructions, we used a previously-generated Flye v2.7-b1585 (Kolmogorov et al. 2019) assembly of trio-binned ONT reads, which had been polished by medaka and Merfin

(Formenti et al. 2022). Draft contigs were then aligned to the Flye assembly with minimap2 to identify the adjacencies along with the missing sequence. Note that since our draft contigs ended at the regions of sequencing biases, rather than unresolved genomic repeats, we were able to unambiguously identify all underlying adjacencies between the contigs.

Inferred gaps between the draft contigs were directly filled by the corresponding Flye assembly sequences. In cases where minimap2 alignments indicated that draft contigs overlapped, the contigs were joined according to an optimal alignment of the flanking 10 kb regions (mismatch/indel penalty of 300), identified using the parasail library (Daily 2016). We additionally checked that identified overlap sizes closely matched the ones inferred from the alignments against the Flye assembly. Apart from one gap of 20.4 kb in the PAR1 of ChrX, all other gaps were below 7 kb and all re-joined overlaps were below 3 kb (2.7 kb).

Source code and scripts used for assembly graph construction, pruning, semi-automated repeat resolution and consensus can be found in https://github.com/snurk/sg_sandbox (commit ver. 19ee5e306f83f8eb5f5a6ac6a3477e2f925b375e). Note that after finishing the T2T-Y assembly, the entire assembly procedure was re-engineered and updated in the Verkko assembler (Rautiainen et al. 2022). Re-assembly of the HG002 genome with Verkko was able to replicate the T2T-Y assembly, further validating our original reconstruction (with the exception of the P5 inversion error described in the main text, which Verkko correctly assembled).

### Error detection and polishing

For detecting errors and making corrections, we used Illumina WGS 2x250bp reads, a separate set of HiFi reads, and a subset of the ONT reads that were re-basecalled with Guppy v5.0.7 (data marked as Validation in **Supplementary Table 1**). The assembled HG002-X and HG002-Y (HG002XYv2.0) were appended to CHM13v1.1 autosomes to generate a reference fasta for mapping. Both ChrX and ChrY sequences were targeted for polishing.

The overall evaluation and polishing pipeline is similar to that used for polishing the initial T2T-CHM13 assembly (v0.9-v1.1) Mc Cartney *et al.* (Mc Cartney et al. 2022), but with adjustments made to account for the presence of both the maternal (X) and paternal (Y) haplotypes and a few updates to the specific tools used in each step (**Extended Data Fig. 2a**).

#### Sequencing read alignment

Illumina reads were aligned with bwa v0.7.17 (Li and Durbin 2009) using the bwa alignment pipeline available at https://github.com/VGP/vgp-assembly/tree/master/pipeline/bwa. Primary read alignments to chrXY were extracted and de-duplicated using samtools v1.9 (Li et al. 2009) with the following commands:

```
samtools view -hb -F0x104 -@12 illumina.bam chrX chrY >
illumina.pri.XY.bam
samtools collate -o namecollate.bam illumina.pri.XY.bam
samtools fixmate -m namecollate.bam fixmate.bam
samtools sort -o positionsort.bam fixmate.bam
```

```
samtools markdup positionsort.bam illumina.pri.XY.mkdp.bam
samtools index illumina.pri.XY.mkdp.bam
```

##### Primary alignments

HiFi and ONT reads were aligned with Winnowmap v2.03 (C. Jain et al. 2022, 2) using the Winnowmap alignment pipeline available at https://github.com/arangrhie/T2T-Polish/tree/master/winnowmap. In brief, this pipeline aligns the long-reads after downweighting the most repetitive 0.02% of 15-mers present in the reference. The aligned bam file is then filtered to only contain primary alignments using samtools v1.9 (view -F0x104 -@12 -hb). Primary read alignments to ChrX and ChrY were extracted and further used for polishing and evaluation.

##### Marker assisted alignments

To obtain conservatively mapped reads supported by paternal or maternal haplotypes, we generated marker-assisted alignments. First, 21-mers from Illumina WGS of HG002, HG003 (paternal), and HG004 (maternal) were collected with meryl v1.3 (Rhie et al. 2020). Inherited, haplotype specific 21-mers (hapmers) were collected using Merqury (Rhie et al. 2020). Given the k-mer frequency distribution of HG002, we inferred k-mer frequency over 15 had higher probability for being a “single-copy in the genome” vs. “error” in HG002 using Genomescope2 (Rscript genomescope.R -i HG002.k21.hist -o gs2 --fitted_hist), with an inferred “single-copy” peak at 32 (Formenti et al. 2022). We defined an upper-bound as peak*1.5, where the k-mers start to occur at higher probability as belonging to the “two-copy” region in the genome. The “single-copy marker k-mers” were then obtained by intersecting the k-mers occurring only once in the assembly to ensure the marker is unique in both the genome and in the assembly.

Unlike the single-copy marker k-mers used for validating CHM13 (Nurk et al. 2022; Mc Cartney et al. 2022), an additional set of marker k-mers was collected including multi-copy k-mers that were unique to a specific haplotype/chromosome, e.g. multi-copy k-mers from the father were excluded from the ChrX set but included in the ChrY set (and vice versa for the mother). In fact, most of the haplotype-specific duplications or multi-copy kmers observed were on ChrY, due to its unique sequence composition (**Extended Data Fig. 2b**).

Meryl 21-mer dbs were generated using the following command lines:

```
$MERQURY/_submit_build.sh 21 HG002.fofn HG002
$MERQURY/_submit_build.sh 21 HG003.fofn pat
$MERQURY/_submit_build.sh 21 HG004.fofn mat
```

Inherited, haplotype specific k-mers were obtained using the following command lines:

```
$MERQURY/trio/hapmers.sh mat.meryl pat.meryl HG002.k21.meryl
```

Single-copy kmers, occurring as single-copy in the genome were obtained using:

```
meryl less-than 48 [ greater-than 15 HG002.k21.meryl/ ] output
HG002.k21.single.meryl
```

Collection of k-mers in the assembly was performed with:

```
meryl count k=$k $fa output $prefix.meryl
```

Exclusive, single-copy kmers with no k-mers from the other haplotype were collected with:

```
meryl difference HG002.k21.single.meryl pat.hapmer.meryl output
HG002.k21.single.mat.meryl
meryl difference HG002.k21.single.meryl mat.hapmer.meryl output
HG002.k21.single.pat.meryl
```

Single-copy marker kmers, similar set used in Mc Cartney et al. are obtained with:

```
meryl intersect [ intersect HG002.k$k.single.mat.meryl [ equal-to 1
$prefix.meryl ] ] mat.hapmer.meryl output $single.mat.meryl
meryl intersect [ intersect HG002.k$k.single.pat.meryl [ equal-to 1
$prefix.meryl ] ] pat.hapmer.meryl output $single.pat.meryl
```

Multi-copy marker kmers are obtained by this time doing a union with haplotype-specific, multi-copy kmers:

```
meryl union     [ intersect HG002.k$k.single.mat.meryl [ equal-to 1
$prefix.meryl ] ] mat.hapmer.meryl output $marker.mat.meryl
meryl union     [ intersect HG002.k$k.single.pat.meryl [ equal-to 1
$prefix.meryl ] ] pat.hapmer.meryl output $marker.pat.meryl
```

Finally, wiggle tracks for visualization and validation were generated:

```
meryl-lookup -wig-depth -sequence ../assembly/$name.X.fasta -mers
$single.mat.meryl >  $single.wig
meryl-lookup -wig-depth -sequence ../assembly/$name.Y.fasta -mers
$single.pat.meryl >> $single.wig

meryl-lookup -wig-depth -sequence ../assembly/$name.X.fasta -mers
$marker.mat.meryl >  $marker.wig
meryl-lookup -wig-depth -sequence ../assembly/$name.Y.fasta -mers
$marker.pat.meryl >> $marker.wig
```

#### Polishing

We polished the initial HG002XYv2.0 assembly in three major steps aiming to polish: 1) short nucleotide variant (SNV)-like errors (v2.1), 2) telomeric and sub-telomeric end sequences, correct structural variant (SV)-like and remaining SNV-like errors (v2.6), and 3) the remaining errors in the telomere (v2.7). Evaluation was performed after each round of polishing (**Extended Data Fig. 2a**).

Short nucleotide variant (SNV) errors were polished in two rounds. The first round used variant calls from HiFi and Illumina “Hybrid” mode of the DeepVariant v1.2 (Poplin et al. 2018) and ONT

Pepper-Margin pipeline of the DeepVariant (Shafin et al. 2021). Then, each variant set was filtered with Merfin v1.1 (Formenti et al. 2022) similarly as in Mc Cartney *et al.*. Unlike the CHM13 assembly polishing, we observed that the inclusion of all “PASS” ONT variants (including indels and multi-allelic variants) increased the overall assembly quality, as Merfin was able to filter false-positive calls. Therefore, for both Hybrid and ONT modes, variants were filtered only for the “PASS” filter flag in the output VCF and filtered with Merfin. In addition, we used the experimental “better” mode in Merfin for polishing, because the autosomes in this reference originate from CHM13 (not HG002) and this breaks Merfin’s assumption that copy-number differences observed in the reads vs. entire assembly (K*) represent errors.

The second round of SNV error correction was performed after observing insertion biases compared to the HPRC-HG002v1 maternal and paternal assembly as well as compared to the GIAB truth set for HG002 (Details described in “Variants called from assembly with respect to GRCh38 and GRCh37” section). Also, we found false-negative error calls after applying the “better” mode, which was too stringent and removed all true variant calls where both the reference path and the alternate path were supported neutrally by k-mers. This motivated the development of “loose” mode, however relying on the more confident variant call set, and only discards variants when applying the edit would increase the number of error k-mers.

Telomere sequence correction and SV calls were simultaneously targeted for polishing with the second round of SNV error correction on HG002XYv2.1. First, telomere length was estimated using the longest aligned ONT read anchored to the closest “marker” k-mer. Sequences with >40% of telomeric motifs were replaced with a perfect telomeric model sequence (repeating the 6-mer canonical telomeric TAACCC motif at p-arm or GGGTTA at q-arm). This version was named HG002XYv2.2, and was polished for 4 more rounds using Racon v1.4.3 (Vaser et al. 2017) to return any true variants within the telomeres. All edits made to v2.2 were kept in respect to v2.1 for the combined polishing (v2.1 to v2.6).

Second, SV errors were identified using multiple SV callers and subjected to manual inspection, similarly as in Mc Cartney *et al.*. In total, 101 SVs were called by the SV callers and subjected to curation. A majority of the SV calls were false-positives due to excessive read coverage in the primary alignment, or were affected by low-coverage due to sequencing biases in one platform. Additionally, the HPRC-HG002v1.0 assembly-based comparison revealed 1 additional SV-like error. All SV error correction candidates required the support of at least 1 single-copy marker within a reasonable distance on either side of the SV. Collectively, 2 SV sites on the X and 2 on the Y were chosen for further correction, which included small and large base error correction. Among them, 3 were correctable with HiFi reads, 1 lacked HiFi coverage and was only correctable with ONT reads. All the flagged large errors were located at the PAR1 region, due to the lower HiFi coverage.

Combining the telomere patches, SV-edits, and SNV-edits, HG002XYv2.1 was polished to HG002v2.5 and re-evaluated.

Later, a developmental version of the HPRC validation and polishing pipeline Flagger v0.1 (https://github.com/mobinasri/flagger/tree/v0.1) (Liao et al. 2022) was run using the ONT reads on the HG002XYv2.1 assembly, accounting for sequencing biases in the HSat region. After manual inspection, we confirmed 2 SNPs from the ONT variant calling pipeline required correction. Re-combining these two edits with the edits above, HG002XYv2.1 was polished to v2.6 and re-evaluated.

In the meantime, a developmental version of Verkko was run on the HiFi reads and ONT reads with an updated consensus module. The newly generated sequence had better read-to-assembly support on the Xp and Yp subtelomeric sequences, and was used to replace these regions.

Below we provide detailed description and parameters used in each step.

##### Small (SNV) error correction on v2.0

SNVs were called using DeepVariant v1.2 on Illumina reads using "Illumina" mode, Illumina and HiFi reads using the "Hybrid" mode, and on ONT reads using "ONT" mode using the following command lines:

```
# DeepVariant for Illumina
run_deepvariant  --model_type=WGS  --ref=v2.fasta  --reads=input.bam  --
output_vcf=illumina.pri.XY.mkdp.vcf.gz                                --
output_gvcf=illumina.pri.XY.mkdp.gvcf.gz --num_shards=72

# DeepVariant for hybrid Illumina & HiFi
run_deepvariant \
 --model_type "HYBRID_PACBIO_ILLUMINA" \
 --ref v2.fasta \
 --reads input.bam \
 --output_vcf "${OUTPUT_DIR}"/"${OUTPUT_VCF}" \
 --num_shards "${THREADS}" \
 --regions chr20 \
 --intermediate_results_dir "${OUTPUT_DIR}"/intermediate_results_dir

# DeepVariant for ONT
run_pepper_margin_deepvariant call_variant \
 -b input.bam \
 -f v2.fasta \
 -o "${OUTPUT_DIR}"/pepper_deepvariant_output \
 -p "${OUTPUT_PREFIX}" \
 -t "${THREADS}" \
 --ont
```

DeepVariant calls for Illumina were used for further evaluation. Hybrid and ONT calls were filtered, normalized, and merged with bcftools v1.10.2, hap.py v0.3.14 and a custom python script (Mc Cartney et al. 2022) using the following command lines:

```
# Filtering
bcftools view -f "PASS" -Oz $HYBRID_DV_VCF > $HYBRID_DV
```

```
bcftools index $HYBRID_DV
bcftools view -f "PASS" -Oz $ONT_DV_VCF > $ONT_DV
bcftools index $ONT_DV

# Normalize with HAPPY
hap.py \
${HYBRID_DV} \
${ONT_DV} \
-r ${REF} \
-o ${HAPPY_OUTPUT_PREFIX} \
--pass-only \
--engine vcfeval \
--threads "${THREADS}"

# Merge
python3
$tools/T2T_polishing_scripts/polishing_merge_script/vcf_merge_t2t.py \
-v1 ${HYBRID_DV} \
-v2 ${ONT_DV} \
-hv ${HAPPY_OUTPUT_PREFIX}.vcf.gz \
-o "${MERGE_VCF_OUTPUT}"
bcftools index $MERGE_VCF_OUTPUT

# Split to chrX and chrY to apply Merfin
MERGE_X=$MERGE_X.vcf.gz
MERGE_Y=$MERGE_Y.vcf.gz
bcftools view -Oz $MERGE_VCF_OUTPUT chrX > $MERGE_X
bcftools view -Oz $MERGE_VCF_OUTPUT chrY > $MERGE_Y
```

To filter out spurious variants, we collected k-mer sets of HG002 excluding paternal (for X) or maternal (for Y) k-mers and applied Merfin v1.1 for polishing each chromosome individually. First, Illumina *k*-mers were collected using k=21 from HG002, HG003 (paternal) and HG004 (maternal) WGS reads using Meryl v1.3 and Merqury v1.3 (Rhie et al. 2020). Next, haplotype specific k-mers were excluded and filtered from HG002 k-mers. Then, GenomeScope2.0 (Ranallo-Benavidez, Jaron, and Schatz 2020, 2) was used for estimating k-mer coverage and obtaining the probability table and used in Merfin. The following command lines were used:

```
# Build
$MERQURY/_submit_build.sh 21 HG002_reads.fofn HG002
$MERQURY/_submit_build.sh 21 HG003_reads.fofn pat
$MERQURY/_submit_build.sh 21 HG004_reads.fofn mat

# Collect haplotype specific k-mers (hapmers)
$MERQURY/trio/hapmers.sh mat.meryl pat.meryl HG002.k21.meryl
```

```
# exclude maternal and paternal k-mers, and only use k-mers with
frequency greater than 1
meryl greater-than 1 [ difference HG002.k21.meryl mat.hapmer.meryl ]
output HG002.k21.no_mat.gt1.meryl
meryl greater-than 1 [ difference HG002.k21.meryl pat.hapmer.meryl ]
output HG002.k21.no_pat.gt1.meryl

# k-mer coverage obtained with GenomeScope2.0
Rscript $tools/genomescope2.0/genomescope.R -k 21 -i
HG002.k21.no_mat.gt1.hist -o HG002.k21.no_mat.gt1.gs --fitted_hist
# Model converged het:0.00308 kcov:32.6 err:0.000577 model fit:1.1
len:2923221256
Kmercov: 32.58

Rscript $tools/genomescope2.0/genomescope.R -k 21 -i
HG002.k21.no_pat.gt1.hist -o HG002.k21.no_pat.gt1.gs --fitted_hist
# Model converged het:0.00301 kcov:32.6 err:0.000577 model fit:1.1
len:2916612912
Kmercov: 32.57

# Merfin using -better mode
merfin -polish-better -sequence chrX.fa -seqmers chrX.fa.meryl -
readmers HG002.k21.no_pat.gt1.meryl -peak 32.57 -prob
HG002.k21.no_pat.gt1.gs/lookup_table.txt -vcf $MERGE_X -output
chrX.merfin

merfin -polish-better -sequence chrY.fa -readmers
HG002.k21.no_mat.gt1.meryl -peak 32.58 -prob
HG002.k21.no_mat.gt1.gs/lookup_table.txt -vcf $MERGE_Y -output
chrY.merfin
```

Lastly, the consensus was generated using the following command lines:

```
for chr in chrX chrY; do
  bcftools view -Oz --threads 8 $chr.merfin.polish.vcf >
$chr.merfin.polish.vcf.gz
  bcftools index  $chr.merfin.polish.vcf.gz
  bcftools consensus -H1  $chr.merfin.polish.vcf.gz -c
$chr.merfin.polish.chain -f $chr.fa > $chr.merfin.fasta
done
```

The polished XY was then combined again with CHM13v1.1 autosomes for additional polishing (noting hereafter HG002v2.1 for brevity).

##### Telomere correction on HG002v2.1

Telomere correction was performed in 3 steps: 1) replace the error-prone assembled sequence with model telomeric sequences, 2) re-introduce true variants near or inside the telomeres, and 3) refine the telomere length.

For estimating the length of the model telomeric sequence (100% canonical telomeric repeats), ONT and HiFi marker assisted alignments passing the closest expected hap-mer (i.e. maternal for the X, paternal markers for the Y) were retrieved from the pairwise alignment file (.paf) and the largest number of soft-clipped bases was taken as the telomere length.

```
# collecting largest clipped bases on the p-telo
zcat hg002XYv2.1_ont_guppy_5.0.7.markersandlength.paf.gz \
  | awk -v chr=$chr hamper=$pos '$6==chr && $9>hamper && $8<5' \
  | awk '{print $2-$4}' | sort -nr
zcat hg002XYv2.1_hifi.markersandlength.paf.gz \
  | awk -v chr=$chr hamper=$pos '$6==chr && $9>hamper && $8<5' \
  | awk '{print $2-$4}' | sort -nr | head -n1

# on the q-telo
zcat hg002XYv2.1_ont_guppy_5.0.7.markersandlength.paf.gz \
  | awk chr=$chr hamper=$pos '$6==chr && $8<hamper && $9>154343790' \
  | awk '{print $2-$4}' | sort -nr | head -n1
zcat hg002XYv2.1_hifi.markersandlength.paf.gz \
  | awk chr=$chr hamper=$pos '$6==chr && $8<hamper && $9>154343790' \
  | awk '{$2-$4}' | sort -nr | head -n1
```

As expected, the longest spanning reads were obtained from the ONT reads. From each end of the chromosomes, the closest canonical telomeric sequence position was selected towards the end of the regions annotated to contain >40% of telomeric sequence motifs ("last non-telomeric position" for brievity).

The model sequence was generated from the closest length to the clipped + last non-telomeric position, and used to replace the assembled sequence from the beginning (end) of the chromosome to the last non-telomeric position for the p (q) arm.

After re-aligning the HiFi and ONT reads with Winnowmap, reads were filtered to generate the marker-assisted alignments. No supplementary alignments were used. Each telomeric end was then subjected to polishing. After visual inspection, regions for polishing were defined where there was inconsistency between the reads and the assembly. Unless the HiFi sequencing depth was very low (e.g. Yp-telo), model sequences were in general in good agreement with both HiFi and ONT reads, except for the sub-telomeric region, where the true sequence diverged from the model. In these cases, HiFi reads spanning the closest hap-mer were extracted and the Racon v1.4.3 "liftover" branch was applied using the `L` option to output edits in VCF. The VCF was post-filtered to include edits only on the selected region.

```
racon -t 20 telo_hifi.filt.fq telo_hifi.sam v2.2.XY.fasta -L
v2.2.XY.racon > v2.2.XY.racon.fasta
```

After initial polishing, the Yq-telo region still showed sequence discrepancies when comparing the spanning reads to the polished consensus, and this region was iteratively polished and re-evaluated 3 more times.

At the final round, both the spanning HiFi and ONT reads were aligned to the polished intermediate consensus to refine the beginning (end) of the model sequence in the p (q) arm. The end position of each chromosome was chosen as the most distal base supported by at least 2 HiFi or ONT reads and were trimmed back so that the first (last) base began (ended) with the canonical telomeric repeat unit.

The resulting polished telomere edits were mapped back to v2.1 to determine the boundaries, then added to the SV edits.

##### Large (SV) error correction on HG002v2.1

The HiFi and ONT primary alignments were re-generated on HG002v2.1 to call structural variants (SVs) using Sniffles v1.0.12a (Sedlazeck et al. 2018) and cuteSV v1.0.12 (T. Jiang et al. 2020). For Sniffles we used default parameters (see below "sniffles"). For cuteSV we used both default and suggested parameters (see code below, "cuteSV" and "cuteSV-sug") for either HiFi or ONT data, which adjusts the "max cluster bias" and the "ratio of merging" for insertions and deletions. cuteSV also utilizes a reference genome, which in this case consisted of the ChrX and ChrY assemblies. Finally, all SV calls were merged using SURVIVOR v1.0.7 (Jeffares et al. 2017) using the following options: a maximum distance between breakpoints of 1 kb, a minimum caller support of one, taking into account the SV type and a minimum SV size of 50 bp (see code below, "survivor").

```
#sniffles
sniffles \
    --mapped_reads hg002XY-[HiFi|ONT]-v2.1.bam \
    --vcf hg002XY-[HiFi|ONT]-v2.1-sniffles.vcf

#cuteSV
cuteSV \
    hg002XY-[HiFi|ONT]-v2.1.bam \
    hg002XY-v2.1.fasta \
    hg002XY-[HiFi|ONT]-v2.1-cuteSV.vcf \
    cuteSV-tmp-cuteSV \
    --genotype

#cuteSV-sug: ONT
cuteSV \
    hg002XY-ONT-v2.1.bam \
    hg002XY-v2.1.fasta \
```

```
    hg002XY-ONT-v2.1-cuteSV-suggested.vcf \
    cuteSV-tmp-cuteSVsugONT \
    --max_cluster_bias_INS    100 \
    --diff_ratio_merging_INS  0.3 \
    --max_cluster_bias_DEL    100 \
    --diff_ratio_merging_DEL  0.3 \
    --genotype
#cuteSV-sug: HiFi
cuteSV \
    hg002XY-HiFi-v2.1.bam \
    hg002XY-v2.1.fasta \
    hg002XY-HiFi-v2.1-cuteSV-suggested.vcf \
    cuteSV-tmp-cuteSVsugHiFi \
    --max_cluster_bias_INS    1000 \
    --diff_ratio_merging_INS   0.9 \
    --max_cluster_bias_DEL    1000 \
    --diff_ratio_merging_DEL   0.5 \
    --genotype

# make survivor file list
ls hg002XY-*.vcf > survivor_file_list.txt

#survivor
MAX_DIST_BREAKPOINTS=1000  # Max distance between breakpoints (0-1
#  percent of length, 1- number of bp)
MIN_SUPP_CALLER=1          # Minimum number of supporting caller
USE_TYPE=1                 # Take the type into account
#  (1==yes, else no)
USE_STRAND=0               # Take the strands of SVs into account
#  (1==yes, else no)
UNUSED_PARAMETER=0         # Disabled.
MIN_SV_SIZE=50             # Minimum size of SVs to be taken
#  into account.
SURVIVOR merge \
    survivor_file_list.txt \
    ${MAX_DIST_BREAKPOINTS} \
    ${MIN_SUPP_CALLER} \
    ${USE_TYPE} \
    ${USE_STRAND} \
    ${UNUSED_PARAMETER} \
    ${MIN_SV_SIZE} \
    hg002XY-HiFi-v2.1-mergeSV.vcf
```

The final SV set contained 21 variants on ChrX and 80 on ChrY. We manually inspected the HiFi and ONT primary and marker-assisted read alignments to validate the SV calls. A large cluster of false-positive SVs was found on the HSat arrays caused by mapping biases in the ONT reads. Three SV-like regions were identified to be correctable, one on ChrX and two on ChrY. The SV calls also identified mis-assemblies in the telomeric sequences, which required additional investigation.

Alternatively, the HG002v2.1 ChrX and ChrY assemblies were also compared against HPRC-HG002v1.0 (Jarvis et al. 2022). Using dipcall v0.3 (Li et al. 2018) with minimap v2.20 (parameter -z200000,10000) to align the two assemblies and call variants. After careful examination of the HiFi and ONT read alignments in the vicinity of variants, we found most of the regions had been correctly assembled in HG002v2.1; however, we did detect one region on ChrX that required further correction.

All four regions flagged for correction were located in PAR1, the coordinates of which were estimated by aligning ChrY to ChrX using Mashmap2 v2.0. We obtained a one-to-one alignment with percent identity >95% at 5 kb segments with options --filter_mode one-to-one -pi 95. The PAR1 and PAR2 regions were identifiable with identity over 98.5%.

For each SV correction candidate region, we extracted read alignments from the marker-assisted alignments. We tried Racon v1.6.0 "liftover" branch and Flye v2.9 in both polishing mode and consensus mode to generate patches for each locus, and then chose the best concordance patch after re-aligning the reads. In brief, we used Racon with HiFi read alignments using options to produce edits in VCF format (-t24 reads.fq reads.sam asm.fa -L asm.out). Flye polish mode was run with HiFi reads using --out-dir out --pacbio-hifi reads.bam --polish-target asm.fa options or ONT reads using --out-dir out --nano-hq reads.bam --polish-target asm.fa options. Flye consensus mode was run with HiFi reads using --threads 24 --out-dir out --pacbio-hifi reads.fq and with ONT reads using --threads 24 --out-dir out --nano-hq reads.fq. The resulting polished sequence patch was used as a target for re-aligning the HiFi and ONT reads using Winnowmap2 as described above. After manual inspection, the best concordant patch was chosen. Patch sequences were aligned to the sequence of HG002XYv2.1 using Winnowmap2 -x asm5 -t 24 --MD -a options, and simple differences (edits) were called using dipcall v0.3 (Li et al. 2018) dipcall/dipcall.kit/htsbox pileup -vcf asm.fa in.bam. One region was missing a large replacement sequence because of the soft-clipped assembly alignment, which has been manually recovered after extracting coordinates from the alignment in the patch sequence. The final edits within the SV-error region were collected for further polishing (**Supplementary Table 2**).

##### Variants called from HPRC assembly with respect to GRCh38 and GRCh37

As part of the polishing evaluations, we evaluated the accuracy of variants called from the HG002-X and HG002-Y assemblies aligned to GRCh38, comparing small variants to a HiFi DeepVariant callset from the precisionFDA Truth Challenge V2 and a scaffolded trio-hifiasm assembly from >100x HiFi reads (Jarvis et al. 2022). To evaluate the accuracy of insertions and deletions >=50bp, we also aligned the HG002v2.1 assembly to GRCh37 and compared its structural variant calls to the GIAB v0.6 SV benchmark.

Variants on ChrX and ChrY were called from HG002XY v2.1, v2.5 and v2.7 assemblies using dipcall v0.3 (Li et al. 2018). We used the -z200000,10000 parameter with minimap2 to improve alignment contiguity, as it has previously been shown to improve variant recall in regions with dense variation like the Major Histocompatibility Complex (Chin et al. 2020). The assembly was treated as "male", using the ChrX assembly for its maternal haplotype and the chromosome Y assembly for its paternal haplotype.

Small variant evaluation was performed using hap.py v3.15 (https://github.com/Illumina/hap.py) and the HG002 ChrX/Y callset output from dipcall as "truth". We compared small variants from a GRCh38 HiFi DeepVariant callset from the precisionFDA Truth Challenge v2 (N. D. Olson et al. 2022) and a GRCh38 dipcall callset from a scaffolded trio-hifiasm assembly from >100x HiFi reads (Jarvis et al. 2022). Targeting was not used for the comparison of the HiFi DeepVariant callset as there was no associated region file. Targeting however was performed for the comparison of the trio-hifiasm callset using the associated dipcall region file (dip.bed). For better comparisons of complex variants, hap.py was run using vcfeval (https://github.com/RealTimeGenomics/rtg-tools). Variant calls were stratified using GIAB stratifications v3.0 (https://doi.org/doi:10.18434/mds2-2499), stratifying true positive, false positive, and false negative variant calls in challenging and targeted regions of the genome. Small variant evaluation was performed for all callsets.

To evaluate accuracy of insertions and deletions ≥50 bp, we also aligned the v2.1 assembly to GRCh37 and compared the assembly's variants to the GIAB v0.6 SV benchmark (Zook et al. 2020). Evaluation was performed with truvari bench v3.1.0 (English et al. 2022) for the HG002 ChrX/Y callset following multiallelic split of alleles in dipcall output using bcftools norm. Comparisons were constrained to regions of the GIAB v0.6 SV benchmark using the `--includebed` parameter. Additional parameters (--multimatch –passonly -r 2000 -C 2000) were used to account for differences in the representation of multiallelic variants and matching variants in long tandem repeats.

The evaluation on v2.1 revealed a strong 1-bp insertion bias in the assembly that could be potentially further polished at a 2nd round of SNV correction. All variants from the evaluation were projected back to v2.1 assembly for further investigation (here, HPRC variants for brevity).

##### Homopolymer and 2-mer evaluation

In order to reduce possible Illumina or HiFi sequencing biases in the HPRC variants, we intersected the calls with DeepVariant calls made with Illumina, HiFi, and Hybrid mode on v2.1 and named them as "HPRC-Illumina" and "HPRC-Hybrid" variant set, respectively). The variants on v2.1 were called in the same way as in the first round of SNV error correction. The intersected variant indel length difference was then compared against the homopolymer and 2-mer length differences in the HiFi and Illumina reads aligned at each variant. This step was performed using the runlength code to compute the run length matrix from the CHM13 assembly evaluation (https://github.com/arangrhie/T2T-Polish/tree/master/runlength) (Mc Cartney et al. 2022).

First, we applied a “PASS” filter to both variant sets in order to remove unreliable reference calls from each intersected variant set, and compared the run length difference observed in HiFi and Illumina reads. Regardless of the DeepVariant mode used, we observed that the HiFi reads suggested a 1-bp insertion bias in the assembly (assembly has an extra base), while the Illumina reads were more in agreement in the homopolymers. In contrast, the 2-mer microsatellite length suggested a 1-bp deletion bias (assembly is missing one base) in both the HPRC-Illumina and HPRC-Hybrid sets, with mixtures of inconsistently supported variants.

In order to apply a more conservative criteria, we chose variants called as homozygous in addition to the “PASS” filter, and compared the run length differences again. With this criteria, the homopolymers showed a cleaner peak in the HPRC-Illumina data and supported the 1-bp insertion bias, as was seen in the HiFi reads. The 2-mer length difference also had a cleaner agreement between the HiFi and Illumina reads, with fewer inconsistent variants.

We confirmed, by applying the “PASS” and heterozygous filter, that the majority of the inconsistent homopolymer and 2-mer patterns were observed in both HiFi and Illumina reads. Therefore, we decided to use the “PASS” and “Homozygous” filter and prepared the candidate variants for further filtering with Merfin.

##### Less conservative filtering with Merfin

Because the small errors we are trying to fix are more likely to be called correctly given the shorter length of the variant, we chose to include the HPRC-Illumina version of the variants in case there was a conflict with the HPRC-Hybrid. However, given that some variants are only accessible with the HiFi reads, we appended variants specific to HPRC-Hybrid set to the candidate set. The merging process was performed with bcftools v1.3 (Danecek et al. 2021). Before obtaining the candidate variant set, we excluded all regions previously curated in telomere and SV correction steps. The merged SNV edits were run with Merfin “loose” mode using the same 21-mer dbs as in the first iteration. This mode ensures that the variants are excluded only if the suggested correction increases the number of missing Illumina k-mers (error k-mers). Lastly, the telomere edits and SV edits were merged into v2.1_sv_edits.vcf, and the Merfin polished edits as v2.1_snv_edits.vcf. Both vcfs were appended as v2.1_to_v2.5.vcf, and used for building the v2.5 consensus.

```
# Concatenate the X and Y calls
for DV in illumina hybrid
do
  bcftools concat -Oz hprc_$DV.sv_masked.PASS.HOM.X.vcf.gz
hprc_$DV.sv_masked.PASS.HOM.Y.vcf.gz >
hprc_$DV.sv_masked.PASS.HOM.vcf.gz
  bcftools index hprc_$DV.sv_masked.PASS.HOM.vcf.gz
done

# Intersect
bcftools isec -p isec hprc_illumina.sv_masked.PASS.HOM.vcf.gz
hprc_hybrid.sv_masked.PASS.HOM.vcf.gz
```

```
# Hybrid-only calls
cd isec
bcftools view -Oz 0001.vcf > hybrid_only.vcf.gz
bcftools index hybrid_only.vcf.gz

# Concat
cd ..
DV=illumina_plus_hybrid
VCF=hprc_$DV.sv_masked.PASS.HOM.vcf.gz
bcftools concat -Oz -a hprc_illumina.sv_masked.PASS.HOM.vcf.gz
isec/hybrid_only.vcf.gz | bcftools sort -Oz - > $VCF
bcftools index $VCF

# Merfin
./merfin_loose.sh $VCF v2.1.XY.fasta HG002.k21.no_pat.gt1.meryl
HG002.k21.no_mat.gt1.meryl 2> merfin_loose.$DV.log
```

##### HPRC Flagger pipeline on v2.1

An updated version of Flagger (https://github.com/mobinasri/flagger) was developed after we found substantial sequencing biases in HiFi and ONT reads, which affected Flagger's coverage-based evaluation. This updated pipeline is identical to v0.1 release, except for step 2. More details are given in Liao *et al.* (Liao et al. 2022) and "Incorporating HSATs biases" (https://github.com/mobinasri/flagger/tree/v0.1/docs/flagger#2-incorporating-hsats-coverage-bias).

Flagger reported 3 unreliable blocks, with two possible SNV corrections called from ONT reads. However, there were 6 additional variants having inconsistent results or not enough support from the marker-based HiFi or ONT alignments that were disregarded.

These two SNV corrections were again added to the merged SV and SNV edits from v2.1, and used to build the v2.6 version of the HG002 XY assemblies.

##### Telomere correction on v2.6

To polish the telomeres, we selected HiFi reads from the marker-assisted mappings that aligned to the p-arm telomere of chromosomes X and Y as performed on v2.1. These reads were re-mapped with winnowmap v2.03:

```
winnowmap -ax map-pb assembly.fasta reads.fastq > reads.sam
```

and the consensus was polished with racon v1.4.3 using the command:

```
racon reads.fastq assembly.fasta polished.fasta -w 50000 -e 0.02
```

This process was repeated for three rounds.

The ChrX telomere still showed indels post polishing so we instead used consensus from a verkko assembly which had better agreement with the reads. For ChrX, the sequence from coordinates 4701 to 5692 was replaced with a 2000 bp sequence and for ChrY the sequence from 1114 to 15797 was replaced with a 14722 bp sequence.

#### Evaluation

Polishing and evaluation are tightly inter-dependent. As such, part of the evaluation procedure overlapped steps performed in the “Polishing” section and is described in detail. Here, we are providing details not covered in the “Polishing” section. Each evaluation method is summarized in **Extended Data Fig. 2a** (left panel).

##### Merqury

The 21-mers from HG002 Illumina reads and the hapmers from HG003 and HG004 generated in “Polishing, Marker assisted alignment” section were re-used for running Merqury v1.3. The entire trio-mode pipeline was run on the Illumina reads for HG002v2.0-v2.7 to obtain QV estimates and evaluate haplotype switches (**Supplementary Table 3**). Likewise, the 21-mers of the HG002 HiFi reads were obtained using the same k-mer building pipeline in Merqury as in the “Marker assisted alignment” section. A hybrid k-mer db was created using the Illumina and HiFi 21-mers, after excluding the unique k-mers observed in each k-mer databases. The HiFi and Hybrid k-mers were also used to evaluate QVs on all assemblies. We note that because of the highly repetitive nature of ChrY, it is more likely for an error k-mer in the assembly to match by chance with a true k-mer in the reads (e.g. a k-mer induced by a one base deletion error in the assembly may actually exist somewhere else in the genome). Thus, the k-mer-based QV estimates are expected to be slightly inflated for more repetitive chromosomes. The corrections applied between v2.5-v2.7 were minor (2 SNPs and edits at the repetitive telomeric sequences), and the QV estimates from v2.6 and v2.7 were identical to v2.5 even though the read-to-assembly agreement was improved.

The hybrid k-mer db was obtained with:

```
meryl greater-than 1 illumina.k21.meryl output illumina.gt1.meryl
meryl greater-than 1 hifi_val.k21.meryl output hifi_val.gt1.meryl
meryl union-sum illumina.gt1.meryl hifi_val.gt1.meryl output hybrid.meryl
```

The hapmer wiggle tracks and error (asm_only) wiggle tracks are used for visually inspecting possible switch errors or true base-pair errors in each assembly version.

##### Coverage analysis

We used the Winnowmap primary and marker assisted alignments for performing coverage analysis. Coverage based issues observed in HiFi and ONT reads were detected using the issues.sh script from Mc Cartney et al. (https://github.com/arangrhie/T2T-Polish/tree/master/coverage). In short, this script generates regions with excessive clippings or low support from a given alignment. The low support regions are further annotated when it overlaps with a hybrid error-kmer at the same position “Low_Error” or overlaps with regions enriched (>80%) for 2-mer microsatellite repeat. In addition, this script generates alignment coverage statistics and multiple alignment summary tracks in wiggle format such as minimum/maximum/average/median of coverage, read-to-assembly identity, mapping quality (MQ), reads per strand, number of clipped reads in every 1024 bp window by default. All of the median wiggle tracks were used in addition to the clipping track for evaluation.

The resulting issues.bed file from the marker-assisted alignment of HiFi and ONT reads were further investigated for cataloging potential remaining issues (**Supplementary Table 2**).

##### VerityMap

Following **Supplementary Note 7** of Nurk et al. 2022 (Nurk et al. 2022), we performed orthogonal validation of the assembly using VerityMap v2.1.0-alpha, https://github.com/ablab/VerityMap (Bzikadze, Mikheenko, and Pevzner 2022). VerityMap first identifies all rare *k*-mers in the assembly, carefully selects a small subset of rare *k*-mers (*solid k-mers*), finds locations of solid *k*-mers in each read, and constructs a *compatibility graph* (Bzikadze, Mikheenko, and Pevzner 2022) with the vertex-set formed by all matches between the selected solid *k*-mers shared by a read and the assembly. Then, VerityMap finds an optimal path in this graph, uses this path for read mapping, and finds misassembly breakpoints using these mappings. In more mathematical terms:

Let $a_R$ and $b_R$ ($a_A$ and $b_A$) be occurrences of solid *k*-mers *a* and *b* in the read *R* (contig *A*) such that $a_R$ precedes $b_R$ ($a_A$ precedes $b_A$). We define $d(a_R, b_R)$ ($d(a_A, b_A)$) as the distance between $a_R$ and $b_R$ in *R* ($a_A$ and $b_A$ in *A*). We refer to the pair of $a_A$ and $a_R$ ($b_A$ and $b_R$) as a *match* $a_M$ ($b_M$) and define $diff(a_M, b_M) = |d(a_R,b_R) - d(a_A,b_A)|$.

VerityMap detects approximate locations of misassembly breakpoints: for example, a deletion of length *L* in an assembly typically triggers a pair ($a_M$, $b_M$) with a surprisingly large difference $diff(a_M, b_M) \cong L$, where $a_M$ and $b_M$ represent solid *k*-mers flanking the deletion breakpoint in such a way that $a_M$ ($b_M$) precedes (follows) the deletion breakpoint. If nearly all reads covering *k*-mers *a* and *b* show a *systematic bias* in values of $diff(a_M, b_M)$, then an indel of approximate size $diff(a_M, b_M)$ is likely contained between $a_A$ and $b_A$. We refer to each read that shows systematic bias in values of $diff(a_M, b_M)$ as a *discordant* read and report the fraction of discordant reads connecting each pair of solid *k*-mers in the assembly. If only a fraction *MinHetFraction* (default value = 30%) of reads containing both *a* and *b* are discordant, the region between $a_A$ and $b_A$ might contain a heterozygous site.

**Supplementary Fig. 2** shows analysis of discordant HiFi and ONT reads mapped to the assembly of chromosome Y in HG002. All putative events were further manually inspected in IGV v2.14.1 for validity. Specifically, a putative heterozygous position at ~59.1 Mb reported by HiFi reads is flanked by a drop in coverage reported by both VerityMap and Winnowmap. Since ONT reads do not show any coverage abnormalities in this region, and no ONT reads are discordant in this region, this heterozygous position might be spurious and explained by HiFi coverage drop-out in **Supplementary Fig. 3**. Manual analysis of read mappings at position ~40 Mb reveals a potential misassembly (**Supplementary Fig. 4**): all but two HiFi reads are clipped and numerous ONT reads report indels of various lengths. HiFi read mappings produced by Winnowmap show reduced coverage in this region and enrichment of nucleotide positions with reference base differing from the read consensus base.

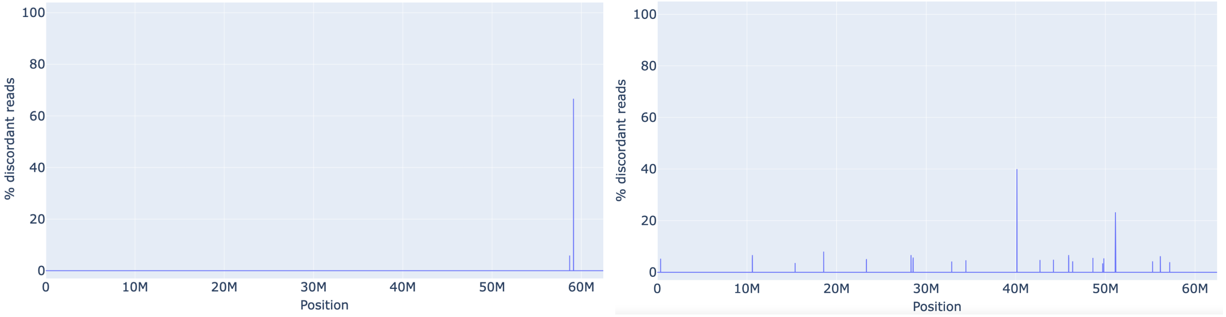


**Supplementary Fig. 2 | VerityMap validation.** The x-axis is chromosome position and the y-axis is percent deviated reads (0-100%). Left (right) subfigure corresponds to mapping of HiFi (ONT) reads.

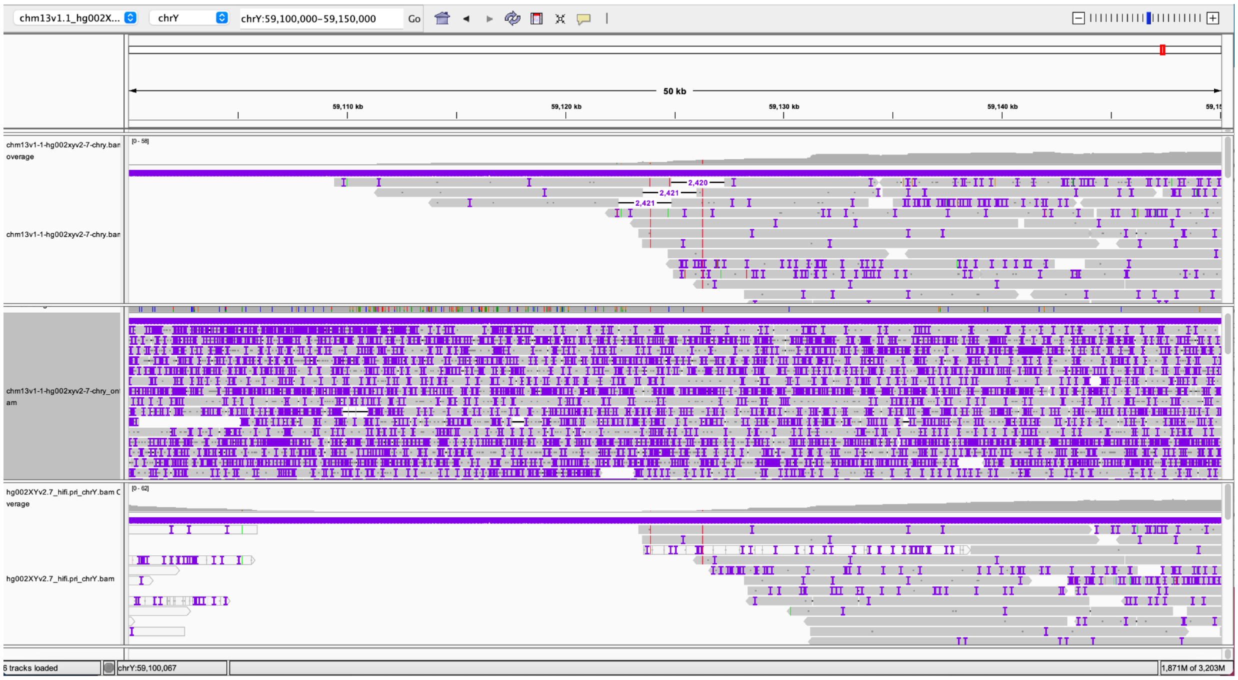


**Supplementary Fig. 3 | VerityMap validation of ChrY around 59.1 Mb using long-read mapping.** Mappings of HiFi reads are produced by VerityMap (top), of ONT reads — by VerityMap (middle), of HiFi reads by Winnowmap (bottom). Both Winnowmap and VerityMap indicate a drop in HiFi coverage depth. Only three HiFi reads mapped by VerityMap show ~2.5kb deletions in assembly. No mapped ONT reads support these deletions.

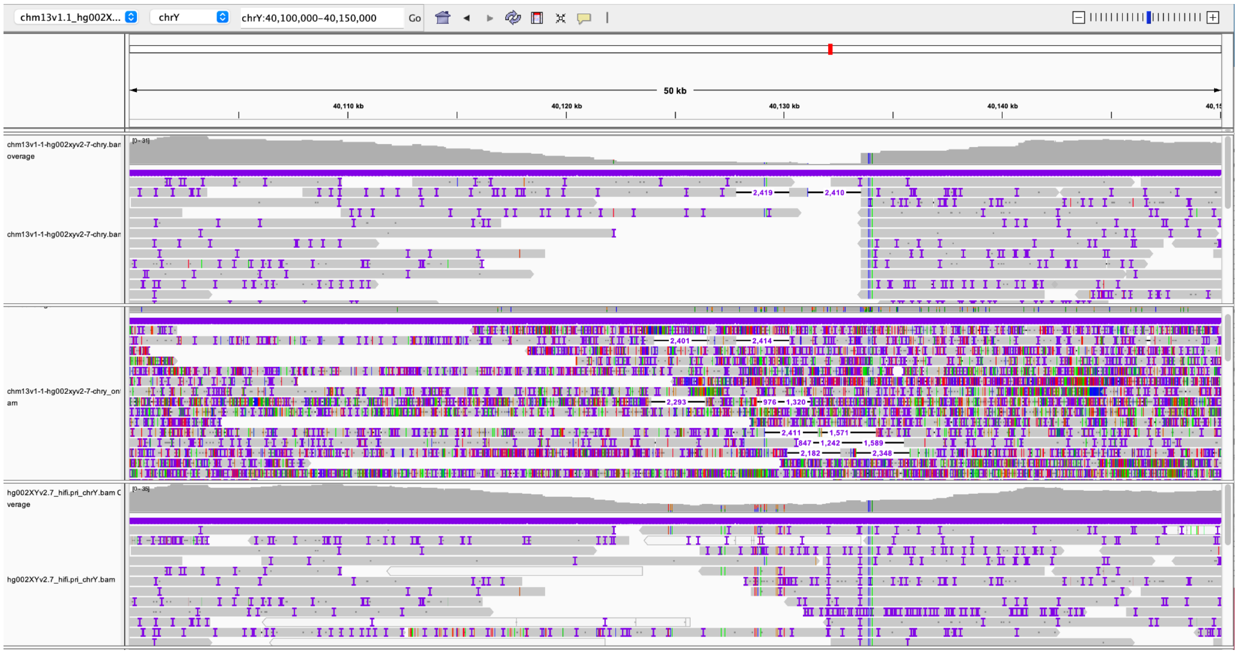


**Supplementary Fig. 4 | VerityMap validation of ChrY around 40 Mb using long-read mapping.** Mappings of HiFi reads are produced by VerityMap (top), of ONT reads by VerityMap (Middle), of HiFi reads by Winnowmap (bottom). Little support in HiFi VerityMap and WinnowMap suggests possible misassembly in this region

#### Second-most frequent allele with NucFreq

The chromosome-wide second-most frequent alleles observed in the HiFi reads and blocks of enriched second alleles were obtained using NucFreq v0.1 (Mitchell R. Vollger et al. 2022; Mc Cartney et al. 2022). NucFreq was run using the following commands to collect regions where the second most frequent allele from the HiFi marker-assisted Winnowmap alignments had an allelic frequency over 30% and sites 1 kb adjacent to one another were merged:

```
python3 NucPlot.py --n HiFi --height 4 -y 100 -t 16 --bed all.bed --obed
hifi.marker.nucfreq.bed --minobed 2 hifi.pri.bam hifi.png
cat hifi.marker.nucfreq.bed \
  | awk 'NR>1 {print $0"\t"100*$5/($4+$5)}' | awk '$NF>30' \
  | bedtools merge -d 1000 -i - > hifi.marker.nucfreq.30.mrg.bed
```

#### Strand-seq evaluation

Strand-seq data were generated as previously described (Sanders et al. 2020; Ebert et al. 2021). Raw FASTQ files from 65 Strand-seq libraries were aligned to both the GRCh38 reference assembly (GCA_000001405.15) and the T2T-CHM13v1.1 + HG002XYv2.7 assembly using the BWA aligner v0.7.17-r1188 (Li and Durbin 2009). Next, each BAM file was processed using the R package breakpointR (Porubsky, Sanders, Taudt, et al. 2020) using the following parameters: windowsize = 2000000, binMethod = 'size', pairedEndReads = TRUE, min.mapq = 10, genoT = 'binom', background = 0.1, minReads = 100. Results of this analysis were used to generate directional composite files as previously described (Sanders et al. 2016; Porubsky, Sanders, Höps, et al. 2020), using breakpointR function “synchronizeReadDir” (Ebert et al. 2021). In order to detect recurrent changes in strand directionality we ran breakpointR again on such composite files with the following parameters: windowsize = 50000, binMethod = 'size', pairedEndReads = TRUE, min.mapq = 10, genoT = 'binom', background = 0.1, minReads = 50. Putative assembly errors would be detectable in composite files as regions where reads overwhelmingly map in minus orientation indicative of a misorientation or unresolved inversion. Putative collapses in the assembly would be visible as regions where reads map in both the minus and plus orientation.

##### Integrity with HG003Y

Additionally, to check the integrity of the HG002 cell line versus its original source, we compared against the HG003 cell line, which was derived from the father of the HG002 donor. If the structure of the Y chromosome in HG002 and HG003 is in agreement, then it is very likely that no structural variants arose during cell culture (otherwise, the exact same variants would have had to independently arise in both lines). To test this, we collected all publicly available long-read data from both HG002 and HG003, which included older HiFi and ONT UL data from HG003 and more recent HiFi (Revio) instruments. Data from HG003 was mapped to the T2T-CHM13+Y assembly, and HG002 to T2T-CHM13 autosomes + HG002v2.7XY as done for polishing and validation.

We observed no evidence for large structural variation between the T2T-Y assembly and any of the additional HG002 or HG003 sequencing data (**Supplementary Fig. 5**). Also, the HSat1/HSat3 coverage bias observed in our sequencing data disappears (e.g. see the uniformity of HG003 HiFi coverage tracks). The newer HG003 HiFi (Revio) data shows even better agreement with the HG002 assembly than the older HG002 HiFi data, and the number of sites flagged as potential assembly issues is fewer when using the newer data. Thus, given the agreement of the HG003 sequencing data mapped to the T2T-HG002-Y assembly, we conclude that the T2T-Y assembly is a faithful reconstruction of a human Y chromosome.

HG003 GIAB HiFi data is available to download at: https://ftp-trace.ncbi.nlm.nih.gov/ReferenceSamples/giab/data/AshkenazimTrio/HG003_NA24149_father/PacBio_CCS_15kb_20kb_chemistry2/CHM13v2.0/GIAB_5mC_CpG/

HG003 ONT UL data was generated by HPRC and is available to download at: https://s3-us-west-2.amazonaws.com/human-pangenomics/index.html?prefix=NHGRI_UCSC_panel/HG003/nanopore/Guppy_4.2.2/

HG002 and HG003 HiFi Revio data is available to download at: https://downloads.pacbcloud.com/public/revio/2022Q4/

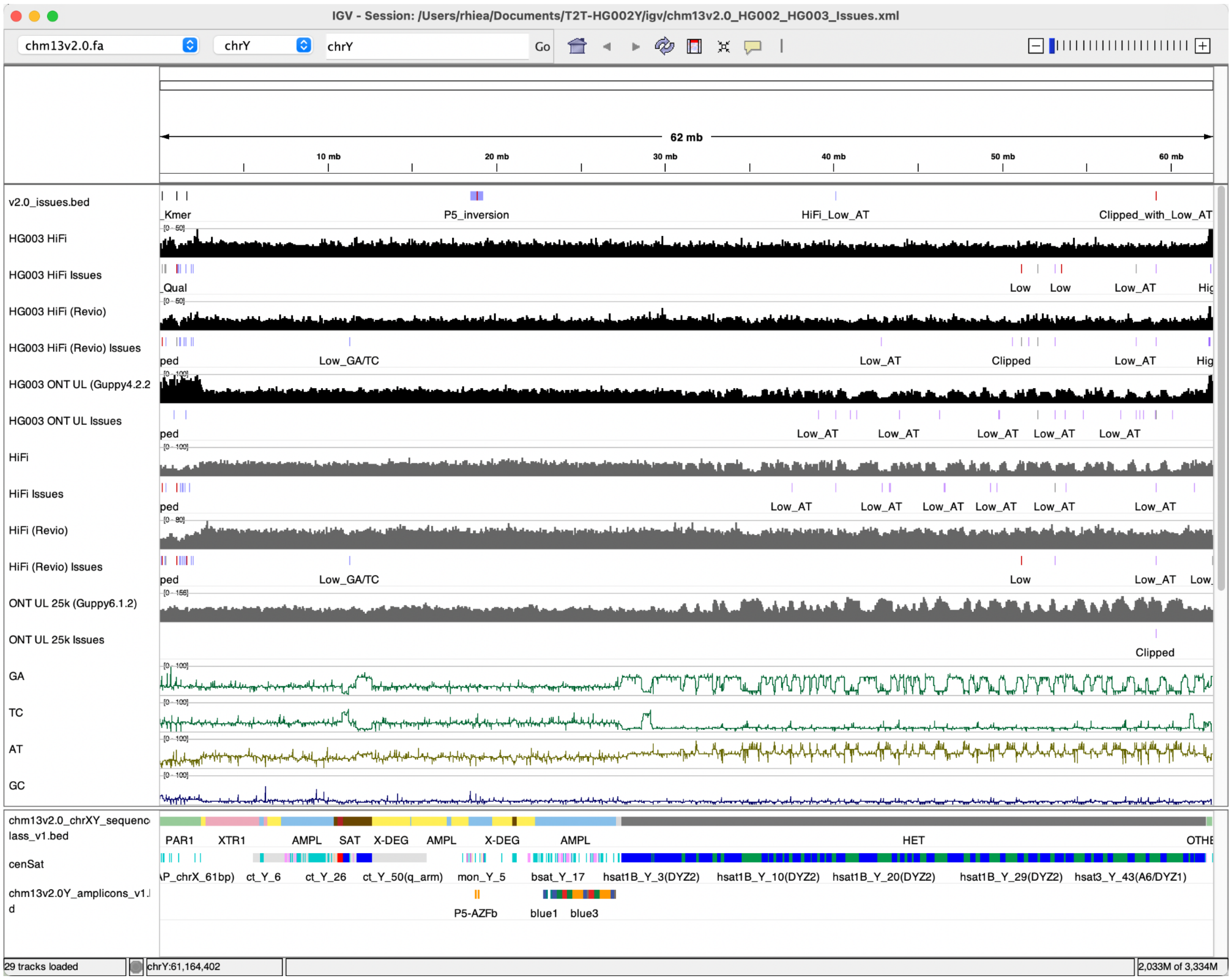


**Supplementary Fig. 5 | Mapping coverage and issue tracks for various read sets from HG003 and HG002.** Datasets from HG003 are labeled on each track with "HG003". The rest are from HG002. All 3 available replicates of HG002 HiFi (Revio) datasets were merged and is shown in one track. Bottom track displays 2-mer microsatellites in each non-overlapping 128 bp window.

#### Chromosome spreads and Fluorescent In-Situ Hybridization (FISH)

GM24385 (HG002) lymphoblastoid cells (LCLs) were obtained from Coriell and cultured at 37°C in 5% CO2 in RPMI 1640 medium (Corning) supplemented with 15% FBS and 1X Glutamax (Gibco). For chromosome spread preparation, cells were arrested in mitosis by the addition of Karyomax colcemid solution (0.1 µg/ml, Life Technologies) to the growth medium for 6 hours. Cells were collected by centrifugation at 200g for 5 minutes and incubated in 0.4% KCl swelling solution for 10 minutes. Swollen cells were pre-fixed by the addition of freshly prepared Methanol: Acetic acid (3:1) fixative solution (~100 µL per 10 ml total volume). Pre-fixed cells were collected by centrifugation at 200 g for 5 minutes and fixed in Methanol: Acetic acid (3:1) fixative solution. Spreads were dropped on a glass slide and incubated at 65°C overnight. Before hybridization, slides were treated with 1mg/ml RNAse A (1:100 from Qiagen) in 2xSSC for at least 45 minutes at 37°C and then dehydrated in a 70%, 80%, and 100% ethanol series for 2 minutes. Denaturation of spreads was performed in 70% formamide/2X SSC solution at 72°C for 1.5 minutes and immediately stopped by immersing slides in ethanol series pre-chilled to -20°C.

The probe for HSAT1B/DYZ2 was generated by PCR using HG002 genomic DNA as a template, KAPA Taq polymerase (Kapa Biosystems), and 50μM Biotin-11-dUTP (Jena Bioscience). Primer sequences: forward - CGCAGCCTAATAACGTGTGGGCTTG, reverse - AATAAAACATAACCATGAAACCTAC. Genomic DNA for PCR was isolated using DNeasy Blood & Tissue Kit (QIAGEN) according to the manufacturer's instructions. The Y centromeric alpha satellite probe (DYZ3) conjugated to TexasRed was from Cytocell, cat# LPE0YcR. The fluorescently labeled oligonucleotide HSAT3/DYZ1 probe 5'FAM-$(AATGG)_7$ was synthesized by IDT.

Labeled DNA probes were denatured in a Hybridization buffer (Empire Genomics) by heating to 80°C for 10 minutes before applying to denatured slides. Spreads were hybridized to probes under HybriSlip hybridization cover (GRACE Biolabs) sealed with Cytobond (SciGene) in a humidified chamber at 37°C for 72 hours. After hybridization, slides were washed in 50% formamide/2X SSC 3 times for 5 minutes at 45°C, then in 1x SSC solution at 45°C for 5 minutes twice, and at room temperature once. Biotin detection was performed using 2 μg/ml Streptavidin - Cy5 (ThermoFisher) in PBST for 3 hours. Slides were then washed in PBST 3 times, rinsed with double deionized H2O, air-dried, and mounted in Vectashield containing DAPI (Vector Laboratories).

Z-stack images were acquired on the Nikon TiE microscope equipped with 100x objective NA 1.45, Yokogawa CSU-W1 spinning disk, and Flash 4.0 sCMOS camera. Image processing was performed in FIJI.

### Comparison to GRCh38Y

#### Y haplogroup identification

Both T2T-Y and GRCh38-Y were aligned to the hg19 ChrY sequence with samtools pileup (Li et al. 2009) to identify SNPs. The software yhaplo v1.1.2 uses phylogenetically significant SNPs to build a tree and then compares that to the ISOGG database to determine the haplogroup (Poznik 2016). The following command was used to determine the haplogroup:

```
yhaplo -i /path/input.txt
```

The Y haplogroup of the 1000 Genomes Project (1KGP) samples were assigned following Poznik et al. (Poznik et al. 2016) if available. For the samples not present in the initial 1,244 set, we used Y-SNP haplogroup hierarchy finder (Tseng et al. 2022) using reliable ("PASS") variants called on GRCh38-Y.

#### Alignments between GRCh38 and HG002 Y assemblies

##### SafFire

Alignments between the GRCh38-Y and T2T-Y assemblies for the purposes of visualization with SafFire were generated with the following minimap2 v2.24 command:

```
minimap2 -t 8 -c --eqx -x asm20 --secondary=no -s 25000 -K 8G \
{input.ref} {input.query} > {output.paf}
```

The PAF was then processed with rustybam v0.1.29 (10.5281/zenodo.6342176) using the following set of commands:

```
rb trim-paf {input.paf}  \
    | rb break-paf --max-size 5000  \
    | rb orient \
    | rb filter --paired-len 100000 \
    | rb stats --paf \
    > {input.for.saffire}
```

and then visualized using SafFire v0.2 (10.5281/zenodo.6376287). DupMasker (Z. Jiang et al. 2008) and dna-brnn (Li 2019) annotations were generated using Rhodonite v0.12 (10.5281/zenodo.6036498).

##### LASTZ

The T2T-Y sequence was aligned to GRCH38-Y using LASTZ v1.04.15 (Harris, Robert S. 2007). Both sequences had been softmasked by the NCBI's assembly submission pipeline using WindowMasker. LASTZ was run in two stages: first to identify filtered ungapped high-scoring sequence pairs (HSPs), second to extend those HSPs to alignments, allowing gaps. Filtering excluded alignments with identity below 80%, ungapped alignments with fewer than 400 matched bases, and gapped alignments with fewer than 1,000 matched bases.

The commands used were:

```
lastz hg002Y.fasta hg38Y.fasta --ungapped --filter=identity:80 --
filter=nmatch:400 --hspthresh=36400 --
format=general-:name1,start1,end1,name2,start2,end2,strand2,nmatch >
hg38Y_onto_hg002Y.anchors
lastz hg002Y.fasta hg38Y.fasta --segments=hg38Y_onto_hg002Y.anchors --
filter=identity:80 --filter=nmatch:1000 --allocate:traceback=800M --
format=general:name1,zstart1,end1,name2,strand2,zstart2+,end2+,nmatch,length1
,id%,blastid% --rdotplot+score=hg38Y_onto_hg002Y.dots > hg38Y_onto_hg002Y.dat
```

Alignments were post-processed to identify, for each position along T2T-Y, the alignment with the highest identity containing that position. This was accomplished using genodsp (http://github.com/rsharris/genodsp) and the following command:

```
cat hg38Y_onto_hg002Y.dat | grep -v "^#" | awk '{ print $1,$2,$3,$10 }' >
hg38Y_onto_hg002Y.identity.dat
```

```
echo "" | genodsp --uncovered:show --precision=3 --chromosomes=hg002Y.lengths
= maxwith hg38Y_onto_hg002Y.identity.dat >
hg38Y_onto_hg002Y.hg002Y_best_identity.dat
```

The alignment dotplot and best identity were plotted using R (https://github.com/arangrhie/T2T-HG002Y/tree/main/alignments/lastz). Regions along T2T-Y were colored according to their class.

Sequence identity was derived from LASTZ alignments, by averaging the identity, over positions that have alignments, of the best alignment through that position. Identity was computed from matches and mismatches only (gaps were ignored): m/(m+mm).

##### Pangenomics Research Tool Kit (PRG-TK)

To visualize the big structural differences of the three ChrY assemblies (GRCh37-Y, GRCh38-Y, and T2T-Y), we use Pangenomics Research Tool Kit v0.4.1 (Chin et al. 2022) to construct the principal bundles that represent the contiguous conserved stretch of sequences among pangenome contigs. Twenty-four contigs over 800 kb from HPRC year one release are used to construct the minimizer anchored pangenome graph with parameters w=128, k=56, r=12, and min_span=28. Two hundred nineteen bundles are identified. In figure **Extended Data Fig. 3c**, we show the sequences of the three assemblies as compositions of the bundles to compare the large-scale rearrangement of the ~30M euchromatin region of ChrY.

#### Gene annotation

##### CAT and Liftoff gene annotation

Because we had gene annotations for CHM13 (Nurk et al. 2022), the CAT annotation was performed as in CHM13, but for the ChrY. Briefly, a Cactus v2.0.5 (Armstrong et al. 2020) alignment to GRCh38 was generated with chimp as an outgroup. Iso-Seq reads were aligned and assembled with Stringtie2 (https://github.com/skovaka/stringtie2, commit 647ab51) (Kovaka et al. 2019, 2), aligned to the assembly with TransMap (as part of Cactus) (Stanke et al. 2008), and used as input for CAT along with the GENCODEv35 (Frankish et al. 2021) annotation.

The following Liftoff v1.6.1 command was run to map genes from the Gencode v35 Y chromosome assembly to the T2T Y chromosome assembly:

```
liftoff v2.7.Y.fasta grch38.fa -g gencode.v35.annotation_Y.gff3 -copies -sc 0.95 -polish
```

The Liftoff output was intersected with the CAT annotation using BEDtools intersect (Dale, Pedersen, and Quinlan 2011) to isolate genes that Liftoff mapped to ChrY that were not in the CAT annotation. Liftoff identified 166 additional genes, of which 152 were extra paralogs of genes already in the CAT annotation and 14 were newly identified.

#### RefSeq gene annotation

The *de novo* annotation of T2T-CHM13v2.0 was performed as previously described for other vertebrate genomes (Rhie et al. 2021; Pruitt et al. 2014), in parallel with the annotation of GRCh38.p14, and released with NCBI *Homo sapiens* Annotation Release v110. The annotation of protein-coding and long non-coding genes was derived from the alignments of RefSeq-curated sequences and primary evidence to the repeat-masked genome. A total of 82,862 curated RefSeq transcripts (with NM_or NR_ prefix), 345,700 cDNAs, 8.65 million ESTs, 9.7 billion RNA-Seq reads, and 83 million PacBio IsoSeq and Oxford Nanopore reads from over thirty distinct tissues were retrieved from SRA and tentatively aligned to the assembly using Splign v2.1.0 (Kapustin et al. 2008) or minimap2 v2.17 (Li 2018, 2). Similarly, 63,836 known RefSeq proteins (NP_ prefix) and 149,352 GenBank proteins were aligned to the genome using ProSplign (NCBI C++ Toolkit r645952).

A total of 20,011 protein-coding genes and 20,716 non-coding genes were annotated on T2T-CHM13v2.0. The vast majority of protein-coding genes (96.3%) and 38.4% of non-coding genes were derived from the placement of curated RefSeq sequences, while the rest were annotated by Gnomon (NCBI C++ Toolkit r645952) based on primary evidence alignments (see details in (Rhie et al. 2021)).

BUSCO v4.1.4 (Seppey, Manni, and Zdobnov 2019) was run in “protein” mode on the longest protein per coding gene. Among the 13780 models in the primates_odb10 lineage dataset, 99.4% were identified as complete (98.8% single copy and 0.7% duplicated copy), 0.1% were found to be fragmented and 0.5% were missing.

#### RefSeq Liftoff gene annotation

The RefSeq Liftoff gene annotation was created by using the Liftoff program v1.6.3, with options -copies -sc 0.95 -polish -exclude_partial -chroms to map across all human genes in RefSeq annotation release v110 from the GRCh38.p14 genome to the T2T-CHM13v2.0 assembly.

#### Ensembl gene annotation

A subset of the genes from GENCODE v38 (Frankish et al. 2021) were mapped to the T2T assembly via a 2-pass alignment process. The subset did not include readthrough genes nor genes on patches or haplotypes, and only one copy of the genes on the ChrX/Y PAR region (only one copy, ChrX, is modeled in the Ensembl representation of the PAR genes).

Firstly, for each reference gene, the sequence of the underlying genomic region was retrieved (including intronic regions), with an additional 500 bp upstream and downstream flanking regions. These were treated as pseudo-long reads and were aligned to the target genome using minimap2 v2.17-r941 (Li 2018) using the ONT settings to allow variability between the reference and target regions. The following command was used:

```
minimap2 --cs --secondary=no -x map-ont [genome_index] [input_file] > [alignment_file]
```

The top hit of the reference region to the target was taken and the equivalent region in the target genome was calculated. If the top hit did not cover the full reference region, the shortfall in coverage was calculated based on the missing 5'/3' sequence and the region in the target genome was adjusted to take the length of the missing sections into account (allowing for issues stemming from gaps/indels or divergent regions).

Once the initial region was identified through minimap2, the two regions were aligned via MAFFT v7.475 (2020/Nov/23) (Katoh and Standley 2014). For each gene, the corresponding exons were retrieved and the coordinates were projected through the alignment of the two regions. Transcripts were then reconstructed from the projected exons. For each transcript, the coverage and identity when aligned to the parent transcript from GRCh38 were calculated.

If the resulting transcript had either a coverage or identity <98%, the parent transcripts were aligned to the target region using minimap2 in splice-aware mode, with the high quality setting for Iso-Seq/cDNA style transcripts enabled. The maximum intron size was set to 100 kb by default, for genes with introns larger than 100 kb in the reference annotation, this value was adjusted to 1.5x their max intron size (to allow some variability).

```
minimap2 --cs --secondary=no -G [max_intron_size] -ax splice:hq -u b [genome_index] [input_file] > [sam_file]
```

For each transcript mapped to the target genome, we then assessed the quality of the mapping based on aligning the original reference sequence with the newly identified target sequence. Again, if the coverage or identity of the aligned sequence was <98%, the reference transcript sequence was re-aligned to the target region using Exonerate v2.0 (Slater and Birney 2005). Exonerate, while slower than minimap2, has the ability to handle very small exons and also can incorporate CDS data to preserve the CDS (introducing pseudo-introns as needed). The following command was used:

```
exonerate -options --model cdna2genome --forwardcoordinates FALSE --softmasktarget TRUE --exhaustive FALSE --score 500 --saturatethreshold 100 --dnawordlen 15 --codonwordlen 15 --dnahspthreshold 60 --bestn 1 --maxintron [max_intron_size] -coverage_by_aligned 1 --querytype dna --targettype [target_type] --query [query_file] --target [target_file] --annotation [annotation_file] > [output_file]
```

Transcripts that still failed the coverage and identity cut-offs using all three approaches were aligned across the whole genome minimap2. This was done to account for rare cases where the target region identification may have been incorrect (perhaps due to a large insertion or an inversion affecting the region). At this point there was generally a single best model for each transcript, defined as the model with the highest combined coverage and identity of all approaches used.

Once the primary mapping process was completed, we searched for potential recent duplications and collapsed paralogues. To search for recent duplications, we took the canonical transcript of each gene (the longest transcript in the case of non-coding genes, or the transcript with the longest translation followed by the longest overall sequence for protein coding genes), and aligned it across the genome using minimap2 in a splice-aware manner. Mappings that overlapped with existing annotations from the primary mapping process on the target genome were removed. For new mappings that did not overlap existing annotations, the quality of the alignment was then assessed by aligning the mapped transcript sequence to the corresponding reference transcript to calculate the coverage and percent identity of the mapping. For these mappings different coverage and percent identity cutoffs were required based on the type of transcript mapped. The cutoffs for retaining transcripts were as follows with %coverage and %identity listed, respectively: protein coding (95, 95), long non-coding (90, 90), small non-coding (95, 95), pseudogene (80, 90).

In order to collapse potential paralogues, where two or more loci in the reference genome map to fewer loci in the target genome, we clustered overlapping genes of the same type (based on exonic overlap) and then collapsed any redundant transcript structures at each locus where the overlap occurred. Non-redundant transcript structures were merged into a single representative gene in the case of protein-coding genes with coding-exon overlap.

Overall, 59,620 of 59,668 (99.92%) genes were mapped from the GENCODE v38 subset to the T2T assembly.

#### Iso-Seq analysis

The generated HQ (Full-length high quality) transcripts, all HQ datasets were mapped to GRCh38.p13 as well as T2T-CHM13v2.0 using three different long read alignment tools: uLTRA v0.0.4.1 (Sahlin and Mäkinen 2021), deSALT v1.5.5 (Liu et al. 2019) and minimap2 v2.24-r1122 (Li 2018, 2). Next, we ran the cDNA_cupcake v28.0.0 (https://github.com/Magdoll/cDNA_Cupcake) workflow to collapse the redundant isoforms from bam, followed by filtering the low counts isoforms by 10 and filter away 5' degraded isoforms that might not be biologically significant.

```
#uLTRA
align --prefix prefix --isoseq --t 4 --index index_dir/
GRCh38.v33p13.primary_assembly.fa HG002.polished.hq.fastq.gz results_dir/
#deSALT
aln -T -o HG002.sam -t 4 -x ccs HG002.polished.hq.fastq.gz
#minimap2
minimap2 -t 8 -ax splice:hq -uf --secondary=no -C5 -O6,24 -B4
GRCh38.v33p13.primary_assembly.fa HG002.polished.hq.fastq.gz
```

To obtain the list of isoforms that are unique to the T2T-CHM13v2.0 genome, we compared the mapping of isoforms between GRCh38 and T2T-CHM13 (https://github.com/unique379r/bioinfo-scripts). We performed this for each replicate of HG002 (NA27730, NA26105 and NA24385). The genomic isoforms as well as ChrY-specific isoforms were generated for further analysis and interpretation. We also generated a merged set of unique T2T-CHM13 isoforms using the agat_sp_merge_annotations.pl script in the AGAT module (https://www.doi.org/10.5281/zenodo.3552717). The program uses the Omniscient parser that takes care of duplicated names and fixes other oddities found across the files. Merged statistics are shown in **Supplementary Fig. 6** for all chromosomes and in **Supplementary Fig. 7** for chromosome Y**.**

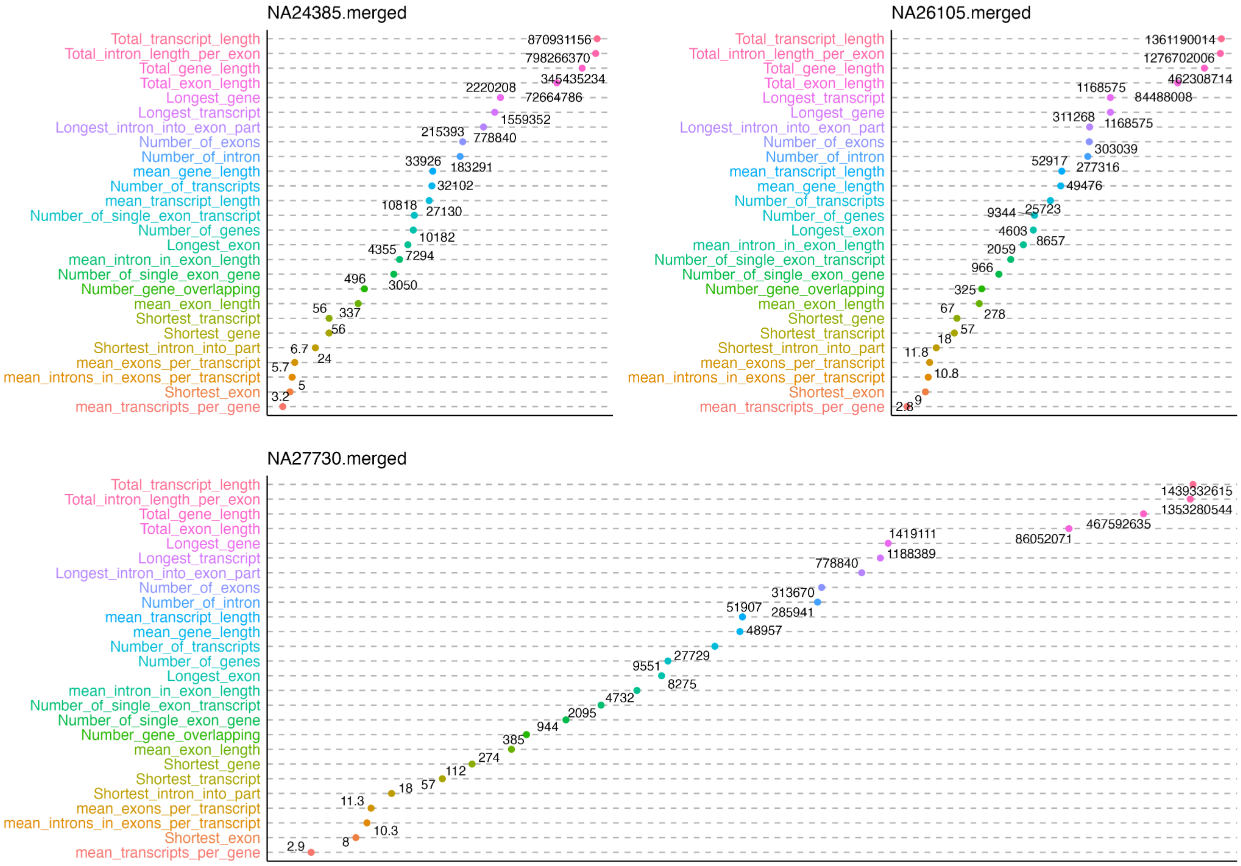


**Supplementary Fig. 6 | Number of isoforms found on all chromosomes across the mapping tools.** Numbers shown are the union set from the 3 different alignment methods.

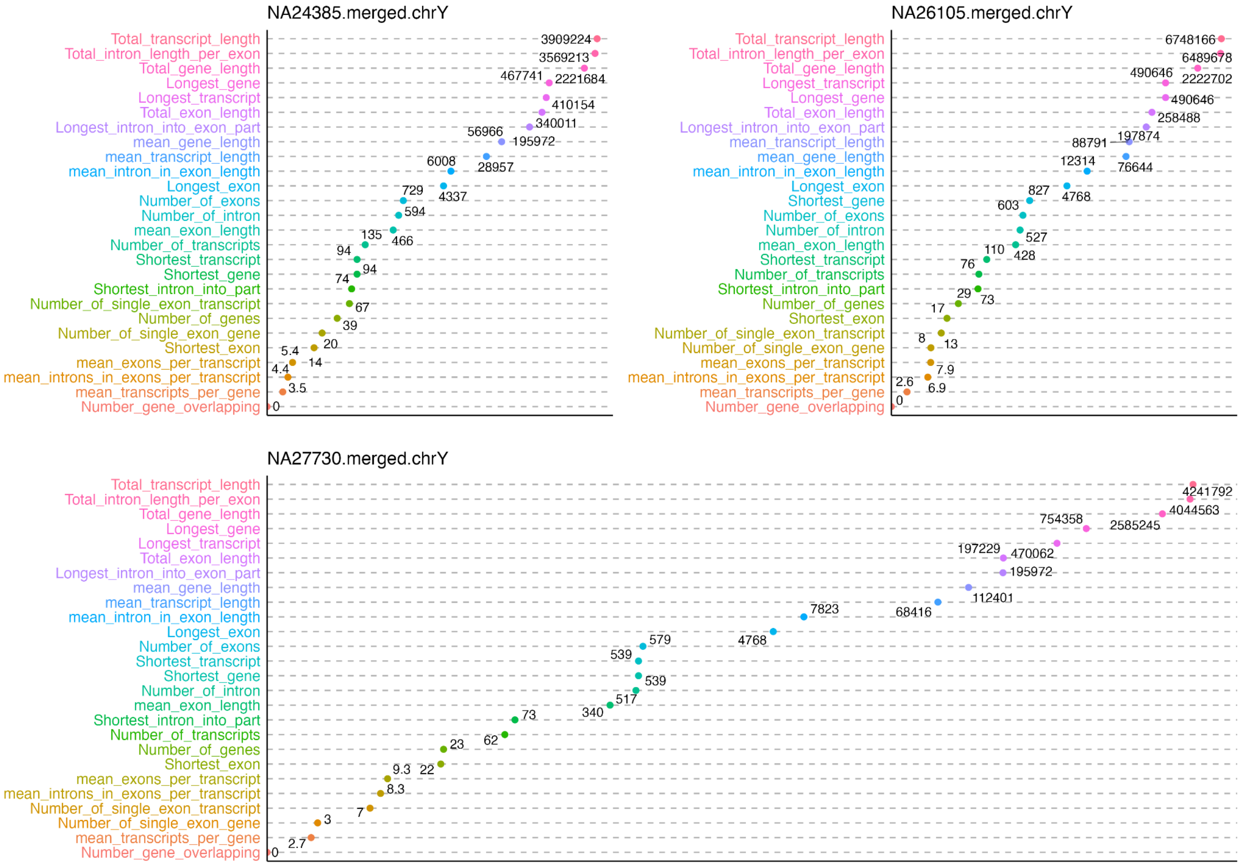


**Supplementary Fig. 7 | Number of isoforms found on the Y chromosome across the mapping tools.** Numbers shown are the union set from the 3 different alignment methods.

#### Ampliconic gene copy number validation

As an additional validation of the T2T-Y assembly, we estimated the copy number of ampliconic genes in the assembly, and compared this to results previously published, for the same individual (Vegesna et al. 2019). The published results had derived estimates from two independent methods: 1) ddPCR and 2) computationally using AmpliCoNE commit c54d9a8, https://github.com/makovalab-psu/AmpliCoNE-tool (Vegesna et al. 2019) on Illumina reads for HG002 from Genome In A Bottle (Zook et al. 2014). To estimate copy number in the assembly, we used AmpliCoNE on simulated Illumina reads extracted from the assembly.

### Repeat annotation

#### Segmental duplications

Segmental duplication (SD) annotations were created using the same methods as in Vollger *et al.* without modification (Mitchell Robert Vollger [2021] 2022). In brief, SDs in T2T-CHM13v2.0 were identified using SEDEF (Numanagić et al. 2018) after repeat masking with Tandem Repeats Finder (Benson 1999) and RepeatMasker (Arian FA, Hubley, Robert, and Green, P 2015). The code is deposited on Zenodo (https://zenodo.org/record/5499093) and made available as a Snakemake pipeline.

#### Repeat model discovery and annotation

##### Repeat model discovery with RepeatModeler and loci identification with RepeatMasker

To assess previously unannotated repetitive regions of ChrY, a RepeatMasker4.1.2-p1 run was completed on the T2T-Y assembly using the Dfam 3.3 library (J. Storer et al. 2021) with the following settings: sensitive setting (-s), using the species tag of human (-species human) and the NCBI BLAST-derived search engine RMBlast (-e ncbi): $ RepeatMasker -s -species human -e ncbi. These regions were then hard-masked and a RepeatModeler2.0.1 analysis was performed on the remaining (unmasked) regions. The resulting repeat model consensuses were extended and subjected to two subsequent levels of filtering:

1. Removal of duplicate models based on a 2% divergence value using a custom perl script
2. Removal of models corresponding to repeat families already found in the Dfam database using cross_match analysis

Since the RepeatModeler2.0.1 algorithm implements a random sampling of the genome, the remaining repeat model consensuses were used as a library for a secondary RepeatMasker run to collect all associated instances for each model across the T2T-Y assembly.

##### Discovery of new satellite models

Satellites and tandem repeats were initially annotated in the above RepeatMasker run, based on a combination of alignments to known satellite sequences in the RepeatMasker library and associated screening with Tandem Repeats Finder (TRF) (trf409.linux64). While RepeatModeler2 identified some new satellites, it is specifically designed for interspersed repeats rather than tandem repeats and therefore, large sections of chrY were left unannotated for tandem repeats that may actually be present.

We expanded annotation coverage of these missing repetitive regions using two methods: ULTRA v1.0 (D. Olson and Wheeler 2018) and NTRprism v1.0.0 (Altemose et al. 2022). The first, ULTRA, is an open-source tool that can annotate and provide statistically consistent scoring for very large repeat units (up to a repeat period of 4000), arbitrarily-long repetitive regions, and ancient repeats that have highly decayed repetitive signals. ULTRA v1.0 was run with a repeat periodicity of 1000 on the unmasked ChrY. The second, NTRprism, creates a Nested Tandem Repeat (NTR) “spectrum” indicating the most abundant tandem repeat periodicities across a given sequence. The higher the score the more likely they are to be present in higher copy numbers and with higher homogeneity. NTRprism v1.0.0 was run in 50 kb bins with a 0.01 column sum score threshold on the unmasked ChrY.

We focused on unannotated regions/gaps greater than 5 kb, which were identified via BEDtools v2.29.0 (Quinlan and Hall 2010) by subtracting both repeat annotations (first: using Dfam library, second: using pre-filtered RepeatModeler consensuses) from the entire T2T-Y sequence. These gaps were manually curated in a UCSC Genome Browser session to check for any feature annotation overlap (e.g. gene annotations). Tandemly repeated sequences for each gap were detected and assessed with a combination of ULTRA, NTRprism, and the TRF v4.09 GUI version using default parameters (Benson 1999) to determine the best monomer consensus for a given satellite model.

The repeat discovery and annotation pipeline was originally run on the entire CHM13 genome, including ChrX and its PAR regions. When running the pipeline on T2T-Y alone, we detected satellites (arrayed tandem repeats) in the PAR regions of ChrY that we had not annotated in CHM13. Following re-annotation of CHM13 and HG002-X with the additional ChrY-derived repeat models, these ChrY-derived satellites were in fact found in the ChrX PARs, as expected (**Supplementary Table 15**). There are a few reasons these ChrY-derived satellites were undetected in the initial screen of CHM13, including a combination of the following:

1. RepeatModeler is designed to detect interspersed repeats, not tandem repeats present at a single locus and/or with low copy number or high sequence variability
2. The models ascertained from RepeatModeler are derived from a random sample of the genome, and therefore the output of every program run is slightly different
3. RepeatModeler and TRF are both more sensitive when run on a smaller sequence, such that the sample of the genome reflects a higher portion of the total sequence, so more satellites were detected when run on the single ChrY compared to the full CHM13 genome

Note, however, that regardless of these considerations, RepeatModeler should *not* be run in parallel on different chromosomes, as the models this practice would produce would not accurately represent the models in the genome as a whole.

##### Manual curation of previously unknown repeat models

Following the production of a secondary RepeatMasker annotation set (using the pre-filtered RepeatModeler consensus), curation steps were implemented to refine previously unknown repeat models. Multiple sequence alignment (MSA) plots of the RepeatModeler consensus sequences aligned to the T2T-Y assembly were used to assess the divergence across associated instances. Overlaps with CAT/LiftOff gene annotations, segmental duplications, and tandem repeats found within gaps were manually curated. RepeatModeler consensuses were screened as potential composite subunits through pattern recognition in both the UCSC browser and RepeatMasker output, while Bedtools closest (-k 2 -iu -io -D ref) and (-k 2 -id -io -D ref) was used to assess the neighboring repeats and their frequency increasing the likelihood that they were part of a larger repeat, or composite. This curation led to the generation of a final repeat library of previously unknown or unannotated satellite monomers (n=14) and subunits of composites (n=15) (**Supplementary Table 11**).

##### Compilation and polishing of T2T-Y repeat annotations

The compilation pipeline laid out in Hoyt *et al.* (Hoyt et al. 2022) was followed to avoid potential false positives by simply masking with a combined library of new repeat models and known repeat models (Dfam library). The pipeline involves a third RepeatMasker run on the original hard-masked ChrY (masked using Dfam library only) using a library which included the Dfam database plus all additional entries (new models from T2T-CHM13 and these ChrY analyses). This third repeat annotation and the first Dfam library only repeat annotation were then processed through a final processing script resulting in a high confidence repeat masker annotation track for the UCSC genome browser.

The direct activity of Transposable Elements (TEs) over long stretches of evolutionary history accounts for a large proportion of the human genome. The annotations of TEs form a fossil record whereby the recursive insertion of TEs within other TEs can be disentangled and visualized. In 2015, we developed a specialized visualization for the UCSC genome browser (Rosenbloom et al. 2015) to specifically address the hierarchy and fragmentation present in TE annotations. The annotation glyphs provided a visual language for simultaneously representing the relative position of fragments of a single TE insertion within the genome and the relative location of these fragments within the sequence of the full-length TE family. Our track was designed to work for UCSC hosted genomes only, and was initially applied to two assemblies (hg38 and mm10). For the T2T and Dfam (J. Storer et al. 2021) projects we developed a trackHub-aware version of our visualization with expanded features for managing the visualization and optionally filtering results. The TE annotations on the T2T assemblies were prepared using a new tool from the RepeatMasker 4.1.3 package (rmToTrackHub.pl) to generate a trackHub for the visualization.

The same three-step repeat annotation pipeline (1. RepeatMasker (Dfam library only), 2. RepeatMasker (Dfam library + new repeat models), 3. compilation pipeline) was applied to GRCh38-Y as well. Repeats were summarized using buildSummary.pl (J. M. Storer et al. 2021) at the class and family level (**Table 1**, **Supplementary Table 12**) and at the subfamily level for new repeats (**Supplementary Table 11**) in both T2T-Y and GRCh38-Y.

#### Composite repeats

Composite elements were defined and characterized as described in Hoyt et al. 2022 (Hoyt et al. 2022) as: a repeating unit consisting of three or more repeated sequences, including TEs, simple repeats, composite subunits, and/or satellites, that is found as a tandem array in at least one location in the genome.

Following the generation of our high confidence repeat masker track with the inclusion of composite subunits, three composite repeats were identified, each of which associates with a gene: *TSPY*, *RBMY*, and *DAZ*. The *TSPY* and *RBMY* composite units are structured with one gene per composite unit, so that an array of composite units includes multiple genes. The *DAZ* composite units are different in that an entire array falls within one gene. BLAT v36.5 (Kent 2002) was used to locate other composite unit copies across HG002 chrY and cross-reference them with their associated gene annotations (CAT/liftoff). **Supplementary Table 18** only reports those copies that are associated with genes (protein-coding or pseudogenes), but fragmented and/or diverged copies were also found without genes for *TSPY* and *RBMY* (not *DAZ*). Generation of the composite consensuses was completed by searching HG002-chrY sequence for full-length composite insertions via BLAT with an exemplar locus as the search query. The insertions were then iteratively aligned using alignAndCallConsensus.pl (J. M. Storer et al. 2021) to the exemplar locus, producing a consensus sequence for each of the three composites. There was sequence diversity of *DAZ* composites across the gene family, with the main diversity occurring within *DAZ3* and *DAZ4*, but a single consensus was sufficient to find the copies, so a single consensus was included.

##### Identification of full-length TEs

The active families in the human genome for SINEs, LINEs, and retroposons are AluY, L1Hs, and SVA_E/F, respectively; the recently active family in the human genome for ERVs is HERV-K. Identification of these potentially active, full length TEs across T2T-Y and GRCh38-Y was done by following the methods laid out in Hoyt et al. 2022 (Hoyt et al. 2022). Full length elements and ERV structural category counts and locations can be found in **Supplementary Table 12**.

##### T2T-Y liftOver analysis

Liftover chains were generated from Minimap2/NEXTflow (See below "**Curated liftover chains**") sequence alignments between GRCh38-Y and T2T-Y. A bed file was generated from the T2T-Y RepeatMasker output and a liftOver (https://genome.ucsc.edu/cgi-bin/hgLiftOver) was performed to the GRCh38-Y assembly. Subsequent analyses were performed on both the unlifted and lifted TEs as laid out in Hoyt et al. 2022 (Hoyt et al. 2022) with the exception of identification of liftover errors instead of polymorphisms. The chm13v2-unique_to_hg38.bed (https://s3-us-west-2.amazonaws.com/human-pangenomics/T2T/CHM13/assemblies/chain/v1_nflo/chm13v2-unique_to_hg38.bed, as plotted in **Fig. 1**) was used to define syntenic regions with GRCh38. Full stats are provided in **Supplementary Table 13**, with data summarized in **Supplementary Fig. 8.**

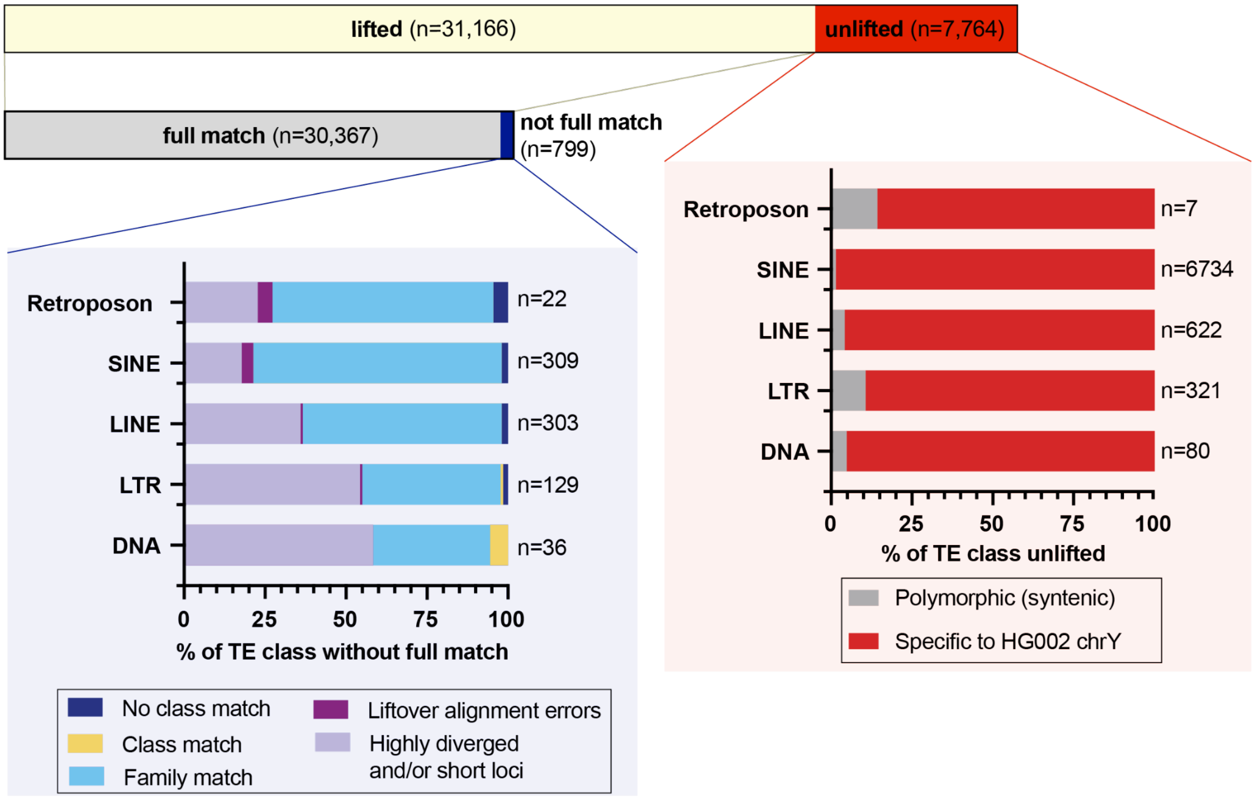


**Supplementary Fig. 8 | TE liftover between T2T-Y and GRCh38-Y.** A minority of T2T-Y TEs (~20%; Top - red) were unable to be lifted to GRCh38. (Red-boxed highlight) Stacked bar plot showing percentage of TEs by class (DNA, LTR, LINE, SINE, and Retroposon) that were unlifted from T2T-Y gap-filled regions (non-syntenic, red) versus syntenic regions (gray). Out of the TEs that were able to be lifted (top, yellow), a minority did not have a full match (~3%; middle, blue); a full match being no change in annotation. (Blue-boxed highlight) Stacked bar plot showing percentage of lifted TEs by class broken down further into discordance categories as follows: 1) no TE class match (dark blue), 2) class match, but family change (yellow), 3) family match, but subfamily change (light blue), 4) liftover alignment errors (dark purple), and 5) highly diverged sequences and/or short fragments (light purple). The first two categories (dark blue and yellow) encompass the most notable differences in lifted TE annotations between T2T-Y and GRCh38-Y. Number of TEs per category shown across figure as *n*. Full stats included in **Supplementary Table 13**.

#### Satellite annotation

Centromeric Satellite (Cen/Sat) annotations were generated as in (Altemose et al. 2022), with a few refinements tailored to ChrY satellites. Rather than restrict the satellite annotation to the region surrounding the centromere, we annotated satellites across the entire chromosome. First, major satellite types were extracted from the RepeatMasker tracks generated for this chromosome, merging features of the same satellite type within 10 kb of each other. However, for HSat2 and HSat3, a specialized annotation tool was used (https://github.com/altemose/chm13_hsat, from (Altemose et al. 2022). Alpha satellite annotations were replaced with the more exact annotations described below. DYZ19 was annotated by RepeatMasker as a 265 kb LTR12B element, consistent with the derivation of its 125 bp repeat from an LTR, as described by (Skaletsky et al. 2003). HSat1B was defined by merging RepeatMasker annotations “HSATI”, “AT-rich”, and “AluY” in the Yq region. Because there were frequently small gaps in RepeatMasker annotations between contiguous blocks of

HSat3 and HSat1B (aka DYZ1 and DYZ2), the exact boundaries between these blocks were later refined manually using alignments to consensus sequences as a guide. Other CenSat track boundaries were manually refined to eliminate overlaps. The "pericentromeric region" was defined at the boundaries of the alpha satellite HOR array ±5 Mb. Within this region, any gaps between satellite annotations were labeled as "ct" for "centric transition."

#### Cytoband annotation

The cytoband track for T2T-Y was produced by first using liftOver on the cytoband track from hg38, but only 4 of the band boundaries lifted over properly (q11.21-q11.221, q11.221-q11.222, q11.222-q11.223). The remaining band boundaries were placed either using the CenSat annotation (described below) or by taking the 1 kb sequence at the boundary from GRCh38 and using BLAST to locate it in T2T-Y (for p11.32-p11.31, p11.31-p11.2, q11.223-q11.23). The p11.2-p11.1, p11.1-q11.1, and q11.1-q11.21 boundaries were placed at the beginning, midpoint, and end of the alpha satellite HOR array, respectively. The q11.23-q12 boundary was placed at the proximal end of the DYZ18 (HSat3) array.

#### Transduction analysis

We utilized the same approach as Hoyt *et al.* (Hoyt et al. 2022) to identify putative DNA transductions mediated by retroelements. Briefly, 100 bp upstream and 3 kb downstream of L1s and SVAs annotated in T2T-Y were searched to detect the target site duplications (TSD) and 3' transduction signatures (for 5' SVA transductions, 3 kb upstream of the elements was investigated) using a modified version of TSDfinder (https://github.com/IOB-Muenster/TSDfinder v1.0) (Szak et al. 2002). Then, we removed transductions residing in segmental duplications and masked the transduced sequences using RepeatMasker v4.1.2-p1 (Arian FA, Hubley, Robert, and Green, P 2015) with -q -species human -xsmall parameters. To find the potential progenitor of each transduction within T2T-CHM13v2.0 and GRCh38, the offspring sequences were aligned to the corresponding databases containing 3 kb sequences downstream of full-length L1s and SVAs (as for 5' SVA transductions, 3 kb upstream of all SVAs were used regardless of their length) using BLAST v2.11.0 (Altschul et al. 1990) with the following command:

```
blastn -query transducedSequences.fa[masked] \
  -db 3KbDownStreamFullLengthRetroTEs.fa \
  -task blastn -evalue 0.05 -max_target_seqs 5 -perc_identity 90
```

Next, a locus was considered a likely source if all of the following criteria were met in the BLAST results: a) hit and subject had the same orientation, b) query and subject were at least 90% identical, c) at least 30% of the query length was aligned, and d) the start coordinates of each query-subject were within 20 bp of each other. Finally, to produce the final call set, the remaining transductions were subjected to manual curation.

#### Non-B DNA motif annotation

To predict sequence motifs with the potential to form alternative DNA structures (non-B DNA), we used nBMST (https://github.com/abcsFrederick/non-B_gfa commit 1c8f963) (Cer et al. 2012) for repeat motifs (A-phased, direct, inverted, and mirror repeats and STRs) and Z-DNA motifs and we removed motifs with a spacer larger than 15 bp (Zou et al. 2017; Svetec Miklenić et al. 2020). We used Quadron with default parameters to detect G4-motifs (https://github.com/aleksahak/Quadron commit 19047e3) (Sahakyan et al. 2017), which also yields a score that predicts the stability of a predicted G4 structure, based on a machine-learning algorithm using empirical datasets. Motifs with a Quadron score ≥19 are considered stable, and those with <19 unstable, respectively. We then intersected this non-B motif annotation with other existing annotations of T2T-Y (gene annotations, satellite repeats, and CpG islands) using *bedtools intersect* (Quinlan and Hall 2010).

### Data visualization

For **Figure 1**, alignment of GRCh38-Y and T2T-Y visualized with SafFire (Mitchell Robert Vollger [2021] 2022). Segmental duplications (SDs) are colored by duplication types defined in DupMasker (Z. Jiang et al. 2008). IGV v2.14.1 was used to draw ideograms, sequence classes, palindromes, inverted repeats, and AZF annotation from **Supplementary Table 21-22**. BEDtools v2.29.0 (Quinlan and Hall 2010) map was used to calculate density (-o count) per 100 kb window across each gene type: protein-coding and pseudogenes (based on CAT/Liftoff annotations). BEDtools coverage was used to calculate bp coverage per 100 kb window across each repeat class from RepeatMasker to represent repeat density across T2T-Y. Similarly, BEDtools coverage was used to calculate bp coverage per 100 kb window for non-B DNA motifs. BEDtools map was used to calculate average (-o mean) methylation frequency per 100 kb window (based on HG002 Nanopore and HiFi methylation data). Rideogram v.0.2.2 (Hao et al. 2020) was used to generate these visualized tracks as well as the three composite repeat tracks. GraphPad Prism v9.1.0 ("GraphPad Prism Version v9.1.0 for Windows, GraphPad Software, San Diego, California, USA. Last Accessed: 2022-11-28." n.d.) was used to generate the TE composition per sequence class plots (corresponding to **Supplementary Table 14**, **Extended Data Fig. 4**, and **Supplementary Fig. 6**, respectively).

### TSPY gene family analysis

#### TSPY copy number estimation from SGDP

Copy number estimates of the TSPY gene was performed as in Vollger et al, 2022 (Mitchell R. Vollger et al. 2022). In brief, we applied the fastCN v0.2 pipeline (Pendleton et al. 2018), which uses sequence read-depth as a proxy. Short-read sequence data were processed into 36 bp non-overlapping fragments and mapped to a masked T2T-CHM13v2.0 reference using mrsFAST v3.4.2 (Hach et al. 2010) with a maximum of two substitution mismatches not allowing for indels. Masking was determined by TRF v4.09 and RepeatMasker v4.1.2-p1. Read-depth across the genome was corrected for GC bias and copy number was determined using linear regression on read-depth versus known fixed copy number control regions. Finally, integer genotypes for *TSPY* were generated by taking a weighted average of the copy number estimates from windows overlapping the locus. A snakemake pipeline for creating this reference and processing short reads into copy number can be found here: https://github.com/mrvollger/fastCN-smk (doi:10.5281/zenodo.8136270).

##### Phylogenetic tree analysis of the TSPY genes

To understand the relationship between the *TSPY* gene copies across T2T-Y, a phylogenetic analysis was performed. All curated protein-coding and pseudogene *TSPY* copies (including introns) from the CAT/Liftoff and RefSeq/Liftoff annotations were used. For outgroup rooting of the tree, *TSPY* sequences were used from *Hylobates moloch* (accession # NW_022611649.1) (Escalona et al. 2023) and *Pongo abelii* (accession # KP141780.1) (Cortez et al. 2014). Alignment was carried out in MAFFT v7.471 (Katoh and Standley 2014) using custom parameters based on manually inspected alignment results, primarily an iterative alignment with the L-INS-i method. Phylogenetic analysis was run in RAxML-NG v0.9.0 (Stamatakis 2014) using the GTR+I+G model based on 10 parsimony-based starting trees. Rapid bootstrap approximation was run to generate 200 bootstrap replicates using the University of Connecticut Health Center server. Consensus bootstrap values were then mapped to the highest likelihood phylogeny in Geneious v2019.2.3 (Geneious n.d.) and visualized in FigTree v1.4.4("FigTree. Last Accessed: 2022-11-28." n.d.).

#### Centromere analysis

##### HOR haplotype and SVs

###### HG002 cenY analysis

The HG002 T2T-Y assembly was processed using the standard alpha-satellite (AS) tools as described in Altemose et al. (Altemose et al. 2022) and the standard panel of UCSC Browser tracks was built as follows (**Supplementary Fig. 9**):

1. The SF-track shows alpha-satellite (AS) monomers indicating to which AS supra chromosomal family (SF) each monomer belongs.
2. The HOR-track shows all monomers which belong to AS higher-order repeats (HORs) and indicates which monomer of which HOR each one is.
3. The structural variant, or StV-track shows each copy of the S4CYH1L HOR in the cenY active array and indicates whether each copy is a regular complete copy or deleted/rearranged.
4. The AS strand track shows whether AS runs along the forward or reverse strand, indicating occasional inversions and/or different AS direction in different sequence domains.

These tracks can be viewed in the UCSC Human Genome Browser: https://genome.ucsc.edu/s/fedorrik/T2T_dev

a

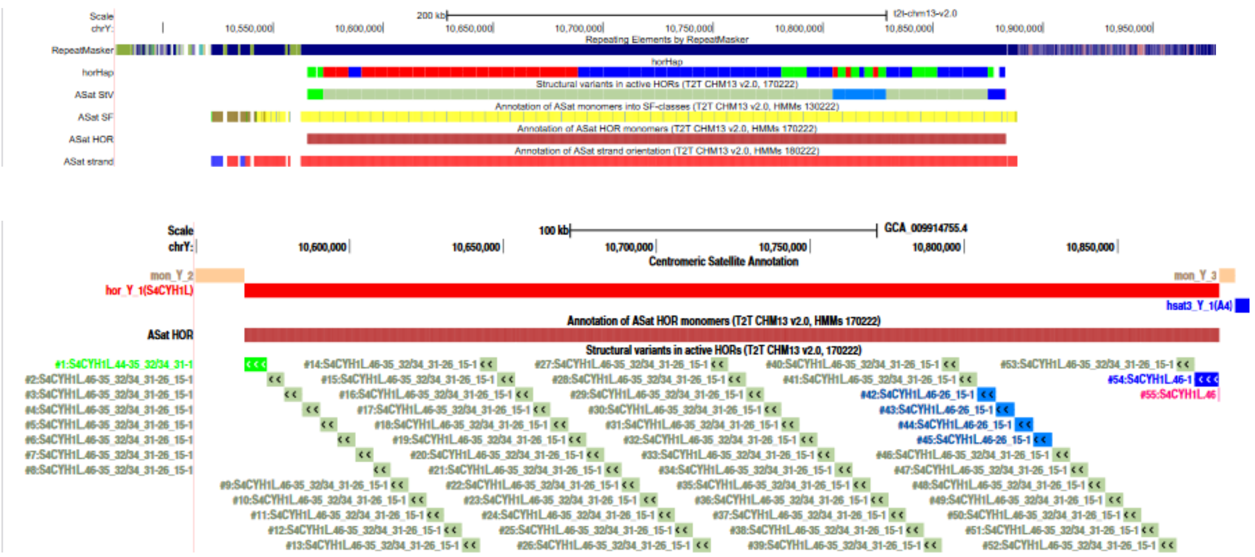


**Supplementary Fig. 9 | Representation of HG002 active HOR array structure, HOR haplotypes and SVs in UCSC Browser tracks. a.** The tracks listed in the section above are shown in “dense” mode; **b.** The SV track is shown in “pack” mode which shows every HOR copy with numbers. 34mers are gray, 36mers are light blue, 44mer is green and the complete 46mer is dark blue.

The S4CYH1L (DYZ3) AS HOR was re-examined and re-defined for this paper to take into account its polymorphic variants both known from the old literature and revealed by the recent complete assemblies. In the previous definition, the HOR which belonged to SF4 (Shepelev et al. 2015) consisted of 34 monomers present in the most numerous and common HOR variant (Tyler-Smith and Brown 1987). The longer and much less frequent version of the HOR known from the old literature (Tyler-Smith and Brown 1987) and found in HG002 has 36 monomers, so that the 34mer may be considered a deleted variant of this 36mer. Finally, at both flanks of cenY arrays in both HG002 and RP11 (see below), even longer HOR copies are present (one copy on each side) which have 10 additional monomers (42mer on the left and 46mer on the right). The 46mer was judged to be the longest evolutionarily-relevant variant and was used as a canonical HOR with its monomers numbered from 1 to 46, and the shorter variants were considered to be deleted variants of this canonical HOR. Commonly in indel variants, the shorter versions feature various hybrid monomers where the in/del border does not align with an arbitrary monomer start site (see (Uralsky et al. 2019; Altemose et al. 2022) for details). Then the structure of 34mer would appear as S4CYH1L.1-15_26-31_32/34_35-46 (notation as in (Altemose et al. 2022)), the structure of 36mer as S4CYH1L.1-15_26-46, the structure of 42mer as S4CYH1L.1-31_32/34_35-44, and the complete 46mer as S4CYH1L.1-46 (**Supplementary Fig. 10**). Monomer S4CYH1L.32/34 is a hybrid which has a part of monomer 32 on the left end and a part of monomer 34 on the right end, and is the result of the deletion of two monomers in a canonical sequence. Note that in cenY the HOR array runs along the reverse strand, so if the monomer order in the track is read in a standard manner (from left to right) the monomers would appear in the reverse order (from 46 to 1). Finally, on the extreme right flank, a solitary monomer 46 (1mer) followed the complete 46mer. The HORs were numbered from left to right and the complete array had 55 HORs as follows: 34mer - 48, 36mer - 4, 42mer - 1, 46mer - 1, 1mer - 1. The order of HORs can be seen in the StV track. The resulting structure of the AS array in HG002 cenY was revealed as shown in **Supplementary Fig. 9**.

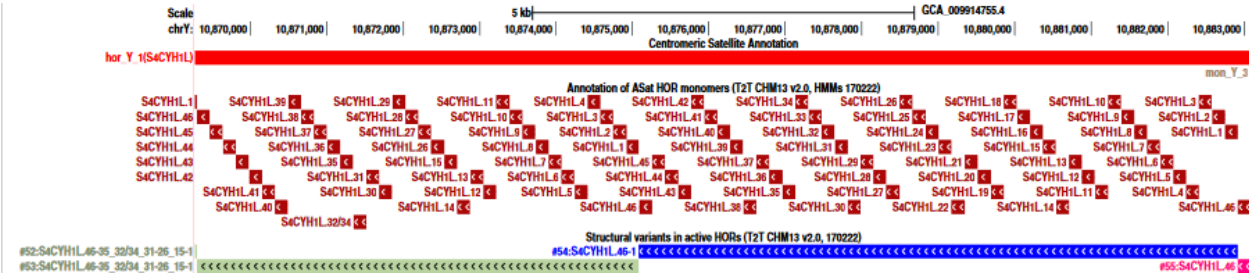


**Supplementary Fig. 10 | The new definition of S4CYH1L (DYZ3) HOR**. The figure shows the right flank of the HG002 active HOR array. As the HORs run in reverse direction, the monomer numbers should be read from right to left. The leftmost HOR (#55) which has only one monomer (monomer 46), the next HOR (#54) is full-length (monomers 1 through 46), HOR#53 is 34mer with 15_26 junctions and 32/34 hybrid.

A summary of the HG002 cenY HOR structure is as follows:

1. The HOR array runs along the reverse strand, as well as the flanking SF4 non-HOR sequences, but two small pieces in the left non-HOR flank are inverted and go in the forward orientation (**Supplementary Fig. 9a**).
2. The left non-HOR AS flank has ~60 kb of AS and is made of SF4 (yellow) and SF6 (brown) in about equal proportions. It is disrupted by multiple L1 elements and by a large (114 kb) chunk of the non-AS sequence which is a part of SD that also overlaps almost all the left non-HOR AS flank. The region is shared to various degrees with acrocentrics (mainly with cens 14 and 15, but also cens 9, 20, etc.). The inverted regions are mostly covered by SDs. The small chunk of SF4 monomeric AS not covered by SD and adjacent to the HOR array has insertion of a cluster of L1PA3 fragments, which has a small piece of AS inside.
3. The right non-HOR flank is just ~5 kb long, is made of SF4 and is disrupted only by a single Alu repeat at chrY:10883788-10884095 (T2T-CHM13v2.0 coordinates here and onwards).
4. AS in the arms. There is possibly a very small (42 bp) isolated chunk of an SF4 (Ga) monomer in the long arm at chrY:17477617-17477658 and a chunk of an ancient Ia monomer at chrY:240673-240779, in the short arm. Also, there is a large (994 kb) AS-containing SD at chrY:18,362,322-19,356,689. The latter has the red, orange and lilac ancient AS families (Ca, Ba and Ja, respectively) totaling ~50 kb disrupted by multiple TEs and is shared with cen 12 which is probably a home site of this sequence, as it is included in the broader AS context there. The identities with cen12 are low-end and only partially recognized in the SD track. The SD is a palindrome, so the AS is represented by 2 symmetrical and almost identical pieces in opposite orientations, ~50 kb long each.
5. StVs. The HOR array occupies the region chrY:10565751-10882579 (316,829 bp) and lists 55 HORs (including the one at the right end containing just one monomer). 48 of these

are the predominant 34mers, and the two flanking long HORs (42mer on the left and a complete 46mer on the right) were described above. That leaves four copies of 36 mer S4CYH1L.46-26_15-1 (HORs 42-45) which are located as a single tandem cluster at chrY:10804299-10828806. This could be an island of the older (since it is longer) cenY HORs which represents the second generation of the HOR evolution by sequential truncation, the first being the long HORs of the flanks and the third being the classical 34mers. Thus, the story starts from the 46mer, which first lost 10 monomers to produce the 36mer, and then another 2 monomers to produce 34mer.

6. HOR-haps. We have extracted all 54 HORs from the array and built the whole-HOR phylogenetic trees, as described in Altemose 2022 (Altemose et al. 2022), to assess HOR haplotypes. We were able to sort the HORs into 3 main haplotypes (**Supplementary Fig. 9a**), and 32 full-length 34mers (**Supplementary Fig. 9b**) were used to construct a HOR-hap HMMER-based tool as described in Altemose *et al.* (Altemose et al. 2022) for cenX. The HOR-haps predicted by this tool in cenY are shown in the HOR-hap track (**Fig. 3**). Briefly, there are two main domains, the red on the left and the blue on the right of about equal length. Additionally, the 36mers and the four HOR copies located symmetrically around the 36mer island (two on each side) apparently belong to the third HOR-hap (green). Next, the consensus HORs were generated from each set of HORs (**Supplementary File 1: Y_HOR-hap cons**) and compared. It appears that there are two major haplotypes, red and green-blue, and two less distinct sub-haps, green and blue (**Supplementary Fig. 9c**). Only a few diagnostic positions discriminate the 3 HOR-hap consensus sequences as follows (the HOR-hap which differs from the other two is shown in parenthesis): positions 219 (red), 759 (red), 2367 (blue), 3011 (red), 3063 (green), 3592 (green), 3762 (green), 3890 (blue), 5722 (red) and 5747 (blue). The intra-array average divergence in the green HOR-hap is dramatically higher (1.5%) than in red and blue HOR-haps (0.35% and 0.32%, respectively), which suggests it is the ancestral variant from which first the blue 34mer and then the red 34mer were generated. This aligns with the fact that the sequence of 36mers matches the green HORhap.

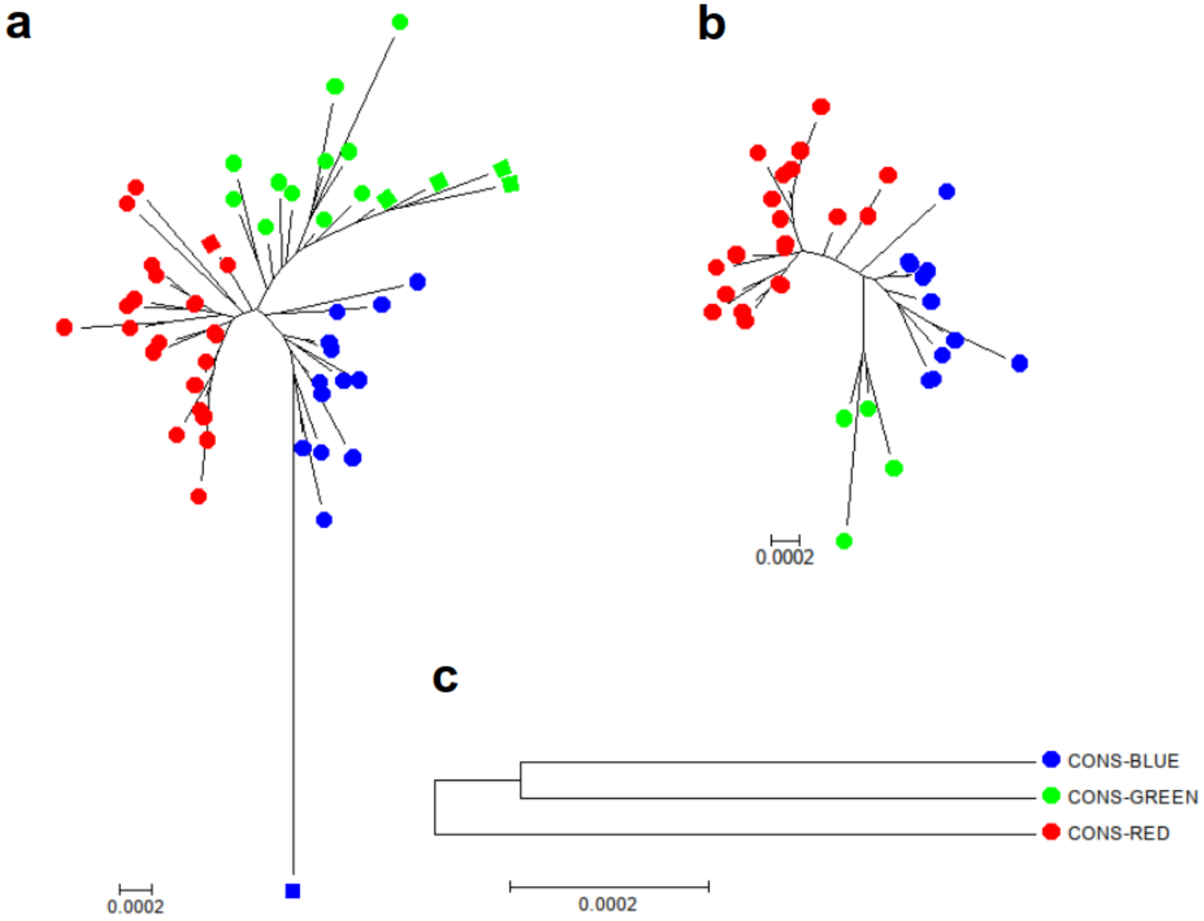


7.

**Supplementary Fig. 11 | HOR haplotype analysis in HG002. a.** A complete minimum evolution tree showing 54 HORs, which were cut to align with 34mer was built using the "pairwise deletion option" in MEGA5. The flanking long HORs and 36mers are marked by squares (44mer is red, 46mer is blue and 36mers are green). Assignment of the 44mer to the red branch indicates its complex rearrangement (see 4 in the RP11 section). **b.** This tree shows only the complete 34mers which were used to build the HMMER-based HOR-hap tool. A number of HORs which sat close to the root branches were filtered out because they could have been hybrid HORs which would obscure the classification. This tree is shown in **Fig. 3** in the main text. **c.** Minimum evolution tree of HOR-hap consensus sequences made from the dataset shown in **b**.

#### RP11 cenY analysis

The RP11 chrY centromere was described in Jain et al. 2018 (M. Jain, Olsen, et al. 2018). Its main contig is MF741337 but it lacks the AS edges of the centromere. Therefore we have merged additional contigs to it as follows: ULGL01000015.1[1 to 331091] + MF741344[136833 to 85827] + MF741337[1 to 280254] + MF741339[115015:end].

The RP11 cenY assembly (four contigs) seems to have some SNP-like errors. We have extracted the HOR sequences of RP11 and HG002 and built a multiple alignment. There are six positions where all the HG002 HORs have one nucleotide and all the RP11 HORs have another nucleotide or deletion. Three out of these six positions (1672, 1441, 162) are in areas where one nucleotide repeats several times and RP11 repeats are one nucleotide shorter than HG002 repeats. Positions 5808 and 7408 are single-nucleotide substitutions and position 6170 is a deletion in the RP11 HORs.

To check if these differences were real, we have created k-mers including these positions for both RP11 and HG002 variants and checked for them in the illumina WGS reads of RP11 (SRR6819611). The oligos from RP11 have 0 or 1 hits while the oligos from HG002 have hundreds of hits (**Supplementary Table 19**). So, we consider these positions to be errors in the ONT consensus calling which is not unexpected, since the work in Jain et al. 2018 (M. Jain, Olsen, et al. 2018) was done during the early days of long-read sequencing and used only the sequences of the whole BACs sequenced in a single read for consensus calling. Since such long reads were infrequent, the sequence was most likely not well polished. Therefore, we chose not to use the RP11 sequence for full-fledged analysis similar to the one we did for HG002, but only annotated the sequence with the tools developed for HG002.

The RP11 assembly was studied the same way as reported for HG002 above and the same set of tracks was generated (**Supplementary Fig. 10**). The HORhaps were established using the HG002-based tool. The RP11 cenY tracks can be viewed here: https://genome.ucsc.edu/s/fedorrik/rp11.cenY

Note that the StVs in the RP11 active array were previously partially studied in Jain et al. 2018 (M. Jain, Olsen, et al. 2018), but this study did not include the extreme flanks of the active array. Also, Vlahović et al. 2019 (Vlahović, Glunčić, and Paar 2019) have studied the StV content of cenY in the hg38 assembly which included the cenY reference model derived from HuRef (Miga et al. 2014) and some flanking traditional contigs derived from RP11.

A summary of the RP11 HOR structure is as follows:

1. The HOR array runs along the reverse strand, as well as the flanking SF4 non-HOR sequences, but two small pieces in the left non-HOR flank are inverted the same way as in CHM13.
2. The left non-HOR AS flank seems to be structured the same way as in T2T-Y (HG002).
3. The right non-HOR flank seems to be the same as in T2T-Y.
4. StVs. The long HOR on the left flank is different in that it is a HOR with monomers 1-44 (i.e. a complete HOR just missing the two monomers at the end). In HG002, it was a HOR with a 2-monomer deletion (marked by 32/34 hybrid and the complete absence of mon 33) typical of the 34mer. Its structure was S4CYH1L.1-31_32/34_35-44. This suggests that the 44mer left flank HOR in RP11 has the unperturbed ancestral structure and the left flank 42mer in HG002 is somehow rearranged in an unclear manner. Also, the 36mers are completely absent in RP11, instead there are nine 35mers with duplicated monomer 31/32 (**Supplementary Fig. 11**), with the structure S4CYH1L.1-15_26-31_31/32_32/34_35-46 marked by 31/32 hybrid. The duplicated sequence is as follows:
   TAAAAACTACACAGAAGCATTCTGAGAAACTTCTCAGTGATGTGAGCATTCTTCTCA
   CAGAGTTGAACTATCTTTTGATTGAGCAGTTTTGAAACACTGTTTTTTTAGAATCTG
   CAAGTGAATATTTGGAGCCTTTTGGGTCTTATTGTGGAAAAGGAAATATCTTCACAT
   AAAAAC
   The right-flank long HOR 1-46 is the same as in HG002.

5. Insertion of 4bp in RP11. 10 HORs in RP11 have 4 bp insertion (INS) in monomer 6 (5 TG dinucleotides instead of 3, revealed by CTTCTTTGTGATGTGTGTGTGTATTC 26mer). No HORs with INS are observed in HG002. The control probe for this region is a 22mer with just 3 TG dinucleotides (CTTCTTTGTGATGTGTGTATTC; probe C1) which appeared to be not specific to cenY and gave multiple hits in autosomes which could be clearly seen upon screening of female genomes (80-100 hits per SRA project or 5-10 hits per diploid asm in 25 women screened; **Supplementary Table 19**). Thus a longer 31mer probe was devised by adding 5 bp on each end (GTATGAATACACACATCACAAAGAAGTTTCT; probe C2). Notably, two INS occur in 35mers and one in 44mer long HOR on the left flank. Thus, the INS does apparently appear in three different structural backgrounds. The likely explanation is that 35mers and 44mer with INS in RP11 are in fact hybrid HORs which fused a part of HOR with mon6 (which contains the INS) with the downstream parts which are the signatures of 35mer and 44mer. An alternative explanation is the repeated independent formation of the INS.
6. HOR-haps in RP11. The cenY RP11 sequence was annotated using the HMMER-based HOR-hap tool built using the HG002 HOR alignment as described above (see **Fig. 3** in the main text). It appears that the blue and green HORs are almost completely absent, and the centromere is formed by the red HOR array.
7. HG002 versus RP11 comparison summary. The above analysis shows that despite the similar number of HORs and similar non-HOR flanks and flanking HORs the bodies of the active arrays differ dramatically in HG002 versus RP11. These differences are summarized in **Supplementary Table 20**. Specifically, the HOR-hap composition is different, as only the red haplotype is shared. Moreover, out of 43 red HORs in RP11, 10 have the INS and 9 have 35mers. As both features are absent in HG002, it is clear that the red region occupied by these HORs is not shared between the two centromeres. That leaves only two small featureless regions in RP11 (see **Fig. 3** in the main text) which could be shared. As many cenYs are known to be much longer than the ones in HG002 and RP11 (Miga et al. 2014), one may hypothesize that the two centromeres are the deleted derivatives of a much longer one ancestral to both, with only a little or no overlap between the two derived deleted variants. Further studies of complete cenY assemblies are needed to prove or disprove this proposition.

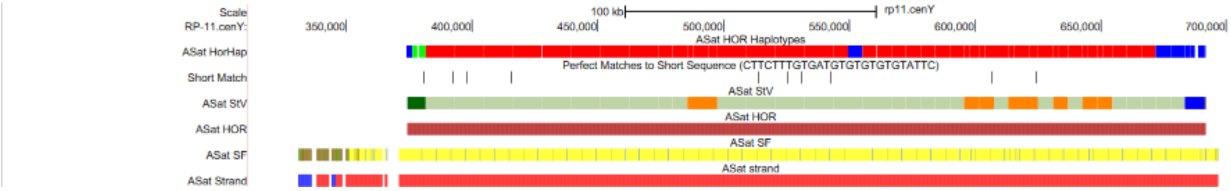


**Supplementary Fig. 12 | RP11 centromere, active HOR array structure, HOR haplotypes and SVs.** Tracks are shown the same way as in **Supplementary Fig. 9**. The short match track (perfect match of INS k-mer) indicates the positions of the HORs with 4 bp insertion. In StV track, the orange boxes indicate 35mer HORs, the dark blue box on the right side is 46mer, and the black box on the right side is an intact 44mer 1-44 (different from the rearranged one in HG002). Graphics from this figure were used to create RP11 panels in **Fig. 3** in the main text.

**a**

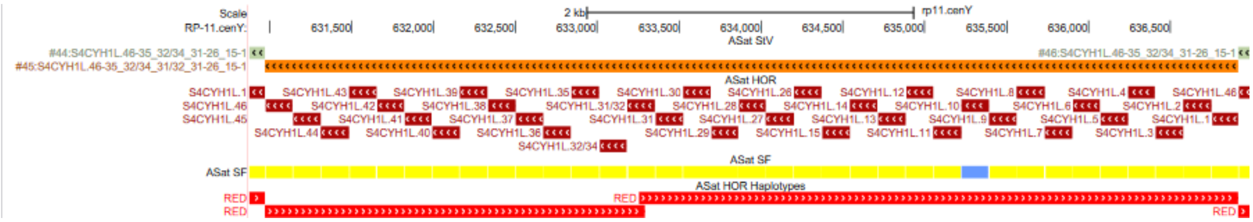

**b**

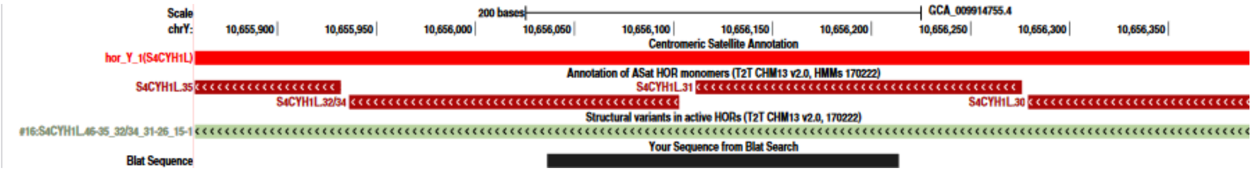

**Supplementary Fig. 13 | The structure of the 35mer HOR. a.** The Browser for RP11. The StV, HOR, SF and HOR-hap tracks are shown, as described above. The monomer structure of the HOR should be read from right to left, as the HOR is on the reverse strand. The 31/32 hybrid monomer (followed by 32/34 hybrid characteristic of the 34mer) is the signature of a 35mer. As the resulting sequence goes 31_32/31_32/34, it indicates the out-of-register duplication of a monomer which contains parts of monomers 31 and 32. Also, note that the yellow color in the SF-track indicates SF4, note that monomer 10 in the S4CYH1L is almost always recognized as SF5 (R2 class), which is very close to SF4. The HOR-hap track at the bottom shows that the 35mer is recognized as a red HOR-hap. **b.** The Browser for HG002. The black bar at the bottom indicates the region duplicated in a 35mer.

The HMM files for the HMMER-based HumAS-HMMER-HOR classification tool and for S4CYH1L HOR-hap classification tool are provided in **Supplementary Files 2-3** (Y_HOR HMM and Y_HOR-hap HMM), respectively. The latter contains a single HMM with profiles for classification of all human AS HORs, it is identical to the one published in Altemose *et al.*(Altemose et al. 2022) except the profiles for S4CYH1L which were updated from 34mer to 46mer HOR.

#### CENP-A

The CENP-A CUT&RUN data was aligned to the T2T-CHM13v2.0 assembly as previously described in Altemose *et al.* (Altemose et al. 2022). The alignments were filtered using the single-copy k-mer locus filtering method as described in Hoyt *et al.* (Hoyt et al. 2022) through the use of the UCSC GenomeBrowser tool overlapSelect. This filtering method is dependent upon location and requires that a given read alignment overlap an entire single copy k-mer, in this case 51-mer (with the parameter “-overlapBases=51”), in the T2T-CHM13v2.0 assembly in order to be retained.

#### Stained glass plot of the DYZ3 array

To generate the StainedGlass plot of the DYZ3 array, we first extracted the sequences corresponding to the DYZ3 array and surrounding regions (chrY:10350001-10950000). Then, we ran StainedGlass v0.4 with the following command: ./StainedGlass.sh --config sample={sample_name} fasta={fasta_path} mm_f=30000 window=5785 --cores {num_of_cores} make_figures. To adjust the color scale in the plot, we used a custom R script to redefine the breaks in the histogram and its corresponding colors. That script is publicly available here: https://eichlerlab.gs.washington.edu/help/glogsdon/Shared_with_Arang/HG002_chrY_StainedGlass_adjustedScale.R. The command used to generate the new plot is: HG002_chrY_StainedGlass_adjustedScale.R -b {output_bed} -p {plot_prefix}.

### Epigenetic profile

#### ONT NanoNOMe sequencing data

HG002 NanoNOMe data generated in Gershman *et al*. (Gershman et al. 2022) was used in this study. Raw sequencing data can be accessed on the Sequence Read Archive BioProject with accession number PRJNA725525 (**Supplementary Table 1**).

#### ONT NanoNOMe alignments

Sequencing reads were indexed with f5c v0.5 (Gamaarachchi et al. 2020) with the "index" command with "--iop 3 -t 48" settings. For alignment, high frequency k-mers were computed using meryl v1.3 (Rhie et al. 2020) for both T2T-CHM13v2.0 and T2T-CHM13v1.1 autosomes + HG002XYv2.7 references with the following commands:

```
meryl count threads=48 k=15 output merylDB_20 chm13v2.0.fa.gz
meryl print greater-than distinct=0.9998 merylDB_20 >
chm13v2.0_repetitive_k15.txt

meryl count threads=48 k=15 output merylDB_27 chm13v1.1_hg002XYv2.7.fasta
meryl print greater-than distinct=0.9998 merylDB_27 >
chm13v1.1_HG002XYv2.7_repetitive_k15.txt
```

HG002 nanoNOME reads were aligned using Winnowmap v2.03 (C. Jain et al. 2022, 2) with the k-mers computed above and alignments were converted from SAM to BAM and sorted using samtools v1.9 (Li et al. 2009). The alignments were indexed using samtools. The following commands were used for both sets of alignments:

```
winnowmap -t 48 -W {genome}_repetitive_k15.txt -ax map-ont {genome_fasta}
{nanonome_fastq} |
samtools view -@48 -Sb |
samtools sort -@48 -o {bam}
samtools index -@48 {bam}
```

Alignments were then merged using the samtools "merge" command with default settings and the combined BAM file was indexed using samtools as above. The combined BAMs were then filtered to retain primary alignments and a list of primary reads greater than 20 kb (as done in (Gershman et al. 2022)) was created for future filtering with the following commands:

```
samtools view -@48 -h -b -F 256 -F 2048 {combined_bam} >
filtered_{combined_bam}
samtools view -@48 filtered_{combined_bam} |
awk 'length($10) > 20000' |
cut -f1 > {genome}_20kb_readIDs.txt
```

#### ONT Nanopolish CpG and GpC methylation calling

Nanopolish v0.13.2 from the "nanonome" branch (https://github.com/jts/nanopolish/tree/nanonome) was used to call methylation on HG002 nanonome data (Simpson et al. 2017). CpG and GpC methylation was called simultaneously with the four-state "cpggpc" model (Lee et al. 2020) for each FASTQ using the following command:

```
nanopolish call-methylation \
--progress \
-b filtered_{combined_bam} \
-r {fastq_file} -g {genome_fasta} \
-q cpggpc \
-t 48 > {methylation_calls}.tsv
```

Methylation calls for each FASTQ were combined and reads greater than 20 kb with primary alignments were retained using the read list generated above. CpG and GpC calls were processed separately with a log-likelihood ratio (LLR) cutoff of –1.5/1.5 for CpG methylation and a LLR cutoff of -1/1 for GpC methylation (Gershman et al. 2022). Using custom scripts (https://github.com/timplab/nanopore-methylation-utilities) methylation calls were made, TSVs were converted to BedGraphs, BedGraphs were sorted, compressed, and indexed, then aggregated methylation frequencies were produced. The following commands were used for each genome:

```
python3 ./nanopore-methylation-utilities/mtsv2bedGraph.py \
-q cpg \
-c 1.5 \
--nome \
-i {cpg_methylation_calls}.tsv \
-g {genome_fasta} > cpg_meth.tmp
      sort cpg_meth.tmp -k1,1 -k2,2n | bgzip > cpg_meth.bed.gz
tabix -p bed cpg_meth.bed.gz
      python3 ./nanopore-methylation-utilities/parseMethylbed.py \
frequency -i cpg_meth.bed.gz > cpg_meth.freq

python3 ./nanopore-methylation-utilities/mtsv2bedGraph.py \
-q gpc \
-c 1.0 \
--nome \
-i {cpg_methylation_calls}.tsv \
-g {genome_fasta} > gpc_meth.tmp
      sort gpc_meth.tmp -k1,1 -k2,2n | bgzip > gpc_meth.bed.gz
tabix -p bed gpc_meth.bed.gz
      python3 ./nanopore-methylation-utilities/parseMethylbed.py \
```

```
frequency -i gpc_meth.bed.gz > gpc_meth.freq
```

#### ONT Remora CpG methylation calling and processing

HG002 ONT nanopore sequencing data was re-basecalled using Guppy v6.1.2 using dna_r9.4.1_450bps_modbases_5mc_cg_sup_prom.cfg model with the following command:

```
guppy_basecaller -i input -s save -c
dna_r9.4.1_450bps_modbases_5mc_cg_sup_prom.cfg -x "cuda:all" -r
```

Resulted unaligned BAM files containing methylation calls (modbams) were downloaded from the Human Pangenome Reference Consortium (https://s3-us-west-2.amazonaws.com/human-pangenomics/index.html?prefix=NHGRI_UCSC_panel/HG002/nanopore/ultra-long/). Modbams were converted to FASTQ files while retaining the modbam MM and ML tags using samtools v1.15.1 using the following command:

```
samtools fastq -@48 -T Mm,Ml {input.bam} > {output.fastq}
```

FASTQ files were aligned to both T2T-CHM13v2.0 and HG002XYv2.7 genomes using Winnomap v2.03 as above for nanoNOMe data with the addition of the “-y” parameter which retains the MM and ML tags. The alignments were then converted to sorted BAMs containing only primary mappings and indexed. The following commands were used:

```
winnowmap -t 48 \
-W {genome}_repetitive_k15.txt \
-ax map-ont -y {genome_fasta} {mod_fastq_list} > {output.sam}
      samtools view -@24 -Sb -F 256 -F 2048 {output.sam} |
samtools sort -@24 -T {temporary_directory} - > {output.bam}
samtools index -@48 {output.bam}
```

Aggregated methylation percentages at all CpGs were then calculated using modbam2bed v0.6.2 (https://github.com/epi2me-labs/modbam2bed) with bases with >0.8 probability called “methylated” and bases with <0.2 probability called “unmethylated.” Bed files were then converted to a format similar to the “cytosine report” produced by the Bismark software v0.23.1dev (Krueger and Andrews 2011). The following commands were used:

```
modbam2bed -t 48 \
-e \
-m 5mC \
--cpg \
-a 0.20 -b 0.80 \
{genome_fasta} {output.bam} > {output.bed}
awk -v OFS='\t' '{{print $1,$3,$6,$13,$12,"C","CG"}}' {output.bed} >
{output.bismark}
```

#### PacBio methylation calling

The probability of methylation for each CpG site in HiFi reads was assigned using primrose v1.3.0 (https://github.com/PacificBiosciences/primrose) with default parameters. Reads were aligned with pbmm2 v1.9.0 (https://github.com/PacificBiosciences/pbmm2). The percent of methylated reads at each reference genome position was calculated using pb-CpG-tools v1.1.0 (https://github.com/PacificBiosciences/pb-CpG-tools) with `-p model`. Resulting modbams were re-processed identically to Remora-called ONT data to collect comparable ("native", to distinguish from pb-CpG-tools results) Bismark-like aggregated methylation data, with probability thresholds of <20% for unmethylated and >80% for methylated.

#### Whole genome bisulfite sequencing (WGBS) and Enzymatic Methyl-seq (EM-seq) processing

WGBS and EM-seq data from HG002 (Foox et al. 2021) were downloaded from SRA with ffq v0.2.1 (Gálvez-Merchán et al. 2022). WGBS and EM-seq reads were trimmed using fastp v0.23.2 (Chen et al. 2018) according to (Foox et al. 2021) for these technologies. Reads were aligned to the CHM13+HG002XYv2.7 genome that was combined with common bisulfite sequencing control genomes, unmethylated lambda phage and methylated pUC19 plasmid. The combined genome was prepared for Bismark v0.23.1dev (Krueger and Andrews 2011) analysis through the use of the "bismark_genome_preparation" command with "--bowtie2 --genome_composition" as key parameters. Trimmed reads were aligned using the "bismark" command with "--bam --bowtie2" as key parameters. Alignments were deduplicated using "deduplicate_bismark" and methylation bias was determined using "bismark_methylation_extractor." Methylation data was extracted using the script "bismark_methylation_extractor" and converted to aggregated methylation data files using the scripts "bismark2bedGraph" and "coverage2cytosine." This analysis was packaged into a Snakemake pipeline (see Code availability for more details).

#### Analysis of methylation detection technologies

Methylation data from long-read and short-read technologies were compared in R (R Core Team 2018). Custom R scripts (see Code availability below) used the R packages bsseq (Hansen, Langmead, and Irizarry 2012), Biostrings (Pagès et al. 2022), the GenomicRanges family of packages (Lawrence et al. 2013), GenomeInfoDb (Arora et al. 2022), gUtils ("Mskilab/GUtils: R Package Providing Additional Capabilities and Speed for GenomicRanges Operations. Last Accessed: 2022-11-28." n.d.), corrplot (https://github.com/taiyun/corrplot), ggplot2 (Wickham, Danielle Navarro, and Thomas Lin Pedersen n.d., 2), readr (Wickham et al. 2022), rtracklayer (Lawrence, Gentleman, and Carey 2009), and tidyverse (Wickham et al. 2019). For comparison between technologies, CpG sites were retained for analysis if the site was a reference CpG site and if the site was covered by ≥5 reads and ≤200 reads in all samples analyzed. Additionally "GCG" sites were excluded from correlation analyses due to the inability of these sites to be accurately called in ONT nanoNOMe data (Lee et al. 2020). Both bigWig and BED files containing aggregated methylation percentages for CpGs and GpCs were created in R with custom scripts (https://github.com/arangrhie/T2T-HG002Y/tree/main/epigenetics). CpG and GpC sites with no called reads were omitted from these files.

Methylation summary files are available to download or load on IGV at https://s3-us-west-2.amazonaws.com/human-pangenomics/index.html?prefix=T2T/CHM13/assemblies/annotation/regulation/. Code for methylation analysis can be found on GitHub (https://github.com/arangrhie/T2T-HG002Y/tree/main/epigenetics).

#### Centromeric Dip Region (CDR)

The Centromeric Dip Region (CDR) was annotated as previously described (Gershman et al. 2022). The CDR was manually annotated as the area where CpG methylation is lower than the flanking active, alpha-satellite, higher order array (HOR) in the centromere. The HG002 cenY CDR shows two “dips” in the average methylation percentage (**Fig. 3**), which is supported by read-level methylation calls (**Supplementary Fig. 14**).

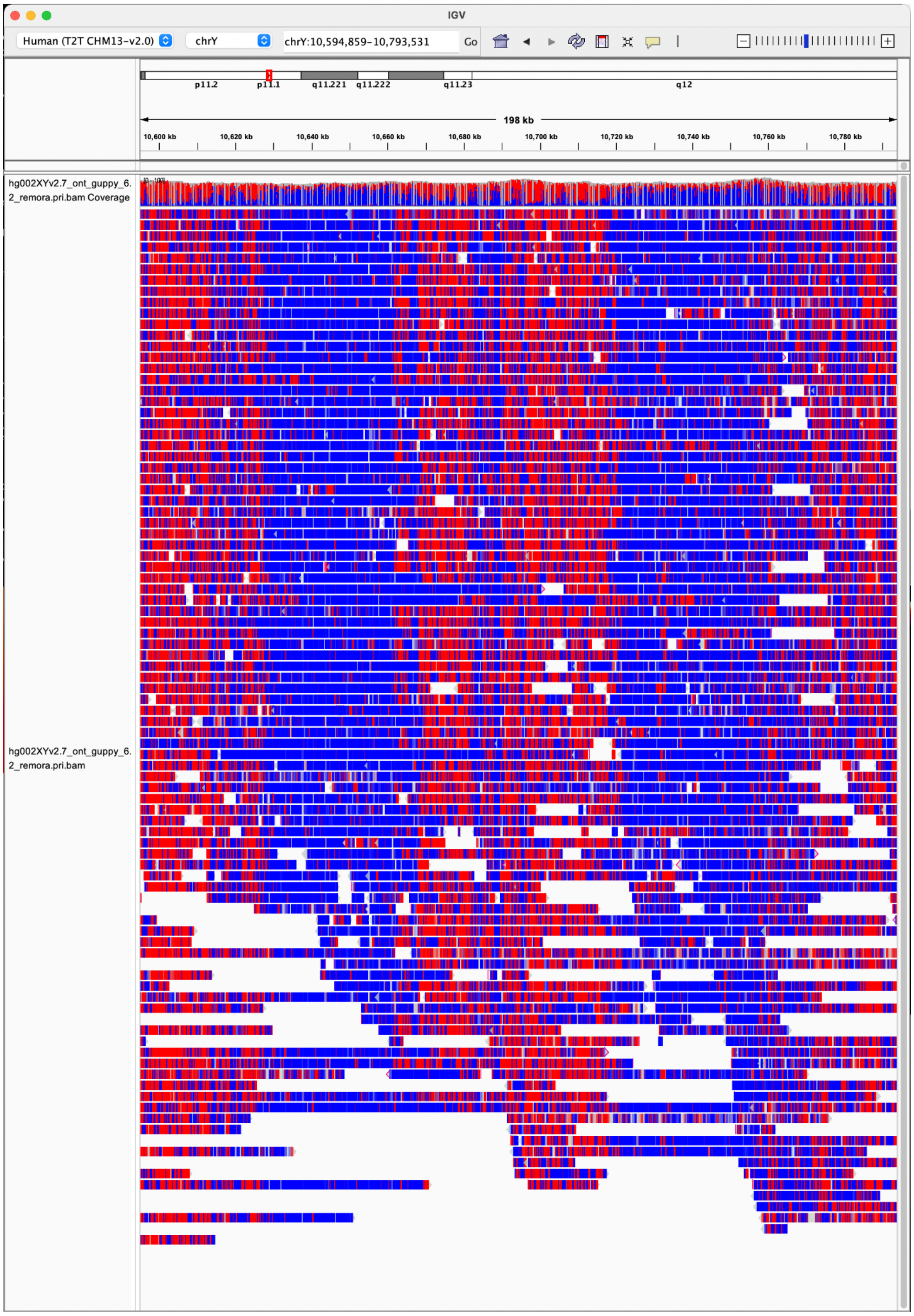


**Supplementary Fig. 14 | Centromeric methylation dips observed in ONT ultra-long reads.** Ultra-long nanopore reads mapped to HG002 cenY and visualized in IGV where unmethylated CpGs are blue and methylated CpGs are red. Two distinct hypomethylated dips are visible at both consensus and single-molecule resolution.

### Sequence classes on the Y chromosome

In order to differentiate between X-degenerate and ampliconic regions, we used either exact boundaries of palindromes or intrachromosomal identity as defined in Skaletsky *et al.* (Skaletsky et al. 2003) with adjusted borders based on the gene annotations. First, the repeat-masked ChrY was split into 5 kb sliding windows with a step size of 1 kb. Each window was mapped back to the T2T-Y chromosome v2.7 with the Winnowmap2 v2.03 (C. Jain et al. 2022). After excluding self-alignments (windows where the distance between the window coordinates and the mapped region was smaller than 5 kb), all alignments were required to span at least 1kb, and the identity between the mapped sequence and the reference was calculated. For each window, the maximum identity to any other window was calculated, and the windows with identity over 50% were considered indicative of ampliconic regions if present consecutively.

#### Palindrome structure, P1-P3

The palindrome P1 underwent a rearrangement consistent with a single event of NAHR. For the schematic representations in **Fig. 4**, amplicons from Teitz *et al.* (Teitz et al. 2018) were mapped to GRCh38-Y and T2T-Y assemblies with Winnowmap2 v2.03 (C. Jain et al. 2022) and the following commands:

```
meryl count k=15 output merylDB chrY_hg002_v2.7.fasta
meryl print greater-than distinct=0.9998 merylDB > repetitive_k15_HG002.txt

winnowmap --MD --eqx -W repetitive_k15_HG002.txt -ax map-pb
chrY_hg002_v2.7.fasta Y_repeats.fasta >Y_repeats_on_HG002.sam
winnowmap --MD --eqx -W repetitive_k15_HG002.txt -ax map-pb chrY.fa
Y_repeats.fasta >Y_repeats_on_hg38.sam
```

#### Palindrome structure, P4-P8

The approximate boundaries of palindrome arms were manually selected using Gepard v2.1 (Krumsiek, Arnold, and Rattei 2007), and further refined based on a self-alignment of palindrome arms with adjacent flanks and the reverse complement of the same sequence using global alignment with Stretcher (Rice, Longden, and Bleasby 2000) (**Supplementary Table 20**). This enabled the extraction of arm sequences that were used to calculate the sequence identity between palindromic arms.

#### Azoospermia Factor (AZF) region

For AZFa, the deletion is known to be caused by recombination between two HERV15 proviruses, and potentially deletes two genes: USP9Y and DDX3Y (Sun et al. 2000). Based on the CAT/Liftoff gene annotation for the T2T-Y, we found the HERV15 surrounding the two genes above. Sequences between the two HERV15s were used to determine the AZFa boundaries. The length of the AZFa region was 791 kb, matching previous estimates of 800 kb. Boundaries of AZFb and AZFc were defined by the amplicon units; by P5/proximal P1 deletion (yel3/yel1) and by the b2/b4 deletion. In brief, we generated a self-dotplot of the T2T-Y assembly using word size of 100 with Gepard v2.1 (Krumsiek, Arnold, and Rattei 2007). Then, breakpoints were identified as illustrated in Fig. 2 of Navarro-Costa *et al.* (Navarro-Costa, Plancha, and Gonçalves 2010) as shown in **Supplementary Fig. 15**.

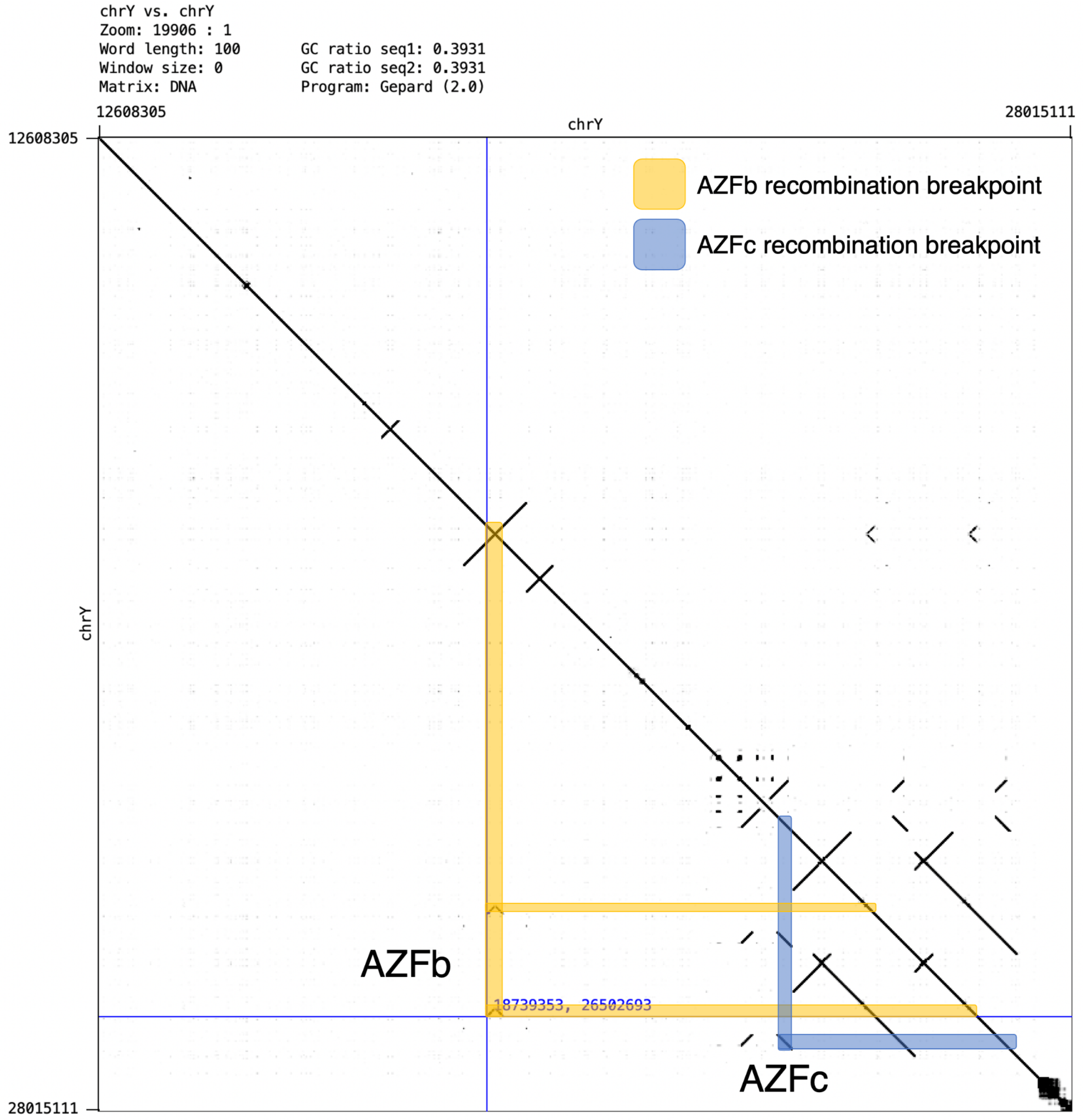


**Supplementary Fig. 15 | Identifying AZFb and AZFc recombination breakpoints.**

#### Pseudoautosomal region (PAR) and X-transposed region (XTR)

Initially, we ran LASTZ v1.04.00 (Harris, Robert S. 2007) to identify the PAR and XTR coordinates on the ChrX and ChrY sequences. LASTZ was run between the unmasked T2T-CHM13v1.1 X chromosome and T2T-Y chromosome, and between the HG002 X chromosome and HG002 Y chromosome. We ran LASTZ in two stages; in the first stage we identified ungapped alignments that were then used in the second stage as anchors for gapped alignments. In the first stage (ungapped alignment step), we filtered out short alignment by keeping anchors with 400 or more matched bases and more than 85% sequence identity. In the second stage (gapped alignment step), we retained alignments with at least 1000 matched bases, testing 96%, 97%, 98%, 99%, and 99.5% identify, showing only 96% and 97% because the PAR boundaries were the same across all boundaries and the XTR differed only between 96% and 97%. We generated and visually assessed the dotplots from these alignments to identify the start and end of the one-to-one alignments (**Extended Data Fig. 9a**). Dotplots were generated in R (R Core Team 2018). The full LASTZ commands with parameters:

```
lastz T2T_chrY.fa[unmask] T2T_chrX.fa[unmask] --ungapped \
--filter=identity:80 --filter=nmatch:400 --hspthresh=36400 \
--format=general-:name1,start1,end1,name2,start2,end2,strand2,nmatch \
--progress=1 > t2t_X_onto_Y.unmasked.anchors
```

```
lastz T2T_chrY.fa[unmask] T2T_chrX.fa[unmask] \
--segments=t2t_X_onto_Y.unmasked.anchors --allocate:traceback=800M \
--filter=identity:96 --filter=nmatch:1000 \
--
format=general:name1,zstart1,end1,name2,strand2,zstart2+,end2+,nmatch,
length1,id%,blastid% \
--rdotplot=t2t_X_onto_Y_identity100.unmasked.dots --progress=1 >
t2t_X_onto_Y_identity96.unmasked.lz
```

For PARs, the coordinates were the same across all identity filters while XTR was slightly shorter when using the 97% identity filter compared to the 96% identity filter (**Supplementary Table 26**). For the X chromosome XTR, there were only 8 base pairs separating the shorter and longer XTR segment so we merged them as one region. The ChrX PAR and XTR coordinates differed between the CHM13-X and HG002-Y alignment and HG002-X and HG002-Y alignment, due to different lengths between CHM13-X and HG002-X; CHM13-X is 154,259,566 bp while HG002-X chromosome is 154,349,815 bp.

Alternatively, we produced an additional set of alignments using HG002-X and HG002-Y and CHM13-X. Alignment was performed with Minimap2 using the following code:

```
minimap2 -cxasm20 --cs -z10000,1000 chrX.fa chrY.fa
```

The resulting pairwise alignment was then filtered by identity > 95% and length > 10 kb. Here are 12 relevant alignment blocks (format: chrY, chrY-len, start, end, strand, chrX, chrX-len, start, end, matching bases, length of alignment block):

```
chrY 62460029    3832  108018     +     chrX  154259566  1     108464
     99556 112676     <- PAR1: block1
chrY 62460029 108607   1328683  + chrX 154259566 109273     1307231
1167409 1246039  <- PAR1: block2
```

```
chrY  62460029  1370293   2458320   + chrX  154259566  1307341    2394410
1080607 1092832  <- PAR1: block3  (end)
chrY  62460029  2727072   2883124   + chrX  154259566  87835350   87991293
153124  157354
chrY  62460029  2888377   3067662   + chrX  154259566  87996364   88173617
171009  183667
chrY  62460029  3067718   3340757   + chrX  154259566  88183921   88447036
259424  273751
chrY  62460029  3343080   4032012   + chrX  154259566  88536875   89223206
672016  694556
chrY  62460029  4165131   4182820   - chrX  154259566  89362279   89379978
17469   17735
chrY  62460029  4400137   5077740   + chrX  154259566  89931381   90624986
655850  707405   <- EDIT: minimap2 is missing alignment
chrY  62460029  5077744   5914561   + chrX  154259566  90736063   91570785
811002  850523
chrY  62460029  6201358   6400875   + chrX  154259566  87642550   87835359
186528  203117   <- this is translocated
chrY  62460029  62122809  62456509 + chrX  154259566  153925834  154259566
333087  333917   <- PAR2
```

The PAR and XTR coordinates from both tools were manually revised based on exact sequence alignments to determine the boundaries (**Supplementary Table 21**).

### Yqh heterochromatin region

#### Yqh DYZ1/DYZ2

To produce **Extended Data Fig. 8a**, the ChrY sequence (except for Yq) was split into non-overlapping 50 kb bins, and each bin was input into NTRprism v0.22 (Altemose et al. 2022); parameters: 1 6000 10 6; https://github.com/altemose/NTRprism. In the Yq region, each entire block of DYZ18, DYZ1, or DYZ2 was input into NTRprism. The top periodicity detected in each bin/region was reported if its normalized column sum from the NTRprism output was greater than 0.01. The dotplots in the **Extended Data Fig. 8b,d** were produced using dottup, part of the EMBOSS software package v6.6.0.0 (Rice, Longden, and Bleasby 2000).

To generate DYZ1 and DYZ2 consensus sequences, their repeat units were first extracted by an in-silico digestion of the Yq region at the EcoRI recognition site for HSat3/DYZ1 and the SpeI recognition site for HSat1B/DYZ2. Resulting fragments were size filtered (keeping HSat3 fragments between 3400-3700 bp; HSat1B fragments between 2300-2500 bp), then a multiple sequence alignment was produced for each family using kalign v3.3.2 (Lassmann 2020). HMMER was used to convert this multiple alignment into a profile HMM v3.3.2 with hmmbuild command (Wheeler and Eddy 2013), then nhmmer was used to scan across the Yq reference sequence and identify additional sequence matches to this profile HMM, which may have been missed by the *in silico* digestion approach. These matches were aligned and a second profile HMM was built, which was fed into nhmmer one more time, followed by alignment and building a third profile HMM, from which a consensus sequence was emitted. This consensus was added to the multiple alignment and fed into HMMER's esl-alipid command, from which each individual repeat unit's percent identity to the consensus sequence was extracted and used to generate the plot in **Fig. 5c**.

#### Phylogenetic analyses of *Alu*Y repeats

To understand the relationship between the *Alu*Y instances across T2T-Y, particularly those associated with HSat, a phylogenetic analysis was performed. *Alu*Y loci from RepeatMasker annotations were subsampled (n=500) for each of the following groups across T2T-Y: HSat1B-associated, HSat3-associated, and non-HSat-associated. To extend this analysis, those *Alu*Y loci that were associated with HSat1B and HSat3 across the acrocentric chromosomes of CHM13 were also included (2,500 total *Alu*Y across the five groups). Lastly, to extend this analysis further across the genome, *Alu*Y loci that were associated with HSat3 across non-chrY/non-acrocentric chromosomes were included as well (n=140 total), but not for HSat1B since they are not found associated beyond chrY and the acrocentrics. HSat annotations are part of the Cen/Sat track from the "Satellite annotation" section. Since *Alu*Y was derived from *Alu*Sc8, this consensus was included as the outgroup (http://www.repeatmasker.org/AluSubfamilies/humanAluSubfamilies.html). Alignment was carried out in MAFFT v7.471 (Katoh and Standley 2014) using custom parameters based on manually inspected alignment results. Phylogenetic analysis was run in RAxML-NG v0.9.0 (Stamatakis 2014). Rapid bootstrap approximation was run to generate 100 bootstrap replicates using the University of Connecticut Health Center server. Consensus trees were then generated in Geneious v2019.2.3 (Geneious n.d.) and visualized in FigTree v1.4.4 ("FigTree. Last Accessed: 2022-11-28." n.d.).

### Short-read variant calling on T2T-CHM13+Y

Short-read alignment and variant calling pipeline used for the 1KGP and SGDP samples is released in https://github.com/schatzlab/t2t-chm13-chry as v1.0.0.

#### Impact of masking PAR and XTR in variant calling

To assess variant calling and filtering strategies on the X and Y chromosomes, we simulated paired-end sequence reads with a read length of 150 base pairs (bp) for 10 XY and 10 XX individuals, using NExt-generation sequencing Analysis Toolkit (NEAT) software (https://github.com/zstephens/neat-genreads, commit a2d7739) (Stephens et al. 2016). Variants were inserted from 10 XY and 10 XX European individuals from high coverage variant calls from 1KGP (Byrska-Bishop et al. 2022) using the -v option in NEAT (**Supplementary Table 25**). NEAT requires a reference genome to sample reads from, so we used the same reference genome that was used in the generation of the 1KGP high-coverage GRCh38 VCFs (ftp://ftp.1000genomes.ebi.ac.uk/vol1/ftp/technical/reference/GRCh38_reference_genome/GRCh38_full_analysis_set_plus_decoy_hla.fa). Additionally, we used the default sequencing error model provided in the NEAT software. For the autosomes, mitochondrial DNA (mtDNA), and X non-PARs (females only) coverage was set to 20x. For the X and Y non-PARs in XY samples, coverage was set to 10x each as both are haploid in a 46,XY genome and expected to have half the sequencing coverage of a diploid autosome. PAR sequences were simulated as diploid on the X chromosome for both XY and XX samples because the PAR variants in 1KGP were called only on the X chromosome. NEAT produces simulated FASTQs, along with a golden VCF with the set of true positive variants and a golden BAM with a golden set of aligned reads.

We performed quality trimming and alignment on the simulated FASTQs. Using BBMap bbduk v35.85 (Bushnell 2014), a high-performance tool for adapter and quality trimming, bases with a Phred score of 20 were trimmed on both the left and right side of reads, reads shorter than 75bp were discarded, and reads with average Phred quality below 20 were removed (bbduk trimming parameters used: qtrim=rl trimq=20 minlen=75 maq=20).

Reads were mapped to a version of 1KGP GRCh38 reference genome where the both X and Y chromosomes were unmasked (default), and versions of 1KGP GRCh38 reference genome informed by the sex chromosome complement (SCC) of the sample using bwa v0.7.17 (Li and Durbin 2009) (**Supplementary Table 27**). Briefly, XX samples are expected to have two X chromosomes and no Y chromosome, while XY samples are expected to have an X and a Y chromosome; therefore, the sex chromosome complement reference genome for XX samples has the Y chromosome hard masked out, and the sex chromosome complement reference genome for XY samples has the pseudoautosomal region (PARs) on the Y chromosome hard masked (Webster et al. 2019). For the simulated XY samples, we additionally aligned reads to the SCC version of the reference genome with the Y chromosome XTR sequence hard masked out: XTR1: 3050044–6235111, XTR2: 6532906–6748713. The Y chromosome XTR was hard masked using BEDtools v2.30.0 (Quinlan and Hall 2010) maskfasta. X and Y PAR coordinates were obtained from Ensembl GRCh38 PAR definitions (Aken et al. 2017): X PAR1: 10001–2781479, Y PAR1:10001–2781479, X PAR2: 155701383–156030895, Y PAR2: 56887903–57217415. XTR and ampliconic coordinates on the Y chromosome were obtained from Poznik *et al.* (Poznik et al. 2013) in hg19 and lifted over to GRCh38 using UCSC liftOver (Kent et al. 2002), X chromosome XTR coordinates were obtained from Webster *et al.* (Webster et al. 2019), and X chromosome ampliconic coordinates were obtained from Cotter *et al.* (Cotter, Brotman, and Wilson Sayres 2016) in hg19 and lifted over to GRCh38 using UCSC liftOver tool (Kent et al. 2002).

To assess the mapping quality between default and SCC alignments, we calculated average MAPQ across chromosome X in 50 kb windows, sliding 10 kb using BEDtools map for one simulated XY and one simulated XX sample. Before running the bedtools map, we converted the bam files to bed using BEDtools bamtobed.

Variant calling was performed using GATK's v4.2.1.0 HaplotypeCaller tool, joint genotyping was performed using GenotypeGVCFs tool, and filtering was performed using SelectVariants and VariantFiltration tools (McKenna et al. 2010). For simulated XX samples and XY samples, PARs were called diploid. XY and XX samples were jointly genotyped separately and filtered using GATKs hard filtering threshold recommendations (filtering parameters: -filter "QD < 2.0" -filter

"QUAL < 30.0" -filter "SOR > 3.0" -filter "FS > 60.0" -filter "MQ < 40.0" -filter "MQRankSum < -12.5" -filter "ReadPosRankSum < -8.0").

We calculated true positives (TP), false positives (FP), and false negatives (FN) for each individual within PAR1 and PAR2 using a custom python script. Using both the simulated VCFs output from NEAT that have the ground truth variants for each individual and the called VCFs, a TP was defined as a SNP that was both called and simulated, a FP was defined as a SNP called but not simulated, and a FN was defined as a SNP that was simulated but not called. For XY samples, where we additionally tested the impact of masking the XTR on the Y. To this we calculated the number of TPs, FPs and FNs on the X chromosome XTR using just the X chromosome golden VCFs for each individual.

#### Mappability comparison and variant calling in 1KGP samples

Using the NHGRI Genomic Data Science Analysis, Visualization, and Informatics Lab-Space (AnVIL) (Schatz et al. 2022), we performed short-read alignment and variant calling for the 3,202 samples in 1KGP (Byrska-Bishop et al. 2022) using the T2T-CHM13v2.0 assembly as a reference. These samples were sequenced to at least 30x coverage by the New York Genome Center (NYGC), and alignment and variant calling was previously performed on the GRCh38 reference.

For our analysis, we largely followed the short-read alignment and variant calling pipeline previously used for analysis of T2T-CHM13v1.0 (Aganezov et al. 2022). This pipeline was initially built to mirror the pipeline used by NYGC on the GRCh38 reference, using updated tools when appropriate. By using a nearly identical pipeline to the one used by NYGC, it allows us to assess differences in alignment and variant calling between GRCh38 and T2T-CHM13, without confounding from the pipeline itself. For analysis of short read alignment and variant calling on T2T-CHM13v2.0, we used the same pipeline as the T2T-CHM13v1.0 analysis, although we updated certain elements of the pipeline to better represent genetic variation in the PARs. Specifically, we used XYalign v1.15 (Webster et al. 2019) to produce two separate karyotype-specific references (the sex chromosome complement described in the previous section). In the XX-specific reference, the entire Y chromosome is masked, whereas in the XY-specific reference, only the Y-PARs are masked. During the alignment step, we aligned XX and XY samples separately, using the appropriate karyotype-specific reference. This effectively forces any reads originating from the PARs to align to the X chromosome PAR. During the haplotype-calling phase of the variant-calling step, samples were again processed independently, using the appropriate karyotype-specific reference. Importantly, for XY samples, variants were called in the X-PAR as diploid, rather than haploid. Thus, the X-PAR effectively represents genetic variation occurring within either the X-PARs or the Y-PARs. While this does make it impossible to determine—for XY samples—whether variation in the PARs is truly originating from the X-PAR or the Y-PAR, it does improve alignment and variant detection in the PARs, as discussed in the main text.

For all analyses, measures of mappability (reads mapped, reads properly paired, mismatch rate) were assessed with samtools v1.16.1 using the “samtools stats” tool, and variant counts and allele frequencies were assessed with bcftools v1.16 using the “bcftools stats” tool (Danecek et al. 2021).

We defined syntenic regions between GRCh38-Y and T2T-Y based on the Minimap2/NEXTflow alignment between GRCh38-Y and HG002-Y described above. Using the GRCh38-to-CHM13v2.0 PAF alignment (https://s3-us-west-2.amazonaws.com/human-pangenomics/T2T/CHM13/assemblies/chain/v1_nflo/grch38-chm13v2.paf), we extracted and merged all alignments between GRCh38-Y and HG002-Y to define 1) regions of GRCh38-Y syntenic with T2T-Y, and 2) regions of T2T-Y syntenic with GRCh38-Y. Using bedtools v2.30.0 (Quinlan and Hall 2010), we subset variant calls in unrelated XY samples to these regions.

Using these variant calls, we then identified variants called on GRCh38-Y that “disappear” on T2T-Y. To do so, we first used GATK release 4.2.4 LiftoverVcf (Picard version 2.25.4) (Van der Auwera GA and O’Connor BD 2020) with the “RECOVER_SWAPPED_REF_ALT” flag to lift over variants from GRCh38-Y to T2T-Y, using the Minimap2/NEXTflow alignment chain file. We then identified the subset of these lifted variants that were not called natively on T2T-Y (based on position and reference allele), and we define this subset as the variants called on GRCh38-Y that “disappear” on T2T-Y. Using their GRCh38-Y coordinates, these variants were then intersected with putative collapsed regions on GRCh38-Y, described below.

#### Putative collapsed regions in GRCh38-Y

Three individual’s variant calls and its corresponding bam files from the 1KGP dataset were downloaded from AnVIL: one individual each from the J1, R1b and E1b haplogroups (HG01130, HG00116 and HG01885, respectively). Variant calls on ChrY syntenic region were subsetted using bcftools v1.13 (Danecek et al. 2021). From the vcf file, allelic read depth (defined as AD field) and reference allele depth (1st value in the DP field) were extracted using a custom script vcfExtractADandDP.jar along with each variant’s chromosomal position.

```
bcftools view -s HG00116,HG01130,HG01885 \
-c1 -R ${REF}_syntenic_to_${TARGET}.noPAR.bed \
-Ov --threads 20 \
-o ${REF}_three_samples.syntenic.vcf ${REF}_unrelated.chrY.pass.variants.vcf
java -jar -Xmx256m vcfExtractADandDP.jar ${REF}_three_samples.syntenic.vcf >
${REF}_three_samples.syntenic.noPAR.txt
```

The output was used for visualization in **Fig. 6e** using a custom R script (https://github.com/arangrhie/T2T-HG002Y). Variants from HG00116 (R1b, similar Y haplogroup with GRCh38-Y, thus least structural variations get called) were further aggregated when non-reference alleles were present, merged within 50 kb.

The bam files were processed to collect coverage tracks using IGVtools v2.14.1 (Robinson et al. 2011) and samtools v1.13 (Danecek et al. 2021):

```
samtools view -hb CHM13v2_crams/$SAMPLE.${VER}.cram -T $FA -@20 -O bam chrY >
${ver}/$SAMPLE.Y.bam
samtools index ${VER}/$SAMPLE.Y.bam
```

```
igvtools count ${VER}/$SAMPLE.Y.bam $SAMPLE.Y.tdf $FA.fai
```

Collected coverage of the three samples, variant calls and the aggregated variants were manually inspected on GRCh38, to agree among all 3 samples to have 1) excessive number of variants called, 2) overlap with known gaps in GRCh38, and 3) does not overlap the palindromic region where we identified substantial rearrangements between the GRCh38-Y and T2T-Y. If the aggregated region overlapped a known sequence class or repeat component, those boundaries were chosen for both the GRCh38-Y and T2T-Y.

#### Mapping and variant calling of the SGDP samples

The Simons Genome Diversity Project (SGDP) presents 279 open-access high-coverage genomes from 130 diverse populations selected to capture greater human genetic, linguistic, and cultural variation (Mallick et al. 2016). Compared to 1KGP, SGDP includes 118 additional populations with samples sequenced to an average of 43x coverage using a shared PCR-free Illumina library. The SGDP samples were aligned and genotyped to T2T-CHM13v2.0 and GRCh38 on AnVIL (Schatz et al. 2022) following the same pipeline as the analysis of 1KGP samples. Read mapping characteristics to T2T-CHM13v2.0 indicated high-quality alignments, similar to the 1KGP results, with a median paired-reads percentage of 97.7% and a median alignment error rate of 0.0044% (**Supplementary Figs. 16-17**). For all samples, mapping to T2T-CHM13v2.0 yielded improvements over GRCh38, with a median error alignment error rate of 0.0053% (**Supplementary Fig. 18)**. The difference (T2T-CHM13v2.0 - GRCh38) in median paired-reads percentage was 0.400%.

Chromosome-level aggregation of PASS variants reveals a varying number of variants called using each reference (**Extended Data Fig. 10d).** For most chromosomes, fewer variants were identified using T2T-CHM13v2.0 than GRCh38, reflecting corrections resulting in fewer artifact variants called from using an inaccurate assembly. For some chromosomes, such as chrY, more variants were identified, capturing variants found in the newly added sequences. Across the entire Y chromosome, roughly 25% more variants were identified using T2T-Y than GRCh38-chrY: 658,636 versus 528,576 variants, respectively. However, within synthetic regions on chrY we find more variants using GRCh38-chrY (375,292 versus 502,088 variants) comparable to what we found for the 1KGP dataset. For both reference genomes, population-level aggregates display an increased count of variants in African samples compared to other non-African populations (**Extended Data Fig. 10e**). Per-sample variant T2T-CHM13v2.0 analysis of all populations reveals a median of 3,463,417 SNPs and 691,222 indels per sample, while subsetting for African samples yields a median of 4,547,156.5 SNPs and 883,054.5 indels. The reduction in variant counts for non-African samples is consistent with 1KGP T2T-CHM13v1.0 variant analysis. As expected, African samples show comparable numbers of variants for T2T-CHM13v2.0 and GRCh38 reflecting greater population genetic diversity relative to these references which have limited African ancestry.

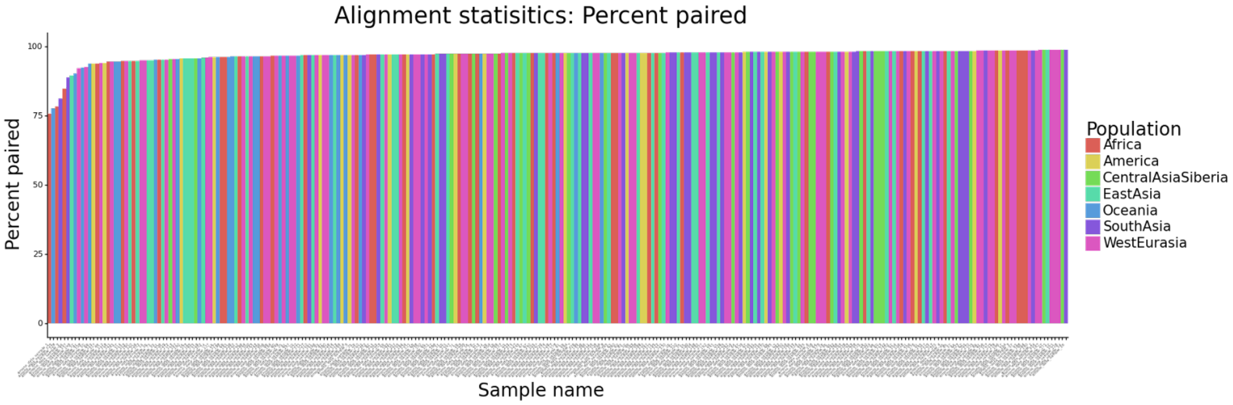


**Supplementary Fig. 16 | Read alignment percentage of paired reads from the SGDP samples.** Each bar represents the percent of proper pairs for individual samples colored by the population of origin.

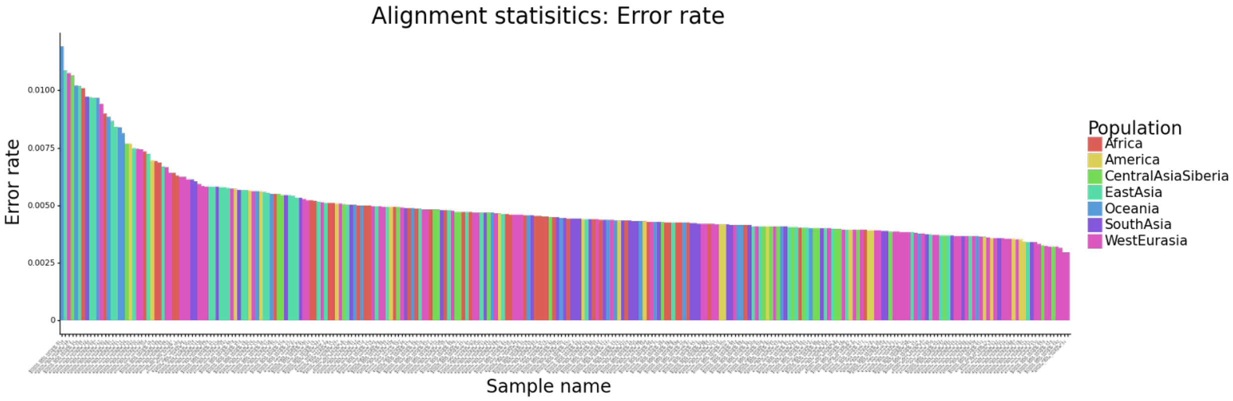


**Supplementary Fig. 17 | Read alignment error rate from the SGDP samples.** Each bar represents the alignment error rate for individual samples colored by the population of origin. Note the alignment error rate represents both sequencing errors in the reads as well as the true biological difference between the sample and the reference genome it is mapped against.

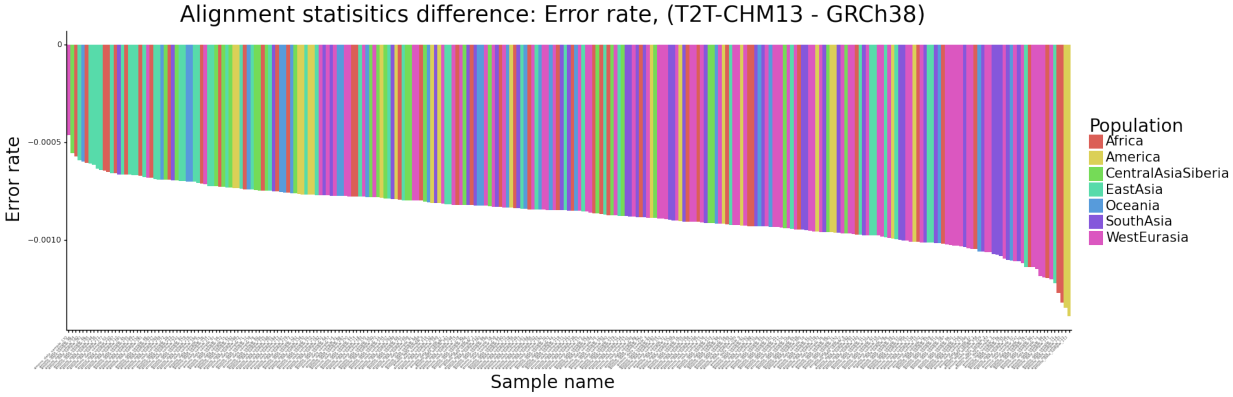


**Supplementary Fig. 18 | Difference (T2T-CHM13v2.0 - GRCh38) in read alignment error rate from the SGDP samples.** Each bar represents the difference in alignment error rate for individual samples colored by the population of origin. All SGDP samples yielded a lower error rate for T2T-CHM13v2.0 as compared to GRCh38.

### Curated syntenic region and liftover chains

#### GRCh38 pre-processing

To prevent ambiguous alignments, all false duplications, as determined by the Genome in a Bottle Consortium (GCA_000001405.15_GRCh38_GRC_exclusions_T2Tv2.bed), as well as the GRCh38 modeled centromeres, were masked from the GRCh38 primary assembly. In addition, unlocalized and unplaced (random) contigs were removed.

#### minimap2-based pipeline

The initial chain file was generated using nf-LO v1.5.1 (Talenti and Prendergast 2021) with minimap2 v2.24 (Li 2018, 2) alignments. These chains were then split at all locations that contained unaligned segments greater than 1 kbp or gaps greater than 10 kbp. Split chain files were then converted to PAF format with extended CIGAR strings using chaintools (https://doi.org/10.5281/zenodo.6342391, v0.1), and alignments between nonhomologous chromosomes were removed. The trim-paf operation of rustybam (https://zenodo.org/record/6342176, v0.1.29) was next used to remove overlapping alignments in the query sequence, and then the target sequence, to create 1:1 alignments. PAF alignments were converted back to the chain format with paf2chain (https://github.com/AndreaGuarracino/paf2chain, commit f68eeca), and finally, chaintools was used to generate the inverted chain file.

Full commands with parameters used were:

```
nextflow run main.nf --source GRCh38.fa --target chm13v2.0.fasta --outdir dir
-profile local --aligner minimap2
python chaintools/src/split.py -c input.chain -o input-split.chain
python chaintools/src/to_paf.py -c input-split.chain -t target.fa -q query.fa
-o input-split.paf
awk '$1==$6' input-split.paf | rb break-paf --max-size 10000  | rb trim-paf -
r | rb invert | rb trim-paf -r | rb invert > out.paf
paf2chain -i out.paf > out.chain
python chaintools/src/invert.py -c out.chain -o out_inverted.chain
```

Rustybam trim-paf uses dynamic programming and the CIGAR string to find an optimal splitting point between overlapping alignments in the query sequence. It starts its trimming with the largest overlap and then recursively trims smaller overlaps (https://mrvollger.github.io/rustybam/#align-once).

Results were validated by using chaintools to confirm that there were no overlapping sequences with respect to both T2T-CHM13v2.0 and GRCh38 in the released chain file. In addition, trimmed alignments were visually inspected with SafFire (https://github.com/mrvollger/SafFire, commit aa16e43) to confirm their quality.

##### wfmash-based pipeline

In addition to the minimap2-based whole genome alignment, we applied a wfmash-based pipeline to validate the chain file. This pipeline starts with whole-genome aligning T2T-CHM13v2.0 and the masked and filtered GRCh38 assembly with the wfmash sequence aligner (https://github.com/ekg/wfmash, commit a36ab5f) (Marco-Sola et al. 2021), requiring 1-to-1 homologous regions at least 5 kb long and nucleotide identity of at least 95%. Similarly, the resulting chain was post-processed to obtain 1:1 alignments using rustybam and the paf2chain tool (https://github.com/AndreaGuarracino/paf2chain, commit f68eeca). All PAF files with full CIGAR strings were then inspected with SafFire for quality investigation. All the instructions to obtain the final PAF and CHAIN files can be found at the following repository: https://github.com/pangenome/chm13-grch38-liftover. The minimap2- and wfmash-based chains showed high consistency over the genomes.

##### Structurally variable region

Structurally variable regions were defined for regions containing structural variants and other regions challenging to do assembly-assembly alignments. Specifically, using alignment of the T2T-Y to GRCh38 with dipcall v0.3 (Li et al. 2018), any SVs > 49 bp and other regions where the alignment may not be correct due to large SVs were collected. Additionally, we added regions likely to be problematic for aligning and calling small variants: (1) Any Segmental duplications, Tandem repeats >10kb in length, and Satellite arrays (including 15 kb flanking sequence on each side) that contain a break in dipcall confident regions (i.e., partially covered repeats); (2) 15 kb on either side of gaps in GRCh38; (3) 15 kb on either side of any other breaks in the dip.bed; (4) SVs >=50 bp and any overlapping tandem repeats +50bp.

##### Variants on GRCh38 that will disappear when calling on T2T-CHM13v2.0

The dipcall results were used again to identify small variant differences between T2T-Y and GRCh38-Y after excluding any small variants overlapping the structurally variable region. After exclusions, the regions included 2,314 SNVs and 1,291 indels smaller than 50 bp in 16.6 Mbp on chrY, which are expected to not be called on CHM13v2.0 if they were called in an individual on GRCh38. The vcf and bed are available at https://ftp-trace.ncbi.nlm.nih.gov/ReferenceSamples/giab/data/AshkenazimTrio/analysis/T2T-HG002-XY-v2.7/.

#### Datasets and resources for T2T-CHM13v2.0

##### Lifting over resources from GRCh38 to CHM13v2.0

Using the curated chain file, we lifted over dbSNP build 155 (ftp.ncbi.nih.gov/snp/archive/b155/) (Sherry, Ward, and Sirotkin 1999), the March 13, 2022 release of Clinvar (ftp://ftp.ncbi.nlm.nih.gov/pub/clinvar/vcf_GRCh38/weekly/) (Landrum et al. 2020; 2018), and GWAS Catalog v1.0 (https://ebi.ac.uk/gwas/home, accessed March 8th, 2022) (Welter et al. 2014; Buniello et al. 2019) from the GRCh38 primary assembly to T2T-CHM13v2.0. To generate a VCF file for portion of the GWAS Catalog that could be lifted over, we first identified all RefSeq IDs (rsIDs) in the v1.0 associations file (https://ebi.ac.uk/gwas/api/search/downloads/full), in the SNPS, SNP_ID_CURRENT, and STRONGEST SNP-RISK ALLELE fields. This yielded 192,121 unique rsIDs, which we then intersected with all dbSNP variants on the primary contigs for Chromosomes 1-22, Chromosomes X and Y, subsetting to those rsIDs that matched only a single record on each chromosome in dbSNP. This resulted in 189,051 unique variants from the GWAS Catalog that were used for liftover.

Liftover was performed as previously described (Aganezov et al. 2022). Specifically, we performed liftover using the GATK v4.1.9 LiftoverVcf, Picard v2.23.3 (Van der Auwera GA and O'Connor BD 2020) tool with the default parameters using the GRCh38 to T2T-CHM13v2.0 chain file provided here. This successfully lifts over variants that map exactly from GRCh38 to T2T-CHM13v2.0 but does not recover variants with swapped reference and alternative alleles. To recover variants with swapped reference/alternative alleles, we ran LiftoverVcf again, with the "RECOVER_SWAPPED_REF_ALT" flag. Notably, this feature does not recover multiallelic variants, so to recover these variants, we first separated them into multiple biallelic variants, performed liftover using the "RECOVER_SWAPPED_REF_ALT" flag, and converted them back to their multiallelic representations. Variants whose position lifted over, but whose reference allele or any alternative alleles did not match the T2T-CHM13v2.0 allele were not recovered.

#### ENCODE

Prior to mapping, reads were obtained from the ENCODE dataset (Dunham et al. 2012) (https://www.encodeproject.org/), and reads originating from a single library were combined. Reads were mapped with Bowtie2 v2.4.1 (Langmead and Salzberg 2012) as paired-end with the arguments "--no-discordant --no-mixed --very-sensitive --no-unal --omit-sec-seq --xeq --reorder". Alignments were filtered using SAMtools v1.10 using the arguments "-F 1804 -f 2 -q 2" to remove unmapped or single end mapped reads and those with a mapping quality score less than 2. PCR duplicates were identified and removed with the Picard tools "mark duplicates" command v2.22.1 and the arguments "VALIDATION_STRINGENCY=LENIENT ASSUME_SORT_ORDER=queryname REMOVE_DUPLICATES = true".

Alignments were then filtered for the presence of unique k-mers. Specifically, for each alignment, reference sequences aligned with template ends were compared to a database of minimum unique k-mer lengths. The size of the k-mers in the k-mer filtering step are dependent on the length of the mapped reference sequence. Alignments were discarded if no unique k-mers occurred in either end of the read. The minimum unique k-mer length database was generated using scripts found here. Alignments from replicates were then pooled.

Bigwig coverage tracks were created using deepTools2 bamCoverage v3.4.3 (Ramírez et al. 2016) with a bin size of 1bp and default for all other parameters. Enrichment tracks were created using deepTools bamCompare with a bin size of 50 bp, a pseudo-count of 1, and excluding bins with zero counts in both target and control tracks.

Peak calls were made using MACS2 v2.2.7.1 with default parameters and estimated genome sizes 3.03e9 and 2.79e9 for CHM13 and GRCh38, respectively. GRCh38 peak calls were lifted over to CHM13 using the UCSC liftOver utility, the chain file created by the T2T consortium, and the parameter "-minMatch=0.2".

##### gnomAD

Genome wide variant data from the Genome Aggregation Database (gnomAD) release v3.1.2 was lifted over from GRCh38 to each assembly using CrossMap v0.6.1 (Zhao et al. 2014). The chain files used were created from the GRCh38-based HAL file, downloaded from the Minigraph-Cactus alignment v1.0 of Liao et al. 2022 (Liao et al. 2022) (https://github.com/human-pangenomics/hpp_pangenome_resources/blob/main/hprc-v1.0-mc.md). The resulting VCFs were annotated with predicted molecular consequence and transcript-specific variant deleteriousness scores from PolyPhen-2 and SIFT using Ensembl Variant Effect Predictor. We created a NextFlow pipeline utilizing a containerised instance of Ensembl VEP to enable efficient annotation at scale (https://github.com/Ensembl/ensembl-vep/tree/release/108/nextflow, commit cb84684). All data can be found at https://projects.ensembl.org/hprc/.

#### Human Y chromosome contamination in bacterial genomes

##### Screening against Chrisman *et al.* study

We used the MUMmer v4.0 package (Marçais et al. 2018) to compare 73,691 bacterial 100-mers reported as enriched in human males by Chrisman *et al.* to the T2T-Y chromosome. We found that, as predicted, more than 95% of the 100-mers had near-perfect matches, defined as an exact match of 50 bp or longer, to the complete T2T-Y sequence. The nucmer program from MUMmer was run with default options, except to specify -l 50 for an exact match length of 50 or more, and -c 50 so that it reported matches as short as 50 bp.

##### Screening with 64-mers

Meryl v1.3 (Rhie et al. 2020) (https://github.com/marbl/meryl) was used to compare 64-mers between NCBI RefSeq release 213 (July 2022) and Chromosome Y from T2T-CHM13v2.0 and GRCh38. Each bacteria contig was annotated with the number of matching k-mers in T2T-CHM13v2.0, in GRCh38, and the number of k-mers in the contig with a match (hits-per-query, hpq). Each position in the reference chromosomes was annotated with the multiplicity of the k-mer at that position in the RefSeq contigs (mers-per-base, mpb), and with the number of contigs containing the k-mer (queries-per-base, qpb).

```
meryl/bin/meryl k=64 count chm13v2.chrY.fasta output chm13v2.chrY.k64.meryl
meryl/bin/meryl k=64 count  grch38.chrY.fasta output  grch38.chrY.k64.meryl

for part in `seq -f %02g 1 1 31` ; do
  meryl/bin/position-lookup \
    -s chm13v2.chrY.fasta \
    -m chm13v2.chrY.k64.meryl \
    -hpq chm13v2/hpq.${part} \
    -mpb chm13v2/mpb.${part} \
    -qpb chm13v2/qpb.${part} \
```

```
    release213/bacteria/bacteria.${part}??.genomic.fna.gz
  > chm13v2/err.${part} 2>&1
  meryl/bin/position-lookup \
    -s genomes/grch38.chrY.fasta \
    -m genomes/grch38.chrY.k64.meryl \
    -hpq grch38/hpq.${part} \
    -mpb grch38/mpb.${part} \
    -qpb grch38/qpb.${part} \
    release213/bacteria/bacteria.${part}??.genomic.fna.gz
  > grch38/err.${part} 2>&1
done
```

The hits-per-query outputs were filtered to retain only contigs with more than 20 k-mer matches or with more than 10% of the contig sequence covered by k-mer matches.

```
cat chm13v2/hpq.* | awk '($2>20) || ($2>0 && $2/$3>0.1)' > chm13v2.matches
cat  grch38/hpq.* | awk '($2>20) || ($2>0 && $2/$3>0.1)' > grch38.matches
```

The queries-per-base outputs were combined and accumulated into 10 kb windows along each reference chromosome.

```
perl coverage.pl chm13v2/mpb.?? > chm13v2-mers-per-10k-window
perl coverage.pl chm13v2/qpb.?? > chm13v2-queries-per-10k-window
perl coverage.pl grch38/mpb.?? > grch38-mers-per-10k-window
perl coverage.pl grch38/qpb.?? > grch38-queries-per-10k-window
```

The per-10k-window files were then converted to an interval wiggle file, with the distinct number of RefSeq entries found in each window for visualization (**Extended Data Fig. 11a**).

Contig length distribution was obtained from the matches files. First, sequence entries were retrieved using seqrequester (version 'r95 fa5bdac1', https://github.com/marbl/seqrequester) to extract each fasta file to query against the 64-mers built from regions annotated as HSat1B and HSat3. No sequences had 64-mers found that were shared between HSat1B and HSat3.

```
# sequence extraction with seqrequester
cut -f4 ${ref}.matches > ${ref}.matches.list
seqrequester extract \
 -sequences ${ref}.matches.list \
 release213/bacteria/bacteria.${part}??.genomic.fna.gz \
> ${ref}.matches.$part.fa

# collect 64-mers from HSat3 and HSat1B
awk '$1=="chrY"' chm13v2.0_censat_v2.0.bed > chrY_censat.bed
awk '$4 ~/hsat1B_Y/ {print $1":"$2+1"-"$3}' chrY_censat.bed > HSat1B.list
awk '$4 ~/hsat3_Y/  {print $1":"$2+1"-"$3}' chrY_censat.bed > HSat3.list

samtools faidx -r HSat3.list v2.7.Y.fasta  > HSat3.fa
samtools faidx -r HSat1B.list v2.7.Y.fasta > HSat1B.fa

meryl count k=64 HSat1B.fa output HSat1B.meryl
meryl count k=64 HSat3.fa  output HSat3.meryl
meryl intersect HSat1B.meryl HSat3.meryl output HSat1B_HSat3.meryl

for seq in grch38.matches chm13v2.matches
do
```

```
 meryl-lookup -existence -sequence $seq.fa \
  -mers HSat1B.meryl HSat3.meryl HSat1B_HSat3.meryl \
  > $seq.hsat
 awk '{if ($8>0) {HSAT="HSat1B_HSat3"} else if ($4>0) {HSAT="HSat1B"} else if
($6>0) {HSAT="HSat3"} else {HSAT="Others"} {print $1"\t"$2+63"\t"HSAT}}'
$seq.hsat > $seq.hsat.category
done
```

Next, sequences found only in T2T-Y were retrieved, by removing matches found in both T2T-Y and GRCh38-Y.

```
cut -f1 grch38.matches.category > grch38.matches.hsat.category.list
java -jar -Xmx256m txtContains.jar \
 chm13v2.matches.hsat.category grch38.matches.hsat.category.list 1 \
 > both-chm13v2-grch38.matches.hsat.category
java -jar -Xmx256m txtGrepv.jar \
 grch38.matches.hsat.category.list chm13v2.matches.hsat.category 1 \
 > chm13v2.matches.hsat.category.chm13v2_only
```

The .category files contain RefSeq sequence entries, number of 64-mers of the sequence, category annotation as “HSat1B”, “HSat3”, or “Others”. Sequence length distribution by categories were then visualized in **Extended Data Fig. 11b**.

Lastly, the complete sequence names of the entries in chm13v2.matches.hsat.category.chm13v2_only were retrieved from the original fasta files, which can be found in **Supplementary Table 32**. The first and second words in the names were extracted to visualize the taxonomic abundance of the microbial genomes in a pie chart using Kronatools v2.8 (Ondov, Bergman, and Phillippy 2011) ktImportText with -q option as the input file does not contain quantity field (**Extended Data Fig. 11c**):

```
# extract organism names for each contig name
perl extract-names.pl < chm13v2only.names \
> chm13v2only.names.labels # <- in Supplementary Table 32
awk -F " " '{print $3"\t"$4}' chm13v2only.names.labels \
> chm13v2only.names.labels.krona

# Kronatools
ktImportText -q chm13v2only.names.labels.krona
```

Output data files and additional scripts are at https://github.com/arangrhie/T2T-HG002Y/refseq-contamination/ .

### Supplementary Notes

#### 1. Transduced genomic segments

Transcriptionally active transposons, especially long interspersed element 1 (L1) and short variable number tandem repeat interspersed elements (SINE-VNTR-*Alu*s, SVA), occasionally co-mobilize downstream DNA into a new locus by bypassing a canonical poly(A) termination signal in a process termed 3' DNA transduction (Hoyt et al. 2022; Goodier, Ostertag, and Kazazian Jr 2000; Xing et al. 2006; Pickeral et al. 2000). In contrast, SVAs can also produce 5' DNA transductions by hijacking alternative promoters (Damert et al. 2009; Hoyt et al. 2022). Transduction activity has been recognized as a driver of genome shuffling that includes protein-coding sequences, regulatory elements, and even whole genes (Xing et al. 2006; Pickeral et al. 2000; Moran, DeBerardinis, and Kazazian 1999). Additionally, transductions can occur in the soma, contributing to tumorigenesis, and frequent 3' L1-mediated DNA transductions have been observed in some cancer types (Tubio et al. 2014). To identify potential DNA transduction activities in T2T-Y, we searched for transductions mediated by L1s and SVAs.

We detected six potential 3' L1 transductions within the Y, yet no SVA-driven DNA transductions (**Supplementary Table 16**). Our results show that four L1s carrying transduced segments are full-length elements (>6 kb), two of which possess a canonical poly(A) termination signal within 30 bases upstream of a poly(A) tail. The other two L1s are truncated elements with 3'-transduction signatures, consistent with prior predictions(Xing et al. 2006) since many retroposed L1s are truncated at the 5' end. We found that five L1 transductions were shared with GRCh38-Y, while one is specific to T2T-Y. Despite a genome-wide investigation of both T2T-CHM13+Y and GRCh38, we were not able to locate the potential donor elements of the transduced segments according to our criteria (e.g., they were not within 20 bases downstream of a full-length L1 in a different locus).

To investigate whether any source elements within T2T-Y gave rise to DNA transductions onto other chromosomes, we probed previously reported L1 and SVA transductions (Hoyt et al. 2022). However, we were not able to detect any sign of the source elements. Given the ancient form of L1s annotated on the Y, we confirm a recent analysis (Halabian and Makałowski 2022) of 1KGP data that found no evidence for DNA transduction between the Y and the rest of the chromosomes. Our analysis revealed that the transduction rate in T2T-Y was 0.096 per 1 Mb, which is much lower than the transduction rate observed in the CHM13 autosomes (avg. 6.9 per 1 Mb) and ChrX (10.19 per 1 Mb) (Hoyt et al. 2022). In conclusion, our results indicate that transposable element driven transductions are not abundant in the Y, and traffic of these events is low between this chromosome and the rest of the genome.

### 2. Non-B DNA motifs

We located Y chromosome motifs capable of forming alternative DNA structures (non-B DNA) such as bent helix, slipped strand, G-quadruplex (G4s), cruciform, triple helix, hairpin, and Z-DNA structures. Non-B DNA structures are known to affect a variety of important cellular processes, such as replication, gene expression, and genome stability (Ghosh and Bansal 2003; A. Jain, Wang, and Vasquez 2008; Wang and Vasquez 2014; Varshney et al. 2020). We found inverted, A-phased, and mirror repeats to be abundant in the centromeric alpha satellite HOR, forming a periodic pattern occurring every 5.7 kb (**Fig. 3,** and **Supplementary Table 17**). An additional pair of A-phased and inverted repeats was present in the extended light blue HOR variant, suggesting possible non-B-form variations per HOR SVs. Moreover, the per-base-pair density of A-phased, direct, inverted, mirror, and short tandem repeats (STRs) is higher in the newly completed regions of the Y chromosome, and the density of Z-DNA and G4 motifs was particularly high in the newly completed *TSPY* gene array (**Extended Data Fig. 5, 6b** and **Supplementary Table 17**). Among the 762 new G4 motifs in T2T-Y, 519 are located within the *TSPY* gene array. Specifically, each *TSPY* composite repeat unit contains 14 G4 motifs, seven of which are stable according to their Quadron score (≥19)(Sahakyan et al. 2017). Among these, one is located ~500 bases upstream of the transcription start site on the same strand, which might indicate a role in transcriptional regulation(Varshney et al. 2020) (**Extended Data Fig. 6b**). Along the entire T2T-Y chromosome, 242 G4 motifs overlapped CpG islands (by at least 1 base), again suggesting a role in transcription.

### 3. PAR and XTR masking for sex chromosome complement analysis

Before using T2T-CHM13+Y as an alternate reference, we investigated whether masking PARs or XTRs on ChrY would improve mapping quality (MQ) and variant calling accuracy on the sex chromosomes. Genetic diversity is higher within PAR1 compared to the non-recombining regions on ChrY (Cotter, Brotman, and Wilson Sayres 2016; Lien et al. 2000); however, the ChrX and ChrY PAR sequences in GRCh38 are represented as identical copies (Genome Reference Consortium n.d.). The perfect identity between GRCh38 PARs reduces MQ, hindering accurate variant detection in this region. Previously, the impact of hard-masking the entire ChrY for samples with an XX karyotype was shown to improve mapping quality and increase the number of variants called; however, this was not tested extensively on samples with an XY karyotype (Webster et al. 2019). Thus, we simulated 20x coverage of 150 base Illumina reads for ten XY samples (10x coverage for each ChrX and ChrY) seeded with variants called from the high coverage samples in 1KGP, and tested if the PAR masking improves MQ and variant calling accuracy (**Supplementary Table 25-26**).

In the XY samples, the simulated read alignments showed a near-zero MQ on the PARs when no masking was applied, with almost no variants called. In comparison, when masking the PARs on ChrY, reads aligned with improved mapping quality across all samples (example of one sample shown in **Extended Data Fig. 9**), calling an average of 4,615 true positive variants on PAR1 and

365 on PAR2, respectively, with almost no false positives (**Supplementary Table 27**). Additionally, we tested the impact of masking the XTR on ChrY. Unlike the PARs, the false positives substantially increased from an average of 1 to 33,345 across the ChrX XTR (**Supplementary Table 28**), indicating that mapping and variant calling was improved by masking PARs but that XTR masking was detrimental.

### 4. Different CpG Methylation pattern in HiFi and ONT at Yq12

In general, 5-methylcytosine (5mC) at CpG sites showed good agreement genome-wide between different technologies when using T2T-CHM13v1.1 autosomes and the XY from HG002XYv2.7 (HG002-X and HG002-Y) as a reference (**Supplementary Fig. 19**). This included CpG methylation calls from PacBio HiFi reads using Primrose v1.3.0, ONT reads from nanoNOMe experiments called with Nanopolish (Guppy 5.0.7, with the methylation calling pipeline described in Gershman *et al*. (Gershman et al. 2022)), and ONT reads called with Remora and Guppy 6.1.2 (**Supplementary Table 1**), as well as methylation calls made using short-read methods: Enzymatic Methyl-seq (EM-seq) and Whole Genome Bisulfite Sequencing (WGBS) generated by Foox *et al*. (Foox et al. 2021). When we examined the correlation between technologies in regions with at least 5 reads in all 6 technologies, we found slightly lower correlation between the ONT Nanopolish CpG calls and WGBS than between ONT Remora and WGBS (**Supplementary Fig. 19a-b,d**). This might be due to Nanopolish using a “four-state” model for nanoNOMe experiments, in which reads have both native CpG methylation as well as exogenous GpC methylation marking open chromatin regions. The “four-state” model takes the methylation status of neighboring CpG and GpC sites into account when calling methylation (Simpson et al. 2017) and this is expected to reduce the accuracy of CpG calls in comparison to calls where no GpC methylation is present. Both PacBio Primrose and ONT Remora calls had good correlation to WGBS, with ONT Remora having a slightly tighter correlation range and higher value of r (0.957 vs. 0.912, **Supplementary Fig. 19c-d**). However, we found inconsistent methylation patterns within the heterochromatic satellite region on the q-arm of ChrY (Yqh region). PacBio Primrose called higher methylation frequency on HSat1, while ONT Nanopolish and Remora showed higher methylation frequency on HSat3. Because of the absence of a known truth, and difficulties in mapping Illumina-reads correctly to these satellites, it is unclear which pattern is correct.

One possible explanation for the low correlation in the Yqh region is that sequencing biases are affecting the methylation calls. Previously, we had observed enriched coverage across HSat3 in the HiFi reads, whereas the coverage was depleted across HSat1 in the ONT reads (Fig. 3 from Nurk et al. (Nurk et al. 2022)). Similarly, we observed coverage biases in the satellites of Yqh (**Supplementary Fig. 19e**). In general, HiFi reads show high sequence similarity to the assembled T2T-Y, with a slight drop in HSat1 (>99.6% in HSat1B vs. >99.9 in HSat3), while ONT shows wider fluctuations, but again with higher sequence identity to HSat3 (>97% in HSat1B vs. >98% in HSat3). As shown in **Fig. 5c**, HSat1B contains AT-enriched microsatellite regions (homopolymer compressed sequence composed of only A and T bases, enrichment here is >80%), which are prone to sequencing errors in both HiFi and ONT. This may have affected these technologies’ ability to detect 5mC in these regions.

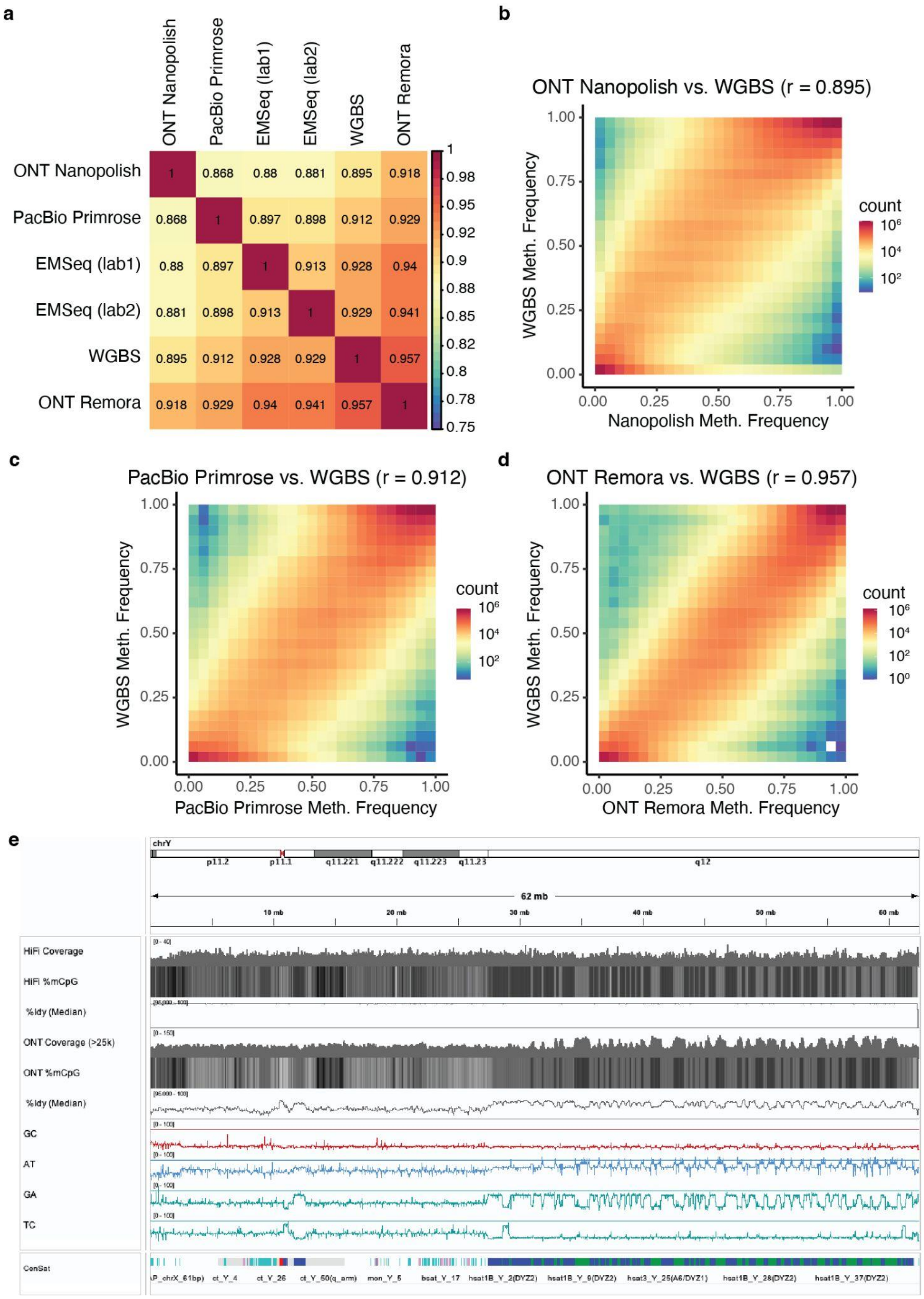


**Supplementary Fig. 19 | Methylation frequency differences in HiFi and ONT reads. a.** Genome-wide 5mC agreement between different technologies at CpG sites. **b.** Correlation between ONT Nanopolish vs. WGBS. **c.** Correlation between PacBio Primrose vs. WGBS. **d.** Correlation between ONT Remora vs. WGBS. **e.** HiFi and ONT sequencing coverage and quality biases illustrated across the Yq12 region. GC/AT/GA/TC tracks show the microsatellite sequence composition.

Chin, Chen-Shan, Sairam Behera, Asif Khalak, Fritz J. Sedlazeck, Justin Wagner, and Justin M.

Zook. 2022. “Multiscale Analysis of Pangenome Enables Improved Representation of Genomic Diversity For Repetitive And Clinically Relevant Genes.” *BioRxiv*, August, 2022.08.05.502980. https://doi.org/10.1101/2022.08.05.502980.
Chin, Chen-Shan, Justin Wagner, Qiandong Zeng, Erik Garrison, Shilpa Garg, Arkarachai Fungtammasan, Mikko Rautiainen, et al. 2020. “A Diploid Assembly-Based Benchmark for Variants in the Major Histocompatibility Complex.” *Nature Communications* 11 (1): 4794. https://doi.org/10.1038/s41467-020-18564-9.
Cortez, Diego, Ray Marin, Deborah Toledo-Flores, Laure Froidevaux, Angélica Liechti, Paul D. Waters, Frank Grützner, and Henrik Kaessmann. 2014. “Origins and Functional Evolution of Y Chromosomes across Mammals.” *Nature* 508 (7497): 488–93. https://doi.org/10.1038/nature13151.
Cotter, Daniel J, Sarah M Brotman, and Melissa A Wilson Sayres. 2016. “Genetic Diversity on the Human X Chromosome Does Not Support a Strict Pseudoautosomal Boundary.” *Genetics* 203 (1): 485–92. https://doi.org/10.1534/genetics.114.172692.
Daily, Jeff. 2016. “Parasail: SIMD C Library for Global, Semi-Global, and Local Pairwise Sequence Alignments.” *BMC Bioinformatics* 17 (1): 81. https://doi.org/10.1186/s12859-016-0930-z.
Dale, Ryan K., Brent S. Pedersen, and Aaron R. Quinlan. 2011. “Pybedtools: A Flexible Python Library for Manipulating Genomic Datasets and Annotations.” *Bioinformatics* 27 (24): 3423–24. https://doi.org/10.1093/bioinformatics/btr539.
Damert, Annette, Julija Raiz, Axel V. Horn, Johannes Löwer, Hui Wang, Jinchuan Xing, Mark A. Batzer, Roswitha Löwer, and Gerald G. Schumann. 2009. “5′-Transducing SVA Retrotransposon Groups Spread Efficiently throughout the Human Genome.” *Genome Research* 19 (11): 1992–2008. https://doi.org/10.1101/gr.093435.109.
Danecek, Petr, James K Bonfield, Jennifer Liddle, John Marshall, Valeriu Ohan, Martin O Pollard, Andrew Whitwham, et al. 2021. “Twelve Years of SAMtools and BCFtools.” *GigaScience* 10 (2): giab008. https://doi.org/10.1093/gigascience/giab008.
Dunham, Ian, Anshul Kundaje, Shelley F. Aldred, Patrick J. Collins, Carrie A. Davis, Francis Doyle, Charles B. Epstein, et al. 2012. “An Integrated Encyclopedia of DNA Elements in the Human Genome.” *Nature* 489 (7414): 57–74. https://doi.org/10.1038/nature11247.
Ebert, Peter, Peter A. Audano, Qihui Zhu, Bernardo Rodriguez-Martin, David Porubsky, Marc Jan Bonder, Arvis Sulovari, et al. 2021. “Haplotype-Resolved Diverse Human Genomes and Integrated Analysis of Structural Variation.” *Science (New York, N.Y.)* 372 (6537): eabf7117. https://doi.org/10.1126/science.abf7117.
English, Adam C., Vipin K. Menon, Richard Gibbs, Ginger A. Metcalf, and Fritz J. Sedlazeck. 2022. “Truvari: Refined Structural Variant Comparison Preserves Allelic Diversity.” *BioRxiv*, February, 2022.02.21.481353. https://doi.org/10.1101/2022.02.21.481353.
Escalona, Merly, Jake VanCampen, Nicholas W Maurer, Marina Haukness, Mariam Okhovat, Robert S Harris, Allison Watwood, et al. 2023. “Whole-Genome Sequence and Assembly of the Javan Gibbon (Hylobates Moloch).” *Journal of Heredity* 114 (1): 35–43. https://doi.org/10.1093/jhered/esac043.
“FigTree. Last Accessed: 2022-11-28.” n.d. Accessed November 28, 2022. http://tree.bio.ed.ac.uk/software/figtree/.
Foox, Jonathan, Jessica Nordlund, Claudia Lalancette, Ting Gong, Michelle Lacey, Samantha Lent, Bradley W. Langhorst, et al. 2021. “The SEQC2 Epigenomics Quality Control (EpiQC) Study.” *Genome Biology* 22 (1): 332. https://doi.org/10.1186/s13059-021-02529-2.
Formenti, Giulio, Arang Rhie, Brian P. Walenz, Françoise Thibaud-Nissen, Kishwar Shafin, Sergey Koren, Eugene W. Myers, Erich D. Jarvis, and Adam M. Phillippy. 2022. “Merfin: Improved Variant Filtering, Assembly Evaluation and Polishing via k-Mer Validation.” *Nature Methods* 19 (6): 696–704. https://doi.org/10.1038/s41592-022-01445-y.

Frankish, Adam, Mark Diekhans, Irwin Jungreis, Julien Lagarde, Jane E Loveland, Jonathan M Mudge, Cristina Sisu, et al. 2021. “GENCODE 2021.” *Nucleic Acids Research* 49 (D1): D916–23. https://doi.org/10.1093/nar/gkaa1087.
Gálvez-Merchán, Ángel, Kyung Hoi (Joseph) Min, Lior Pachter, and A. Sina Booeshaghi. 2022. “Metadata Retrieval from Sequence Databases with Ffq.” bioRxiv. https://doi.org/10.1101/2022.05.18.492548.
Gamaarachchi, Hasindu, Chun Wai Lam, Gihan Jayatilaka, Hiruna Samarakoon, Jared T. Simpson, Martin A. Smith, and Sri Parameswaran. 2020. “GPU Accelerated Adaptive Banded Event Alignment for Rapid Comparative Nanopore Signal Analysis.” *BMC Bioinformatics* 21 (1): 343. https://doi.org/10.1186/s12859-020-03697-x.
Geneious. n.d. “Bioinformatics Software for Sequence Data Analysis. Last Accessed: 2022-11-28.” Geneious. Accessed November 28, 2022. https://www.geneious.com/.
Genome Reference Consortium. n.d. “Assembly Terminology. Last Accessed: 2022-11-29.” Accessed March 27, 2022. https://www.ncbi.nlm.nih.gov/grc/help/definitions/.
Gershman, Ariel, Michael E. G. Sauria, Xavi Guitart, Mitchell R. Vollger, Paul W. Hook, Savannah J. Hoyt, Miten Jain, et al. 2022. “Epigenetic Patterns in a Complete Human Genome.” *Science* 376 (6588): eabj5089. https://doi.org/10.1126/science.abj5089.
Ghosh, A., and M. Bansal. 2003. “A Glossary of DNA Structures from A to Z.” *Acta Crystallographica Section D: Biological Crystallography* 59 (4): 620–26. https://doi.org/10.1107/S0907444903003251.
Goodier, John L., Eric M. Ostertag, and Haig H. Kazazian Jr. 2000. “Transduction of 3′-Flanking Sequences Is Common in L1 Retrotransposition.” *Human Molecular Genetics* 9 (4): 653–57. https://doi.org/10.1093/hmg/9.4.653.
“GraphPad Prism Version v9.1.0 for Windows, GraphPad Software, San Diego, California, USA. Last Accessed: 2022-11-28.” n.d. Accessed November 28, 2022. https://www.graphpad.com.
Hach, Faraz, Fereydoun Hormozdiari, Can Alkan, Farhad Hormozdiari, Inanc Birol, Evan E. Eichler, and S. Cenk Sahinalp. 2010. “MrsFAST: A Cache-Oblivious Algorithm for Short-Read Mapping.” *Nature Methods* 7 (8): 576–77. https://doi.org/10.1038/nmeth0810-576.
Halabian, Reza, and Wojciech Makałowski. 2022. “A Map of 3′ DNA Transduction Variants Mediated by Non-LTR Retroelements on 3202 Human Genomes.” *Biology* 11 (7): 1032. https://doi.org/10.3390/biology11071032.
Hansen, Kasper D., Benjamin Langmead, and Rafael A. Irizarry. 2012. “BSmooth: From Whole Genome Bisulfite Sequencing Reads to Differentially Methylated Regions.” *Genome Biology* 13 (10): R83. https://doi.org/10.1186/gb-2012-13-10-r83.
Hao, Zhaodong, Dekang Lv, Ying Ge, Jisen Shi, Dolf Weijers, Guangchuang Yu, and Jinhui Chen. 2020. “RIdeogram: Drawing SVG Graphics to Visualize and Map Genome-Wide Data on the Idiograms.” *PeerJ Computer Science* 6 (January): e251. https://doi.org/10.7717/peerj-cs.251.
Harris, Robert S. 2007. “Improved Pairwise Alignmnet of Genomic DNA.” Penn State. https://etda.libraries.psu.edu/catalog/7971.
Hoyt, Savannah J., Jessica M. Storer, Gabrielle A. Hartley, Patrick G. S. Grady, Ariel Gershman, Leonardo G. de Lima, Charles Limouse, et al. 2022. “From Telomere to Telomere: The Transcriptional and Epigenetic State of Human Repeat Elements.” *Science* 376 (6588): eabk3112. https://doi.org/10.1126/science.abk3112.
Hui, Joseph, Ilan Shomorony, Kannan Ramchandran, and Thomas A. Courtade. 2016. “Overlap-Based Genome Assembly from Variable-Length Reads.” In *2016 IEEE International Symposium on Information Theory (ISIT)*, 1018–22. https://doi.org/10.1109/ISIT.2016.7541453.
Jain, Aklank, Guliang Wang, and Karen M. Vasquez. 2008. “DNA Triple Helices: Biological Consequences and Therapeutic Potential.” *Biochimie*, Targeting DNA Part II, 90 (8):

1117–30. https://doi.org/10.1016/j.biochi.2008.02.011.
Jain, Chirag, Arang Rhie, Nancy F. Hansen, Sergey Koren, and Adam M. Phillippy. 2022. “Long-Read Mapping to Repetitive Reference Sequences Using Winnowmap2.” *Nature Methods* 19 (6): 705–10. https://doi.org/10.1038/s41592-022-01457-8.
Jain, Miten, Sergey Koren, Karen H. Miga, Josh Quick, Arthur C. Rand, Thomas A. Sasani, John R. Tyson, et al. 2018. “Nanopore Sequencing and Assembly of a Human Genome with Ultra-Long Reads.” *Nature Biotechnology* 36 (4): 338–45. https://doi.org/10.1038/nbt.4060.
Jain, Miten, Hugh E. Olsen, Daniel J. Turner, David Stoddart, Kira V. Bulazel, Benedict Paten, David Haussler, Huntington F. Willard, Mark Akeson, and Karen H. Miga. 2018. “Linear Assembly of a Human Centromere on the Y Chromosome.” *Nature Biotechnology* 36 (4): 321–23. https://doi.org/10.1038/nbt.4109.
Jarvis, Erich D., Giulio Formenti, Arang Rhie, Andrea Guarracino, Chentao Yang, Jonathan Wood, Alan Tracey, et al. 2022. “Semi-Automated Assembly of High-Quality Diploid Human Reference Genomes.” *Nature* 611 (7936): 519–31. https://doi.org/10.1038/s41586-022-05325-5.
Jeffares, Daniel C., Clemency Jolly, Mimoza Hoti, Doug Speed, Liam Shaw, Charalampos Rallis, Francois Balloux, Christophe Dessimoz, Jürg Bähler, and Fritz J. Sedlazeck. 2017. “Transient Structural Variations Have Strong Effects on Quantitative Traits and Reproductive Isolation in Fission Yeast.” *Nature Communications* 8 (1): 14061. https://doi.org/10.1038/ncomms14061.
Jiang, Tao, Yongzhuang Liu, Yue Jiang, Junyi Li, Yan Gao, Zhe Cui, Yadong Liu, Bo Liu, and Yadong Wang. 2020. “Long-Read-Based Human Genomic Structural Variation Detection with CuteSV.” *Genome Biology* 21 (1): 189. https://doi.org/10.1186/s13059-020-02107-y.
Jiang, Zhaoshi, Robert Hubley, Arian Smit, and Evan E. Eichler. 2008. “DupMasker: A Tool for Annotating Primate Segmental Duplications.” *Genome Research* 18 (8): 1362–68. https://doi.org/10.1101/gr.078477.108.
Kapustin, Yuri, Alexander Souvorov, Tatiana Tatusova, and David Lipman. 2008. “Splign: Algorithms for Computing Spliced Alignments with Identification of Paralogs.” *Biology Direct* 3 (1): 20. https://doi.org/10.1186/1745-6150-3-20.
Katoh, Kazutaka, and Daron M. Standley. 2014. “MAFFT: Iterative Refinement and Additional Methods.” In *Multiple Sequence Alignment Methods*, edited by David J Russell, 131–46. Methods in Molecular Biology. Totowa, NJ: Humana Press. https://doi.org/10.1007/978-1-62703-646-7_8.
Kent, W. James. 2002. “BLAT—The BLAST-Like Alignment Tool.” *Genome Research* 12 (4): 656–64. https://doi.org/10.1101/gr.229202.
Kent, W. James, Charles W. Sugnet, Terrence S. Furey, Krishna M. Roskin, Tom H. Pringle, Alan M. Zahler, and and David Haussler. 2002. “The Human Genome Browser at UCSC.” *Genome Research* 12 (6): 996–1006. https://doi.org/10.1101/gr.229102.
Kolmogorov, Mikhail, Jeffrey Yuan, Yu Lin, and Pavel A. Pevzner. 2019. “Assembly of Long, Error-Prone Reads Using Repeat Graphs.” *Nature Biotechnology* 37 (5): 540–46. https://doi.org/10.1038/s41587-019-0072-8.
Kovaka, Sam, Aleksey V. Zimin, Geo M. Pertea, Roham Razaghi, Steven L. Salzberg, and Mihaela Pertea. 2019. “Transcriptome Assembly from Long-Read RNA-Seq Alignments with StringTie2.” *Genome Biology* 20 (1): 278. https://doi.org/10.1186/s13059-019-1910-1.
Krueger, Felix, and Simon R. Andrews. 2011. “Bismark: A Flexible Aligner and Methylation Caller for Bisulfite-Seq Applications.” *Bioinformatics* 27 (11): 1571–72. https://doi.org/10.1093/bioinformatics/btr167.
Krumsiek, Jan, Roland Arnold, and Thomas Rattei. 2007. “Gepard: A Rapid and Sensitive Tool for Creating Dotplots on Genome Scale.” *Bioinformatics* 23 (8): 1026–28.

https://doi.org/10.1093/bioinformatics/btm039.
Landrum, Melissa J, Shanmuga Chitipiralla, Garth R Brown, Chao Chen, Baoshan Gu, Jennifer Hart, Douglas Hoffman, et al. 2020. “ClinVar: Improvements to Accessing Data.” *Nucleic Acids Research* 48 (D1): D835–44. https://doi.org/10.1093/nar/gkz972.
Landrum, Melissa J, Jennifer M Lee, Mark Benson, Garth R Brown, Chen Chao, Shanmuga Chitipiralla, Baoshan Gu, et al. 2018. “ClinVar: Improving Access to Variant Interpretations and Supporting Evidence.” *Nucleic Acids Research* 46 (D1): D1062–67. https://doi.org/10.1093/nar/gkx1153.
Langmead, Ben, and Steven L. Salzberg. 2012. “Fast Gapped-Read Alignment with Bowtie 2.” *Nature Methods* 9 (4): 357–59. https://doi.org/10.1038/nmeth.1923.
Lassmann, Timo. 2020. “Kalign 3: Multiple Sequence Alignment of Large Datasets.” *Bioinformatics* 36 (6): 1928–29. https://doi.org/10.1093/bioinformatics/btz795.
Lawrence, Michael, Robert Gentleman, and Vincent Carey. 2009. “Rtracklayer: An R Package for Interfacing with Genome Browsers.” *Bioinformatics* 25 (14): 1841–42. https://doi.org/10.1093/bioinformatics/btp328.
Lawrence, Michael, Wolfgang Huber, Hervé Pagès, Patrick Aboyoun, Marc Carlson, Robert Gentleman, Martin T. Morgan, and Vincent J. Carey. 2013. “Software for Computing and Annotating Genomic Ranges.” *PLOS Computational Biology* 9 (8): e1003118. https://doi.org/10.1371/journal.pcbi.1003118.
Lee, Isac, Roham Razaghi, Timothy Gilpatrick, Michael Molnar, Ariel Gershman, Norah Sadowski, Fritz J. Sedlazeck, Kasper D. Hansen, Jared T. Simpson, and Winston Timp. 2020. “Simultaneous Profiling of Chromatin Accessibility and Methylation on Human Cell Lines with Nanopore Sequencing.” *Nature Methods* 17 (12): 1191–99. https://doi.org/10.1038/s41592-020-01000-7.
Li, Heng. 2018. “Minimap2: Pairwise Alignment for Nucleotide Sequences.” *Bioinformatics* 34 (18): 3094–3100. https://doi.org/10.1093/bioinformatics/bty191.
———. 2019. “Identifying Centromeric Satellites with Dna-Brnn.” *Bioinformatics* 35 (21): 4408–10. https://doi.org/10.1093/bioinformatics/btz264.
Li, Heng, Jonathan M. Bloom, Yossi Farjoun, Mark Fleharty, Laura Gauthier, Benjamin Neale, and Daniel MacArthur. 2018. “A Synthetic-Diploid Benchmark for Accurate Variant-Calling Evaluation.” *Nature Methods* 15 (8): 595–97. https://doi.org/10.1038/s41592-018-0054-7.
Li, Heng, and Richard Durbin. 2009. “Fast and Accurate Short Read Alignment with Burrows–Wheeler Transform.” *Bioinformatics* 25 (14): 1754–60. https://doi.org/10.1093/bioinformatics/btp324.
Li, Heng, Bob Handsaker, Alec Wysoker, Tim Fennell, Jue Ruan, Nils Homer, Gabor Marth, Goncalo Abecasis, Richard Durbin, and 1000 Genome Project Data Processing Subgroup. 2009. “The Sequence Alignment/Map Format and SAMtools.” *Bioinformatics* 25 (16): 2078–79. https://doi.org/10.1093/bioinformatics/btp352.
Liao, Wen-Wei, Mobin Asri, Jana Ebler, Daniel Doerr, Marina Haukness, Glenn Hickey, Shuangjia Lu, et al. 2022. “A Draft Human Pangenome Reference.” *BioRxiv*, July, 2022.07.09.499321. https://doi.org/10.1101/2022.07.09.499321.
Lien, Sigbjørn, Joanna Szyda, Birgit Schechinger, Gudrun Rappold, and Norm Arnheim. 2000. “Evidence for Heterogeneity in Recombination in the Human Pseudoautosomal Region: High Resolution Analysis by Sperm Typing and Radiation-Hybrid Mapping.” *The American Journal of Human Genetics* 66 (2): 557–66. https://doi.org/10.1086/302754.
Liu, Bo, Yadong Liu, Junyi Li, Hongzhe Guo, Tianyi Zang, and Yadong Wang. 2019. “DeSALT: Fast and Accurate Long Transcriptomic Read Alignment with de Bruijn Graph-Based Index.” *Genome Biology* 20 (1): 274. https://doi.org/10.1186/s13059-019-1895-9.
Mallick, Swapan, Heng Li, Mark Lipson, Iain Mathieson, Melissa Gymrek, Fernando Racimo, Mengyao Zhao, et al. 2016. “The Simons Genome Diversity Project: 300 Genomes from

142 Diverse Populations." *Nature* 538 (7624): 201–6. https://doi.org/10.1038/nature18964.
Marçais, Guillaume, Arthur L. Delcher, Adam M. Phillippy, Rachel Coston, Steven L. Salzberg, and Aleksey Zimin. 2018. "MUMmer4: A Fast and Versatile Genome Alignment System." *PLOS Computational Biology* 14 (1): e1005944. https://doi.org/10.1371/journal.pcbi.1005944.
Marco-Sola, Santiago, Juan Carlos Moure, Miquel Moreto, and Antonio Espinosa. 2021. "Fast Gap-Affine Pairwise Alignment Using the Wavefront Algorithm." *Bioinformatics* 37 (4): 456–63. https://doi.org/10.1093/bioinformatics/btaa777.
Mc Cartney, Ann M., Kishwar Shafin, Michael Alonge, Andrey V. Bzikadze, Giulio Formenti, Arkarachai Fungtammasan, Kerstin Howe, et al. 2022. "Chasing Perfection: Validation and Polishing Strategies for Telomere-to-Telomere Genome Assemblies." *Nature Methods* 19 (6): 687–95. https://doi.org/10.1038/s41592-022-01440-3.
McKenna, Aaron, Matthew Hanna, Eric Banks, Andrey Sivachenko, Kristian Cibulskis, Andrew Kernytsky, Kiran Garimella, et al. 2010. "The Genome Analysis Toolkit: A MapReduce Framework for Analyzing next-Generation DNA Sequencing Data." *Genome Research* 20 (9): 1297–1303. https://doi.org/10.1101/gr.107524.110.
Miga, Karen H., Yulia Newton, Miten Jain, Nicolas Altemose, Huntington F. Willard, and W. James Kent. 2014. "Centromere Reference Models for Human Chromosomes X and Y Satellite Arrays." *Genome Research* 24 (4): 697–707. https://doi.org/10.1101/gr.159624.113.
Moran, John V., Ralph J. DeBerardinis, and Haig H. Kazazian. 1999. "Exon Shuffling by L1 Retrotransposition." *Science* 283 (5407): 1530–34. https://doi.org/10.1126/science.283.5407.1530.
"Mskilab/GUtils: R Package Providing Additional Capabilities and Speed for GenomicRanges Operations. Last Accessed: 2022-11-28." n.d. Accessed November 28, 2022. https://rdrr.io/github/mskilab/gUtils/.
Navarro-Costa, Paulo, Carlos E. Plancha, and João Gonçalves. 2010. "Genetic Dissection of the AZF Regions of the Human Y Chromosome: Thriller or Filler for Male (In)Fertility?" *BioMed Research International* 2010 (June): e936569. https://doi.org/10.1155/2010/936569.
Numanagić, Ibrahim, Alim S Gökkaya, Lillian Zhang, Bonnie Berger, Can Alkan, and Faraz Hach. 2018. "Fast Characterization of Segmental Duplications in Genome Assemblies." *Bioinformatics* 34 (17): i706–14. https://doi.org/10.1093/bioinformatics/bty586.
Nurk, Sergey, Sergey Koren, Arang Rhie, Mikko Rautiainen, Andrey V. Bzikadze, Alla Mikheenko, Mitchell R. Vollger, et al. 2022. "The Complete Sequence of a Human Genome." *Science* 376 (6588): 44–53. https://doi.org/10.1126/science.abj6987.
Nurk, Sergey, Brian P. Walenz, Arang Rhie, Mitchell R. Vollger, Glennis A. Logsdon, Robert Grothe, Karen H. Miga, Evan E. Eichler, Adam M. Phillippy, and Sergey Koren. 2020. "HiCanu: Accurate Assembly of Segmental Duplications, Satellites, and Allelic Variants from High-Fidelity Long Reads." *Genome Research*, August, gr.263566.120. https://doi.org/10.1101/gr.263566.120.
Olson, Daniel, and Travis Wheeler. 2018. "ULTRA: A Model Based Tool to Detect Tandem Repeats." In *Proceedings of the 2018 ACM International Conference on Bioinformatics, Computational Biology, and Health Informatics*, 37–46. BCB '18. New York, NY, USA: Association for Computing Machinery. https://doi.org/10.1145/3233547.3233604.
Olson, Nathan D., Justin Wagner, Jennifer McDaniel, Sarah H. Stephens, Samuel T. Westreich, Anish G. Prasanna, Elaine Johanson, et al. 2022. "PrecisionFDA Truth Challenge V2: Calling Variants from Short and Long Reads in Difficult-to-Map Regions." *Cell Genomics* 2 (5): 100129. https://doi.org/10.1016/j.xgen.2022.100129.
Ondov, Brian D., Nicholas H. Bergman, and Adam M. Phillippy. 2011. "Interactive Metagenomic

Visualization in a Web Browser.” *BMC Bioinformatics* 12 (1): 385. https://doi.org/10.1186/1471-2105-12-385.
Pagès, H., P. Aboyoun, R. Gentleman, and S. DebRoy. 2022. “Biostrings: Efficient Manipulation of Biological Strings.” Bioconductor version: Release (3.16). https://doi.org/10.18129/B9.bioc.Biostrings.
Pendleton, Amanda L., Feichen Shen, Angela M. Taravella, Sarah Emery, Krishna R. Veeramah, Adam R. Boyko, and Jeffrey M. Kidd. 2018. “Comparison of Village Dog and Wolf Genomes Highlights the Role of the Neural Crest in Dog Domestication.” *BMC Biology* 16 (1): 64. https://doi.org/10.1186/s12915-018-0535-2.
Pickeral, Oxana K., Wojciech Makałowski, Mark S. Boguski, and Jef D. Boeke. 2000. “Frequent Human Genomic DNA Transduction Driven by LINE-1 Retrotransposition.” *Genome Research* 10 (4): 411–15. https://doi.org/10.1101/gr.10.4.411.
Poplin, Ryan, Pi-Chuan Chang, David Alexander, Scott Schwartz, Thomas Colthurst, Alexander Ku, Dan Newburger, et al. 2018. “A Universal SNP and Small-Indel Variant Caller Using Deep Neural Networks.” *Nature Biotechnology* 36 (10): 983–87. https://doi.org/10.1038/nbt.4235.
Porubsky, David, Ashley D. Sanders, Wolfram Höps, PingHsun Hsieh, Arvis Sulovari, Ruiyang Li, Ludovica Mercuri, et al. 2020. “Recurrent Inversion Toggling and Great Ape Genome Evolution.” *Nature Genetics* 52 (8): 849–58. https://doi.org/10.1038/s41588-020-0646-x.
Porubsky, David, Ashley D Sanders, Aaron Taudt, Maria Colomé-Tatché, Peter M Lansdorp, and Victor Guryev. 2020. “BreakpointR: An R/Bioconductor Package to Localize Strand State Changes in Strand-Seq Data.” *Bioinformatics* 36 (4): 1260–61. https://doi.org/10.1093/bioinformatics/btz681.
Poznik, G. David. 2016. “Identifying Y-Chromosome Haplogroups in Arbitrarily Large Samples of Sequenced or Genotyped Men.” bioRxiv. https://doi.org/10.1101/088716.
Poznik, G. David, Brenna M. Henn, Muh-Ching Yee, Elzbieta Sliwerska, Ghia M. Euskirchen, Alice A. Lin, Michael Snyder, et al. 2013. “Sequencing Y Chromosomes Resolves Discrepancy in Time to Common Ancestor of Males Versus Females.” *Science* 341 (6145): 562–65. https://doi.org/10.1126/science.1237619.
Poznik, G. David, Yali Xue, Fernando L. Mendez, Thomas F. Willems, Andrea Massaia, Melissa A. Wilson Sayres, Qasim Ayub, et al. 2016. “Punctuated Bursts in Human Male Demography Inferred from 1,244 Worldwide Y-Chromosome Sequences.” *Nature Genetics* 48 (6): 593–99. https://doi.org/10.1038/ng.3559.
Pruitt, Kim D., Garth R. Brown, Susan M. Hiatt, Françoise Thibaud-Nissen, Alexander Astashyn, Olga Ermolaeva, Catherine M. Farrell, et al. 2014. “RefSeq: An Update on Mammalian Reference Sequences.” *Nucleic Acids Research* 42 (Database issue): D756-763. https://doi.org/10.1093/nar/gkt1114.
Quinlan, Aaron R., and Ira M. Hall. 2010. “BEDTools: A Flexible Suite of Utilities for Comparing Genomic Features.” *Bioinformatics* 26 (6): 841–42. https://doi.org/10.1093/bioinformatics/btq033.
R Core Team. 2018. “R: A Language and Environment for Statistical Computing. R Foundation for Statistical Computing, Vienna, Austria.” https://www.R-project.org/.
Ramírez, Fidel, Devon P Ryan, Björn Grüning, Vivek Bhardwaj, Fabian Kilpert, Andreas S Richter, Steffen Heyne, Friederike Dündar, and Thomas Manke. 2016. “DeepTools2: A next Generation Web Server for Deep-Sequencing Data Analysis.” *Nucleic Acids Research* 44 (W1): W160–65. https://doi.org/10.1093/nar/gkw257.
Ranallo-Benavidez, T. Rhyker, Kamil S. Jaron, and Michael C. Schatz. 2020. “GenomeScope 2.0 and Smudgeplot for Reference-Free Profiling of Polyploid Genomes.” *Nature Communications* 11 (1): 1432. https://doi.org/10.1038/s41467-020-14998-3.
Rautiainen, Mikko, and Tobias Marschall. 2020. “GraphAligner: Rapid and Versatile Sequence-to-Graph Alignment.” *Genome Biology* 21 (1): 253. https://doi.org/10.1186/s13059-020-

02157-2.
Rautiainen, Mikko, Sergey Nurk, Brian P. Walenz, Glennis A. Logsdon, David Porubsky, Arang Rhie, Evan E. Eichler, Adam M. Phillippy, and Sergey Koren. 2022. "Verkko: Telomere-to-Telomere Assembly of Diploid Chromosomes." *BioRxiv*, June, 2022.06.24.497523. https://doi.org/10.1101/2022.06.24.497523.
Rhie, Arang, Shane A. McCarthy, Olivier Fedrigo, Joana Damas, Giulio Formenti, Sergey Koren, Marcela Uliano-Silva, et al. 2021. "Towards Complete and Error-Free Genome Assemblies of All Vertebrate Species." *Nature* 592 (7856): 737–46. https://doi.org/10.1038/s41586-021-03451-0.
Rhie, Arang, Brian P. Walenz, Sergey Koren, and Adam M. Phillippy. 2020. "Merqury: Reference-Free Quality, Completeness, and Phasing Assessment for Genome Assemblies." *Genome Biology* 21 (1): 245. https://doi.org/10.1186/s13059-020-02134-9.
Rice, Peter, Ian Longden, and Alan Bleasby. 2000. "EMBOSS: The European Molecular Biology Open Software Suite." *Trends in Genetics* 16 (6): 276–77. https://doi.org/10.1016/S0168-9525(00)02024-2.
Robinson, James T., Helga Thorvaldsdóttir, Wendy Winckler, Mitchell Guttman, Eric S. Lander, Gad Getz, and Jill P. Mesirov. 2011. "Integrative Genomics Viewer." *Nature Biotechnology* 29 (1): 24–26. https://doi.org/10.1038/nbt.1754.
Rosenbloom, Kate R., Joel Armstrong, Galt P. Barber, Jonathan Casper, Hiram Clawson, Mark Diekhans, Timothy R. Dreszer, et al. 2015. "The UCSC Genome Browser Database: 2015 Update." *Nucleic Acids Research* 43 (D1): D670–81. https://doi.org/10.1093/nar/gku1177.
Sahakyan, Aleksandr B., Vicki S. Chambers, Giovanni Marsico, Tobias Santner, Marco Di Antonio, and Shankar Balasubramanian. 2017. "Machine Learning Model for Sequence-Driven DNA G-Quadruplex Formation." *Scientific Reports* 7 (1): 14535. https://doi.org/10.1038/s41598-017-14017-4.
Sahlin, Kristoffer, and Veli Mäkinen. 2021. "Accurate Spliced Alignment of Long RNA Sequencing Reads." *Bioinformatics* 37 (24): 4643–51. https://doi.org/10.1093/bioinformatics/btab540.
Sanders, Ashley D., Mark Hills, David Porubský, Victor Guryev, Ester Falconer, and Peter M. Lansdorp. 2016. "Characterizing Polymorphic Inversions in Human Genomes by Single-Cell Sequencing." *Genome Research* 26 (11): 1575–87. https://doi.org/10.1101/gr.201160.115.
Sanders, Ashley D., Sascha Meiers, Maryam Ghareghani, David Porubsky, Hyobin Jeong, M. Alexandra C. C. van Vliet, Tobias Rausch, et al. 2020. "Single-Cell Analysis of Structural Variations and Complex Rearrangements with Tri-Channel Processing." *Nature Biotechnology* 38 (3): 343–54. https://doi.org/10.1038/s41587-019-0366-x.
Schatz, Michael C., Anthony A. Philippakis, Enis Afgan, Eric Banks, Vincent J. Carey, Robert J. Carroll, Alessandro Culotti, et al. 2022. "Inverting the Model of Genomics Data Sharing with the NHGRI Genomic Data Science Analysis, Visualization, and Informatics Lab-Space." *Cell Genomics* 2 (1): 100085. https://doi.org/10.1016/j.xgen.2021.100085.
Sedlazeck, Fritz J., Philipp Rescheneder, Moritz Smolka, Han Fang, Maria Nattestad, Arndt von Haeseler, and Michael C. Schatz. 2018. "Accurate Detection of Complex Structural Variations Using Single Molecule Sequencing." *Nature Methods* 15 (6): 461–68. https://doi.org/10.1038/s41592-018-0001-7.
Seppey, Mathieu, Mosè Manni, and Evgeny M. Zdobnov. 2019. "BUSCO: Assessing Genome Assembly and Annotation Completeness." In *Gene Prediction: Methods and Protocols*, edited by Martin Kollmar, 227–45. Methods in Molecular Biology. New York, NY: Springer. https://doi.org/10.1007/978-1-4939-9173-0_14.
Shafin, Kishwar, Trevor Pesout, Pi-Chuan Chang, Maria Nattestad, Alexey Kolesnikov, Sidharth Goel, Gunjan Baid, et al. 2021. "Haplotype-Aware Variant Calling with PEPPER-Margin-

DeepVariant Enables High Accuracy in Nanopore Long-Reads." *Nature Methods* 18 (11): 1322–32. https://doi.org/10.1038/s41592-021-01299-w.
Shafin, Kishwar, Trevor Pesout, Ryan Lorig-Roach, Marina Haukness, Hugh E. Olsen, Colleen Bosworth, Joel Armstrong, et al. 2020. "Nanopore Sequencing and the Shasta Toolkit Enable Efficient de Novo Assembly of Eleven Human Genomes." *Nature Biotechnology* 38 (9): 1044–53. https://doi.org/10.1038/s41587-020-0503-6.
Shepelev, V. A., L. I. Uralsky, A. A. Alexandrov, Y. B. Yurov, E. I. Rogaev, and I. A. Alexandrov. 2015. "Annotation of Suprachromosomal Families Reveals Uncommon Types of Alpha Satellite Organization in Pericentromeric Regions of Hg38 Human Genome Assembly." *Genomics Data* 5 (September): 139–46. https://doi.org/10.1016/j.gdata.2015.05.035.
Sherry, Stephen T., Minghong Ward, and Karl Sirotkin. 1999. "DbSNP—Database for Single Nucleotide Polymorphisms and Other Classes of Minor Genetic Variation." *Genome Research* 9 (8): 677–79. https://doi.org/10.1101/gr.9.8.677.
Simpson, Jared T., Rachael E. Workman, P. C. Zuzarte, Matei David, L. J. Dursi, and Winston Timp. 2017. "Detecting DNA Cytosine Methylation Using Nanopore Sequencing." *Nature Methods* 14 (4): 407–10. https://doi.org/10.1038/nmeth.4184.
Skaletsky, Helen, Tomoko Kuroda-Kawaguchi, Patrick J. Minx, Holland S. Cordum, LaDeana Hillier, Laura G. Brown, Sjoerd Repping, et al. 2003. "The Male-Specific Region of the Human Y Chromosome Is a Mosaic of Discrete Sequence Classes." *Nature* 423 (6942): 825–37. https://doi.org/10.1038/nature01722.
Slater, Guy St C., and Ewan Birney. 2005. "Automated Generation of Heuristics for Biological Sequence Comparison." *BMC Bioinformatics* 6 (1): 31. https://doi.org/10.1186/1471-2105-6-31.
Šošić, Martin, and Mile Šikić. 2017. "Edlib: A C/C ++ Library for Fast, Exact Sequence Alignment Using Edit Distance." *Bioinformatics* 33 (9): 1394–95. https://doi.org/10.1093/bioinformatics/btw753.
Stamatakis, Alexandros. 2014. "RAxML Version 8: A Tool for Phylogenetic Analysis and Post-Analysis of Large Phylogenies." *Bioinformatics* 30 (9): 1312–13. https://doi.org/10.1093/bioinformatics/btu033.
Stanke, Mario, Mark Diekhans, Robert Baertsch, and David Haussler. 2008. "Using Native and Syntenically Mapped CDNA Alignments to Improve de Novo Gene Finding." *Bioinformatics* 24 (5): 637–44. https://doi.org/10.1093/bioinformatics/btn013.
Stephens, Zachary D., Matthew E. Hudson, Liudmila S. Mainzer, Morgan Taschuk, Matthew R. Weber, and Ravishankar K. Iyer. 2016. "Simulating Next-Generation Sequencing Datasets from Empirical Mutation and Sequencing Models." *PLOS ONE* 11 (11): e0167047. https://doi.org/10.1371/journal.pone.0167047.
Storer, Jessica, Robert Hubley, Jeb Rosen, Travis J. Wheeler, and Arian F. Smit. 2021. "The Dfam Community Resource of Transposable Element Families, Sequence Models, and Genome Annotations." *Mobile DNA* 12 (1): 2. https://doi.org/10.1186/s13100-020-00230-y.
Storer, Jessica M., Robert Hubley, Jeb Rosen, and Arian F. A. Smit. 2021. "Curation Guidelines for de Novo Generated Transposable Element Families." *Current Protocols* 1 (6): e154. https://doi.org/10.1002/cpz1.154.
Sun, C., H. Skaletsky, S. Rozen, J. Gromoll, E. Nieschlag, R. Oates, and D. C. Page. 2000. "Deletion of Azoospermia Factor a (AZFa) Region of Human Y Chromosome Caused by Recombination between HERV15 Proviruses." *Human Molecular Genetics* 9 (15): 2291–96. https://doi.org/10.1093/oxfordjournals.hmg.a018920.
Svetec Miklenić, Marina, Nikolina Gatalica, Angela Matanović, Bojan Žunar, Anamarija Štafa, Berislav Lisnić, and Ivan Krešimir Svetec. 2020. "Size-Dependent Antirecombinogenic Effect of Short Spacers on Palindrome Recombinogenicity." *DNA Repair* 90 (June): 102848. https://doi.org/10.1016/j.dnarep.2020.102848.

Szak, Suzanne T., Oxana K. Pickeral, Wojciech Makalowski, Mark S. Boguski, David Landsman, and Jef D. Boeke. 2002. "Molecular Archeology of L1 Insertions in the Human Genome." *Genome Biology* 3 (10): research0052.1. https://doi.org/10.1186/gb-2002-3-10-research0052.

Talenti, Andrea, and James Prendergast. 2021. "Nf-LO: A Scalable, Containerized Workflow for Genome-to-Genome Lift Over." *Genome Biology and Evolution* 13 (9): evab183. https://doi.org/10.1093/gbe/evab183.

Teitz, Levi S., Tatyana Pyntikova, Helen Skaletsky, and David C. Page. 2018. "Selection Has Countered High Mutability to Preserve the Ancestral Copy Number of Y Chromosome Amplicons in Diverse Human Lineages." *The American Journal of Human Genetics* 103 (2): 261–75. https://doi.org/10.1016/j.ajhg.2018.07.007.

Tseng, Bill, Hsiao-Lin Hwa, Chun-Yen Lin, Yu-Jen Yu, Adrian Linacre, and James Chun-I. Lee. 2022. "Y-SNP Haplogroup Hierarchy Finder: A Web Tool for Y-SNP Haplogroup Assignment." *Journal of Human Genetics* 67 (8): 487–93. https://doi.org/10.1038/s10038-022-01033-0.

Tubio, Jose M. C., Yilong Li, Young Seok Ju, Inigo Martincorena, Susanna L. Cooke, Marta Tojo, Gunes Gundem, et al. 2014. "Extensive Transduction of Nonrepetitive DNA Mediated by L1 Retrotransposition in Cancer Genomes." *Science* 345 (6196): 1251343. https://doi.org/10.1126/science.1251343.

Tyler-Smith, Chris, and William R. A. Brown. 1987. "Structure of the Major Block of Alphoid Satellite DNA on the Human Y Chromosome." *Journal of Molecular Biology* 195 (3): 457–70. https://doi.org/10.1016/0022-2836(87)90175-6.

Uralsky, L. I., V. A. Shepelev, A. A. Alexandrov, Y. B. Yurov, E. I. Rogaev, and I. A. Alexandrov. 2019. "Classification and Monomer-by-Monomer Annotation Dataset of Suprachromosomal Family 1 Alpha Satellite Higher-Order Repeats in Hg38 Human Genome Assembly." *Data in Brief* 24 (June): 103708. https://doi.org/10.1016/j.dib.2019.103708.

Van der Auwera GA and O'Connor BD. 2020. *Genomics in the Cloud: Using Docker, GATK, and WDL in Terra*. 1st ed. O'Reilly Media. https://www.oreilly.com/library/view/genomics-in-the/9781491975183/.

Varshney, Dhaval, Jochen Spiegel, Katherine Zyner, David Tannahill, and Shankar Balasubramanian. 2020. "The Regulation and Functions of DNA and RNA G-Quadruplexes." *Nature Reviews Molecular Cell Biology* 21 (8): 459–74. https://doi.org/10.1038/s41580-020-0236-x.

Vaser, Robert, Ivan Sović, Niranjan Nagarajan, and Mile Šikić. 2017. "Fast and Accurate de Novo Genome Assembly from Long Uncorrected Reads." *Genome Research* 27 (5): 737–46. https://doi.org/10.1101/gr.214270.116.

Vegesna, Rahulsimham, Marta Tomaszkiewicz, Paul Medvedev, and Kateryna D. Makova. 2019. "Dosage Regulation, and Variation in Gene Expression and Copy Number of Human Y Chromosome Ampliconic Genes." *PLOS Genetics* 15 (9): e1008369. https://doi.org/10.1371/journal.pgen.1008369.

Vlahović, Ines, Matko Glunčić, and Vladimir Paar. 2019. "Rich Polymorphic Variants of Alpha Satellite 34mer Higher Order Repeats in Hg38 Assembly of Human Chromosome Y." bioRxiv. https://doi.org/10.1101/768861.

Vollger, Mitchell R., Xavi Guitart, Philip C. Dishuck, Ludovica Mercuri, William T. Harvey, Ariel Gershman, Mark Diekhans, et al. 2022. "Segmental Duplications and Their Variation in a Complete Human Genome." *Science* 376 (6588): eabj6965. https://doi.org/10.1126/science.abj6965.

Vollger, Mitchell Robert. (2021) 2022. "SafFire. Last Accessed: 2022-11-29." July 18, 2022. https://github.com/mrvollger/SafFire.

Wang, Guliang, and Karen M. Vasquez. 2014. "Impact of Alternative DNA Structures on DNA

Damage, DNA Repair, and Genetic Instability." *DNA Repair*, Cutting-edge Perspectives in Genomic Maintenance, 19 (July): 143–51. https://doi.org/10.1016/j.dnarep.2014.03.017.

Webster, Timothy H, Madeline Couse, Bruno M Grande, Eric Karlins, Tanya N Phung, Phillip A Richmond, Whitney Whitford, and Melissa A Wilson. 2019. "Identifying, Understanding, and Correcting Technical Artifacts on the Sex Chromosomes in next-Generation Sequencing Data." *GigaScience* 8 (7): giz074. https://doi.org/10.1093/gigascience/giz074.

Welter, Danielle, Jacqueline MacArthur, Joannella Morales, Tony Burdett, Peggy Hall, Heather Junkins, Alan Klemm, et al. 2014. "The NHGRI GWAS Catalog, a Curated Resource of SNP-Trait Associations." *Nucleic Acids Research* 42 (D1): D1001–6. https://doi.org/10.1093/nar/gkt1229.

Wheeler, Travis J., and Sean R. Eddy. 2013. "Nhmmer: DNA Homology Search with Profile HMMs." *Bioinformatics* 29 (19): 2487–89. https://doi.org/10.1093/bioinformatics/btt403.

Wickham, Hadley, Mara Averick, Jennifer Bryan, Winston Chang, Lucy D'Agostino McGowan, Romain François, Garrett Grolemund, et al. 2019. "Welcome to the Tidyverse." *Journal of Open Source Software* 4 (43): 1686. https://doi.org/10.21105/joss.01686.

Wickham, Hadley, Danielle Navarro, and Thomas Lin Pedersen. n.d. *Ggplot2: Elegant Graphics for Data Analysis*. 3rd ed. Accessed November 28, 2022. https://ggplot2-book.org/.

Wickham, Hadley, Jim Hester, Romain Francois, Jennifer Bryan, Shelby Bearrows, RStudio, https://github com/mandreyel/ (mio library), Jukka Jylänki (grisu3 implementation), and Mikkel Jørgensen (grisu3 implementation). 2022. "Readr: Read Rectangular Text Data." https://CRAN.R-project.org/package=readr.

Xing, Jinchuan, Hui Wang, Victoria P. Belancio, Richard Cordaux, Prescott L. Deininger, and Mark A. Batzer. 2006. "Emergence of Primate Genes by Retrotransposon-Mediated Sequence Transduction." *Proceedings of the National Academy of Sciences* 103 (47): 17608–13. https://doi.org/10.1073/pnas.0603224103.

Zhao, Hao, Zhifu Sun, Jing Wang, Haojie Huang, Jean-Pierre Kocher, and Liguo Wang. 2014. "CrossMap: A Versatile Tool for Coordinate Conversion between Genome Assemblies." *Bioinformatics* 30 (7): 1006–7. https://doi.org/10.1093/bioinformatics/btt730.

Zook, Justin M., Brad Chapman, Jason Wang, David Mittelman, Oliver Hofmann, Winston Hide, and Marc Salit. 2014. "Integrating Human Sequence Data Sets Provides a Resource of Benchmark SNP and Indel Genotype Calls." *Nature Biotechnology* 32 (3): 246–51. https://doi.org/10.1038/nbt.2835.

Zook, Justin M., Nancy F. Hansen, Nathan D. Olson, Lesley Chapman, James C. Mullikin, Chunlin Xiao, Stephen Sherry, et al. 2020. "A Robust Benchmark for Detection of Germline Large Deletions and Insertions." *Nature Biotechnology* 38 (11): 1347–55. https://doi.org/10.1038/s41587-020-0538-8.

Zou, Xueqing, Sandro Morganella, Dominik Glodzik, Helen Davies, Yilin Li, Michael R. Stratton, and Serena Nik-Zainal. 2017. "Short Inverted Repeats Contribute to Localized Mutability in Human Somatic Cells." *Nucleic Acids Research* 45 (19): 11213–21. https://doi.org/10.1093/nar/gkx731.
